# Synaptic high-frequency jumping synchronises vision to high-speed behaviour

**DOI:** 10.1101/2025.08.20.671248

**Authors:** Neveen Mansour, Jouni Takalo, Joni Kemppainen, Alice D. Bridges, HaDi MaBouDi, Ali Asgar Bohra, Kaja Anielska, Vera Vasas, Théo Robert, Bruce Yi Bu, Shashwat Shukla, Yiyin Zhou, Maike Kittelmann, Joke Ouwendijk, Judith Mantell, Matthew Lawson, Gonzalo de Polavieja, Elizabeth Duke, Aurel A. Lazar, Paul Verkade, Lars Chittka, Mikko Juusola

## Abstract

During high-speed behaviour, animals must synchronise perception and action despite rapid environmental and self-generated motion. How neural systems achieve such precision remains unclear. Here we show how the housefly (*Musca domestica*) maintains visual accuracy during fast motion. Using intracellular and photomechanical recordings during saccade-like stimulation, we traced information flow from photoreceptors to large monopolar cells (LMCs). Visual neurons achieved record-high information sampling (∼2,500 bits·s^-1^) and synaptic transmission (∼4,100 bits·s^-1^), far exceeding previous estimates. We identify a previously unknown mechanism - *synaptic high-frequency jumping* - in which photoreceptor-LMC synapses dynamically shift transmission toward higher frequencies during saccades, extending visual bandwidth to ∼1,000 Hz, effectively eliminating synaptic delays, and quadrupling classical flicker-fusion limits (∼230 Hz). Behavioural experiments show flies respond synchronously within ∼13-20 ms, even before photoreceptor responses peak. A biophysically realistic model reveals how photomechanical-stochastic-refractory quantal sampling co-adapts with saccadic behaviour: through self-motion, flies efficiently translate image motion into temporally-precise, predictive high-speed vision.

## Introduction

Animals moving at high speeds must process visual information rapidly to avoid motion blur. How neural circuits achieve this - and whether self-motion helps or hinders perception - remains unclear. Answering this requires examining how innate and learned behaviours refine sensory perception to support survival. Brains, constrained by thermodynamics, genetics, and cellular biophysics, dynamically harness electrochemical, kinetic, and thermal energy to respond rapidly and accurately to internal and external signals^1^. Yet prevailing models often oversimplify neural signalling by treating neurons as static, unidirectional transmitters, neglecting the role of rapid physical movements at the ultrastructural level^2–8^. Such *morphodynamic interactions* - ultrafast, activitylzldependent mechanical and structural changes in neural elements - include photoreceptor microsaccades^1,2,9–15^, quantal neurotransmitter release^6,7,14–16^, and synaptic feedback mechanisms^17–20^. These phenomena collectively accelerate and enhance neural signalling, enabling rapid and precise perception and action^11,16^.

Houseflies exemplify aerial agility^21^, suggesting exceptional visual capabilities shaped by strong evolutionary pressures^22^. Historically, however, flies were presumed incapable of resolving fine visual details during rapid movements (**Fig. 1a-c**). Fast body and head movements (saccades) produce high angular velocities^21,23–26^, which, coupled with presumedly slow photoreceptor responses, were thought to blur vision^27,28^. Although compensatory head and thorax adjustments^25,26,29,30^ and specialised retinal zones^31,32^ partially mitigate this blur, rapid saccades were still assumed to momentarily render flies “blind”^33^. Yet this longstanding assumption contradicts flies’ remarkable ability to evade threats: how could flies buzzing around your head in summer, effortlessly dodging every swat, truly have impaired vision?

**Fig. 1.**
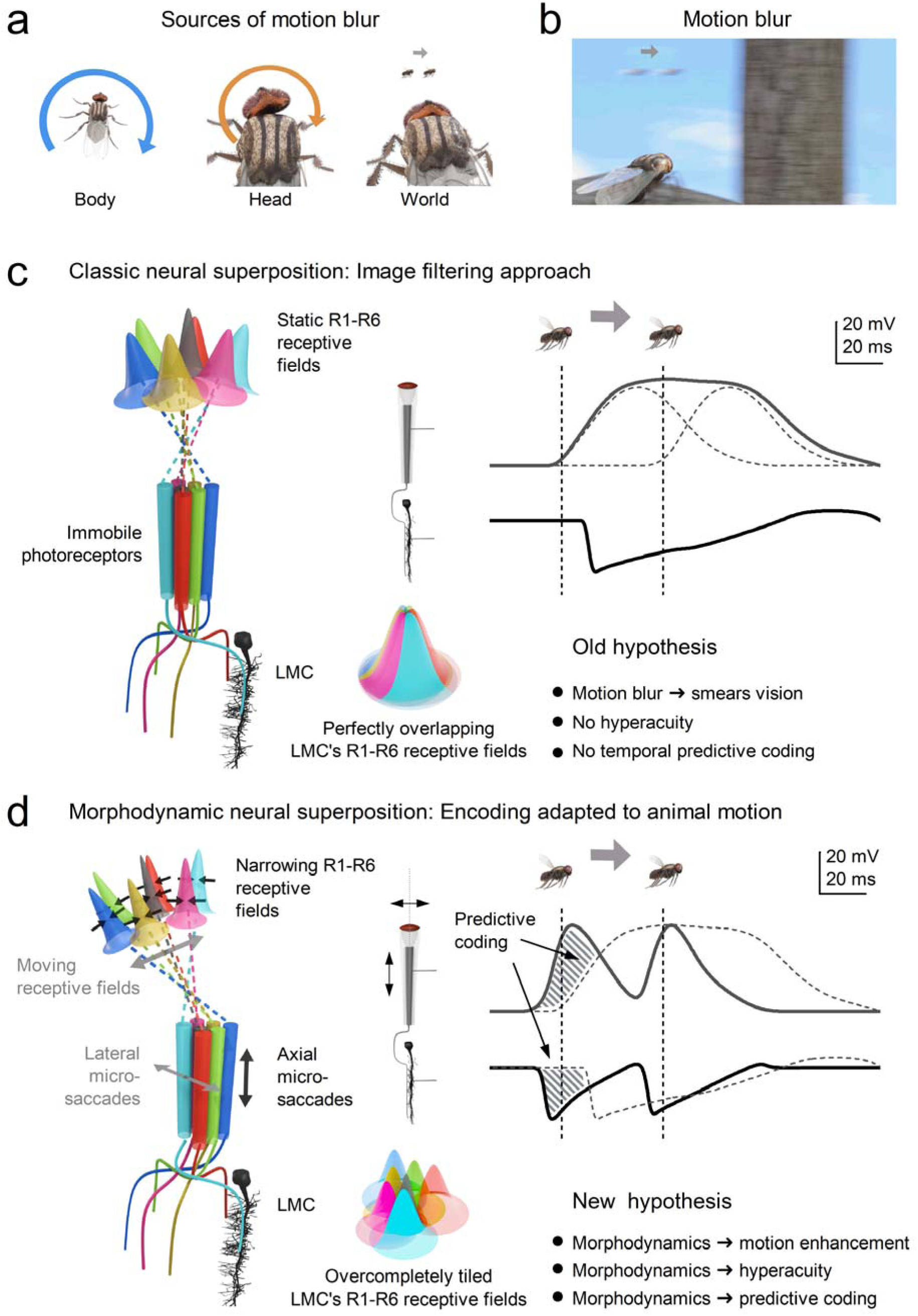
Comparing the predictions of classic static and morphodynamic models of neural superposition in fly vision. a, Sources of motion blur in natural vision arise from the animal’s own fast movements (rapid, saccadic body and head rotations) and from fast motion in the external world^27,28^. During flight or walking, these rapid turns and environmental motions produce abrupt, burst-like shifts of the retinal image, posing a fundamental challenge for neural encoding. **b,** Under classical assumptions, rapid image motion produces spatial and temporal smearing at the retinal level, degrading positional accuracy and limiting the ability to resolve fast-moving objects^27,28,33^. **c, Classic neural superposition model** (image-filtering hypothesis). In the traditional view^33–39^, R1-R6 photoreceptors within each ommatidium are assumed to be immobile and functionally identical, with fixed, static receptive fields that overlap only marginally in visual space. Large monopolar cells (LMCs) pool histaminergic^40–45^ (hyperpolarising) inputs from corresponding R1-R6 photoreceptors in neighbouring ommatidia, producing compound receptive fields that represent a single spatial location^37,46^. Illustratively, the LMC receptive field is depicted as a colour-sectored dome formed by identical R1-R6 receptive fields that perfectly overlap in the classic model. In this framework, neural encoding is modelled as a superposition of static spatial receptive fields convolved with relatively slow temporal impulse responses^27,28^ (typically estimated using white-noise analysis). Because receptive fields are comparatively broad and temporal integration is slow and noisy^39^, responses to moving stimuli are predicted to be strongly blurred in both space and time^33,47^. Consequently, nearby moving objects cannot be resolved independently, neural response peaks lag behind the true object position, and the model provides no intrinsic mechanism for hyperacute localisation or phase-advanced (predictive) encoding^27,28,47^. **d, Morphodynamic neural superposition model** (motion-adapted encoding hypothesis). In contrast to the classic view, the morphodynamic model incorporates the measured sizes and systematic misalignments of R1-R6 rhabdomere visual axes that underlie neural superposition^10,11,48^. Consequently, each R1-R6 photoreceptor has a slightly different receptive field in size and orientation, and together they tile visual space in an overcomplete manner^10,11,48^. Crucially, these photoreceptors are not static. During phototransduction, photoreceptors in neighbouring ommatidia undergo ultrafast axial and lateral photomechanical microsaccades^9–11^ that transiently shift and narrow their effective receptive fields. This motion dynamically reshapes spatial sampling. While receptive fields remain overcomplete at the level of LMCs, spatial pooling is no longer static: it becomes time-dependent and motion-coupled. Through stochastic, quantal, and refractory sampling, this morphodynamic reshaping reduces noise and sharpens spatial resolution^1,9^. For example, photomechanical microsaccades can shift and narrow receptive fields against object motion (dot - fly), causing neural responses to occur earlier^1,9,11^. In **c** and **d**, vertical dashed lines indicate when the fast-moving object (fly) is at the geometric centre of the photoreceptors’ receptive fields. The resulting responses enhance acuity, reduce effective motion blur, and generate phase-advanced temporal structure. Here, predictive coding (**d**, striped area) refers specifically to the biophysical time-locking of moving stimuli to their retinotopic neural representation. In this framework, predictive coding emerges directly from morphodynamic sampling and distinguishes LMC responses in the morphodynamic model (**d**) from those predicted by the classic static image-filtering model (**c**). Importantly, this form of predictive coding differs from hierarchical error-minimisation frameworks^49–51^ and from efference-copy or corollary-discharge mechanisms that cancel self-generated sensory input^52–54^. Instead, prediction here is intrinsic and biophysical: morphodynamic sampling and synaptic high-frequency dynamics reduce phase lag and preserve temporal precision during rapid self-motion.

To resolve this paradox, we examined how fly visual circuits respond to rapid self-motion by presenting controlled saccadic stimuli - light patterns that approximate the temporal statistics of natural saccades (**Fig. 1a**) - while isolating visual processing from the complexities of voluntary flight. We hypothesised that the fly visual system is not merely tolerant of saccades, but has evolved to exploit them, enabling exceptionally fast, accurate, and low-latency visual processing with minimal noise. Using a combination of intracellular recordings, photoreceptor microsaccade measurements, ultrastructural analyses, and biophysically realistic computational modelling, we explore whether visual information processing in the compound eye is not performed by static circuits constrained by optics or degraded by noise. Instead, it emerges from synergetically interacting, moving neural components that use motion itself to sample, encode, and predict visual inputs.

At the core of this process is the *morphodynamic neural superposition* architecture (**Fig. 1d**) of the compound eye^1,11,48^, formed by R1-R6 photoreceptors and large monopolar cells (LMCs). We show how the structure-function relationships within this circuit have co-evolved with saccadic behaviour to maximise coding efficiency during rapid image motion. By integrating empirical data with simulations of photoreceptor-LMC interactions within a novel theoretical framework - morphodynamic information processing^1^ - we identify and mechanistically explain a new phenomenon - *synaptic high*!zl*frequency jumping* - arising from coordinated morphodynamic sampling of rapid saccadic light changes, which enables hyperacute predictive vision.

## Results

### Multiscale Experimental Analysis

Experimentally, we assessed the signalling performance of housefly compound eyes - that is, how accurately, rapidly, and reliably they encode visual information - by studying both static and dynamic properties (**Fig. 2a**; **Supplementary Notes I-III**). To examine static structure, we used synchrotron X-ray imaging (**i**) and electron microscopy (**ii**) on fixed preparations to characterise the optical and ultrastructural adaptations. This included measuring the size and positioning of R1-R6 rhabdomeres across the eye and quantifying how many microvilli - photon-sampling units - each photoreceptor contains.

**Fig. 2.**
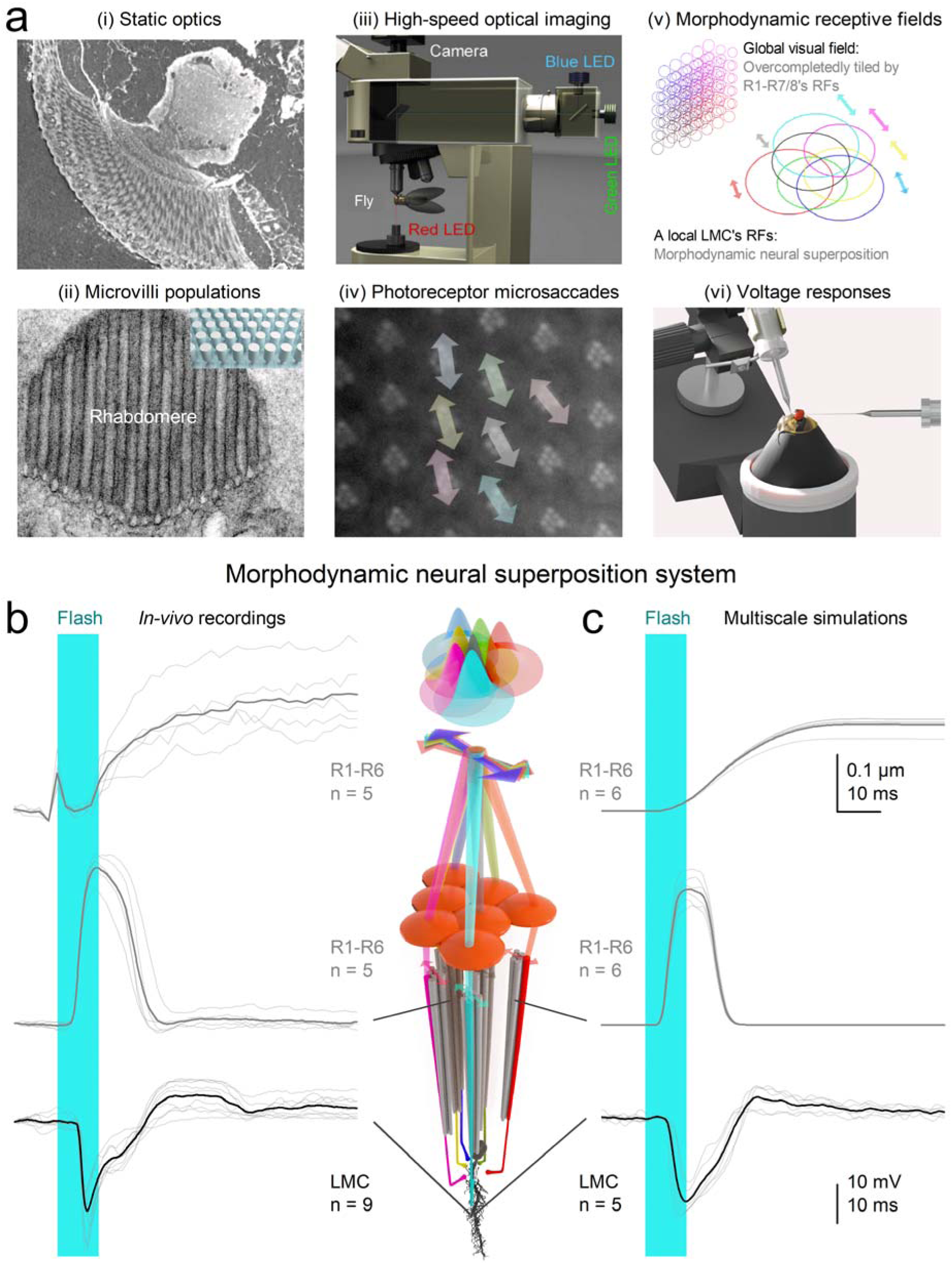
Investigating morphodynamic neural superposition system in the housefly compound eye. a, Key structural and functional components underlying the housefly’s morphodynamic vision were studied using both static imaging and live-recording approaches. The optical architecture of ommatidial structures was analysed in fixed eyes using synchrotron X-ray imaging. (i) The number of independent photon-sampling units (microvilli) in R1-R6 photoreceptor rhabdomeres was estimated via electron microscopy. (ii) High-speed infrared optical imaging enabled in vivo tracking of photoreceptor movements in response to light flashes. An infrared LED was used for movement monitoring without activating the cells, while green/blue light triggered photomechanical microsaccades. (iii) These recordings revealed photomechanical microsaccades - small, directionally diverse shifts of photoreceptors within ommatidia. (iv) Microsaccades shift photoreceptor receptive fields (RFs; circles - obtained through beam-tracing), which overlap to tile the compound eye’s visual field in an overcomplete pattern^11,48^ - i.e., overlapping but not perfectly aligned, unlike earlier assumptions^35,46^ (Fig. 1c, d). Each local LMC’s receptive field combines the photomechanically moving RFs of R1-R6 within a morphodynamic neural superposition system. (v) Intracellular voltage responses of photoreceptors and large monopolar cells (LMCs) to light stimuli were recorded using sharp microelectrodes. **b,** Representative experimental responses to a 10 ms UV flash (cyan bar): submicrometre photomechanical microsaccade measurements (top; including the flash-onset artefact), in vivo recordings of rapid voltage changes in photoreceptors (middle), and downstream LMCs (bottom). R1-R6 photoreceptors in neighbouring ommatidia sample the scene independently, producing variable phase-shifted voltage responses (spatiotemporally overcomplete sampling of the light stimulus) that converge onto a shared LMC, generating a morphodynamically shaped, transient voltage response (thin traces = individual responses, thick traces = mean responses). **c,** Simulated responses from a biophysically realistic model of the morphodynamic neural superposition system, in which over-completely tiled photomechanically moving photoreceptor receptive fields^11,48^, replicate the observed microsaccades and voltage responses. Informed by anatomical and physiological measurements in (**a**), the model accurately predicts receptive field movements and signal propagation through the photoreceptor-LMC circuit. The simulated R1-R6 responses are derived from the same ommatidium and incorporate rhabdomere size differences among these cells (that is, differences in microvillar number - photon sampling units). The simulated LMC responses likewise exhibit normal variability comparable to the real recordings. Responses are larger and faster, with more prominent off-responses, when driven by R1–R6 photoreceptors in the anteriofrontal region of the eye (**Supplementary Fig. 22**). There, increased retinal thickness corresponds to higher microvillar numbers (that is, more sampling units) in the rhabdomeres, enhancing the R1-R6 signal-to-noise ratio and accelerating their voltage responses^9,55^.

To assess dynamic properties, we investigated neural morphodynamics in intact living flies. Using high-speed infrared microscopy^9,11^ (**iii**), we recorded photomechanical microsaccades within neighbouring ommatidia (**iv**), and applied beam-propagation modelling^11^ (**v**) to estimate how these microsaccades move and narrow R1-R6 receptive fields locally. Finally, we used sharp microelectrodes (**vi**) inserted through a small corneal opening to record intracellular voltage responses of photoreceptors and LMCs to light stimuli.

### Biophysical Model of Morphodynamic Encoding

We used experimental findings to construct a biophysically-accurate multiscale model of the R1-R6 photoreceptor-LMC network (**Fig. 2c**; **Supplementary Notes IV**). Beneath each ommatidial lens, each modelled photoreceptor sampled changes in photon flux within its anatomically realistic receptive field via photomechanical movements^2,9,11^, rapidly contracting and elongating along its optical axis while shifting laterally in a complex piston-like motion. These microsaccades continuously reshaped and repositioned receptive fields in response to visual stimuli, depending on the size, eccentricity, and motion axis of each photoreceptor’s light-sensing structure, the rhabdome. Each R1-R6 rhabdomere contains a distinct number of microvilli, ranging from ∼41,000 to ∼74,000 depending on eye location (**Supplementary Notes II**), each functioning as an individual photon-sampling unit^56^. The model generated macroscopic R1-R6 photoreceptor responses from quantal photon absorptions and integrated these dynamic inputs through feedforward and feedback synapses to form a morphodynamic neural superposition system that continuously adapted the information flow between photoreceptors and LMCs to maximise visual encoding.

In the real eye, thousands of these morphodynamic neural superposition units tile the visual surface, forming overcomplete, localised encoding channels (**Figs. 2a**-iv,v, **2b**). Our model replicates how these units sample and process quantal visual information - i.e., changes in photon absorption rate - without requiring adjustable parameters. This framework links neural morphodynamics^1,9,11^ and active sampling dynamics^57–59^ to emergent coding strategies, such as network synchronisation, that enable high-speed vision and visually guided behaviour.

### Saccadic Bursts Amplify Neural Signalling

We performed electrophysiological experiments (**Fig. 3a**) using ‘saccadic’ light stimuli that mimic the rapidly changing intensity patterns experienced by photoreceptors during natural visual behaviours, such as rapid head and body rotations (Methods). These stimuli featured rapid contrast fluctuations, ranging from moderate (c ≈ 0.6) to high (c ≈ 1.5), across multiple temporal frequencies (20, 50, 100, 200, and 500 Hz 3 dB cut-offs; **Fig. 3c**). As a control and to benchmark our findings against classical studies^60–64^, we also applied lowlzlcontrast (cLJ≈LJ0.3), bandwidthlzllimited Gaussian white noise (GWN) stimuli (**Fig. 3c**). Here, we focus on diurnal conditions, when houseflies are most active. Additional experiments using these and further stimuli - extending to even higher 3 dB cut-off frequencies and spanning a million-fold intensity range from darkness to bright daylight - are described in **Supplementary Notes I** (**Supplementary Figs. 11-13**).

**Fig. 3.**
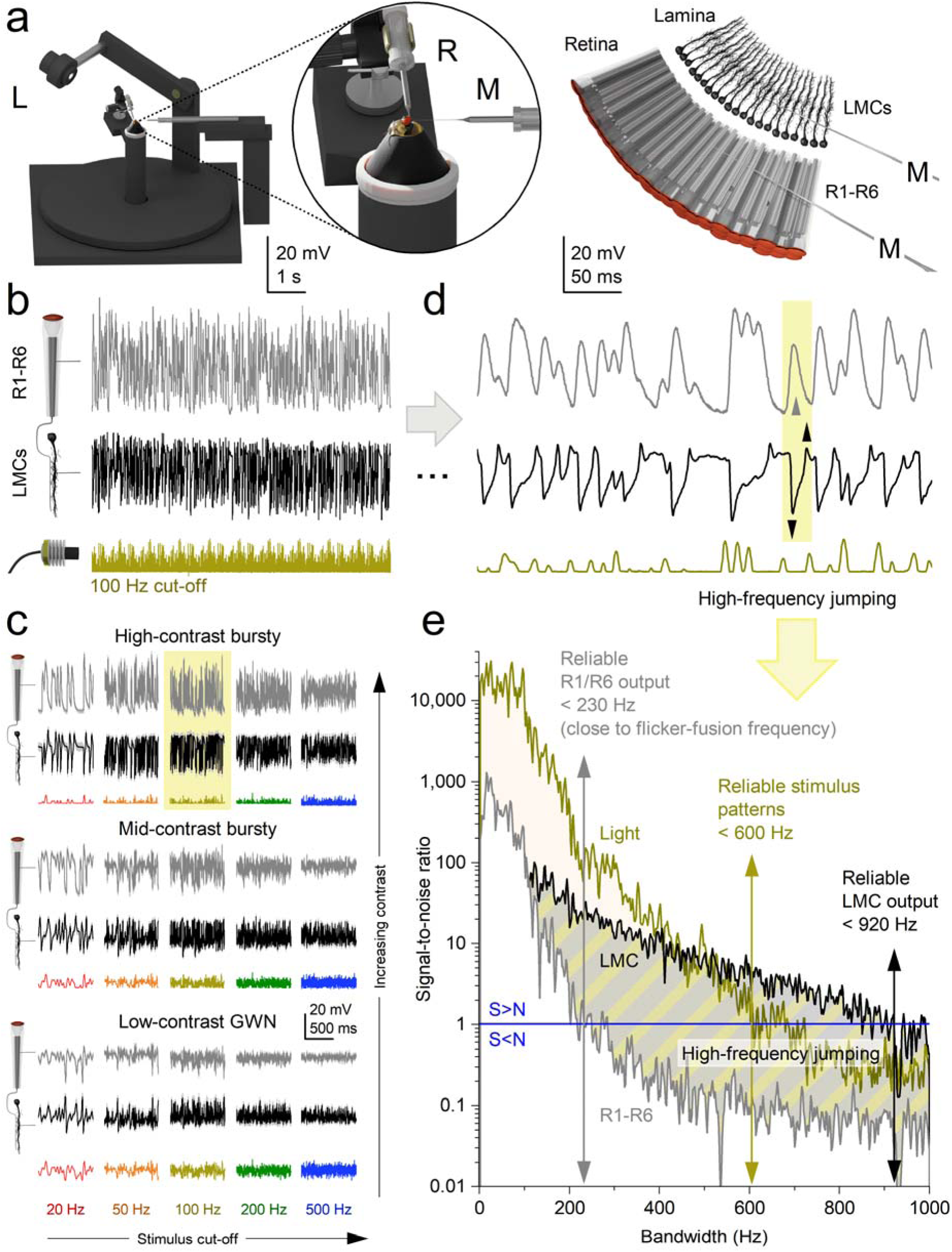
Electrophysiological recordings: *synaptic high-frequency jumping* transfers saccadic light information from photoreceptors to higher-frequency LMC responses. a, Left-schematic: The Cardan arm system positions light-point stimulus (L) at the centre of the recorded neuron’s receptive field, while micromanipulator-controlled measurement (M) and reference (R) microelectrodes record intracellular voltage responses. Right-schematic: R1-R6 photoreceptor and LMC recordings are conducted separately from the retina and lamina of intact, head-fixed flies (with intact eyes, red in the inset). Data from a single, stable photoreceptor and LMC with high signal-to-noise (see **Supplementary Figs. 2-13** for the corresponding population data and **Supplementary Tables** 1-6 for the corresponding statistics). **b,** Typical photoreceptor and LMC voltage responses to 30 repetitions of 2-second-long 100 Hz highcontrast bursts. **c,** Photoreceptor and LMC responses to varying stimuli, from saccade-like high-contrast bursts to the maximum-size Gaussian white-noise (GWN) stimuli (of low contrast) across specific bandwidths (20, 50, 100, 200 and 500 Hz). For details, see Methods (Fig. 10). Mean (thick traces) and individual responses (thin) to repeated stimuli (coloured). The yellow box highlights the most varying responses to 100 Hz highcontrast bursts. Recordings are from the same photoreceptor and LMC. **d,** A 600□ms segment of the recordings (from **b**) illustrates how an LMC generates ultrafast, inverted biphasic on/off responses (down-and up-arrows) to the slower, more monotonically rising and falling photoreceptor responses. While the photoreceptor outputs reflect low-frequency changes in light intensity, the LMC transforms this input into high-frequency transients, effectively *transposing* the slow signal into a faster carrier band. **e,** Signal-to-noise ratios of the bursty light stimulus and the resulting photoreceptor and LMC responses (from **b** and **d**) demonstrate how synaptic high-frequency jumping causes LMC responses to effectively double the base frequencies of the original light stimulus patterns. This results in LMC responses displaying an effective signalling bandwidth of approximately 920 Hz (signal-to-noise ratio > 1), compared to the photoreceptor responses and the light stimulus at approximately 230 and 600 Hz, respectively.

GWN stimuli, traditionally used for estimating neural information capacity^60–64^, are known to reduce encoding efficiency in neurons such as photoreceptors^9,55,65–67^ and LMCs^16^, which integrate quantal events (e.g., photon and neurotransmitter arrivals) through refractory sampling^9,55,65^. Each photoreceptor microvillus, a discrete photon-sampling unit, generates a quantum bump only after fully recovering from its previous phototransduction event^55,65,68^. Because GWN lacks the temporal structure of natural stimuli, it drives these sampling units into prolonged refractory states, impairing ability to track rapid photon-rate changes, and thereby reducing response amplitude and information content^9,55,65,69^.

Consistent with this, photoreceptors and LMCs exhibited their strongest responses to high-contrast saccadic bursts (**Fig. 3c**). These stimuli comprise brief, bright events separated by short, darker intervals, which allow photoreceptors to recover from refractoriness and integrate more photon quanta efficiently^9^. In contrast, responses to low-contrast GWN stimuli were significantly smaller and decreased further as stimulus frequency increased (bottom row, right column). The 500 Hz GWN stimulus was particularly ineffective, revealing a limit in how well these neurons can track rapidly changing, randomly ordered light contrast changes.

Together, our results from individual neurons and across populations (**Supplementary Figs. 2-13**; **Supplementary Tables 1**-**6**) show that housefly photoreceptors and LMCs preferentially encode fast, burst-like changes in light intensity, similar to those encountered during high-speed saccadic movements^9^. These findings highlight a fundamental limitation of neural coding under unnatural steady-state conditions (e.g., prolonged exposure to bright backgrounds with GWN stimulation), which artificially elevate refractoriness, reduce quantal integration efficiency, and suppress neural responses^55,66,67^.

### Synaptic High-Frequency Jumping Accelerates Vision

Because rapid saccadic flight behaviours are often thought to momentarily “blind” flies through motion blur^33^, we investigated how accurately the housefly photoreceptor-LMC superposition system encodes fast-changing visual inputs. **Fig. 3d** illustrates, with high temporal resolution, how typical R1-R6 photoreceptors and LMCs respond to a bursty sequence of saccadic light fluctuations. Both photoreceptor and LMC responses were nearly noise-free (**Supplementary Figs. 2b, 5b**), faithfully tracking repeated bursts of contrast. However, their voltage waveforms differed markedly.

Photoreceptors responded to stimulus intensity with relatively smooth, continuous signals. In contrast, LMC responses consisted of sharp, ultrafast transient signals, precisely aligned to the rising and falling phases of the photoreceptor response (**Fig. 3d**, yellow box with up/down arrows). These transients were temporally locked to contrast changes and effectively segmented the input signal into a string of biphasic events.

Comparing stimulus and response bandwidths revealed that LMC signals consistently shifted toward significantly higher frequencies, reliably encoding information up to ∼1,000 Hz (signal-to-noise ratio > 1; **Fig. 3e**; **Supplementary Fig. 11**). This *synaptic high-frequency jumping* far exceeded both the reliable stimulus bandwidth (∼600 Hz) and the photoreceptor’s encoding limit (∼230 Hz in this example). Thus, the biphasic nature of LMC responses (**Fig. 3d**, below) can quadruple or more the frequency content of their photoreceptor inputs (above), efficiently and instantaneously accentuating transitions in the photoreceptor signal to enhance temporal precision, enabling the synapse to resolve events with ∼0.5 ms precision (**Supplementary Fig. 11**; **Supplementary Movie 1**).

Thus, during rapid saccadic visual input, the photoreceptor-LMC circuit employs highlzlfrequency jumping to accelerate vision - shifting neural signals into higherlzlfrequency carrier bands where fast transients can be more effectively represented and transmitted - a strategy that mitigates motion blur and supports highlzlspeed, predictive control of behaviour.

### High-Frequency Jumping Maximises Neural Information

We next investigated how high-frequency jumping affects information transfer between R1-R6 photoreceptors (**Fig. 4a**) and LMCs (**Fig. 4b**) under diverse visual stimuli. To simulate realistic conditions encountered during rapid, cluttered flight, we used ten saccadic light patterns and five randomised Gaussian-white-noise (GWN) light patterns as controls (**Fig. 3c**).

**Fig. 4.**
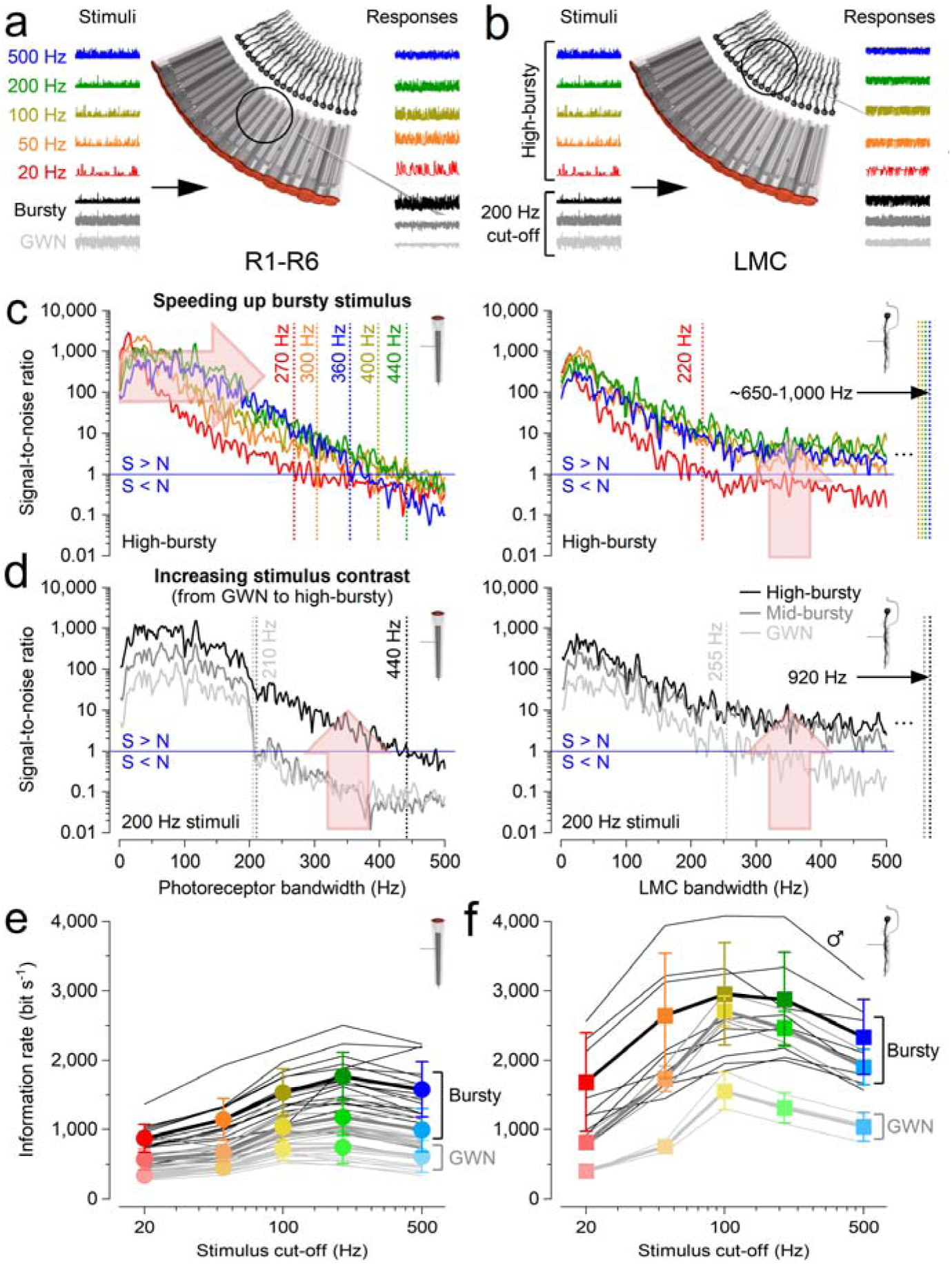
Electrophysiological recordings: high-frequency jumping maximises information transfer to LMCs during fast, high-contrast saccadic bursts. **a-d**, To ensure consistent depiction of stimulus-dependent adaptation trends, we present data from a single, highly stable photoreceptor and LMC recording series. **e-f**, Population data from the corresponding recording series of photoreceptors (n = 17) and LMCs (n = 5); cf. **Supplementary Figs. 2-13. a,** Analysis of R1-R6 photoreceptor signalling performance was based on intracellular voltage responses to repeated 2-s light patterns across 15 different stimuli. **b,** Similar analyses were conducted for LMCs to compare their responses to those of photoreceptors across all the stimuli. **c, Left**: in R1-R6s, higher saccadic stimulation cut-off (right arrow) expands effective signalling bandwidth up to ∼440 Hz (indicated by dotted lines, where the signal-to-noise ratio >1), increasing information content. **Right**: in LMCs, high-frequency jumping reallocate this enhanced information across their entire bandwidth (up arrow), elevating the signal-to-noise ratio and achieving effective signalling up to ∼1,000 Hz (excluding the slowest 20 Hz bursts, red trace). **d, Left**: Photoreceptors’ effective signalling bandwidth massively expands with stimulus contrast. For high-contrast saccadic bursts (c ≈ 1.29), the maximum signal-to-noise ratio reaches ∼2,000, and the bandwidth doubles to ∼440 Hz, compared to low-contrast Gaussian white noise (GWN; c ≈ 0.33) where the maximum signal-to-noise ratio is ∼100 with an effective bandwidth of ∼210 Hz. **Right**: LMCs’ effective signalling bandwidth also increases and broadens with stimulus contrast. High-frequency jumping is notably more effective during high-contrast bursts than with GWN. As a result, during saccadic stimuli, LMCs’ effective signalling range extends to over twice that of photoreceptors, reaching ∼920 Hz (for 200 Hz high-contrast bursts). Whereas with GWN, the LMCs’ effective signalling range is only slightly wider, ∼255 Hz, compared to the photoreceptors’ ∼210 Hz. **e,** Photoreceptors’ information transfer rates peaked for 200 Hz high-contrast “saccadic” (bursty) stimulation, with the highest estimates reaching about 2,500 bits·s^-1^. Their information transfer rates during GWN stimulation were 2-to-3-times lower. **f,** LMCs’ information transfer rates were 2-to-3-times higher than those of photoreceptors, reaching up to 4,000 bits·s^-1^ (in one male fly). These estimates typically peaked for 100 Hz or 200 Hz high-contrast “saccadic” bursts. The corresponding information transfer rates during GWN stimulation were 2-to-3 times lower. (**e**, **f**) Thin line, individual cells; thick, mean ± SD. The cell-to-cell variations in information transfer rate estimate likely reflect variable microelectrode recording locations and the eye’s sexual dimorphism.

Photoreceptor responses (**Fig. 4a**) showed that faster saccadic stimuli maximally broadened their effective signalling bandwidth (signal-to-noise ratio > 1) to ∼440 Hz, with this range being limited by each photoreceptor’s total number of microvilli (sampling-units)^9,55^. Variability across recordings (**Fig. 3e**) reflected natural differences in microvillar number, which depend on the rhabdomere length/thickness and vary across the compound eye (**Supplementary Figs. 21-23**). This bandwidth expansion substantially increased photoreceptor information content (**Fig. 4c**, left). Notably, these intracellular recordings typically exceeded the classic flicker-fusion frequency for *Musca* (∼230 Hz)^70^, which was originally derived from electroretinograms. Such extracellular field potential measurements underestimate local neural performance by averaging spatial and temporal signal variations, background activity, and noise across the eyes^18^.

LMC responses (**Fig. 4b**) exhibited even stronger phasic activity under the same conditions. Synaptic high-frequency jumping redistributed photoreceptor signals into higher-frequency carrier bands at the photoreceptor-LMC synapse, greatly extending LMC bandwidth (**Fig. 4c**, right). This effect was strongest during high-frequency, high-contrast bursts, where photoreceptor signal-to-noise ratios reached ∼2,000 (**Fig. 4c**, left). Under these conditions, LMC bandwidth (∼1,000 Hz) more than doubled that of the corresponding photoreceptors (∼440 Hz), whereas GWN stimuli (**Fig. 4d**) produced only modest increases (photoreceptors: ∼210 Hz; LMCs: ∼255 Hz).

The degree of LMC bandwidth extension depended on stimulus intensity, contrast, and cut-off frequency, with high-frequency jumping reaching maximal bandwidths during 50–200 Hz high-contrast bursts (∼920–1,000 Hz) at daylight intensities (**Supplementary Figs. 11-13**). For the LMC shown (**Fig. 4c**), all high-contrast bursty stimuli except the slowest (20 Hz) evoked high-frequency jumping, whereas low-contrast GWN never did. These results underscore GWN’s limitations in evaluating naturalistic neural coding^1,9,55,67^, particularly for fast, behaviourally relevant inputs.

With improved high-frequency signal-to-noise ratios, photoreceptor information transfer rates (**Fig. 4e**, left) peaked at ∼1,200-2,500 bits·s^-1^ during 200 Hz saccadic stimulation - compared to ∼600-1,000 bits·s^-1^ under GWN. Corresponding LMC rates (**Fig. 4e**, right) were 2-3 times higher, reaching ∼2,500-4,100 bits·s^-1^. These are likely the highest neural information rates reported to date and more than double those previously measured in *Calliphora* photoreceptors and LMCs under GWN^60^. Thus, the housefly’s morphodynamic neural superposition system appears explicitly tuned to encode fast saccadic inputs with exceptional efficiency and minimal noise, far surpassing conventional expectations^63,71,72^.

While our intracellular recordings clearly demonstrate the crucial role of synaptic high-frequency jumping in maximising information transfer during saccadic stimulation - mimicking information flow during high-speed behaviours - they cannot fully explain the underlying biophysical mechanisms. To address this, we systematically tested our multiscale model of the morphodynamic neural superposition system (**Figs. 2c**, **5**), directly comparing its predictions with experimental intracellular recordings and performance analyses.

**Fig. 5.**
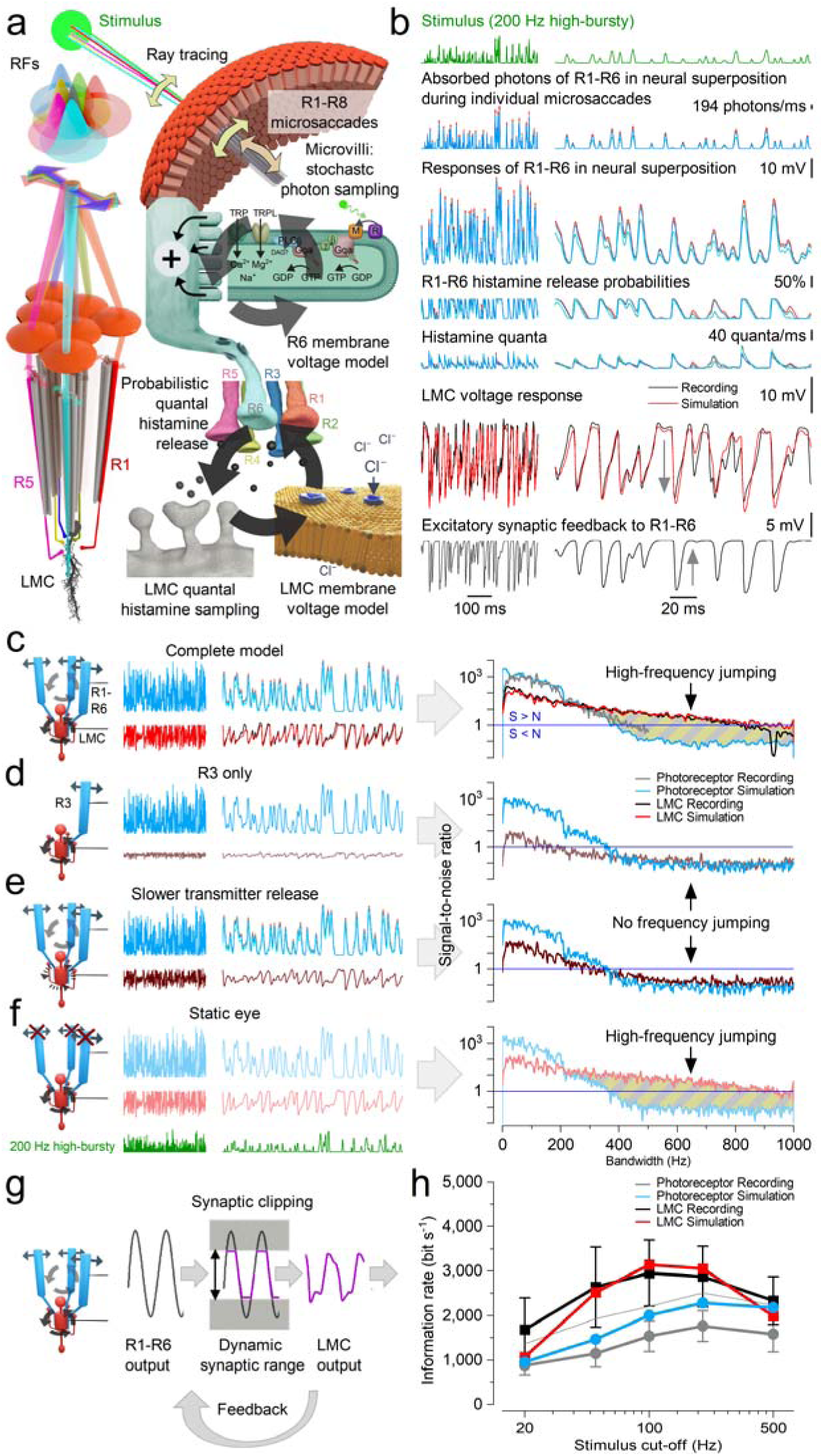
Morphodynamic neural superposition model: high-frequency jumping and hyperacuity emerge from parallel photoreceptor inputs shaped by multilayered interactions. a, Model architecture. R1-R6 photoreceptors from neighbouring ommatidia sample light through a “flower pattern” of partially overlapping receptive fields (RFs). **Feedback loop 1** (top): Individual microsaccades dynamically shift RFs in response to local contrast changes, driving stochastic quantum bump sequences and adapting the receptive field positions. **Feedback loop 2** (bottom): Voltage differences between R1-R6 photoreceptors trigger quantal histamine release, which binds to postsynaptic chloride channels on LMCs, generating hyperpolarising responses^40–45^. This engages excitatory synaptic feedback from LMCs to photoreceptors, balancing synaptic loads and enabling fast, phasic signal transmission^18,19^. **b, Biophysical signal flow during high-speed stimulation (200**LJ**Hz).** Top to bottom: photon absorption in R1-R6 during microsaccades; neural responses; histamine release probabilities and quantal output (6 individual traces in each case); resulting LMC voltage; and depolarising synaptic feedback^18–20^. Simulated responses (blue/red) closely match in vivo recordings (black/grey), demonstrating the model’s precision. c, Complete model reproduces high-frequency jumping. Signal-to-noise ratios (SNRs) of both recordings and simulations show high reliability across a broad frequency range under bursty stimulation, confirming the emergence of high-frequency jumping in LMCs. Note: L1/L2 LMC-subtypes receive “identical” histaminergic input from R1-R6^18,73,74^, and their On/Off polarities^75^ only emerge downstream in the medulla, likely shaped by their distinct neurotransmitters and local circuitry. d, Single input disrupts high-frequency jumping. Driving the LMC with input from only one photoreceptor (here R1) removes high-frequency jumping and reduces SNR. e, Slower transmitter release suppresses high-frequency jumping. Reducing synaptic speed lowers temporal resolution and disrupts high-frequency structure in LMC output. f, Phase locking persists without microsaccades. A static-eye model (no photomechanical input) still supports high-frequency jumping due to intrinsic morphodynamic (stochastic-quantal-refractory) sampling. **g, Excitatory Feedback from LMCs**^18,19^ **modulates photoreceptor responses, minimising the time signal transmission is clipped** by the limited synaptic output range - making the synapse differentiate more. h, Information rates match recordings. Simulated R1-R6 and LMC responses across stimulus bandwidths carry similar information to real recordings, peaking near 200 Hz. Mean and SD of the recordings in Fig. 4e,**f** (thin grey trace, R1-R6 recording with the highest rates). Note that the apparent difference between the stimulus frequencies at which photoreceptors and LMCs show maximal responses reflects a functional transformation rather than a mismatch. Photoreceptors respond most strongly to ∼200 Hz bursty stimulation because this frequency range maximally engages refractory stochastic quantal sampling under bright, saccade-like conditions, thereby sharpening temporal contrast and increasing signal-to-noise ratios. In contrast, following synaptic high-frequency jumping, LMCs exhibit maximal responses to burst envelopes around ∼100 Hz because the synapse redistributes power from these lower-frequency burst structures into much higher-frequency carrier components (extending toward ∼1,000 Hz). Thus, the stimulus frequency that maximally drives photoreceptors is not the same as the frequency band in which LMCs transmit information most efficiently.

### Multiscale Interactions Induce High-Frequency Jumping

The morphodynamic photoreceptor-LMC neural superposition model (**Fig. 5a-b**) accurately replicates experimental data and elucidates the mechanistic origins of high-frequency jumping (**Fig. 5c**; **Supplementary Figs. 27-38**). Simulations show that LMCs’ transient responses (**Fig. 3b**) - and thus high-frequency jumping - arise during the parallel tonic^16–18,76^ quantal histamine release from six photoreceptors (R1-R6) into a shared LMC^40–45^. Importantly, high-frequency jumping is not the result of a single mechanism but emerges from concurrent adaptive interactions between pre-and postsynaptic processes, as illustrated by two major circular feedback loops (**Fig. 5a**).

The model reflects the compound eye anatomy: R1-R6 photoreceptors from neighbouring ommatidia have partially overlapping receptive fields, forming a “flower pattern”^11,48^ (**Fig. 5a**). In the first feedback loop (top arrow circle), these receptive fields react to a spatiotemporal light stimulus (green disk) with microsaccades - tiny, directionally varied shifts driven by stochastic refractory photon sampling and photomechanical transduction in ∼41,000-74,000 microvilli per photoreceptor. This morphodynamic sampling generates diverse quantum bump sequences and voltage responses across R1-R6 (**Fig. 5b**, top three traces), dynamically tuning each photoreceptor’s receptive field to local stimulus changes.

In the second feedback loop (bottom arrow circle), differences in R1-R6 voltages modulate their tonic histamine release probabilities. Histamine binding to LMC receptors triggers Cl^-^ influx (left) and produces hyperpolarising postsynaptic responses^16,18,19,40–45^. The LMC thus integrates quantal input from six photoreceptors with partially overlapping receptive fields, boosting visual information flow to the brain^16,18,75,77^ (while suppressing aliasing^9,11^). Simultaneously, LMC signals provide excitatory feedback to photoreceptors, maintaining their tonic readiness. By continuously balancing excitatory and inhibitory loads^17–19,74,78^, this feedback ensures phasic and undelayed synaptic transmission.

To identify the components essential for high-frequency jumping, we systematically disabled or modified key mechanisms within the model. Simulations, consistent with the data processing theorem^66,79^, revealed that pooling signals from all six photoreceptors is critical (**Fig. 5d**). A single photoreceptor-LMC synapse, even under ideal noise-free conditions, transmits no more information than the photoreceptor itself. Tonic quantal release alone - e.g. in a single R1-LMC connection-introduces background noise, reducing the LMC’s signal-to-noise ratio and preventing high-frequency jumping.

In contrast, pooling slightly variable conductance changes from six photoreceptors (**Fig. 5b**), each with a signall⍰ltolzlnoise ratio >1,000 (**Fig.4**), naturally cancels this noise^9^. Pooling increases the collective photoreceptor output sixl⍰lfold, driving the synapse to progressively “clip” the extremes of these nearl⍰lnoiselzlfree bursts (**Supplementary Notes IV.11**). This clipping produces a squarelzllike waveform, injecting highlzlfrequency (>500⍰Hz) components that extend the LMC response bandwidth to nearly 1,000⍰Hz and enable in vivol⍰llike high-frequency jumping (**Fig. 5c**; **Supplementary Notes II.8**; **Supplementary Figs. 24-25**).

Another crucial requirement is rapid quantal transmitter release in photoreceptors (**Fig. 5e**). Slower release weakens modulation of histamine release probability, rendering LMC responses more low-pass and photoreceptor-like, reducing high-frequency signal-to-noise ratios and impairing high-frequency jumping. Moreover, removing excitatory feedback from lamina interneurons to photoreceptor terminals degrades both photoreceptor and LMC signal fidelity, consistent with experimental findings (**Fig. 5g**)^18,80^. However, synaptic high-frequency jumping still persists, albeit with altered adaptive properties (**Supplementary Fig. 33d**).

This feedback limits the time that large photoreceptor voltage responses - evoked by high-contrast saccadic bursts - spend outside the operational range of the histaminergic output synapse. When photoreceptors transiently hyperpolarise below this range, feedback phasically depolarises them; when voltages rise above it, the feedback is reduced or switched off^18,19^. Under daylight conditions, this feedback therefore acts as an additional high-pass mechanism, broadening synaptic bandwidth and improving both photoreceptor and LMC signal fidelity.

Interestingly, the model suggests that microsaccades contribute only indirectly to high-frequency jumping. Even in simulations with static (non-moving) photoreceptors, high-frequency jumping persists if synaptic feedback is preserved, though with altered dynamics (**Fig. 5f**). Microsaccades introduce variability due to asymmetric, stochastic and slightly asynchronous R1-R6 movements. In a neural superposition system, this results in small timing offsets - where one photoreceptor may activate before another. This temporal variability reduces the estimated information transfer rates of both photoreceptors and LMCs by ∼10%. For example, simulated LMCs transmit an average of 3,481 bits·s^-1^ with static photoreceptors, compared to 3,142 bits·s^-1^ with microsaccadic sampling (for 100 Hz high-contrast bursts: **Fig. 5h**).

### Photomechanical Interactions Accentuate Predictive Hyperacute Responses

We therefore asked whether this modest reduction in information transmission represents a small trade-off for enhanced spatiotemporal acuity.

The visual world is not a flat surface of stationary 1/f contrast patterns, but a complex three-dimensional environment in which objects occlude one another, *shaping the visual input animals experience as they move through space* (**Fig. 6a**). Visual systems must therefore encode dynamic spatial structure with sufficient acuity to guide rapid behaviour under self-motion.

**Fig. 6.**
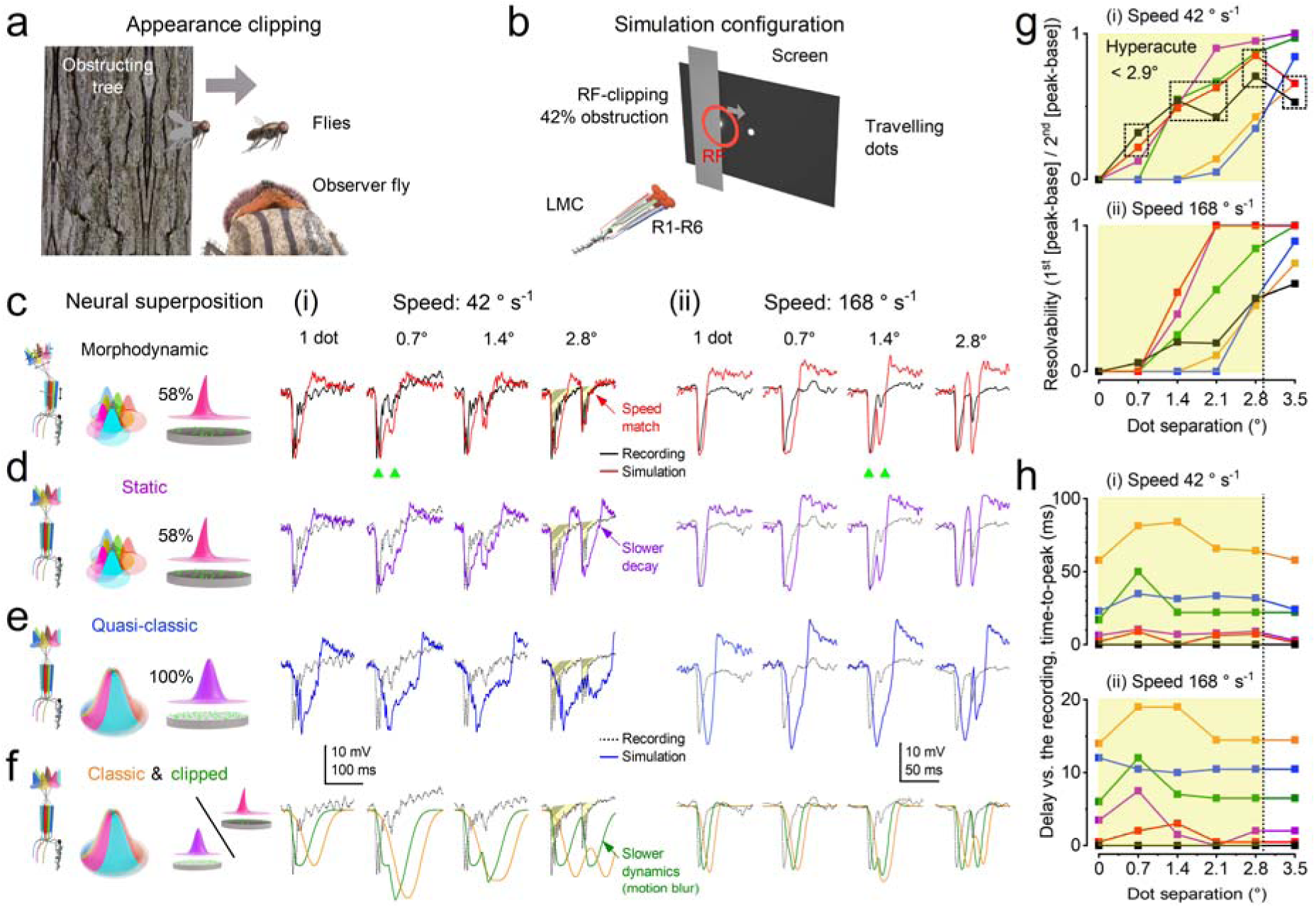
Electrophysiological recordings and simulations: predicted and recorded LMC responses to occlusion-revealed moving objects under different neural superposition models. We compared simulated outputs of large monopolar cells (LMCs) under progressively reduced neural superposition models with corresponding intracellular LMC recordings during controlled visual stimulation. In both recordings and simulations, the geometric centre of the LMC receptive field (RF) was positioned at the edge of a dark occluding plate that covered 42% of the RF (RF centre 0.8° from the plate edge). One or two small dots, mimicking sunlit flying insects, emerged from behind the occluding plate, reproducing the natural situation in which motion becomes visible after occlusion. **a**, Conceptual illustration of the behavioural scenario. **b**, Experimental and simulation configuration. Travelling dots moved laterally and appeared from behind an occluding screen, producing receptive-field clipping at emergence. LMCs integrate inputs from R1-R6 photoreceptors. **c-f**, Intracellular recordings and simulated LMC voltage responses under progressively reduced neural superposition models: **c**, Full morphodynamic neural superposition with 42% clipped receptive fields (red). Simulations closely match recordings, showing rapid, well-timed phasic rise and decay. **d**, Static neural superposition (no photoreceptor microsaccades) with overcomplete receptive fields, synaptic feedback, and stochastic refractory quantal sampling (purple). Responses decay more slowly than in recordings and the full model, indicating reduced spatiotemporal resolution. **e**, Quasi-classical neural superposition (no microsaccades), in which LMCs receive input from perfectly aligned photoreceptor receptive fields of identical size (classical assumption), while retaining stochastic, refractory quantal sampling and synaptic feedback (blue traces). The simulated responses are slower than in **c** and **d**. However, stochastic refractory quantal sampling together with synaptic feedback accentuates transient changes^9,11,17–19,55,68^, producing faster dynamics and reduced lag relative to the stimulus (i.e. less motion blur) compared with classic static-eye predictions (**f**). **f**, Classic stationary neural superposition filter model (orange and olive traces: full and 42% clipped RFs, respectively) following the assumptions of pioneering studies^27,33,35,37,39^. These simulations produce the slowest responses and poorest spatiotemporal resolution, consistent with static-eye predictions^27,28,33,39^, and - in the conventional full-RF case - exhibit little or no hyperacuity. Responses are shown for dots moving at 42° s⁻¹ (i) and 168° s⁻¹ (ii, saccadic velocity). For each speed, responses are shown for one dot and for two dots separated by 0.7°, 1.4°, and 2.1°. Coloured traces indicate model predictions; black (or dotted) traces show corresponding intracellular recordings. **g**, Resolvability (first peak/second peak ratio) as a function of dot separation for 42° s⁻¹ (top) and 168° s⁻¹ (bottom). Dotted boxes highlight the close match between recordings (black) and the full model (red). **h**, Delay (peak-to-peak) between simulated and recorded responses. Near-zero delays indicate accurate temporal prediction. The average interommatidial angle of the Musca compound eye is ∼2.9° (**Supplementary Table 10**), defining the static anatomical resolution limit. The morphodynamic model (**c**) resolves separations well below this limit, even at saccadic velocities^9,23^, demonstrating that motion-coupled morphodynamic sampling preserves hyperacute discrimination when objects emerge from occlusion. Among the tested models, only the morphodynamic model reproduces the ultra-brief, precisely timed LMC responses observed experimentally, indicating that photoreceptor microsaccades are essential for motion-blur reduction and predictive coding. **Supplementary Fig. 18** shows the corresponding case when dots disappear behind the occluding plate.

We hypothesised that cell-intrinsic photomechanical photoreceptor microsaccades^1,2,9–11^ - arising from asymmetric and rotated rhabdomere arrangements across neighbouring ommatidia - enhance compound-eye spatiotemporal acuity at the level of LMC output by transforming motion into structured temporal cues, thereby improving the resolution of moving objects^9,11^ (**Supplementary Figs. 14-18,24-25**). To test this, we presented *Musca* with moving dot stimuli at different speeds; complementary photoreceptor experiments using narrowing stripe stimuli are reported in **Supplementary Note I.7**. Here, we focus on LMC responses, which represent the output of the morphodynamic neural superposition system transmitted to the brain.

In these experiments, head-fixed *Musca* viewed two nearby dots moving at different speeds, up to saccadic velocities^9,23^ (>100°·s⁻¹), which abruptly appeared behind a screen (**Fig. 6b**). We compared how well the full morphodynamic neural superposition model (**Fig. 6c**) and progressively reduced variants (**Fig. 6d,e**), culminating in the classic stationary neural superposition model (**Fig. 6f**), reproduced the acuity observed in recorded LMC responses.

These experiments test the extent to which peripheral morphodynamic visual processing counteracts motion blur and exhibits predictive coding (**Fig. 1**). The full morphodynamic neural superposition model is identical to that used in **Fig. 5** - fixed and without free parameters - with only the test stimuli differing.

Across all tested conditions, intracellular recordings and the full morphodynamic neural superposition model exhibited ultra-brief response waveforms, minimal delays, and high spatiotemporal resolvability, outperforming all reduced variants. This performance is consistent with our hypothesis (**Fig. 1**) of morphodynamic, stochastic, refractory, quantal sampling and synaptic high-frequency jumping in the photoreceptor-LMC circuitry jointly enabling predictive coding and motion-blur minimisation.

These results demonstrate that during fast manoeuvres in cluttered natural environments with occluding objects, *Musca* can achieve hyperacute visual discrimination under conditions that would otherwise be severely degraded by motion blur. Mathematically, this **slit effect** - where an occluding object transiently narrows a photoreceptor’s receptive field and sharpens spatial resolution-resembles **synaptic clipping** (**Fig. 5g**), a temporal mechanism that boosts high-frequency signals (**Supplementary Fig. 25**).

At slower velocities (42°·s^-1^), information sampling and processing by the morphodynamic neural superposition system enhance the compound eye’s spatial acuity^9,11^ from the anatomical limit of ∼2.9° (the average interommatidial angle; **Supplementary Table 10**) to below 0.7°, representing a more than fourfold improvement (**Fig. 6**). Remarkably, even at saccadic speeds (168°·s^-1^), *Musca* LMCs retain hyperacute resolution, resolving details as fine as 1.4° (green triangles).

Together with independent photoreceptor experiments (**Supplementary Figs. 15-17**) showing that *Musca* photoreceptors can resolve moving edges separated by 0.9°, the conclusion is striking: the morphodynamic neural superposition system resolves moving visual details at or below the diffraction limit of the average *Musca* ommatidial lens (B= l. l°; **Supplementary Notes II.8**).

In summary, the second (synaptic) feedback loop in the morphodynamic neural superposition model plays the primary role in enabling temporal high-frequency jumping by coordinating phasic LMC responses to pooled inputs (**Supplementary Movie 1**). The first (photoreceptor) feedback loop, by contrast, primarily enhances spatial acuity, generating asymmetric “flower pattern” receptive field motion (**Supplementary Movie 2**) that sharpens object resolution, especially for moving objects (**Fig. 6**; **Supplementary Figs. 15-18**), and combats spatial aliasing^9,11^.

### Morphodynamics Matching Visual Lifestyles

In houseflies, as in *Drosophila*^9^ and honeybees^10^, photoreceptors contract photomechanically in response to changes in light intensity. However, in *Musca*, these movements - driven by refractory photon sampling reactions^1,2,9–11,56^ - occur much more rapidly, reducing saturation and more effectively maximising phasic information^9^ (**Fig. 7a-c**). This faster refractory quantal sampling improves the signal-to-noise ratio at higher stimulus frequencies (**Fig. 7e**). For example, *Musca* R1-R6 photoreceptors integrate voltage signals three to four times faster than those in *Drosophila*, as reflected in their effective signalling bandwidths: ∼308 Hz versus ∼72 Hz for the same 20 Hz bursty stimulus.

**Fig. 7.**
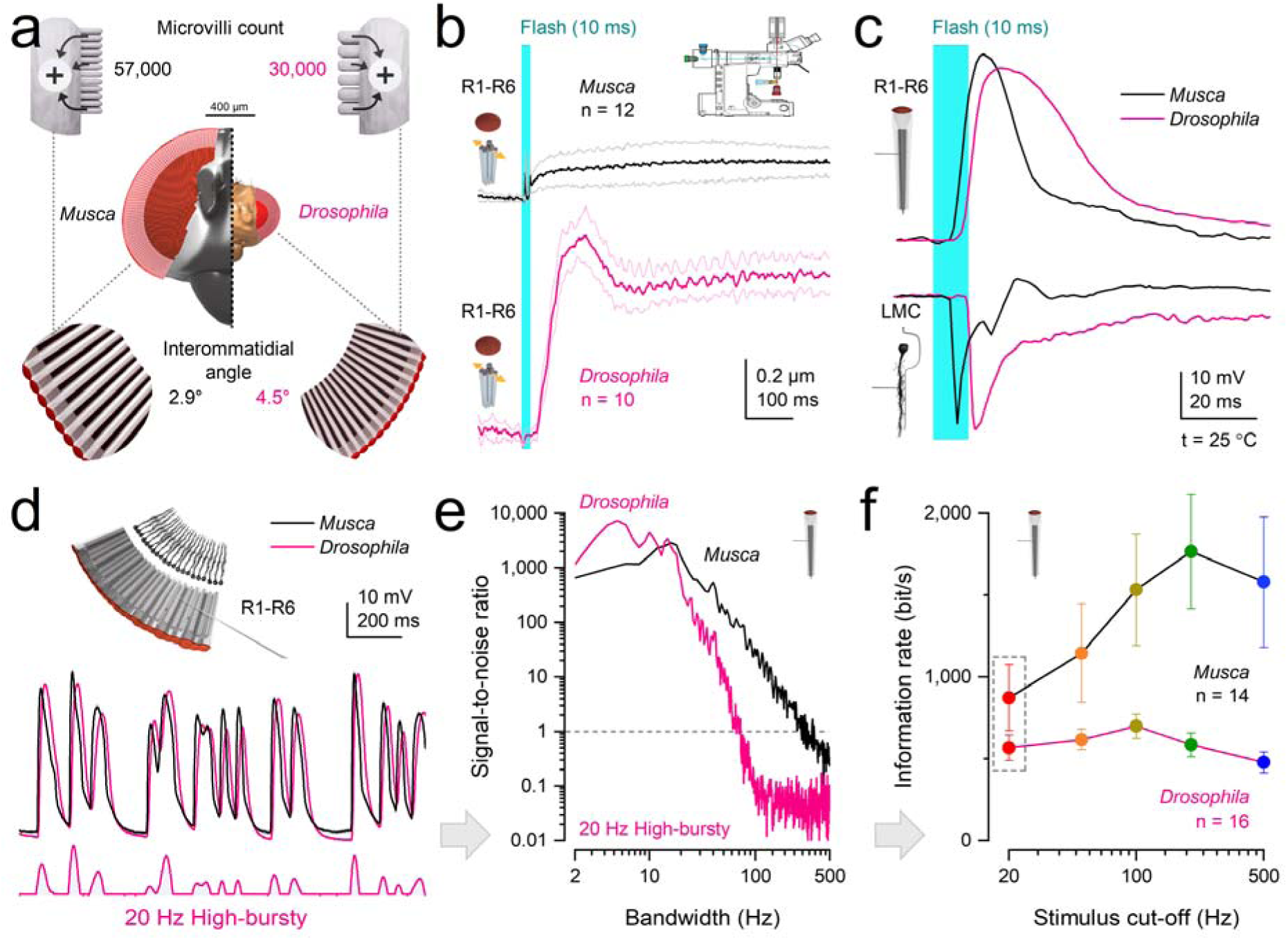
Electrophysiological recordings: morphodynamic sampling and synaptic transmission are adapted to species-specific visual behaviours. a, Anatomical specialisations. *Musca* photoreceptors contain ∼54,000 microvilli - nearly twice as many as *Drosophila* (∼30,000) - providing greater capacity for light sampling. The average interommatidial angle is also smaller in *Musca* (2.9° vs. 4.5°), enhancing spatial acuity. **b, Photomechanical microsaccades**. *Musca* R1-R6 photoreceptors exhibit smaller, faster microsaccades than those in *Drosophila*, enabling higher temporal resolution. **c, Ultrafast neural transmission**. In both species, LMC responses are biphasic and peak earlier than photoreceptor responses, with no detectable synaptic delay (**Supplementary Fig. 1**). In *Musca*, R1-R6 responses to a 10-ms flash reach their peak within 10-16 ms, aligning with the onset of visually guided behaviours (Fig. 8). Example recordings; population statistics in **Supplementary Fig. 1. d, Temporal structure of saccadic responses**. R1-R6 voltage responses to bursty (20 Hz) saccadic contrast changes show clear, phasic dynamics in both species, but *Musca* responses lead those of *Drosophila* in phase. **e, Signal-to-noise ratio (SNR)**. *Musca* photoreceptors maintain reliable signalling (SNR > 1) across a broader frequency range than *Drosophila*, supporting high-speed visual processing. **f, Information transfer**. *Musca* R1-R6 photoreceptors transmit up to three times more information than those in *Drosophila*, peaking at 200 Hz for high-contrast bursts - twice the optimal frequency observed in *Drosophila* (100 Hz). Dotted box highlights the information rates for the data in (**d** and **e**). **d**-**f**: Example recordings; population statistics in **Supplementary Figs. 2-3**. Corresponding *Drosophila* data and analyses from9.

Consequently, a typical R1-R6 photoreceptor in a fast-flying housefly samples approximately three times more information from the same stimulus than its slow-flying *Drosophila* counterpart, which has roughly half as many microvilli (∼30,000/photoreceptor) and exhibits slower refractory and quantum bump dynamics^9,55,61,62,68^. During high-contrast saccadic bursts, *Musca* photoreceptors reach maximal information rates of ∼2,510 bits·s^-1^ (**Fig. 4e**), compared to ∼850 bits·s^-1^ in *Drosophila*^9^ (**Fig. 7f**).

When six neural superposition photoreceptor outputs, each modulated by its own microsaccades, are combined through synaptic feedback, visual information is shifted into biphasic, aliasing-free, phase-locked LMC responses. These transiently amplify even the smallest changes in environmental contrast with minimal delay.

In both species, this rapid, bidirectional information flow eliminates classical synaptic delay: photoreceptor and LMC responses begin rising simultaneously (*Musca* ∼3.5LJms; *Drosophila* ∼6.5LJms after stimulus onset) (**Fig. 7c**). Yet LMC responses peak significantly earlier than their corresponding photoreceptor inputs - by ∼4LJms in *Musca* and ∼13LJms in *Drosophila* - consistent with predictive coding mechanisms operating at the synapse (**Supplementary Fig. 1a,b**).

In signal-processing terms, the photoreceptor-LMC synapse minimises phase lag relative to incoming motion trajectories, thereby improving forward encoding of dynamic input. This prediction is computational - phase-advanced and lag-minimised - rather than cognitive or expectation-based. *Musca*’s faster LMC responses (**Fig. 7a-c**) align with its quicker behavioural reactions and the need to time-lock visual processing to rapid motor outputs.

These findings suggest that evolution has tuned visual information processing to meet each species’ behavioural demands - adjusting microvillus numbers, refractory periods, quantum bump dynamics, photomechanical responses, membrane conductances, and synaptic connectivity - while also constraining metabolic cost^9,55,65^ (**Supplementary Notes IV**). As a result, housefly photoreceptors and LMCs jointly encode contrast changes at roughly twice the speed of those in *Drosophila*^9^ (**Fig. 7a-c**).

### Predictive Coding Enables Ultrafast Behaviours

Next, we investigated whether ultrafast morphodynamic processing, combined with high-frequency-jumping-induced acceleration of neural signalling, is also reflected in the speed of *Musca*’s visually triggered behaviours (**Fig. 8**). To test this, we employed two behavioural paradigms that yielded similar results. First, binocular light flashes elicited rapid antennal movements (**Fig. 8a-d**), potentially allowing the fly to gather additional olfactory, auditory, or thermal information to reduce stimulus-related uncertainty. Second, when startled by a rapidly looming dark object, a tethered *Musca* exhibited a synchronised, rapid lift of all six legs (**Fig. 8e-h**; **Supplementary Movie 3**).

**Fig. 8.**
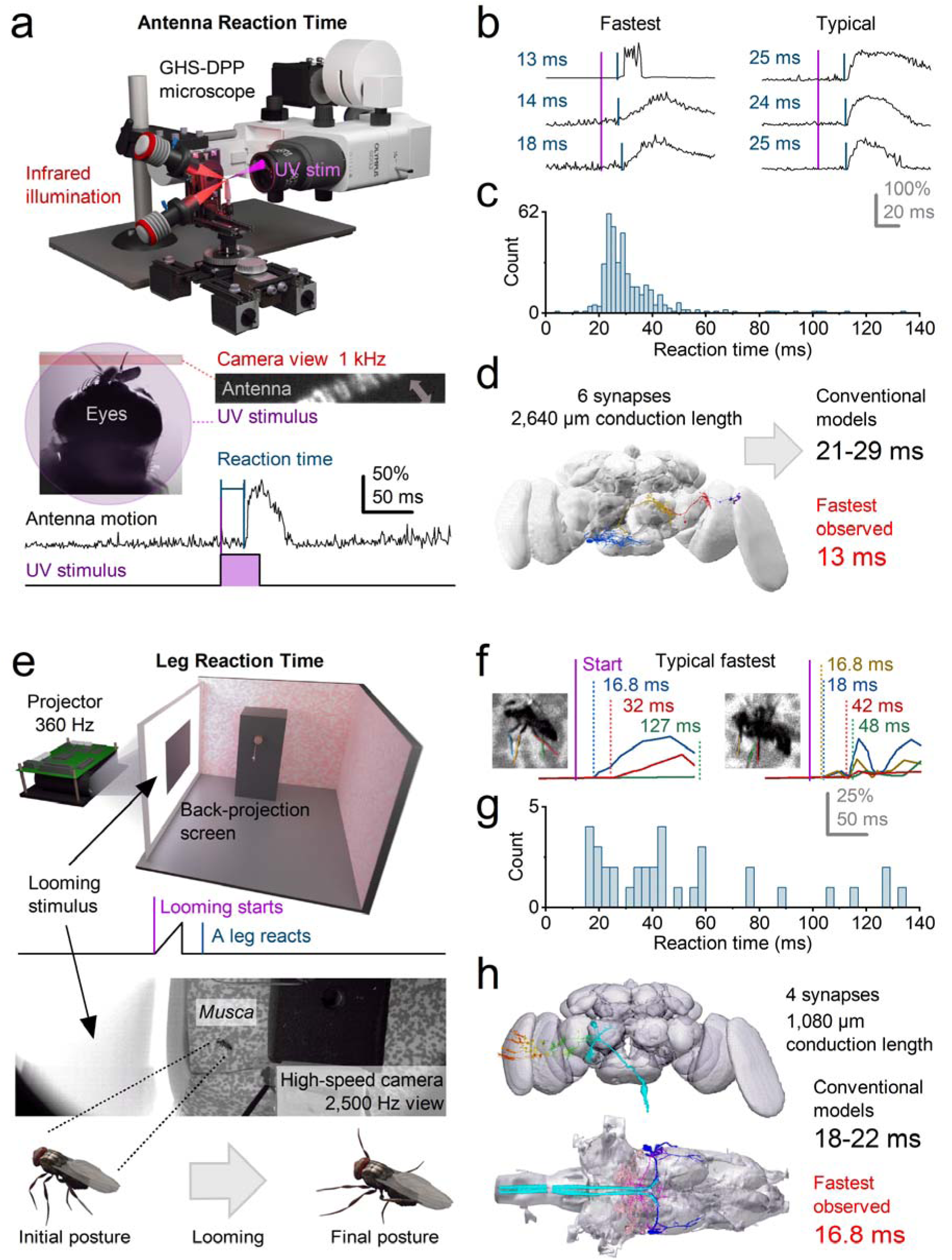
Voluntary behavioural responses in flies outpace classical conduction-delay predictions, even along the shortest neural pathways - revealing accelerated sensorimotor processing. a, High-speed infrared videography (1□kHz) was used to track *Musca* antennae responses to a brief UV light flash. Antennal motion was recorded relative to stimulus onset to measure reaction times. **b,** Cross-correlation of antennal motion traces revealed that the shortest reaction latencies occurred within 13-18□ms (left), while more typical responses peaked around 24-25□ms (right), indicating variability across trials. **c,** The broad distribution of reaction times suggests that antennal responses are not reflexive but instead voluntarily modulated. Some flies responded consistently, others sporadically, and some not at all. **Supplementary Notes V** (**Supplementary Fig. 41**) present the corroborating statistical analysis. **d,** A candidate antennal response circuit comprising 5 synapses was reconstructed using the *Drosophila* connectome and scaled to *Musca* using X-ray microtomography. Standard estimates predict a minimal unidirectional pathway latency of ∼21-29□ms. This is 8-16□ms (up to two times) slower than the fastest observed 13□ms antennal responses, implying the involvement of in vivo acceleration mechanisms (e.g., synaptic high-frequency jumping). **e,** In a separate setup, tethered flies were presented with high-speed looming stimuli via a 360□Hz projector and back-projection screen, while leg-lift responses were recorded using a 2.5□kHz high-speed camera (see **Supplementary Fig. 39**). **f,** Reaction times to looming stimuli varied substantially across individuals and trials. The fastest responses were detected after 16.8-18□ms, with others occurring at 32, 42, or even >100□ms poststimulus, suggesting a mixture of voluntary and non-responses. **g,** Distribution of leg reaction times reveals a multimodal pattern, again consistent with voluntary rather than reflexive control. Supplementary Notes V (**Supplementary Fig. 40**) present the corroborating statistical analysis. **h,** The leg-lift pathway includes at least 4 synapses and spans ∼1.1□mm in conduction length to giant fibres (GFs). Using standard assumptions (Methods), the fastest expected motor response would take ∼18-22□ms. Yet observed leg responses at 16.8□ms are 1.2-5.2 ms faster than predicted, reinforcing the idea that classical serial conduction models underestimate the true speed of visual processing in active fly behaviour.

Strikingly, the shortest response latencies ranged from 13 to 16.8LJms. These exceptionally brief delays are remarkable because the behaviours appear voluntary and decision-based, conditions typically associated with greater response variability and a higher likelihood of no response^1,81^. In both cases, neural signals must first be generated in the photoreceptors and then transmitted through brain circuits (**Supplementary Movie 4**). Even the most direct “reflex-like” pathway - bypassing central decision-making - would involve at least five synapses (**Supplementary Notes VI**) before reaching the muscles. Yet both responses were highly variable in timing and often absent, distinguishing them from classic, involuntary reflexes^82^.

To estimate the minimal possible reaction time, we extrapolated from the complete *Drosophila* brain connectome, adjusting for *Musca*’s brain being approximately three times larger (**Supplementary Notes VI**). Using standard values for synaptic transmission, neural charging, and conduction delays within a unidirectional signalling model (**Fig. 8h**; Methods), we predicted a minimum reaction time of 18-29LJms for reflex-like pathways, containing four to six synapses. These predicted delays are 5-16LJms (38%-123%) longer than the shortest observed voluntary response latencies.

This discrepancy suggests that *Musca*’s neural processing is not strictly feedforward, but operates within recurrent feedforward–feedback loops across brain circuits^17,18,81^. The reduction in phase lag arises locally from synaptic and morphodynamic mechanisms. However, within this broader predictive framework, the fly’s internal state - including behavioural context and arousal - can modulate sensory gain and temporal coordination^1,83–85^. We propose that this global recurrent architecture supports the binding of object features across space and time, enabling predictive, phasic information flow powered by quantal, refractory, and morphodynamic mechanisms - mirroring those observed at the photoreceptor-LMC synapse (**Fig. 5**).

## Discussion

Integrating multiscale experimental and modelling approaches, we uncover *synaptic high-frequency jumping* and explain how it emerges. This previously undescribed mechanism enables peripheral visual neurons to shift information into higher carrier frequencies in response to high-speed saccadic input, thereby minimising communication delays and increasing the coding speed of reliable vision. Remarkably, housefly LMCs can transmit information at rates exceeding 4,000LJbits·s^-1^ and operate at bandwidths approaching 1 kHz - far beyond classical flicker-fusion limits^70^. Using ultrahigh-speed videography, we further show that houseflies initiate voluntary, stimulus-triggered behaviours at a time when photoreceptor responses are only just reaching their peak.

These findings challenge long-standing models of sequential neural transmission and reveal how vision dynamically adapts to behavioural demands. Rather than passively processing visual information, houseflies actively shape their sensory input through high-speed flight behaviours, generating the spatiotemporal structures that drive high-frequency jumping, predictive coding, hyperacute perception and rapid neural synchronisation with environmental dynamics.

By highlighting the critical roles of saccadic visual behaviours, morphodynamic mechanisms, and bidirectional synaptic interactions in enabling fast, parallel, low-latency information sampling and processing, our results have broad implications for understanding efficient encoding and predictive coding. In particular, synaptic high-frequency jumping suggests a neurophysiological solution to the binding problem - that is, how information encoded across distinct brain circuits is synchronised to produce unified perception, decision, and action - within the physical constraints of neural computation.

*How does active, high-speed self-motion enhance visual acuity and information throughput?* Houseflies maintain superior visual performance during rapid saccadic turns generated by flight manoeuvres. These self-induced movements do not impair vision; rather, they enhance it. Morphodynamic neural superposition enables photoreceptors and downstream circuits to extract temporally structured, behaviourally relevant features with minimal delay, enabling simultaneous efficient processing of high-speed visual information and hyperacute perception. The resulting phasic signals are rapidly amplified and undergo synaptic high-frequency jumping, emerging as transient, biphasic LMC responses that broaden bandwidth and shorten latency. These signals are further accelerated by the brain’s bidirectional information flow, where tonic feedforward (inhibitory) and feedback (excitatory) interactions help balance synaptic load^1,17–19,78^. As a result, they become synchronised with internal motor states, generating a predictive, time-locked encoding of environmental dynamics - facilitating high-speed decision-making and stable visual perception, even under variable lighting conditions.

Information throughput increases with the number of samples, assuming constant conditions^1,65,66,79^. Photoreceptors continuously adapt to fluctuating light intensities - reflecting logarithmic changes in environmental photon rates - through *refractory quantal sampling*, which dynamically adjusts the quantum efficiency of microvilli^1,9,55,68^. This mechanism enables signal-to-noise ratios of ∼1,000-8,000 (**Fig. 7e**) in response to fast saccadic contrast changes under bright daylight conditions. Approximately 54,000 microvilli per cell engage in stochastic sampling, effectively absorbing ∼10LJ-10LJ photons per second. In bright illumination (>10^7^ photons/s), microvillar refractoriness prevents most absorbed photons from eliciting a quantal response^1,9,55,65,68^. Under these conditions, photoreceptor microsaccades - together with the fly’s saccadic flight behaviours - selectively enhance the sampling of contrast changes^1,9,11^. As a result, photoreceptors produce accurate anti-aliased estimates of the dynamic visual scene without saturating their biophysically limited amplitude or frequency ranges, while averaging out residual noise^1,9,55,68^. Their histaminergic synapses can thus respond reliably and efficiently to even the slightest contrast changes.

Each LMC integrates slightly variable inputs from six photoreceptors, or seven in the male’s frontal-dorsal “love spot” region^70^, via overlapping, photomechanically moving receptive fields (**Fig. 9**). This morphodynamic pooling of parallel photoreceptor outputs enhances spatiotemporal resolution, enabling hyperacute pattern detection and object tracking well below the ∼2.9° limit imposed by the average interommatidial angle, which defines the static pixel resolution of the compound eye. Remarkably, our recordings and simulations show that the morphodynamic neural superposition system, evident in LMC voltage responses, can resolve moving objects separated by just 0.7° - narrower than the ommatidial lens’s airylzldisk angle (1.1°), the theoretical diffraction limit (**Fig. 6**, **Supplementary Fig. 24**) - a performance once thought impossible^33^.

**Fig. 9.**
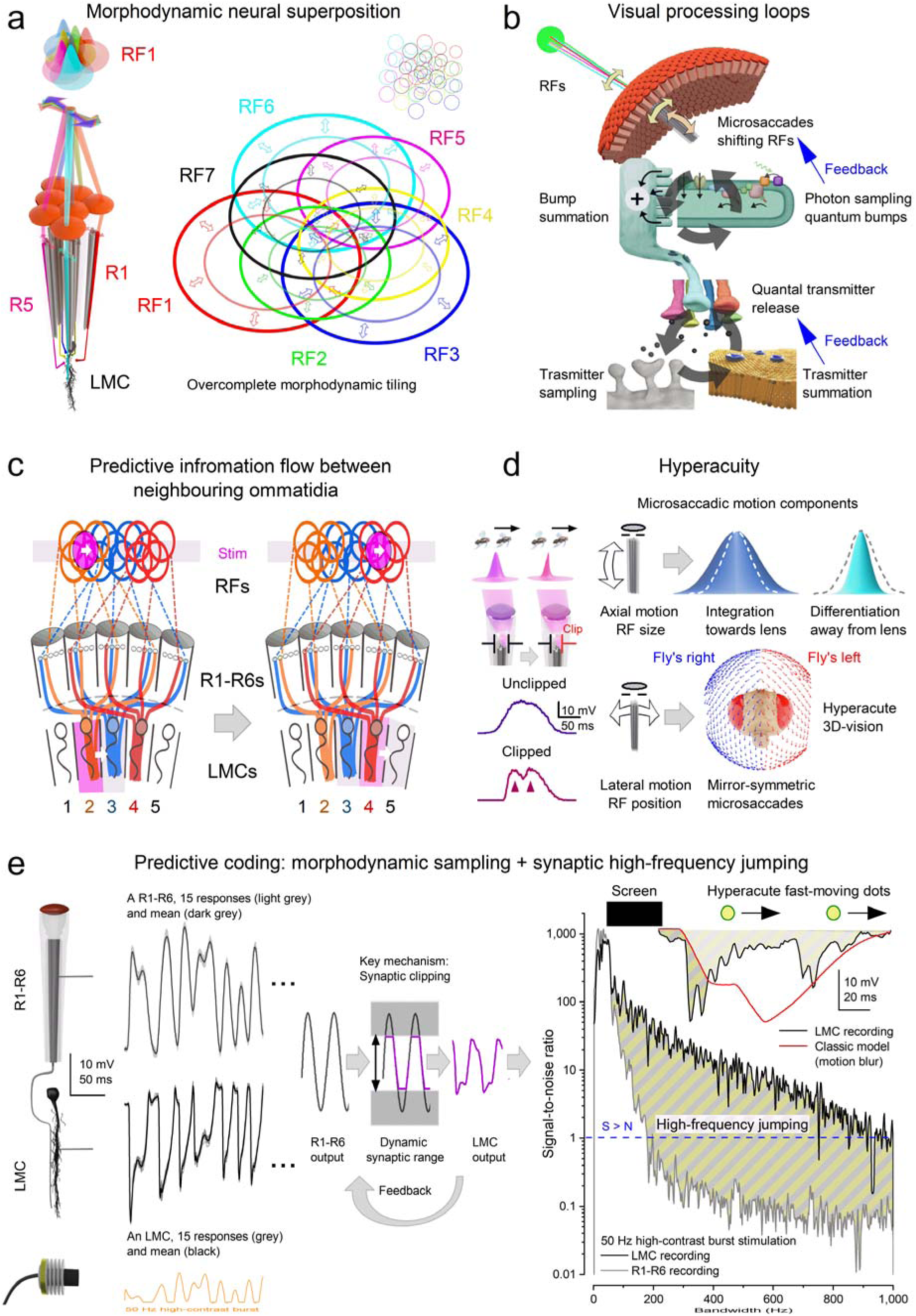
A biophysically realistic morphodynamic neural superposition model of a fly’s peripheral vision explains hyperacute predictive coding. **a,** Overcomplete morphodynamic neural superposition wiring. R1-R7/8 photoreceptors from neighbouring ommatidia sample light through a “flower-like” arrangement of partially overlapping receptive fields (RFs), collectively feeding information to a single LMC^11,48^. Each LMC’s receptive field therefore represents the combined morphodynamic sampling matrix of these inputs, centred on the R7 photoreceptor’s receptive field (RF7). It shifts and narrows morphodynamically in response to the spatiotemporal light patterns detected by the corresponding photomechanical R1-R7/8 receptive fields (RF1-7/8)^1,9–11^. See photoreceptor microsaccade components in (**d**). **b,** Photomechanical and synaptic feedback loops sharpen fly vision. Photomechanical feedback (top): Microsaccades generated by light-activated microvilli dynamically shift RFs in response to local contrast changes, driving stochastic quantum bump sequences and adapting receptive-field size, position, and motion. Synaptic feedback (bottom): Voltage differences between R1-R6 photoreceptors modulate quantal histamine release^40–45^, which binds to postsynaptic chloride channels on LMCs, generating hyperpolarising responses. This, in turn, drives excitatory synaptic feedback from LMCs to photoreceptors, balancing synaptic loads and enabling fast, phasic signal transmission. **c,** Predictive lateral signal spread. Because of the flower-patterned, overcomplete LMC receptive fields (**a**), when a moving object (purple disk) activates one LMC, it simultaneously begins to stimulate photoreceptors feeding into neighbouring LMCs. Thus, information about the object spreads laterally as a wavefront (purple columns spatiotemporally locked to the moving object) and predictively across the network. Moreover, as the R1-R7/8 photoreceptors within an ommatidium are mechanically coupled, a microsaccade in one can shift its neighbour into the light path, thereby activating it^1,9–11^. The newly engaged photoreceptor then drives an LMC in an adjacent neuro-ommatidium whose receptive field would otherwise miss this change, revealing a second, photomechanical mechanism of predictive lateral signal spread. **d,** Morphodynamics underlying hyperacuity. Because of the lateral component of photoreceptor microsaccades (sideways motion), the light beam entering an ommatidium can be clipped by its aperture (the neck formed by surrounding cone and pigment cells^1,9,11^), morphodynamically narrowing the moving photoreceptor’s receptive field. Similar neural enhancement occurs when a moving object is occluded by another (Fig. 6). This enables photoreceptors and the LMCs they innervate to differentiate two nearby moving objects (flies). Resolvability increases further when the rhabdomere moves photomechanically inward, away from the lens, thereby collecting light from a narrower angle^10,11^. Finally, mirror-symmetrical photoreceptor microsaccades in the two eyes further enhance resolvability, giving the flies’ frontal binocular range augmented hyperacute stereovision^10,11^. **e,** Synaptic high-frequency jumping accelerates transmission and minimises delay. During saccadic or high-contrast light-flash stimulation (left), photoreceptor responses can exceed the adaptive “floating” synaptic range (middle), resulting in partial signal clipping (mathematically analogous to the aperture clipping shown in panel **d**). Synaptic high-frequency jumping (right) redistributes power to higher frequencies, extending bandwidth up to ∼1,000 Hz. The resulting phasic LMC responses, supported by morphodynamic model simulations, show no measurable synaptic delay and peak before the corresponding photoreceptor responses. Inset: Morphodynamic refractory quantal sampling and synaptic frequency jumping enable predictive coding and motion-blur reduction in natural 3D environments, allowing LMCs to resolve hyperacute object motion. In contrast, classic stationary filter models produce temporally blurred responses and fail to reproduce this performance.

Notably, high-frequency jumping is absent in recordings using Gaussian white noise (GWN; **Fig. 4d**), which increases refractoriness and desensitises phototransduction^9,55^. It is also missing from responses to linearly presented naturalistic image time series, dominated by slow frequencies (1/f statistics)^9,18,66,86^. Our results therefore suggest that animals’ *active vision* - combining high-speed saccadic movements with brief fixation pauses^9^, including photoreceptor microsaccades^11,12,27^, intraocular muscle contractions^87^, and eye, head, and body rotations^57^ - actively drives high-frequency jumping. This, in turn, enhances the phase congruency of visual features (such as edges, occluding objects, changing textures, and outlines), making them stand out instantaneously in the scene.

*How does behaviour shape neural synchrony to enable ultrafast, predictive sensorimotor processing?* During wakefulness, and especially during active behaviour, flies exhibit heightened visual responsiveness^81,84,88^. Photoreceptor-LMC synapses operate tonically, maintaining continuous interactions between bottom-up sensory input and top-down modulation to support attentional readiness^16–18,76,89^. This dynamic state allows the generation of widespread time-locked neural responses in reaction to behaviourally relevant environmental stimuli.

To understand how high-frequency jumping supports predictive sensorimotor processing, we examined the timing of photoreceptor and LMC responses to rapid stimuli. For example, *Musca* LMCs generate maximum responses within 6.5-9 ms of light onset (mean: 7.6 ± 0.8 ms, n = 10) - well before the associated photoreceptor voltage reaches its peak 5-9.5 ms later (p = 0.012), at 9-16 ms (mean: 11.6 ±1.9 ms, n = 20, **Supplementary Fig. 1a**). Similarly, the fastest reaction times during voluntary vision-driven behaviours fall within 13-20 ms (**Fig. 8**; **Supplementary Fig. 1c**), far shorter than expected under conventional unidirectional transmission models. These findings suggest that high-frequency jumping supports both local synaptic efficiency and global network synchronisation, enabling ultrafast, predictive sensorimotor responses.

To assess whether these principles generalise beyond the photoreceptor-LMC synapse, we reexamined response timings deeper in the fly brain. Supporting high-frequency jumping and morphodynamic synchronisation as general neural strategies, minimal-delay responses were also observed downstream in the *Drosophila* visual system^1,81^. In tethered flying flies, electrical activity recorded from the lobula and lobula plate (**Supplementary Fig. 1d**) - at least three synapses downstream of photoreceptors - appeared within ∼15-20 ms of stimulus onset^81^, closely time-locked to *Drosophila* LMC transients (**Fig. 7c**; **Supplementary Fig. 1b**). Likewise, an 18 ± 1.5 ms (mean ± SD) delay was recorded in the firing of *Drosophila* giant fibres - large command interneurons, four synapses downstream of photoreceptors and involved in collision-avoidance reactions - in response to light-off stimuli^90,91^. This ultrafast signal propagation indicates that latency does not accumulate strictly in proportion to synapse number. Instead, transient light changes evoke near-synchronous activity across successive stages of the optic lobe. These observations challenge classical models that assume slow, strictly sequential processing with substantial cumulative phototransduction and synaptic delays.

Thus, information processing in vivo appears more synchronised and integrated, with signals coordinated across multiple brain regions. This is reflected in the fly brain’s broadly distributed and dynamic energy use during activity^92^. Rather than conveying information sequentially like falling dominoes, neurons are coupled through morphodynamic and bidirectional synaptic interactions - with high-frequency jumping providing interlinked “strings” that cause the dominoes to fall together. Such synchronised, minimal-delay processing - from sensing to decision-making - is likely essential for supporting complex behaviours in real time.

*How do neural systems encode space through time to support accurate perception during rapid self-motion?* Analogous to recent concepts of human eye movements^93–95^, synaptic high-frequency jumping dynamically shifts neural processing of saccadic inputs into higher-frequency domains, enhancing predictive power and visual acuity. Our findings support this broader framework of encoding space through time^9,11,65,93–98^. Specifically, intracellular recordings and biophysically realistic modelling demonstrate how neural circuits actively transform transient visual signals - such as those elicited by saccades - to synchronise perception precisely with high-speed behaviours. This highlights a conserved principle of dynamic, spatiotemporal encoding across diverse visual systems.

Recent studies suggest that synaptic transmission involves ultrastructural mechanical movements^1,5–8,14,15^. Building on this, we propose that high-frequency jumping may be sensitised by stochastic ultrastructural oscillations - morphodynamic jitter - driven by tonic transmitter release. This process may help maintain neural processing and perception in an attentive, ready state at synapses transmitting both bottom-up and top-down signals^1^. Morphodynamic jitter, a form of mechanical stochastic resonance^99,100^, could enable interconnected circuits to respond in phase (i.e., synchronise) to dynamic or behaviourally relevant inputs, selectively amplifying salient signals while suppressing irrelevant ones. Additionally, jitter could temporally align bottom-up sensory signals with top-down motor predictions^101–103^, facilitating faster error correction and behavioural adaptation.

This study underscores the value of an integrative, multi-scale approach to understanding neural systems. By linking molecular, cellular, and systems-level dynamics with high-speed saccadic behaviour, we demonstrate how form, function, and behaviour co-adapt to support robust, adaptive vision in rapidly changing environments - enabling advanced computations already at the level of information sampling. This framework reveals emergent properties - such as high-frequency jumping, efficient coding, hyperacute vision, fast adaptive gain control, and predictive time-locking - that remain obscured when neural components are studied in isolation. For instance, morphodynamic sampling (encompassing photomechanical, stochastic, refractory, and quantal processes) and high-frequency jumping cannot be reproduced by conventional high-level reductionist models that treat photoreceptors and LMCs as static, unidirectional filters.

Beyond insect vision, our findings point to fundamental principles of neural computation. They offer new insights into enduring challenges, such as the neural binding problem, by showing how distributed, time-sensitive signals can be synchronised to generate unified percepts and high-speed, purposeful behaviour. More broadly, these principles could inform the design of next-generation artificial systems that, like biological vision, must operate efficiently under real-time constraints in noisy and dynamic environments.

Looking ahead, uncovering how morphodynamic high-frequency jumping generalises across sensory modalities and species may reveal fundamental laws of biological intelligence - laws that could drive the next revolution in adaptive, real-time artificial systems, from autonomous robots to predictive neuromorphic architectures.

To summarise, our results demonstrate that correctly explaining key performance limits of neural computation - including low effective synaptic delays, wide dynamic range^61,62,65,68^, and hyperacute spatiotemporal resolution^1,9,11^ - requires models that incorporate biological details usually abstracted away: quantal stochastic sampling, refractoriness and latency variability, ultrastructural motion, and naturalistic input statistics.

Using fly vision as an explicit case, we integrated experimental observations with biophysically realistic modelling of phototransduction and the photoreceptor-LMC synapse. In microvillar photoreceptors, vision is built from tens of thousands of discrete sampling units, where each absorbed photon triggers a stochastic quantum bump and transiently reduces local availability through refractoriness. Early cascade events also produce ultrafast photoreceptor microsaccades^2,9–11,56^, coupling ultrastructural motion directly to the timing and statistics of quantal responses. When driven by the bursty contrast dynamics characteristic of naturalistic behaviour, this quantal-stochastic-mechanical substrate naturally produces robust normalisation and gain control (supporting large dynamic range^1,9,65,68^) while sharpening timing relationships that enable hyperacute resolution. Extending the same mechanistic framework to synaptic transmission resolves a long-standing speed puzzle: under naturalistic contrast bursts, the photoreceptor-LMC synapse exhibits *high-frequency jumping*, redistributing transmission toward higher frequencies such that postsynaptic signals become effectively near delay-free, with bandwidths reaching ∼1,000 Hz and information transfer rates at thousands of bits·s^-1^.

Phenomenological filter-plus-noise or fixed-bandwidth descriptions can often be tuned to match selected response statistics, but they typically do so by imposing auxiliary assumptions - such as externally specified noise models, bandwidth caps, or efficiency constraints - rather than deriving speed, dynamic range, and hyperacuity from identified biophysical mechanisms. Here, these properties instead emerge as coupled consequences of quantal-stochastic-refractory sampling, ultrastructural morphodynamics and synaptic high-frequency jumping under naturalistic inputs, providing a direct mechanistic route from cellular microstructure to behaviourally relevant performance (**Fig. 9**).

## Methods

We provide here a brief overview of the main methods; full details of the multiscale experimental and theoretical approaches are in the **Supplementary Information (Supplementary Notes I–VI). Supplementary Note I** covers analyses of intracellular voltage responses from R1-R6 photoreceptors and LMCs under extended experimental paradigms (**Supplementary Figs. 1-18** and **Supplementary Tables 1-9**). **Supplementary Note II** describes high-resolution X-ray and electron microscopy (EM) analyses of *Musca* compound-eye optics and the photomechanical microsaccadic sampling underlying hyperacuity (**Supplementary Figs. 19-25** and **Supplementary Tables 10-12**). **Supplementary Note III** explains in vivo high-speed optical imaging of photoreceptor microsaccades (**Supplementary Fig. 26**). **Supplementary Note IV** details the mathematical modelling of the morphodynamic neural superposition system, incorporating adaptive optics (**Supplementary Figs. 27-38** and **Supplementary Tables 13-16**). **Supplementary Note V** presents the behavioural experiments (**Supplementary Fig. 39**). **Supplementary Note VI** describes the functional connectomics of Musca (**Supplementary Figs. 40-45** and **Supplementary Tables 17-20**).

### Fly stocks

Adult wild-type houseflies (*Musca domestica*) were used in the experiments. Housefly larvae and pupae were sourced from a commercial supplier (Blades Biological Ltd, Cowden, Kent, UK). They were cultured in a standard laboratory incubator (60% humidity) at the School of Biosciences and fed liver and sugar water. Flies were maintained at ∼22 °C under a 12:12 h light:dark cycle. In some experiments, adult wild-type *Drosophila* (Canton-S) reared separately at 25 °C served as controls^9^.

### In vivo intracellular recordings

Fly preparation and intracellular recordings were performed as described previously^9,104^. Briefly, houseflies were anaesthetised on ice. Once immobilised, their wings and legs were removed, and they were fixed to a conical holder (brass/plastic) using beeswax, securing the thorax, proboscis, and right eye to minimise movement artefacts. A small hole (covering 6-10 ommatidia) was cut in the dorsal cornea of the left eye to allow electrode access, and sealed with Vaseline to prevent drying^104^.

Voltage responses of R1-R6 photoreceptors and L1-L3 lamina monopolar cells (LMCs) were recorded using sharp, filamented borosilicate microelectrodes (Sutter Instruments; 1.0 mm outer diameter, 0.5 mm inner diameter), with resistances of 100-250 MΩ, pulled with a P-2000 horizontal laser micropipette puller. The tip of the reference electrode was cracked to reduce its resistance.

Photoreceptors and LMCs were recorded in separate sessions. Electrodes were back-filled with 3 M KCl for photoreceptors, and 3 M potassium acetate with 0.5 mM KCl for LMCs to maintain the chloride battery. The reference electrodes (blunt-tipped) were filled with fly Ringer solution (120□mM NaCl, 5□mM KCl, 5□mM TES, 1.5□mM CaCl₂, 4□mM MgCl₂, and 30□mM sucrose)^61^. Under a Nikon SMZ645 stereomicroscope, a remote-controlled micromanipulator (PM10, Mertzhauser) was used to position the electrodes. Thanks to the system’s stability, a single recording electrode often penetrated multiple photoreceptors - or occasionally LMCs - in sequence, yielding high-quality recordings from many cells within the same eye.

The fly’s temperature was maintained at 25□±□1□°C using a feedback-controlled Peltier device^61,104^. Only stable, high-quality recordings were analysed. In darkness, R1-R6 resting potentials were <□-60□mV, with responses ≥□45□mV to 100□ms saturating light pulses. L1-L3 cells showed dark resting potentials <□-30□mV and maximum responses ≥□20□mV. As LMCs were blindly penetrated and not stained, individual cell identities could not be confirmed; however, most were likely L1 or L2 due to their larger size^16–18^. Data from all recorded LMCs were pooled, given their similar response properties, including dark resting potential, hyperpolarisation to light increments, and response amplitudes.

### Ruling out efference copy interference in intracellular recordings

Stable intracellular recordings require firmly head-fixed flies to eliminate movement artefacts caused by bodily functions and to minimise recording noise. As a result, the fly cannot perform overt visual behaviours, such as head or body saccades, during stimulation. The saccadic bursty stimuli used in this study, therefore, only mimicked the light intensity time-series patterns (see ***Visual stimuli*** below) that such behaviours would normally generate. However, the fly could still produce internal motor-related signals - *efference copies* - typically associated with these behaviours^85,101–103^.

In our recording paradigm, the fly has no control over the stimulus. Thus, any efference copies generated by its motor circuits and transmitted downstream to photoreceptors or LMCs would not be synchronised with, or predictive of, the stimulus-driven responses. Instead, any such top-down signals would appear as sporadic, uncorrelated noise in the recordings.

We have refined our bespoke intracellular recording systems to achieve exceptionally low noise levels^9,16,61,62,65,80,104^. This enables us to accurately quantify neural responses and distinguish signal from noise under various visual conditions. Our signal-to-noise analyses of photoreceptor and LMC responses (see ***Data analysis*** below) have never revealed extrinsic noise patterns consistent with efference copies.

Indeed, all observed photoreceptor noise (variability in processing) can be fully explained and reconstructed from four well-characterised sources (see **Supplementary Notes IV**):

- *Quantum bump noise*^61,62^, modelled as a Lorentzian derived from the Gamma-shaped bump profile
- *Microsaccade noise*^9^, introducing a low-frequency hump
- *Synaptic feedback noise*^18^ from LMCs and amacrine cells, adding high-frequency Poisson-like variability shaped by the LMC waveform
- *1/f instrumental recording noise*^61,62^

Given this comprehensive account of known noise sources, the likelihood that efference copies influenced our recorded data under this specific experimental paradigm is exceptionally small. Moreover, our morphodynamic neural superposition model - lacking any built-in top-down circuitry - replicates both photoreceptor and LMC response dynamics (e.g. **Fig. 5**). Therefore, the present study does not provide evidence for efference copy influence on neural signalling at this early sensory sampling stage.

### Visual point-source stimuli

A high-intensity “white” LED (Seoul Z-Power P4 Star, 100 lumens) was used to deliver light stimuli centred on the receptive field via a randomised quartz fibre-optic bundle (180-1,200□nm transmission range) mounted on a rotatable Cardan-arm, subtending a 3° homogeneous light field^104^. Output was controlled by an OptoLED driver (Cairn Research Ltd, UK).

To characterise temporal encoding, five Gaussian white noise (GWN) stimuli were presented with varying 3 dB cut-off frequencies (20, 50, 100, 200, and 500 Hz; **Fig. 10**). Stimuli were 2 s long and sampled at 1,000 Hz, or 2,000 Hz where noted. Stimuli were generated using MATLAB’s *randn* function, low-pass filtered (MATLAB Filter Toolbox), and scaled to have flat power spectra and equal peak-to-peak modulation (two units). Each bandwidth was tested under three contrast conditions on a linear intensity scale: high (BG0, 0 background light units), mid (BG0.5, 0.5 units), and low (BG1, 1 unit). To capture the widest range of stimulus dynamics that *Musca* photoreceptors and LMCs may encounter during fast acrobatic flight, a subset of cells was additionally tested with higher-bandwidth stimuli with 3 dB cut-offs at 300, 600, and 750 Hz (**Supplementary Fig. 11**).

**Fig. 10.**
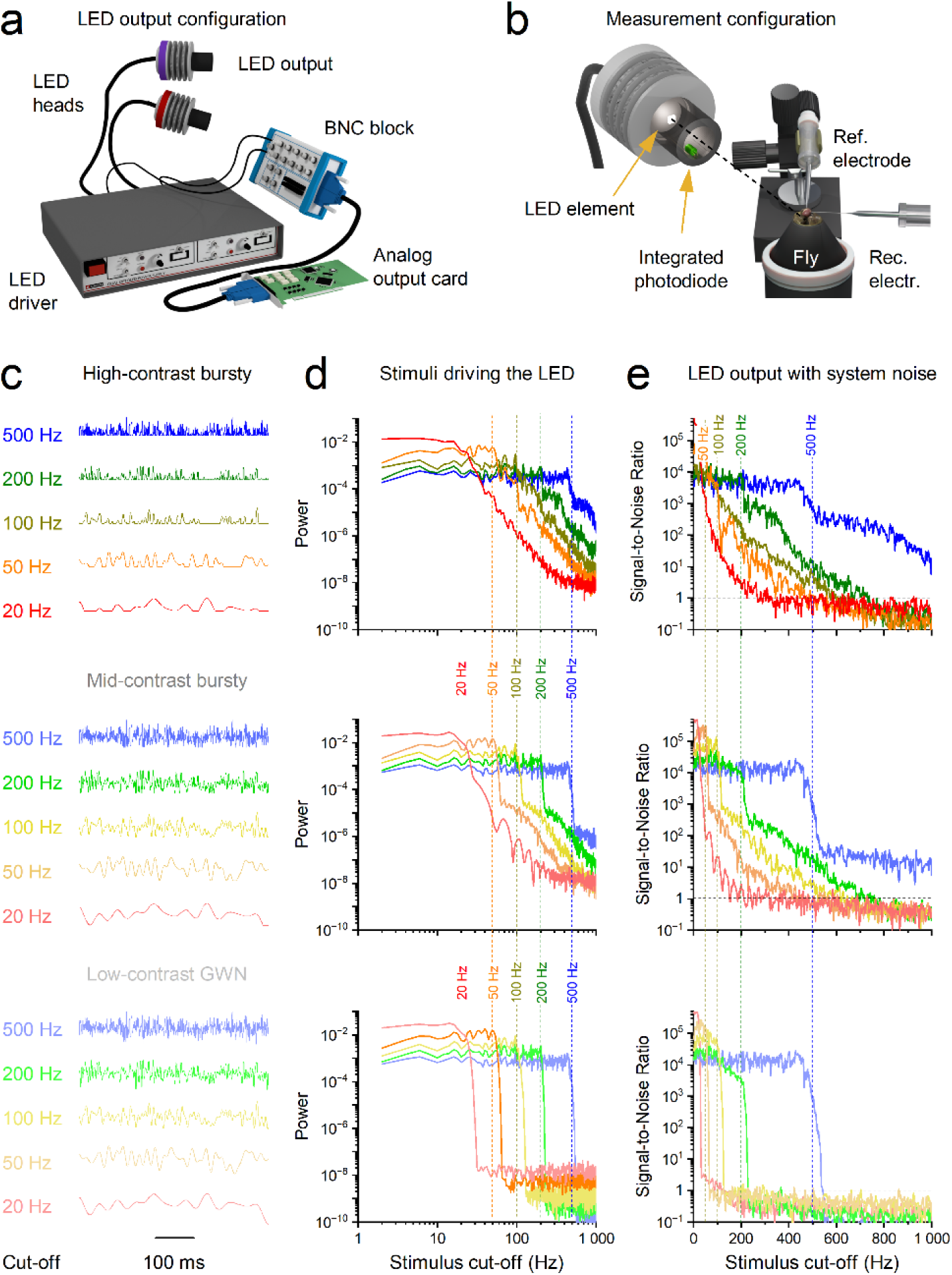
Generation, measurement, and signal-to-noise characterisation of Gaussian whitenoise and bursty visual stimuli. a, LED output configuration used for visual stimulation. A high-intensity white LED was driven by an OptoLED controller via an analog output card and BNC block, delivering temporally modulated light through fibre-optic LED heads. **b,** Measurement configuration. LED output was monitored directly using an integrated photodiode positioned at the fibre output, while intracellular recordings were made simultaneously from the fly photoreceptor or LMC using sharp microelectrodes and a reference electrode. This configuration allowed direct comparison between commanded stimulus waveforms and measured light output. **c,** Example time-domain waveforms of the three stimulus classes used: high-contrast bursty (“saccadic”) stimuli (top), mid-contrast bursty stimuli (middle), and low-contrast Gaussian white noise (GWN; bottom). For each class, stimuli were generated with 3-dB cut-off frequencies of 20, 50, 100, 200, and 500 Hz. All stimuli were scaled to equal peak-to-peak modulation and presented for 2 s. **d,** Power spectra of the stimuli driving the LED for each contrast class and cut-off frequency, showing flat spectra up to the specified 3-dB cut-off (vertical dashed lines) and rapid attenuation at higher frequencies. Colours correspond to stimulus cut-off frequencies as indicated. **e,** Signal-to-noise ratios (SNRs) of the measured LED output for each stimulus condition. Dashed vertical lines indicate nominal stimulus cut-off frequencies. Across all conditions, SNR remained high well beyond the 3-dB cut-off for bursty stimuli, confirming accurate temporal delivery of high-frequency components and defining the usable stimulus bandwidth for subsequent neural analyses.

Contrast was defined using Weber’s law:

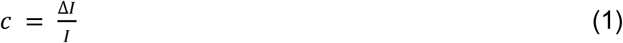

where *Δl* is the intensity change (standard deviation of the stimulus), and *l* is the mean background intensity. Measured contrasts were:

- High-contrast “saccadic” bursts: C(BC0) = 1.29 ± 0.13
- Mid-contrast bursts: C(BC0.5) = 0.61 ± 0.10
- Low-contrast GWN: C(BC1) = 0.33 ± 0.05

Stimuli were presented from the lowest to the highest adapting background. Prior to each stimulus, cells were dark-adapted for 20-30 s. Only cells with stable recordings across all 15 stimulus patterns were analysed. However, because LMC recordings are more difficult to maintain than photoreceptor recordings, in some cases only the five bursty stimuli were used for LMCs. Each stimulus was repeated at least 30 times per cell.

Stimuli and responses were low-pass filtered at 500□Hz or 1,000 Hz (KEMO VBF/23 elliptic filter, UK) and digitised at 1,000 or 2,000□Hz using a 12-bit A/D converter (National Instruments, USA). Data acquisition and stimulus control were handled via custom-written software (Biosyst, M. Juusola, 1997-2020) in MATLAB (MathWorks, USA)^61,66^, interfaced via the MATDAQ package (H.P.C. Robinson, 1997-2005) for National Instruments boards.

In a series of previous studies^9,55,65,66,68,69,105,106^, we systematically examined how fly photoreceptors encode naturalistic visual signals by directly comparing intracellular voltage responses and information transfer rates across multiple stimulus classes. These included: (i) linearly scanned light-intensity sequences extracted from natural images; (ii) saccadically scanned intensity sequences derived from natural images using measured body movements of freely walking flies^9,23^; (iii) randomly scanned intensity sequences from the same images; (iv) bandwidth-limited Gaussian white noise (GWN) stimuli with 3 dB cut-off frequencies spanning 20-500 Hz; (v) mid-and high-contrast bursty stimuli; and (vi) naturalistic light-intensity time series recorded directly from outdoor environments using van Hateren’s calibrated photometric dataset^86^.

Across these comparisons, photoreceptors consistently exhibited the highest encoding efficiency, signal-to-noise ratios, and information transfer rates when driven by high-contrast, bursty stimuli that reproduce the rapid, intermittent intensity transients generated during saccadic movements. In contrast, linearly scanned natural images, randomised scans, and GWN - despite having matched or higher mean intensity and comparable nominal bandwidth - elicited weaker responses and lower information throughput.

These experimental findings were further validated using a biophysically realistic stochastic, refractory, quantal photoreceptor model, which reproduced the same performance advantages under bursty stimulation. In the present study, we extend this experimentally validated modelling framework from *Drosophila* photoreceptors (which contain ∼30,000 microvilli) to *Musca* photoreceptors (∼54,000 microvilli), while preserving the same photomechanical phototransduction principles and stimulus-response relationships.

Thus, the saccadic light stimuli (mid-and high-contrast bursts) used here are not ad hoc abstractions but are grounded in direct comparisons with natural image statistics, measured fly behaviour, and independently validated biophysical models, ensuring that they accurately capture and adequately span the range of rapid intensity dynamics experienced during natural saccadic vision.

### Visual moving grating and dot stimulation

The moving visual-field and moving object stimulus methods and protocols, including the use of occlusions to clip the R1-R6s’ receptive fields to test photoreceptors’ and LMCs’ visual acuity, are explained in **Supplementary Notes I.7-8** (**Supplementary Figs. 14-18 and 24-25**).

## Data analysis

To ensure all the studied cells had reached a similar adaptation state, the first five responses (10 seconds of data) to the repeated stimulus were excluded from both signal and noise analyses. This left at least 25 responses to the same repeated stimulus pattern. The signal was defined as the mean response, and the noise as the deviation of individual traces from this mean^61,66^. Thus, n repetitions (n = 30) yielded one signal and 25 noise traces.

The signal s(t) and noise n(t) traces were segmented into 50%-overlapping stretches and windowed with a Blackman-Harris 4-term window. Each window produced six 500-point-or 1,000-point-long samples, corresponding to 1 or 2 kHz sampling, respectively. Fast Fourier transforms (FFTs) were applied to compute the frequency-domain signal and noise spectra, S(f) and N(f), respectively. The signal-to-noise ratio in the domain SNR(f) was calculated as:

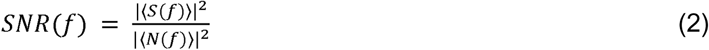

where |〈S(f)〉|^2^ and |〈N(f)〉|^2^ are the power spectra of signal and noise, respectively. Here v denotes voltage, || the absolute value, and O the average over all signal and noise windows^61,66^.

Information transfer rate (*R*) was calculated from the SNR(f) using Shannon’s information formula^79^, which is widely applied in this context^9,55,66^:

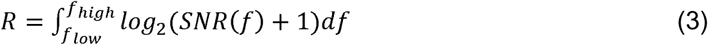

Signals were sampled at either 1 or 2 kHz and windowed accordingly (1,000-or 2,000-point Blackman-Harris window). Therefore, the integration bounds were 2-500 Hz (for 1 kHz sampling) or 1-1,000 Hz (for 2 kHz sampling), not 0 to ∞.

However, for LMC recordings sampled at the lower rate (1□kHz), Eq.□3 underestimates the true information transfer rates—particularly for mid-and high-contrast “saccadic” burst responses—because these evoked high-frequency jumping, with *SNR*(f) » 1 at 500 Hz (*cf*. **Fig. 3e**), indicating that frequencies above 500□Hz contributed non-negligible information. In contrast, this underestimation was not observed for responses to low-contrast Gaussian white noise (GWN) stimuli, which did not evoke high-frequency jumping and exhibited substantially lower response bandwidths (**Supplementary Fig. 7e**).

To correct for the high-frequency jumping effect, information losses in the 1□kHz recordings were estimated by comparison with matched 2□kHz recordings using the same stimuli. For photoreceptors (n□=□2), the mean information loss was approximately 5% and consistent across cells. For LMCs (n□=□2), the loss ranged from 5-23%, with the largest deficits observed for stimuli peaking near 200□Hz. The information transfer rate estimates for both photoreceptors and LMCs were corrected using stimulus-specific factors, calculated as the percentage difference between R_2-*500Hz*_ and R*_2-1,000Hz_* as defined by Eq.□3.

Moreover, information transfer rate estimates for LMCs during high-contrast bursts were less reliable than those for photoreceptors because the recorded voltage signals typically deviated from a Gaussian distribution—except when tested with a 500 Hz Gaussian White Noise stimulus (**Supplementary Notes I**, **Supplementary Fig. 4b**). Consequently, Shannon-based information estimates are most accurate under mid-and low-contrast conditions, where voltage responses more closely approximate Gaussian distributions (**Supplementary Fig. 7b**). Applying Shannon’s method to non-Gaussian responses, which violates its assumptions, may inflate estimates by ∼12%, as verified against the assumption-free triple-extrapolation method^66^ (**Supplementary Fig. 8**).

However, several factors contribute to underestimating the true capacity of the system (**Supplementary Fig. 9**). Microelectrode penetrations inherently damage the recorded cells, reducing signal fidelity. Additionally, responses cannot be measured in a true steady state, as they continually reflect ongoing adaptation, network dynamics^17,18,78^ and top-down eye-muscle activity^107^. Because recordings are never fully ergodic - trial-to-trial variability arises from adaptive network processes rather than pure noise - standard stationary information analyses mistakenly classify intrinsic or network-driven adaptations as additive noise. Thus, the actual information transfer rates and visual performance of housefly vision likely surpass our conservative estimates.

These analyses demonstrate that estimates derived from 25 repetitions (excluding the first five responses) are conservative rather than inflated; adaptive trends cause underestimation, not overestimation, of synaptic information transfer capacity. Therefore, our reported values represent a robust lower bound on the actual encoding precision of LMCs.

Flies also counteract motion blur during saccadic behaviours via multiple mechanisms: predictive stabilisation of head and body movements^25,26^, enhanced processing in acute zones^108^, and photomechanical, refractory light information sampling^9,11^. However, in our experiments, *Musca* were fixed in conical holders, eliminating head and body movement. We also used female flies and did not intentionally target the male acute zone, ensuring that neither electrode placement nor sex biased the results. Thus, the enhanced responsiveness of both photoreceptors and LMCs to fast, saccade-like stimuli reflects high-performance sampling and transmission dynamics^9–11^, which have evolved to support the fly’s high-speed visual behaviours and lifestyle.

### Morphodynamic Neural Superposition System

In the fly’s neural superposition eye (as viewed statically), six R1-R6 photoreceptors from different ommatidia converge onto shared downstream neurons (LMCs and an amacrine cell)^109^. Each photoreceptor is optically aligned to sample light from approximately the same region of space - but due to slight angular offsets and biological variability, their receptive fields do not perfectly overlap^11,48^. Instead, they sample from a small, fuzzy area, creating an over-complete, spatially jittered representation of the visual scene (**Fig. 2b**).

In a living fly, however, this system operates as a spatiotemporally dynamic morphodynamic network^11^. The receptive fields are not fixed; they shift in space and time due to photomechanical microsaccades - tiny, light-driven rhabdomere movements (**Fig. 2b-c**). This adds a temporal dimension to the over-complete spatial sampling, allowing receptive fields to sweep across fine spatial details and generate richer, decorrelated input patterns.

The result is a morphodynamic neural superposition system that enhances information encoding by:

- Dynamically refining receptive field alignment,
- Exploiting redundancy for noise suppression and error correction,
- Supporting high-frequency jumping and predictive coding aligned with behaviour.

This system transforms passive optical overlap into an active, synchronised sampling strategy, optimised for high-speed saccadic vision. **Supplementary Notes IV** details how we modelled the photoreceptor and LMC responses of this sophisticated morphodynamic system.

### Extrapolating *Musca* reaction times

To estimate the minimal reaction time of *Musca* antennal and leg responses to visual stimuli, we focused on two fast sensorimotor pathways: the light-induced antennal movement and the looming-induced leg escape response (see **Supplementary Notes VI**, **Supplementary Figs. 40-45**). Although no complete connectomic or genetic dataset exists for *Musca*, the neural architecture underlying the visual system is highly conserved between *Drosophila* and *Musca*^110^. We therefore used *Drosophila melanogaster*, for which complete connectomic data is available. Using the FlyBrainLab platform (see **Supplementary Notes VI** for details), we identified the shortest reflex-like pathways linking the retina to motoneurons controlling antennal and leg responses (**Fig. 7d,h**), with minimal synaptic gaps and conduction distances. Examining these pathways in Drosophila allows us to extrapolate *Musca*’s visual-motor response time.

We accumulate processing delays along the two *Drosophila* pathways, attributed to phototransduction and voltage integration, synaptic transmission delay, and conduction speed (see **Supplementary Notes VI**). While *Musca* has a larger brain, the main contributors to processing delay, phototransduction and synaptic transmission, are likely conserved, while a small difference in conduction delays may exist and we consider that the *Musca* has 3 times the body size as that of the fruit fly.

Together, these stages yield an estimated reaction time of 22.5-29.5 ms for the light-induced antennal movement and 18.6-22 ms for the looming induced leg escape response. We propose this as a reasonable lower-bound estimate for these visually induced motor responses in *Musca*, based on highly compact reflex-like pathways and conventional fast signalling assumptions. These estimates are at least 5-9 ms slower than our experimental observations of voluntary visual responses in flies (13□ms), suggesting the involvement of in vivo acceleration mechanisms (e.g., synaptic high-frequency jumping).

## Statistics

Statistical analyses were carried out in Python, Origin and MATLAB. Maximum information rates and visual acuity between male and female photoreceptors, and the whole population of photoreceptors and LMCs, were compared. The behavioural data were tested against statistical models. The statistical methods are explained in the **Supplementary Notes I-VI**.

### Illustrations and Videos

Apart from a small number of hand-drawn illustrations, most three-dimensional visualisations, Supplement Movies, and illustrative artwork were created using the open-source software Blender (Blender Foundation; https://www.blender.org), across multiple software versions.

## Acknowledgements

We thank B. Webb, R.C. Hardie, A. Nikolaev, A. Lin and J. Stone for discussions and comments, members of the Juusola laboratory for discussions, and the Duke X-ray imaging team and staff at EMBL Hamburg for their assistance. SBFSEM data of *Musca* ommatidia was generated in the Oxford Brookes University Centre for Bioimaging. Funding: This work was supported by Jane and Aatos Erkko Foundation Fellowships (MJ and JT), The Leverhulme Trust (RPG-2024-016: MJ and PV), the Biotechnology and Biological Sciences Research Council (BB/F012071/1, BB/D001900/1, and BB/H013849/1: MJ), the Engineering and Physical Sciences Research Council (EP/P006094/1: MJ) and Horizon Europe Framework Programme grant NimbleAI - Ultra energy-efficient and secure neuromorphic sensing and processing at the endpoint (LC).

## Author Contributions

Conceptualisation (MJ, JT, and NM), Investigation (NM, JT, MJ, JK, ADB, HM, AAB, KA, TR, BB, SS, YZ, MK, ML and PV), Methodology (MJ, JT, MK, AL, ED, PV), Project administration (MJ, ED, AL, PV, LC), Resources (MJ, PV, AL, LC), Software (JT, JK, HM, BB, SS, YZ, AL, and MJ), Supervision (MJ), Writing - main paper original draft (MJ), Writing - review & editing (MJ, JT, ADB, JK, HM, MK, AL, PV, MK, GdP, ED, LC and NM).

## Reporting summary

Further information on research design is available in the Nature Portfolio Reporting Summary linked to this article.

## Data availability

The data supporting the findings of this study are available from the corresponding authors upon request.

## Code availability

All the software code, including the complete *Musca* morphodynamic neural superposition model, can be downloaded from: https://github.com/JuusolaLab/High-frequency-jumping and https://github.com/JuusolaLab.

