## Supplementary Notes for "Synaptic high-frequency jumping synchronises vision to high-speed behaviour"

**This PDF file includes:**

**Supplementary Notes**

Supplementary Figs. 1-47 (as embedded in the relevant places in the Supplementary Notes)

Supplementary Tables 1-20 (as embedded in the relevant places in the Supplementary Notes)

Legends for Supplementary Movies 1 to 5

Supplementary References

**Supplementary Notes**

The compound eyes of the housefly (*Musca domestica*) capture visual information and transmit it to the brain through a layered neural network. In these Supplementary Notes, we describe our approach to investigating and modelling morphodynamic refractory quantal information sampling and processing in *Musca* photoreceptors and large monopolar cells (LMCs). Our aim is to understand how visual signals are transformed at the photoreceptor–LMC synapse and to compare these cells’ measured activity with model predictions. We also examine how the speed of these early processes - measured at the surface of the fly’s visual system - aligns with the demands of rapid, active visual behaviours. By combining intracellular voltage recordings, ultrastructural analyses of the compound eye, high-speed optical imaging, and biophysically accurate multiscale models, we show how early visual sampling and processing are tuned to saccadic behaviours, shaping the speed and flow of neural information and enabling time-locked, high-speed visual perception with minimal delay.

**Supplementary Notes** are organised in six **Sections** (**I-VI**), which explain the multiscale experimental and theoretical approaches used to study high-frequency jumping and the synchronisation of vision to high-speed behaviour. Their order broadly follows the presentation in the main paper.

I. **Analysing Intracellular Photoreceptor and LMC Voltage Responses**, pp. 3-40
II. **Analysing Compound Eye Static and Morphodynamic Optics**, pp. 41-54
III. **In vivo high-speed optical imaging of photoreceptor microsaccades**, pp. 55-56
IV. **Modelling Morphodynamic Neural Superposition System with Adaptive Optics**, pp. 57-101
V. **Behavioural Experiments**, pp. 102-105
VI. **Functional Connectomics**, pp. 106-116

Because **Sections I-VI** present new experimental approaches and theoretical models - many of which have not previously been combined to study collaborative morphodynamic neural encoding or insect vision holistically - we provide in-depth supporting evidence to illustrate their strengths and limitations in generating in vivo results and integrating new knowledge. To help readers follow and evaluate these sections, **Supplementary Fig. 1-47** and **Supplementary Tables 1-20** are embedded at relevant points throughout the text.

All models (written in MATLAB or Python) can be downloaded from: <https://github.com/JuusolaLab>

**Supplementary Notes Glossary**, pp. 117-120

**Supplementary Movie Legends**, pp. 121

**Supplementary References**, pp. 122-129

**I. Analysing Intracellular Photoreceptor and Large Monopolar Cells (LMCs) Voltage Responses**

**I.1. Both *Musca*’s and *Drosophila*’s Responses Indicate Morphodynamic Predictive Coding**

Photoreceptors in the fruit fly (*Drosophila melanogaster*) sample visual information efficiently during bursty (“saccadic”) light stimuli^1^. Ex vivo^1,2^ and in vivo^1,3,4^ recordings show that phototransduction triggers rapid and robust photomechanical contractions in response to transient light changes. Driven by refractory photon sampling^1-4^, these photomechanical adaptations enable receptive fields to dynamically scan the environment - allowing flies to resolve hyperacute spatial details beyond the optical limits of their compound eyes, while minimising motion blur during rapid saccades^1,4^. Photoreceptors extract most information from temporal light patterns resembling those experienced during natural saccadic motion^1^. These findings are also reproduced by biophysically realistic photoreceptor simulations^1,5,6^.

Photoreceptor output synapses to LMCs engage concurrently with feedback synapses^7,8^ - both top-down and lateral^9,10^ - as well as with gap junctions^11,12^, forming a highly interconnected circuit optimised to maximise visual information throughput^8,13-16^. The architecture of the compound eye - from phototransduction to synaptic integration - has thus co-evolved to efficiently represent the visual environment and support survival. However, species-specific differences in lifestyle and ecology^17-19^ mean that *Drosophila*’s visual performance cannot predict that of other flies. *Drosophila* is a small slow-flying, crepuscular fructivore, whereas the housefly (*Musca domestica*) is a larger, fast-flying diurnal omnivore. *Musca* exhibit faster phototransduction kinetics^20,21^ and possess optical adaptations - including larger lenses and smaller interommatidial angles^22-24^ - supporting higher spatial acuity. These differences suggest that *Musca* achieves even faster and more acute vision than *Drosophila*.

To illustrate these differences, **Supplementary Fig. 1** compares intracellular voltage responses of *Musca* (panel **a**) and *Drosophila* (panel **b**) R1-R6 photoreceptors and LMCs to a saturating 10-ms bright light flash, recorded in vivo at 25 °C. The responses emerge and develop more rapidly in *Musca*, consistent with its adaptation for faster vision and behaviour. In both species, photoreceptor and LMC responses begin rising simultaneously (*Musca* ~3.5 ms; *Drosophila* ~6.5 ms after stimulus onset), indicating no detectable synaptic transmission delay. Yet, in both cases, postsynaptic LMC responses reach their maxima significantly earlier than the corresponding photoreceptor inputs (by ~4 ms in *Musca* and ~13 ms in *Drosophila*), consistent with synaptic predictive coding mechanisms.

Remarkably, the fastest light-evoked voluntary antennae movements - driven by motor neurons, a minimum of six synapses away from the photoreceptors (see **Section VI**, **Supplementary Fig. 43**) - occur in close synchrony with photoreceptor responses (**Supplementary Fig. 1c**). *Musca* photoreceptors peak 9-16 ms after light onset, while the shortest latencies measured for antennae movements range from 13 (dotted line) to 20 ms. This tight correspondence between sensory input and motor output is striking - especially given that this behaviour is voluntary: in many instances, flies either choose not to respond or respond only after a considerable delay - and underscores the exceptional speed and efficiency of the housefly’s visual-motor processing.

Further evidence comes from local field potential (LFP) and extracellular spike recordings from the optic lobes of tethered, flying *Drosophila*^25^ (**Supplementary Fig. 1d**), exposed to abrupt motion stimuli. These signals show minimal delays, closely matching photoreceptor and LMC responses to light flashes. This suggests that higher-order motion processing in the lobula and lobula plate - at least three synapses downstream from the photoreceptors - remains tightly synchronised with phototransduction, which constitutes the absolute spatiotemporal bottleneck: the earliest, quantally constrained step in the neural sampling and integration of visual information.

Together, these findings reveal a ***dynamic form of predictive coding*** during synaptic information transfer, in which phasic neural responses are time‑locked to moving objects and rapidly changing temporal patterns - sharply contrasting with classical models where interneurons rely on static centre‑surround antagonism to exploit spatial correlations in natural scenes^26^.

Morphodynamic predictive coding dynamics are also reproduced in biophysically realistic simulations of the morphodynamic neural superposition system (see **Section IV** for biophysical explanations and modelling).

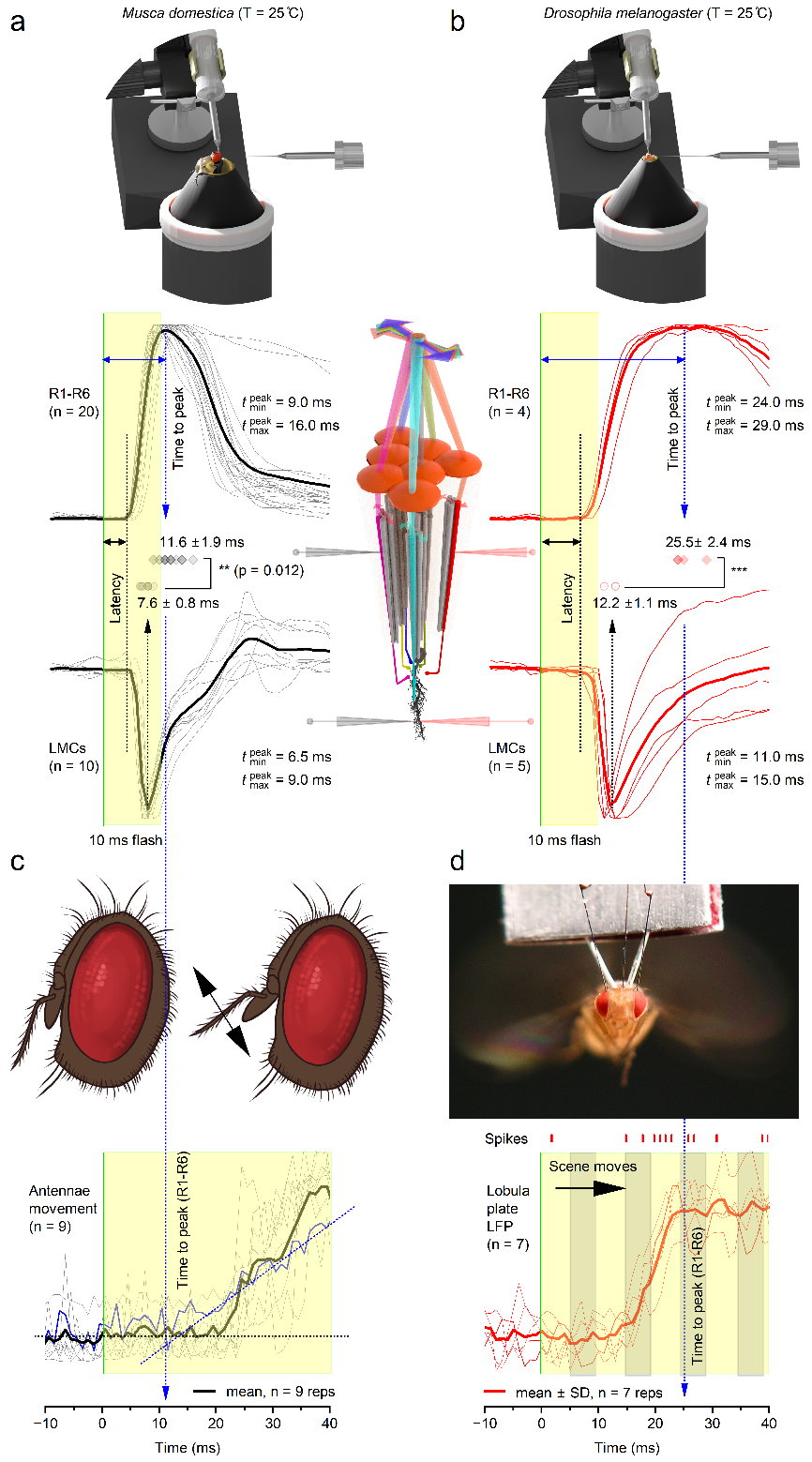
**Supplementary Fig. 1**. **Intracellular voltage responses of housefly (*Musca domestica*) and fruit fly (*Drosophila melanogaster*) R1-R6 photoreceptors are slower than those of their postsynaptic large monopolar cells (LMCs). Strikingly, LMC responses show minimal delays relative to stimulus onset, aligning closely with fast visual behaviours and higher-order neural processing.**

(**a**) *Musca* photoreceptor (top) and LMC (bottom) responses to a 10-ms white light flash following brief (5-15 s) dark adaptation, except for two photoreceptors with broader waveforms that were dark-adapted for longer (~5 min). However, extended dark adaptation had little effect on response rise time or time-to-peak. Both cell types show an identical response latency of ~4 ms from stimulus onset, indicating no measurable synaptic transmission delay. Notably, LMC responses peak significantly earlier (~7.6 ms) than photoreceptor responses, by approximately 4 ms.

(**b**) In *Drosophila*, photoreceptor and LMC responses are slower overall, consistent with the species’ slower, crepuscular lifestyle. Nevertheless, their relative response timing mirrors that in Musca: both cell types exhibit similar latencies (~6-7 ms) with no measurable synaptic delay, and LMCs again peak earlier than photoreceptors.

(**c**) Light-evoked voluntary antennal movements occur in close synchrony with Musca R1-R6 photoreceptor responses. Photoreceptors peak 9-16 ms after light onset; for this example, the fly's shortest antennal movement latencies range from ~13 ms (the extrapolated dotted blue line) to 20 ms (see **Figure 6** in the main paper for the fastest response distribution of all tested flies).

(**d**) Local field potentials (LFPs) and action potentials were recorded from the left optic lobe (most likely from the lobula or lobula plate, based on the strong visual motion sensitivity) of a tethered *Drosophila* flying in a flight simulator. In seven repeated experiments, LFP and spike activity consistently emerged ~10 ms after motion stimulus onset, earlier than the peak of photoreceptor responses to light flashes. LFP traces are sign-inverted for easier comparison. Data adapted from Tang and Juusola^25^, which provides full experimental details. All responses are amplitude-normalised to facilitate waveform comparison.

Next, in **Sections I.1-I.4**, we present a detailed analysis and statistical characterisation of the complete set of R1-R6 and LMC voltage responses to 15 dynamically changing light intensity time series, which contribute to the results shown in **Figures 2**, **3**, and **5** of the main paper.

**I.1. Individual Photoreceptor Responses Are More Consistent Than Population Variability**

**Supplementary Fig. 2** shows voltage responses from a single R1-R6 photoreceptor to repeated presentations of 15 different light stimuli, while **Supplementary Fig. 3** shows the average responses of 14 photoreceptors to the same stimuli.

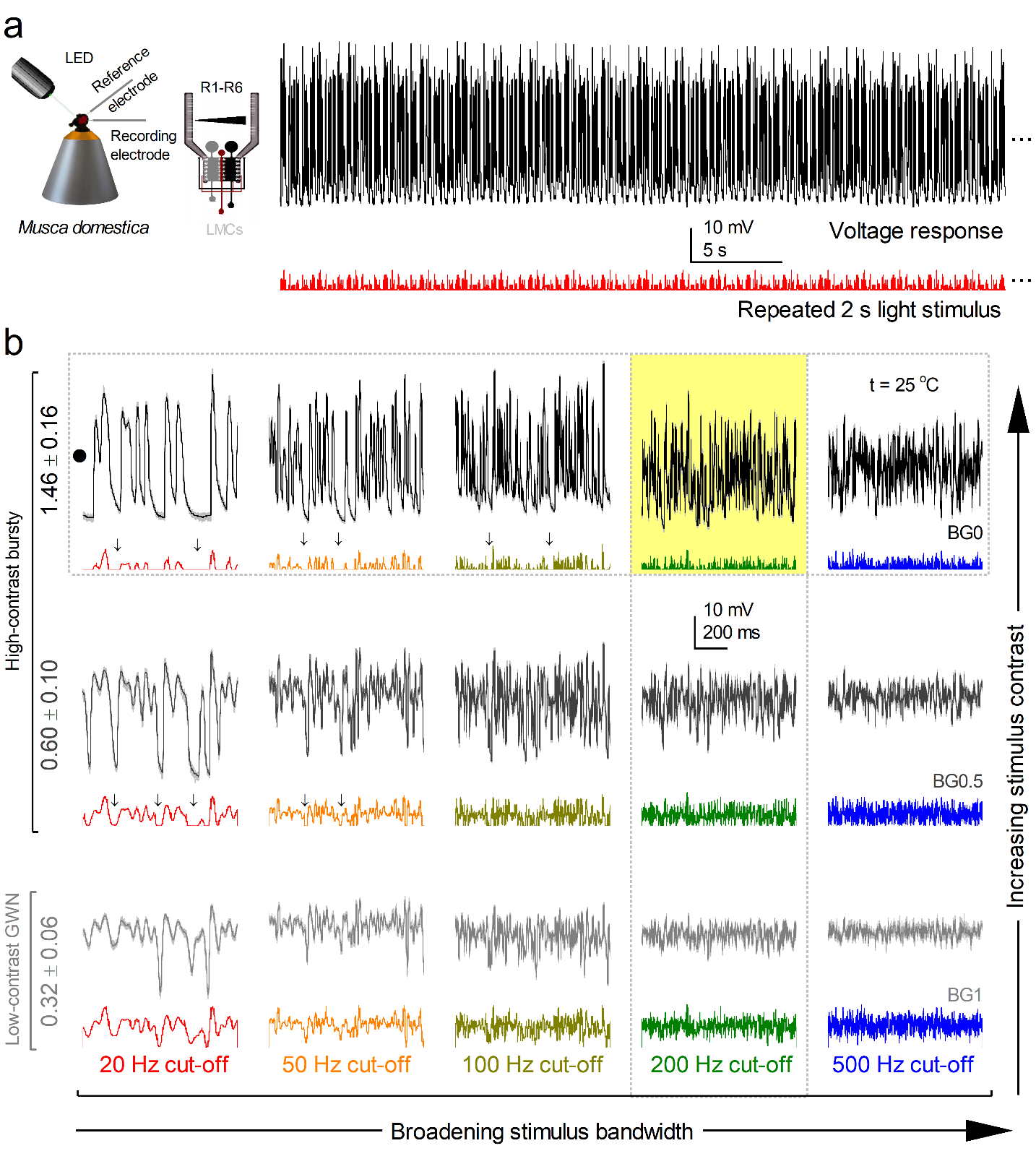

**Supplementary Fig. 2. High-contrast “saccadic” bursts maximise photoreceptor responses with minimal noise.**

(**a**) Left: Schematic of an in vivo intracellular recording setup targeting an R1-R6 photoreceptor in the Musca eye. Right: Example trace showing the repeated high-contrast bursty stimulus (red) and the corresponding photoreceptor voltage response (black) at 20 Hz bandwidth.

(**b**) R1-R6 responses to a range of stimuli varying in both contrast and temporal bandwidth. Stimuli range from high-contrast, saccade-like bursts (BG0) to low-contrast Gaussian white noise (GWN, BG1), each delivered at cut-off frequencies of 20, 50, 100, 200, and 500 Hz. Black and grey traces show the mean responses; thin, light grey traces represent 20 individual responses to different stimulus exemplars. Coloured traces below indicate the 15 different stimulus waveforms. The yellow box highlights the stimulus condition (200 Hz bandwidth) that elicits maximal information transmission. Arrows indicate some dark intervals embedded in the saccadic stimuli. The dotted horizontal and vertical boxes indicate stimulus conditions used in **Supplementary Fig. 4a-d** and **4e-h**, respectively. All data were recorded from the same photoreceptor cell, in a systematic sequence starting with low-contrast GWN (20-500 Hz) and ending with high-contrast bursts (20-500 Hz).

The antialiasing hypothesis^1,4,6,27^ proposes that small positional, structural, and morphodynamic differences among photoreceptors within the retinal sampling matrix reduce aliasing and enhance the reliability of visual information processing^28-30^. Consistent with this idea, intracellular recordings from in vivo houseflies reveal greater response variability across different photoreceptors (**Supplementary Fig. 3**; n = 14 cells) than within a single photoreceptor repeatedly stimulated with the same patterns (**Supplementary Fig. 2**; one representative cell). **Supplementary Tables** **1** and **2** give the corresponding statistics. In **Section IV**, we use a biophysically realistic morphodynamic model of refractory quantal information sampling and processing in the *Musca* neural superposition system to replicate (**Supplementary Figs. 34-35**) and mechanistically explain these results.

**Supplementary Table 1.** Characterising the R1-R6 responses reported in **Supplementary Fig. 1.** Each table entry consists of three descriptive statistics for the mean response trace of a single neuron under the specified contrast background (BG0-BG1) and stimulation bandwidth (20-500 Hz): the 95% percentile range (mV), the total number of peaks, and the median width of peaks.

|  | 20 Hz | 50 Hz | 100 Hz | 200 Hz | 500 Hz |
| --- | --- | --- | --- | --- | --- |
| BG0  (Bursts) | 95% range: 41.21 no. peaks: 20 peak width: 31.03 | 95% range: 41.22 no. peaks: 65 peak width: 11.78 | 95% range: 39.79 no. peaks: 210 peak width: 6.32 | 95% range: 32.97 no. peaks: 163 peak width: 4.63 | 95% range: 24.94 no. peaks: 167 peak width: 4.75 |
| BG0.5  (Mid) | 95% range: 35.00 no. peaks: 29 peak width: 42.02 | 95% range: 27.64 no. peaks: 67 peak width: 13.56 | 95% range: 28.19 no. peaks: 116 peak width: 7.24 | 95% range: 18.47 no. peaks: 152 peak width: 5.36 | 95% range: 12.25 no. peaks: 154 peak width: 5.52 |
| BG1  (GWN) | 95% range: 19.11 no. peaks: 30 peak width: 37.46 | 95% range: 13.95 no. peaks: 70 peak width: 13.20 | 95% range: 15.66 no. peaks: 120 peak width: 7.28 | 95% range: 9.22 no. peaks: 152 peak width: 5.49 | 95% range: 6.41 no. peaks: 152 peak width: 5.84 |

Instrumental noise was minimal, as evidenced by the high repeatability of responses (**Supplementary Fig. 2A**), consistent with earlier findings from fruit fly (*Drosophila*) photoreceptors^31^. This suggests that the observed waveform and amplitude differences (**Supplementary Fig. 3A**) likely reflect biological variation. Contributing factors include slight differences in recording position (frontal rhabdomeres are the longest^32^), sex-specific adaptations such as the male “love spot” - a high-acuity region in the male (male) eyes^33^ - and inherent distinctions among R1-R6 cells in rhabdomere size^1^, connectivity^9,10^ , and functional tuning^1,11^ (see **Section II**, **Supplementary Fig. 13**).

Despite this population-level variability, all photoreceptors adapted in a consistent manner to changes in stimulus contrast and bandwidth (**Supplementary Fig. 2B** and **3B**), aligning with predictions from morphodynamic refractory quantal photon-sampling theory^1,27^.

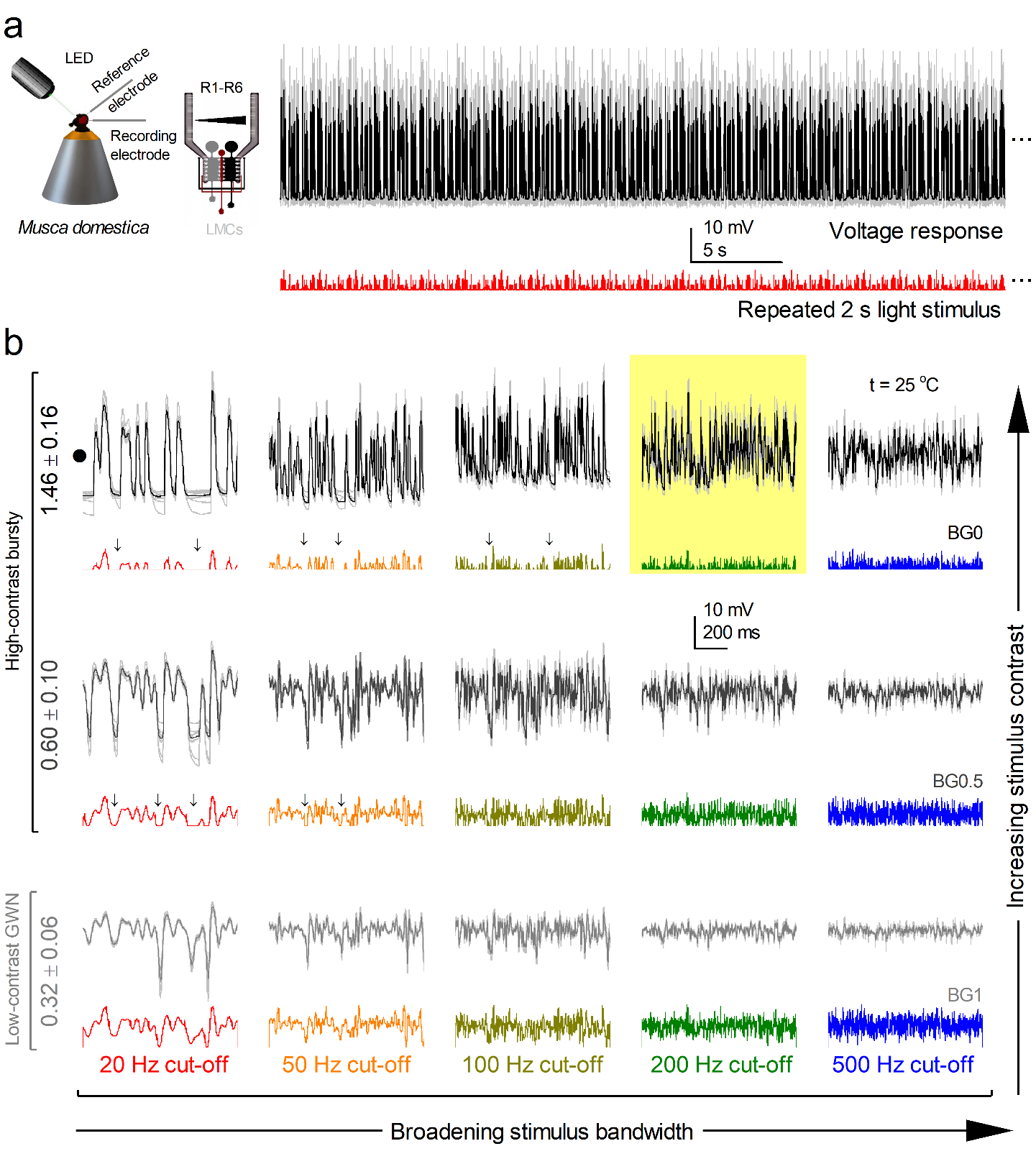

**Supplementary Fig. 3. R1-R6 photoreceptors respond most effectively to high-contrast saccadic stimuli.**

(**a**) Left: Schematic of the in vivo intracellular recording setup targeting R1-R6 photoreceptors in the *Musca* eye. Right: Example trace showing a repeated high-contrast saccadic stimulus (red) and the averaged photoreceptor responses (black trace) across all recorded cells (n = 14) at 20 Hz bandwidth.

(**b**) Population-level R1-R6 responses (n = 14) to a stimulus set varying in contrast and temporal bandwidth, ranging from high-contrast saccadic bursts (BG0) to low-contrast Gaussian white noise (GWN, BG1), delivered at 20, 50, 100, 200, and 500 Hz. Black and grey traces show the mean responses across all photoreceptors, while thin, light grey lines represent individual responses. Coloured traces below indicate the 15 different stimulus waveforms. The yellow box highlights the stimulus condition (200 Hz bandwidth) that elicits maximal information transmission across the population. Arrows mark dark intervals in the saccadic stimuli.

**Supplementary Table 2.** Characterising the R1-R6 responses reported in **Supplementary Fig. 3.** Each table entry summarises three descriptive statistics for the mean response trace of 15 photoreceptors under the specified contrast background (BG0-BG1) and stimulation bandwidth (20-500 Hz): the 95% percentile range (mV), the total number of peaks, and the median width of peaks.

|  | **20 Hz** | **50 Hz** | **100 Hz** | **200 Hz** | **500 Hz** |
| --- | --- | --- | --- | --- | --- |
| **BG0**  **(Bursts)** | 95% range: 29.45 no. peaks: 20 peak width: 30.53 | 95% range: 28.16 no. peaks: 62 peak width: 11.80 | 95% range: 28.24 no. peaks: 95 peak width: 6.83 | 95% range: 23.10 no. peaks: 127 peak width: 5.74 | 95% range: 16.41 no. peaks: 137 peak width: 5.95 |
| **BG0.5**  **(Mid)** | 95% range: 25.25 no. peaks: 26 peak width: 48.74 | 95% range: 19.87 no. peaks: 66 peak width: 13.03 | 95% range: 19.97 no. peaks: 110 peak width: 7.22 | 95% range: 11.96 no. peaks: 126 peak width: 6.45 | 95% range: 7.97 no. peaks: 137 peak width: 6.11 |
| **BG1**  **(GWN)** | 95% range: 17.10 no. peaks: 26 peak width: 44.09 | 95% range: 10.54 no. peaks: 65 peak width: 13.20 | 95% range: 10.41 no. peaks: 104 peak width: 7.57 | 95% range: 5.77 no. peaks: 121 peak width: 6.84 | 95% range: 4.08 no. peaks: 125 peak width: 6.24 |

**I.2. R1-R6 Photoreceptors Capture Information Maximally from 200 Hz "Saccadic" Bursty Light Stimuli**

Because the efficiency and accuracy of photoreceptor sampling ultimately limit visual perception during behaviours^7,11,34^, we quantified what the observed response dynamics mean for the photoreceptor signalling performance.

**Stochastic Sampling by ~54,000 Microvilli Minimises Neural Noise**

We analysed each photoreceptor’s signal-to-noise ratio, ${SNR}_{PR}(f)$, across our full stimulus set (Methods), varying both bandwidth (**Supplementary Fig. 4a**) and contrast (**Supplementary Fig. 4e**). As anticipated, ${SNR}_{PR}(f)$ rises in direct proportion to the voltage response amplitude, which itself scales with contrast (**Supplementary Fig. 2b** and **3b**), matching observations in *Drosophila*^1^ and *Calliphora*^35^. At every bandwidth tested, high-contrast “saccadic” bursts outperformed Gaussian white-noise, with ${SNR}_{PR}(f)$ peaking during 20 Hz bursts. In some cells, ${SNR}_{PR}^{max}(f)$ exceeded 3,000 (red trace), indicating near-identical responses on successive trials - any remaining variability must therefore arise from instrumental noise^31^, long-term adaptation^36^, or intraocular muscle activity^1,37,38^, rendering intrinsic photoreceptor noise negligible during bright bursts.

Remarkably, even under high-contrast bursts at 100 Hz and 200 Hz, ${SNR}_{PR}(500 Hz)$ remains around 1 in some cells, indicating reliable encoding up to 500 Hz, or over twice the ~230 Hz flicker-fusion limit suggested by early ERG studies^39^. However, when stimulus bandwidth extends the photoreceptor’s temporal limits - especially at low contrast - ${SNR}_{PR}^{max}(f)$ declines, reaching its minimum for low-contrast, 500 Hz white-noise (**Supplementary Fig. 4e**, light grey trace).

This extraordinary fidelity arises from ∼54,000 microvilli per photoreceptor (ranging from 41,000 to 74,000 across the Musca compound eyes, as detailed in **Section II**), each acting as an independent, refractory‐limited photon sampler during brief, high‐contrast bursts such as rapid saccades^1,6,34,40^. These microvilli generate stochastic quantum bumps - single-photon responses - which are then dynamically integrated by the photoreceptor’s voltage-sensitive membrane. This integration averages out microscale variability to produce robust macroscopic voltage signals, suppresses intrinsic noise, and amplifies phasic transients, thereby broadening the photoreceptor’s operational bandwidth^1,6,34,40^.

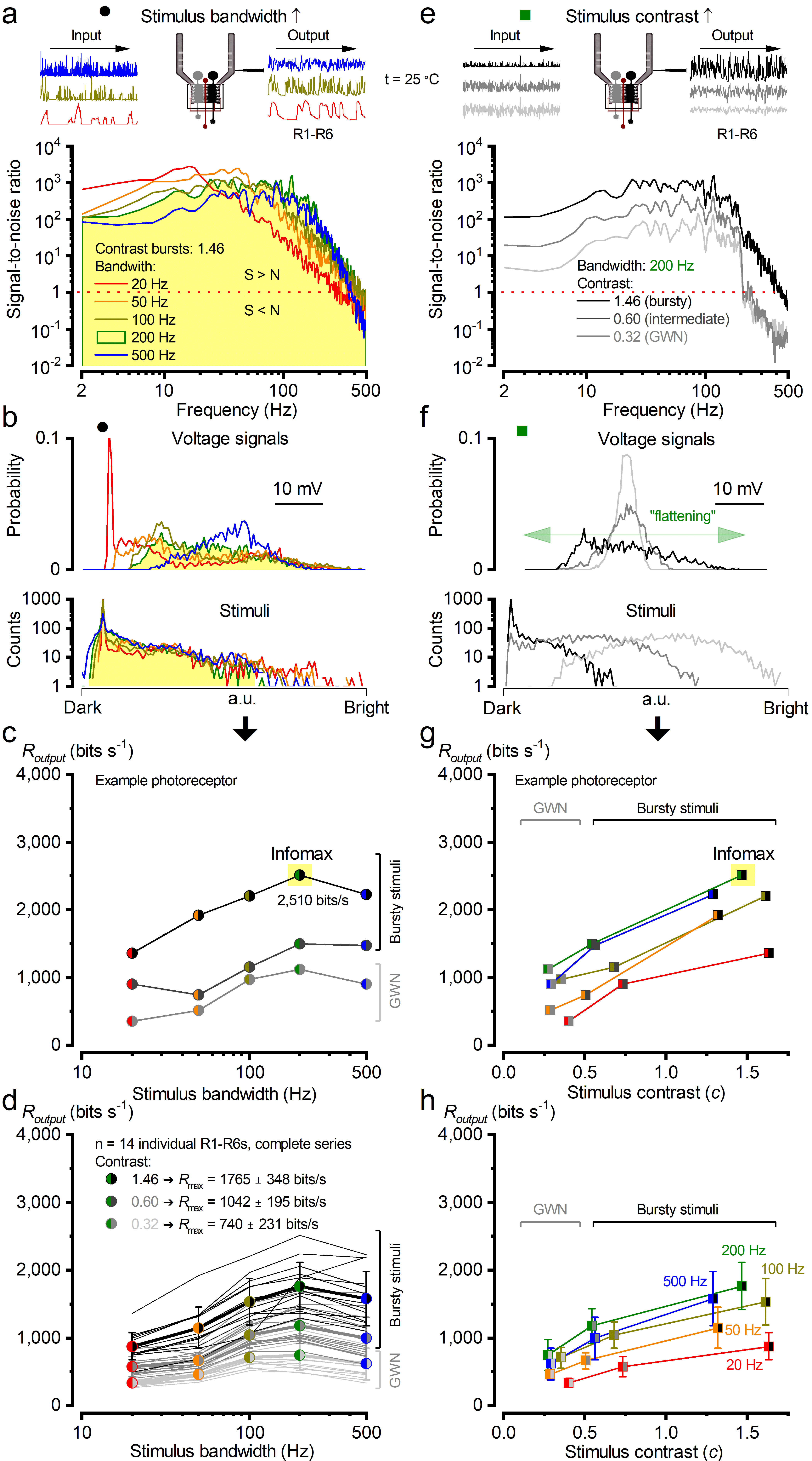
**Supplementary Fig. 4**. **Photoreceptors' information sampling peaks at 200 Hz and is about 2.4-times larger for high-contrast bursty stimuli than for GWN.**

(**a**) Response signal-to-noise ratio, ${SNR}_{PR}(f)$, when increasing the bandwidth from 20 Hz to 500 Hz.

(**b**) Top: All response probability density functions, PDFs, are nearly Gaussian except for 20 Hz and 50 Hz. Bottom: Stimulus intensity distributions are skewed. At 200 Hz, the photoreceptor had the broadest frequency and voltage distributions.

(**c**) The information transfer rates, $R_{output}$ (from a representative photoreceptor in **Supplementary Fig. 2**) were best for high-contrast "saccadic" stimuli, and it peaked at 200 Hz bandwidth (marked with a yellow box). The information transfer rates were calculated using the Shannon formula^41^.

(**d**) $R_{output}$ was calculated for all the photoreceptors (n = 14) to all the stimulus bandwidths. "Saccadic" bursts drove the maximal information transfer, which peaked at 200 Hz.

(**e**) Response ${SNR}_{PR}(f)$ at 200 Hz stimulus bandwidth when increasing the contrast from *c* ~0.32 to *c* ~1.46, being the highest for high-contrast bursty stimuli.

(**f**) Top: Response PDFs are approximately Gaussian for low- and mid-contrast stimuli, but become skewed under high-contrast bursty stimulation. This skew expands and flattens the photoreceptor output range, enhancing signalling efficiency. Bottom: Stimulus intensity distribution is Gaussian for low-contrast stimuli but slightly skewed for mid and high-contrast stimuli.

(**g**) Same as in **c**, but comparing against different contrast levels.

(**h**) Same as in **d**, but comparing against different contrast levels. Data in **a-c** and **e-g** are from the same photoreceptor (presented in **Supplementary Fig. 2**), whereas **d** and **h** show the results for all the photoreceptors (presented in **Supplementary Fig. 3**). Error bars show the standard deviation around the mean values.

**Supplementary Table 3.** A mixed-effects model predicts the information transfer rate in photoreceptors (using the complete datasets in **Supplementary Fig. 4**; n = 210 recordings from 14 cells).

| **Term** | **Estimate** | **Std. Error** | **z** | **p-value** | **95% CI (Lower-Upper)** |
| --- | --- | --- | --- | --- | --- |
| **Intercept (BG0 bursts)** | 6486.93 | 1031.39 | 6.29 | <0.001 | [4465.45, 8508.42] |
| **Contrast: BG0.5 Mid** | −96.18 | 103.18 | −0.93 | 0.351 | [−298.41, 106.06] |
| **Contrast: BG1 GWN** | −115.33 | 103.18 | −1.12 | 0.264 | [−317.56, 86.90] |
| **log10(bandwidth)** | −10589.10 | 1650.21 | −6.42 | <0.001 | [−13823.45, −7354.76] |
| **BG0.5 × log10(bandwidth)** | −197.86 | 50.16 | −3.95 | <0.001 | [−296.16, −99.55] |
| **BG1 × log10(bandwidth)** | −346.87 | 50.16 | −6.92 | <0.001 | [−445.18, −248.56] |
| **log10(bandwidth)²** | 6306.70 | 851.13 | 7.41 | <0.001 | [4638.52, 7974.88] |
| **log10(bandwidth)³** | −1130.70 | 141.64 | −7.98 | <0.001 | [−1408.32, −853.08] |
| **Random effect (Cell ID)** | Var = 40855.41 |  |  |  |  |

Mixed-effects regression revealed a significant (p < 0.001) nonlinear relationship between stimulus bandwidth and information transfer rate in photoreceptors. Interaction terms showed that the effect of frequency varied with contrast conditions (p < 0.001); with the interaction terms accounted for, the contrasts alone had no significant main effect. Cell ID was included in the model as a random effect to capture baseline differences between recordings (Var = 40,855). The final model demonstrated a strong overall fit (mean squared error = 18,400.88; R² = 0.92), suggesting that the combination of stimulus bandwidth and contrast level accounted for most of the variability in information transfer rates in photoreceptors.

**Information Capture Scales with Quantum-Bump Rate**

We estimated the information-sampling rate, $R_{output}$, for each photoreceptor by applying the Shannon formula^1,35,40-42^ to the measured $SNR(f)$^41^ spectra (Methods, **Supplementary Fig. 4**; statistics in **Supplementary Table 3**). This method assumes additive, Gaussian-distributed signal and noise. Indeed, most measured voltage‐response and noise broadly matched a Gaussian profile, especially under mid- to high-contrast burst stimuli (100-500 Hz) and low-contrast Gaussian white-noise (**Supplementary Fig. 4b**). These findings align with previous results from *Drosophila*^1^, and predictions from refractory photon sampling theory^1^. Using entropy-based, assumption-free extrapolation^1,35^, we previously demonstrated that residual skew in the stimulus distributions has a nominal effect (see also **Supplementary Fig. 8** and **9** below, which extend this analysis to LMC recordings).

Information rates increased markedly with stimulus contrast, aligning with our measured ${SNR}_{PR}(f)$ results. Higher quantum bump rate changes are integrated into larger, highly reproducible voltage responses^1,6,34,40^ (**Supplementary Fig. 2b** and **3b**), maximising information capture during high-contrast, 200 Hz "saccadic" bursts - when ${SNR}_{PR}(f)$ was highest across the broadest frequency and amplitude ranges (**Supplementary Fig. 4a** and **4e**). Under these conditions, individual photoreceptors achieved information transfer rates of approximately 1,300-2,500 bits/s, nearly threefold greater than the 400-800 bits/s rates obtained with Gaussian white-noise stimuli (**Supplementary Fig. 4c** and **4g**). This threefold enhancement was consistent across all 14 photoreceptors tested (**Supplementary Fig. 4d** and **3-4h**), closely matching previous findings in *Drosophila* (~850 bits/s for "saccadic" bursts compared to ~270 bits/s for GWN^1^).

The considerable variability in maximum information rates among photoreceptors corresponds to anatomical differences in rhabdomere length, which can vary 1.8-fold across regions of the eye (see **Section II**; **Supplementary Fig. 22g**, estimated microvillar counts range from ~41,000 to ~74,000 per photoreceptor). These findings underscore how the integration of tens of thousands of stochastic quantum bumps substantially enhances visual information capture - particularly under rapidly changing, high-contrast conditions such as those generated by houseflies’ high-speed saccadic movements in confined environments.

**The Myth of Slow, Noisy Photoreception**

Contrary to the traditional view that insect photoreceptors are sluggish integrators - “blinded” by the rapid light changes of saccadic eye movements^43^ - *Musca*’s R1-R6 cells actually excel at sampling fast, high-contrast bursts. Their responses peak in both amplitude and information rate when exposed to rapidly changing “saccadic” light contrast bursts, rather than being blurred or attenuated. This challenges the classic assumptions about noisy visual sampling^44,45^ and slow response integration^46,47^ impairing (“blurring”) compound eye vision during high-speed behaviours.

**A Simple Refractory Sampling Model Explains Information Capture from Photons**

We can reproduce these high-fidelity responses using a minimal stochastic photon sampling model governed by only four parameters (**Section IV**: 4-parameter photoreceptor model):

- Number of microvilli (∼54,000 independent sampling units per rhabdomere)
- Quantum-bump waveform (average shape and duration at the ambient intensity)
- Latency distribution (timing jitter before each bump)
- Refractoriness distribution (how long each microvillus stays unresponsive after a bump)

In this framework^1,6^, each absorbed photon triggers a quantum bump in one microvillus, after which the microvillus enters a brief refractory period^1,6,34,40^. Bursty stimuli, characterised by intermittent dark intervals, allow microvilli to recover between light contrast bursts, resulting in large and rapid increases in quantum bump rates and thus maximising information transfer^1^. In contrast, Gaussian white-noise stimuli hold photoreceptors near a steady mean brightness, causing many microvilli to remain refractory. This limits both bump rate and information throughput^1,34^. As a result, fewer microvilli generate quantum bumps during Gaussian white-noise stimulation, producing smaller responses that carry less information^6,34^.

For detailed explanations and model simulations, see **Section IV** below, which expands this framework to include photomechanical movements of microvilli (driving photoreceptor microsaccades) and synaptic feedback to photoreceptors that dynamically adjust information throughput.

**I.3. LMCs Respond Most Vigorously to "Saccadic" Bursts**

To assess how faithfully photoreceptor signals propagate, we recorded LMC voltage responses to the same light stimuli (**Supplementary Fig. 5** and **6**; **Supplementary Tables 4** and **5**). As expected, LMC outputs were polarity-inverted; histaminergic input hyperpolarises the cell in response to light increments and depolarises it following light decrements^13,48^. Under high-contrast conditions, particularly during 20-500 Hz bursts, LMC responses were dominated by large, phasic transients.

Together, the inhibitory feedforward and excitatory feedback photoreceptor↔LMC synapses^8,9,16,49^ act as a dynamic differentiator, selectively boosting rapid “bursty” contrast changes^8,13,16,31,48,50^ (*cf*. **Supplementary Fig. 2a** and **5a**)^48^. By pooling inputs from six photoreceptors^8,13,16,50,51^ - each with partially overlapping, dynamically shifting receptive fields^4,52^ - this synaptic ensemble amplifies fast fluctuations into higher-frequency voltage deflections, further sharpening and accelerating LMC responses (**Supplementary Fig. 5b** and **6b**).

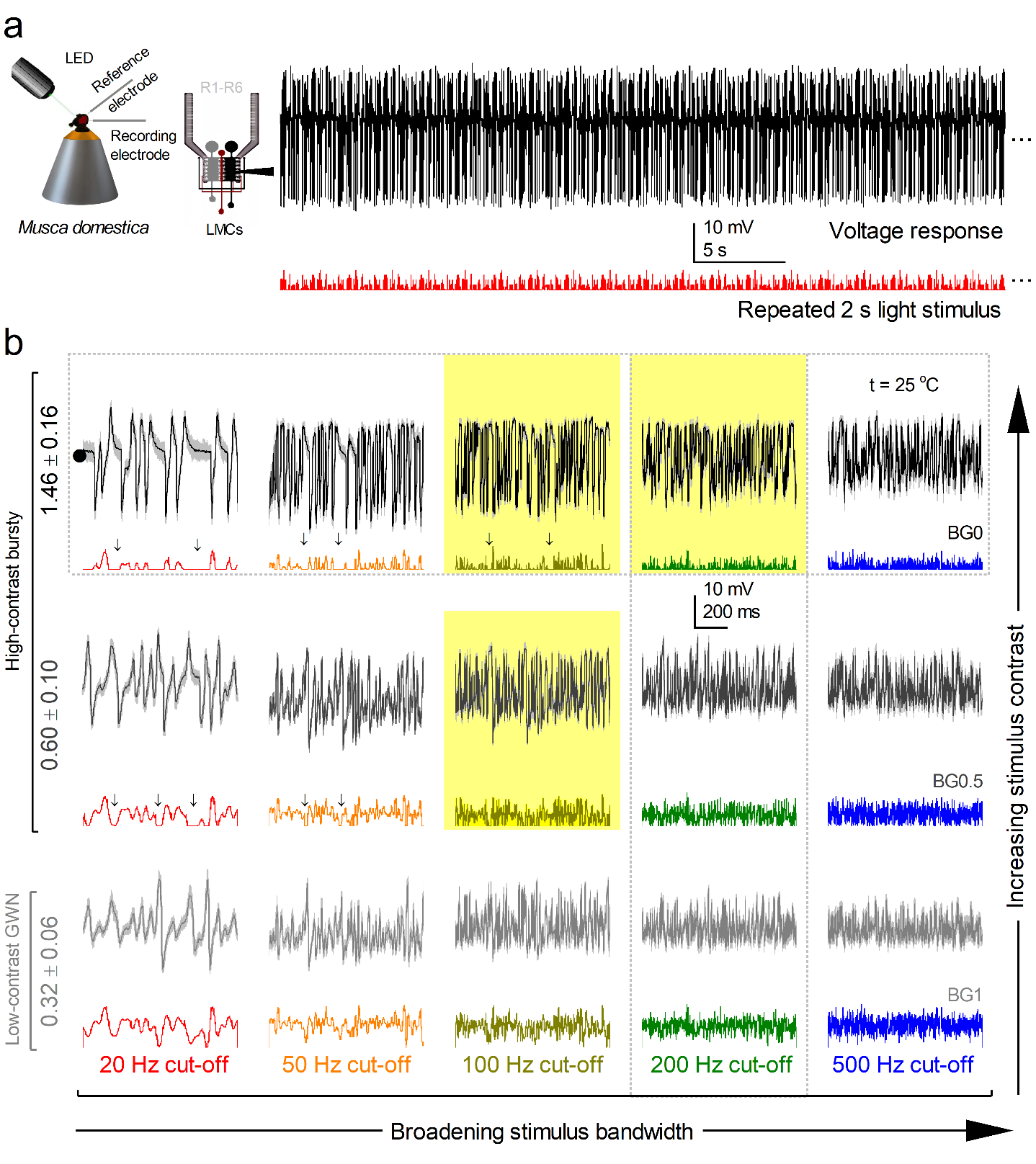

**Supplementary Fig. 5**. **Mid and high-contrast "saccadic" bursts maximise LMC's response.**

(**a**) Left: A schematic of in vivo intracellular recording from Musca eye. Right: An example of a repeated high-contrast bursty stimulus (red trace) and a response (black trace) at 20 Hz bandwidth.

(**b**) An L1-L3 response ranging from high-contrast "saccadic" bursts (BG0) to low-contrast GWN stimuli (BG1) in different cut-off frequencies, i.e. bandwidth patterns (20, 50, 100, 200 and 500 Hz). Mean (thick black and grey traces) and 25 individual responses (thin, lightly coloured) to 15 different stimuli (colourful traces beneath the responses). Yellow boxes: similar maximum information rates were carried by three sets of responses to different bursty stimulus patterns. Arrows: dark intervals in saccadic stimuli. Vertical dotted and horizontal rectangle: responses for bandwidth and contrast used in **Supplementary Fig. 7a** and **7e**, respectively. The recordings are from the same LMC.

**Supplementary Table 4*.*** Characterising the LMC responses reported in **Supplementary Fig. 5.** Each table entry summarises three descriptive statistics for the mean response trace of an individual LMC under the specified contrast background (BG0-BG1) and stimulation bandwidth (20-500 Hz): the 95% percentile range (mV), the total number of peaks, and the median width of peaks.

|  | **20 Hz** | **50 Hz** | **100 Hz** | **200 Hz** | **500 Hz** |
| --- | --- | --- | --- | --- | --- |
| **BG0**  **(Bursts)** | 95% range: 30.57 no. peaks: 34 peak width: 23.52 | 95% range: 31.03 no. peaks: 82 peak width: 11.66 | 95% range: 29.42 no. peaks: 107 peak width: 8.14 | 95% range: 24.49 no. peaks: 110 peak width: 7.13 | 95% range: 22.10 no. peaks: 131 peak width: 5.65 |
| **BG0.5**  **(Mid)** | 95% range: 26.82 no. peaks: 26 peak width: 27.71 | 95% range: 23.93 no. peaks: 73 peak width: 10.49 | 95% range: 23.41 no. peaks: 123 peak width: 5.39 | 95% range: 20.59 no. peaks: 156 peak width: 4.48 | 95% range: 16.72 no. peaks: 168 peak width: 4.28 |
| **BG1**  **(GWN)** | 95% range: 20.23 no. peaks: 27 peak width: 28.64 | 95% range: 18.75 no. peaks: 73 peak width: 10.35 | 95% range: 19.24 no. peaks: 130 peak width: 5.41 | 95% range: 13.87 no. peaks: 164 peak width: 4.23 | 95% range: 11.35 no. peaks: 172 peak width: 4.41 |

As expected^8,13,16,50^, LMC response amplitude grew with increasing contrast (**Supplementary Fig. 5b** and **6b**, rows) - broadly mirroring photoreceptor inputs (**Supplementary Fig. 2b** and **3b**). However, whereas photoreceptors show large amplitude gains with contrast, LMCs exhibit much smaller modulation differences, consistent with *dynamic* (not static^53^) *synaptic gain normalisation* observed in *Drosophila* LMCs^8,16,54^, and predicted by divisive normalisation models based on rapid synaptic feedback regulation^8,55^. Likewise, increasing stimulus bandwidth reduces amplitude modulation in both cell types, but the decline is less pronounced in LMCs (**Supplementary Fig. 5b** and **6b** vs **Supplementary Fig. 2b** and **3b**, columns). Crucially, despite these adaptive trends, LMCs responded most vigorously to both mid- and high-contrast "saccadic" bursts (**Supplementary Fig. 5B** and **6B**, yellow area) and far less to GWN.

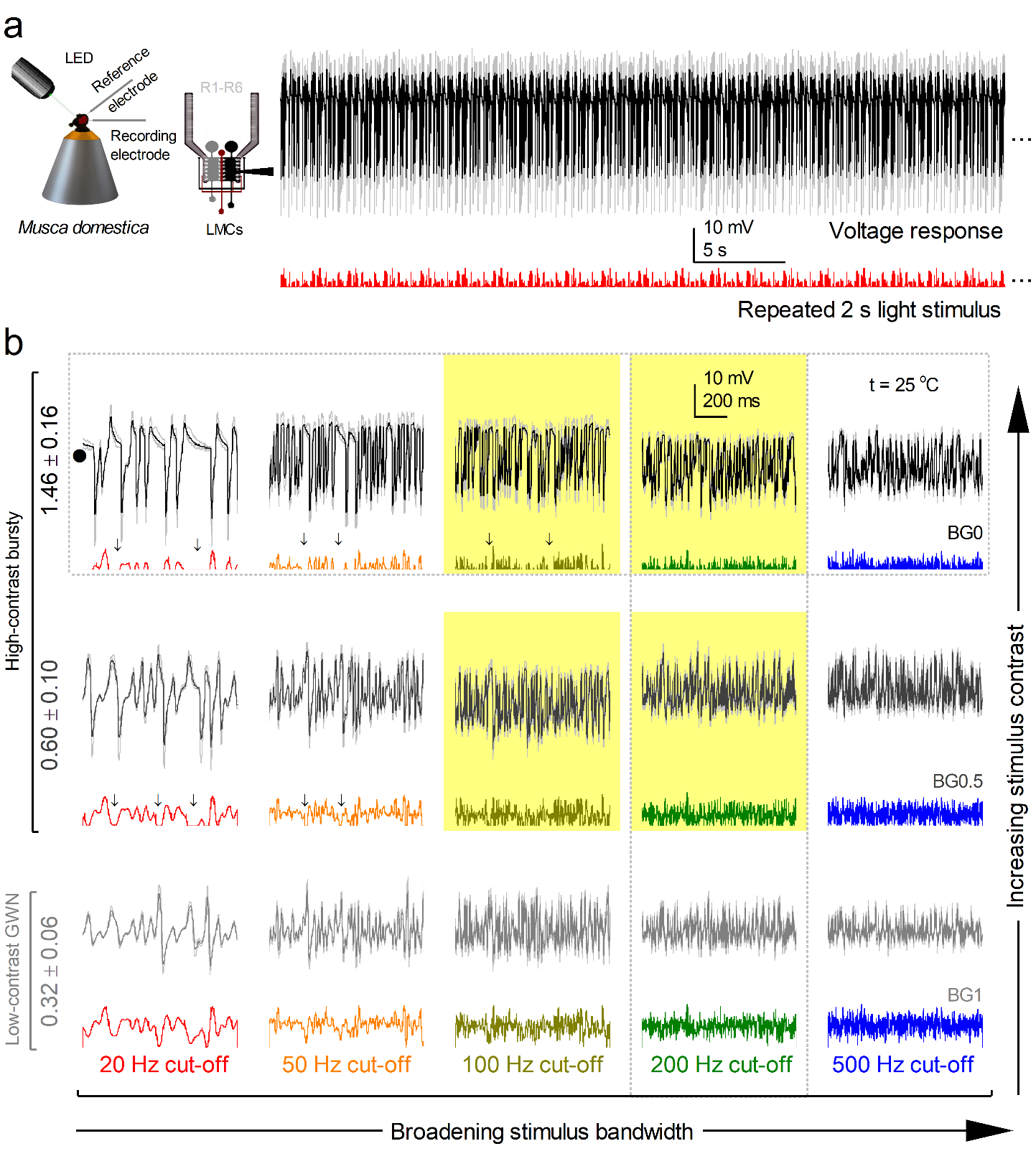

**Supplementary Fig. 6. All the LMCs respond best to mid and high-contrast saccadic stimuli.**

(**a**) Left: A schematic of in vivo intracellular recording from the *Musca* eye. Right: An example of a repeated high-contrast bursty stimulus (red trace) and responses of all the recorded LMCs (n = 3, black trace) at 20 Hz bandwidth.

(**b**) L1-L3 responses (n = 3) ranging from high-contrast “saccadic” bursts (BG0) to low-contrast GWN stimuli (BG1) in different cut-off frequencies, i.e. bandwidth patterns (20, 50, 100, 200 and 500 Hz). Mean of all the recorded LMC responses (thick black and grey traces) and individual LMC responses (thin, lightly coloured) to 15 different stimuli (colourful traces beneath the responses). Yellow box: maximum information responses of all the LMCs. Arrows: dark intervals in saccadic stimuli.

**Supplementary Table 5*.*** Characterising the L1-L3 responses reported in **Supplementary Fig. 6.** Each table entry summarises three descriptive statistics for the mean response trace of three LMCs under the specified contrast background (BG0-BG1) and stimulation bandwidth (20-500 Hz): the 95% percentile range (mV), the total number of peaks, and the median width of peaks.

|  | **20 Hz** | **50 Hz** | **100 Hz** | **200 Hz** | **500 Hz** |
| --- | --- | --- | --- | --- | --- |
| **BG0**  **(Bursts)** | 95% range: 27.36 no. peaks: 20 peak width: 57.02 | 95% range: 24.62 no. peaks: 65 peak width: 14.02 | 95% range: 22.99 no. peaks: 91 peak width: 9.98 | 95% range: 20.48 no. peaks: 108 peak width: 7.54 | 95% range: 18.49 no. peaks: 128 peak width: 6.12 |
| **BG0.5**  **(Mid)** | 95% range: 25.00 no. peaks: 25 peak width: 30.30 | 95% range: 21.09 no. peaks: 72 peak width: 10.82 | 95% range: 20.33 no. peaks: 120 peak width: 5.81 | 95% range: 17.46 no. peaks: 146 peak width: 4.60 | 95% range: 14.36 no. peaks: 157 peak width: 4.46 |
| **BG1**  **(GWN)** | 95% range: 11.86 no. peaks: 117 peak width: 4.31 | 95% range: 16.82 no. peaks: 72 peak width: 10.90 | 95% range: 16.79 no. peaks: 130 peak width: 5.67 | 95% range: 12.23 no. peaks: 156 peak width: 4.53 | 95% range: 10.13 no. peaks: 158 peak width: 4.59 |

**LMCs Show Variable Responses but Consistent Adaptation Dynamics**

Across the population (**Supplementary Fig. 6**, n = 3 LMCs tested with all the stimuli; **Supplementary Table 6**), LMC response amplitudes varied more between cells (**Supplementary Fig. 6**) than within a single cell (**Supplementary Fig. 5**), echoing the heterogeneity seen in R1-R6 photoreceptors. Since LMCs are smaller than R1-R6s^9,10^, and their largest signals arise in fine dendritic arbours, long‐term recordings inevitably sample different compartments (soma, dendrite, or axon)^16,54^. Thus, the inter-cell differences likely reflect both variable electrode placement and subtle functional distinctions^8,13,16,31,54^ among LMC subtypes^56^, despite all cells sharing the same adaptation kinetics (*cf*. **Supplementary Fig. 5b** and **6b**).

Overall, the relatively minor differences observed in response waveforms (when peak-to-peak normalised) across the different LMCs we recorded (n ≈ 20) support the idea that synaptic inputs to L1 and L2 cells are virtually identical at the dendritic level in the lamina^8,16,54^ (*cf*. **Supplementary Fig. 1a**), consistent with both cell types receiving the same number of histaminergic synapses^9,49^. This situation changes markedly at the medulla terminals, where L1 and L2 interact with different neighbouring cells and use distinct neurotransmitters^57^ to initiate the light On and Off visual channels^58^, respectively.

**LMCs’ Information Transfer Peaks during 100-200 Hz “Saccadic” Bursts**

We quantified each LMC’s adaptive encoding by computing its frequency-dependent ${SNR}_{LMC}(f)$ and Shannon information rates across all stimuli (Methods; **Supplementary Fig. 7**). Our core dataset comprised three LMCs challenged with the full stimulus battery, supplemented by two additional cells recorded during only the bursty sequences (all from different flies) to bolster our analysis of “saccadic” inputs.

Under bursty stimulation (**Supplementary Fig. 7a-b**), ${SNR}_{LMC}(f)$ was especially high - far above 100 - and rose steadily with the stimulus bandwidth up to 50-100 Hz, reflecting the synapse’s high repeatability and adaptive gain (**Supplementary Fig. 5**), much like the upstream photoreceptor signals (**Supplementary Fig. 4**). In our best recordings, ${SNR}_{LMC}(f)$ peaked at ~1,250 (**Supplementary Fig. 7a**), indicating nearly noise-free synaptic transmission for those burst patterns, before rolling off at still higher frequencies. Similarly, increasing stimulus contrast boosted LMCs ${SNR}_{LMC}(f)$ (**Supplementary Fig.. 7e**), whereas low-contrast white-noise produced the smallest gains.

Remarkably, even at 500 Hz - beyond the spectral content of our stimuli - ${SNR}_{LMC}(500Hz)$ remained well above 1 (**Supplementary Fig. 7a**), revealing reliable high-frequency jumping; unexpectedly fast synaptic encoding (e.g. **Figure 3**). Limited by 1 kHz acquisition of this subset (with a 500 Hz low-pass filter), we estimated an 18-26 % undersampling loss by comparing with our 2 kHz recording subset (**Figures 3**, **5**). These higher-bandwidth data confirm that LMCs can faithfully transmit phasic responses close to 1,000 Hz - over three times *Musca*’s classic 250 Hz flicker-fusion limit^39^.

Because information rate scales directly with ${SNR}_{LMC}(f)$, it too is maximised for mid- to high-contrast, 100-200 Hz bursts and declines under low-contrast or ultra-high-bandwidth stimuli. Together, these findings show that LMC synapses are exquisitely tuned to relay rapid, high-contrast “saccadic” signals with very high fidelity.

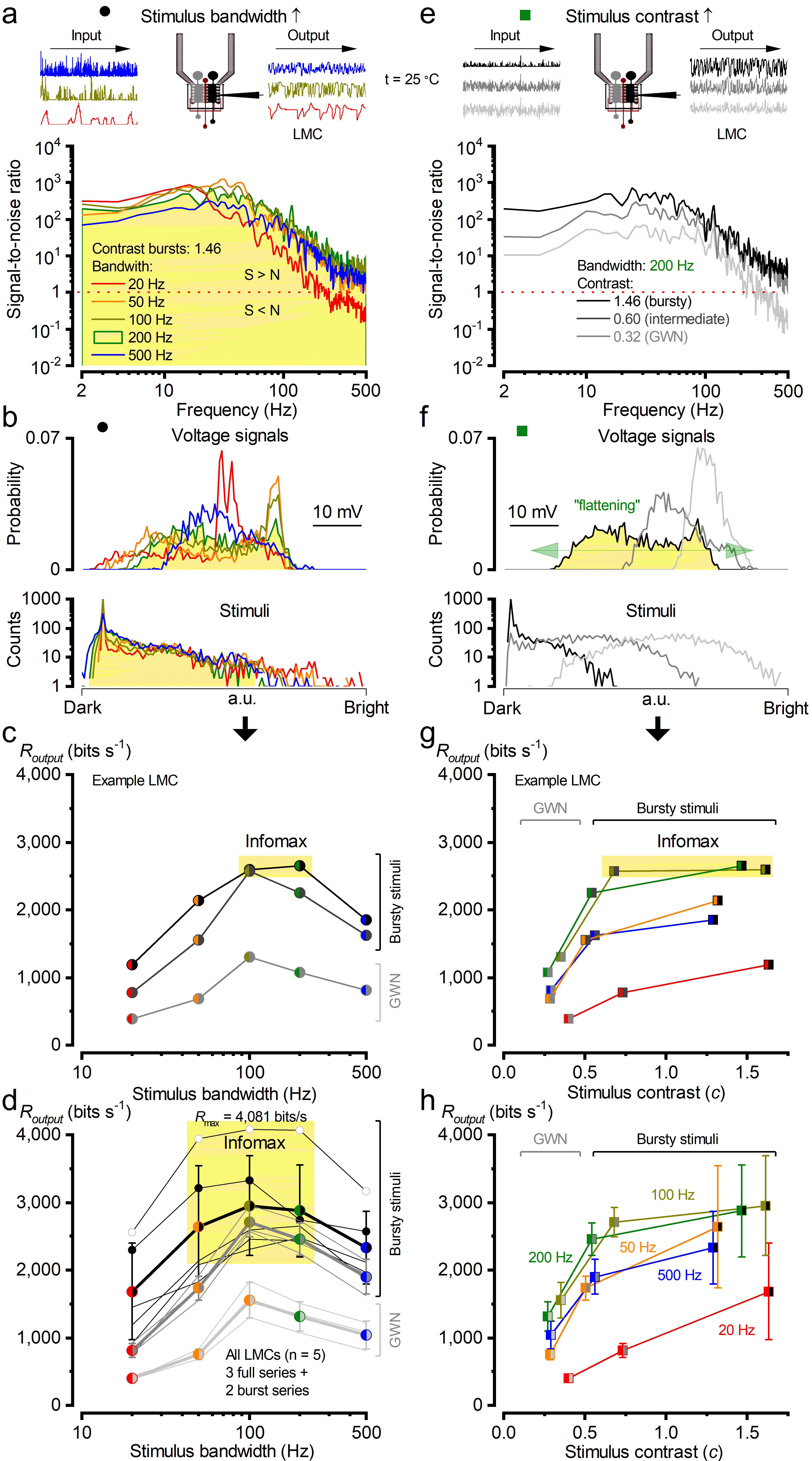
**Supplementary Fig. 7. LMCs' encode maximally 100 Hz and 200 Hz "saccadic" stimuli (both mid and high-contrast bursts).**

(**a**) Response signal-to-noise ratio, ${SNR}_{LMC}(f)$, to high-contrast bursty stimuli when increasing the bandwidth from 20 Hz to 500 Hz.

(**b**) Top: Response probability density functions (PDFs) to high-contrast bursty stimuli are not Gaussian except for 500 Hz. Bottom: Stimulus intensity distributions are skewed Gaussian. At 100 Hz and 200 Hz, the photoreceptor had the widest frequency and Gaussian voltage distributions.

(**c**) The information transfer rates, $R_{output}$, from a representative LMC in **Supplementary Fig. 5**) were best for mid-contrast and high-contrast "saccadic" stimuli, and they peaked at 100 Hz and 200 Hz (marked with a yellow box). $R_{output}$was calculated using the Shannon formula^41^.

(**d**) $R_{output}$ was calculated for three LMCs measured to all the stimuli, and two more cells were used to test the bursty bandwidth stimuli only. Mid- and high-contrast bursty "saccadic" stimuli drove maximal encoding, peaking at 100 Hz and 200 Hz, with a single LMC from a male fly transmitting over 4,000 bits/s. Due to high-frequency jumping, $R_{output}$ values from responses acquired at 1 kHz sampling rate (500 Hz bandwidth) were corrected using corresponding 2 kHz recordings (1,000 Hz bandwidth).

(**e**) Response ${SNR}_{LMC}(f)$ at 200 Hz stimulus bandwidth when increasing the contrast from *c* ~0.33 to *c* ~1.29. ${SNR}_{LMC}(f)$ is the highest for high-contrast bursty stimuli.

(**f**) Top: Response PDFs are approximately Gaussian for low- and mid-contrast stimuli, but become skewed under high-contrast bursty stimulation. This skew expands and flattens the LMC output range, enhancing signalling efficiency. Bottom: Stimulus intensity distribution is Gaussian for low-contrast stimuli but slightly skewed for mid and high-contrast stimuli.

(**g**) Same as in **c** but now comparing against different contrast levels.

(**h**) Same as in **d** but now comparing against different contrast levels. Data in **a-c** and **e-g** show results from the same LMC (**Figure 5**), while **d** and **h** pool all LMC results.

**Supplementary Table 6**. A mixed-effects model predicts the information transfer rate in LMCs (using the complete datasets in **Supplementary Fig. 7**; n = 45 recordings from 3 cells).

| **Term** | **Estimate** | **Std. Error** | **z** | **p-value** | **95% CI (Lower-Upper)** |
| --- | --- | --- | --- | --- | --- |
| **Intercept: BG0 Bursts** | −5976.22 | 671.42 | −8.90 | <0.001 | [−7292.18, −4660.26] |
| **Contrast: BG0.5 Mid** | −106.77 | 89.69 | −1.19 | 0.234 | [−282.56, 69.03] |
| **Contrast: BG1 GWN** | −1018.66 | 89.69 | −11.36 | <0.001 | [−1194.45, −842.86] |
| **log_10_(bandwidth)** | 7756.13 | 693.95 | 11.18 | <0.001 | [6396.02, 9116.24] |
| **log_10_(bandwidth)²** | −1773.70 | 172.44 | −10.29 | <0.001 | [−2111.68, −1435.72] |
| **Random effect (Cell ID)** | Var = 4119.75 |  |  |  |  |

Mixed-effects regression revealed a significant (p < 0.001) nonlinear relationship between stimulus bandwidth and information transfer rate in LMCs and a pronounced contrast effect for BG1 (but not for BG0.5), indicating the difference between bursty (BG0 and BG0.5) and Gaussian (BG1) stimuli. Cell ID was included in the model as a random effect to capture baseline differences between recordings (Var = 4119.75). The final model exhibited a strong fit (MSE = 52,274.12; R² = 0.90), indicating that stimulus bandwidth and the type of contrast (bursty vs. Gaussian) accounted for the majority of variance in information rates in LMCs.

**Evaluating LMCs’ information estimates**

Non-Gaussianity in signal and noise can bias information transfer rate estimates obtained with the Shannon method^41^. To assess this effect for the skewed LMC responses to 200 Hz saccadic burst stimulation (**Supplementary Fig. 7b**), we compared Shannon estimates with those from the *triple extrapolation method*^35,59^ (**Supplementary Fig. 8**), which makes no distributional or additivity assumptions and applies to any continuous signals of finite duration. Using the same dataset (2 kHz sampling) and systematically varying the triple extrapolation input parameters (data size, word length, voltage resolution), we found that the Shannon method yielded, on average, ~12% higher estimates (**Supplementary Fig. 8g**).

***Triple extrapolation***: In brief, *Musca* LMC responses to the same repeated 200 Hz bursty stimulus were digitised by dividing the recordings into time intervals, $T_{w}$, which were further subdivided into smaller bins of $t_{w}$ = 0.5 ms. This procedure defines “words” of length $T_{w}$, consisting of $T_{w}/t_{w}$ “letters”.

The mutual information between the response $S$ and the stimulus is then given by the difference between the total entropy, $H_{s}$:

$H_{S}=-\sum_{i} P_{S}(s_{i}){log}_{2} P_{S}(s_{i})$	(1)

where $P_{S}(s_{i})$ is the probability of finding the $i$-th word in the response, and the noise entropy, $H_{N}$:

$H_{N}=-\left\langle\sum_{i=1} P_{i}(\tau){log}_{2} P_{i}(\tau) \right\rangle_{\tau}$	(2)

where $P_{i}(\tau)$ denotes the probability of finding the *i*-th word at a time *t* after the initiation of the trial. This probability $P_{i}(\tau)$ was calculated across trials of using the same bursty stimulus pattern.

The values of the digitized entropies depend on the length of the “words” $T$, the number of voltage levels $v$ (upsilon) and the *size* (as %) of the data file, $H^{T,\nu,size}$. The rate of information transfer was obtained taking the following three successive limits (**Supplementary Fig. 8c-e**):

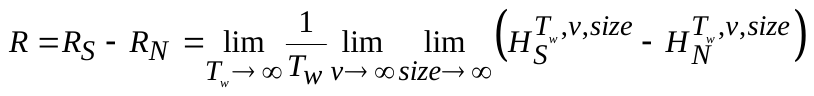
(3)

These limits were calculated by extrapolating the values of the experimentally obtained entropies. Here, a response matrix for the analysis contained 2,000 points x 25 trials. The total entropy and noise entropy of the recording was then obtained from the response matrices using linear extrapolation within the parameter ranges, as specified in **Supplementary Fig. 8e**. For estimating the information transfer rates, the total entropy and noise entropy extrapolation for $size$ and $v$ (**Supplementary Fig. 8c-d**) were performed with 2^nd^-order Taylor series, as such fits approximated these limits more accurately.

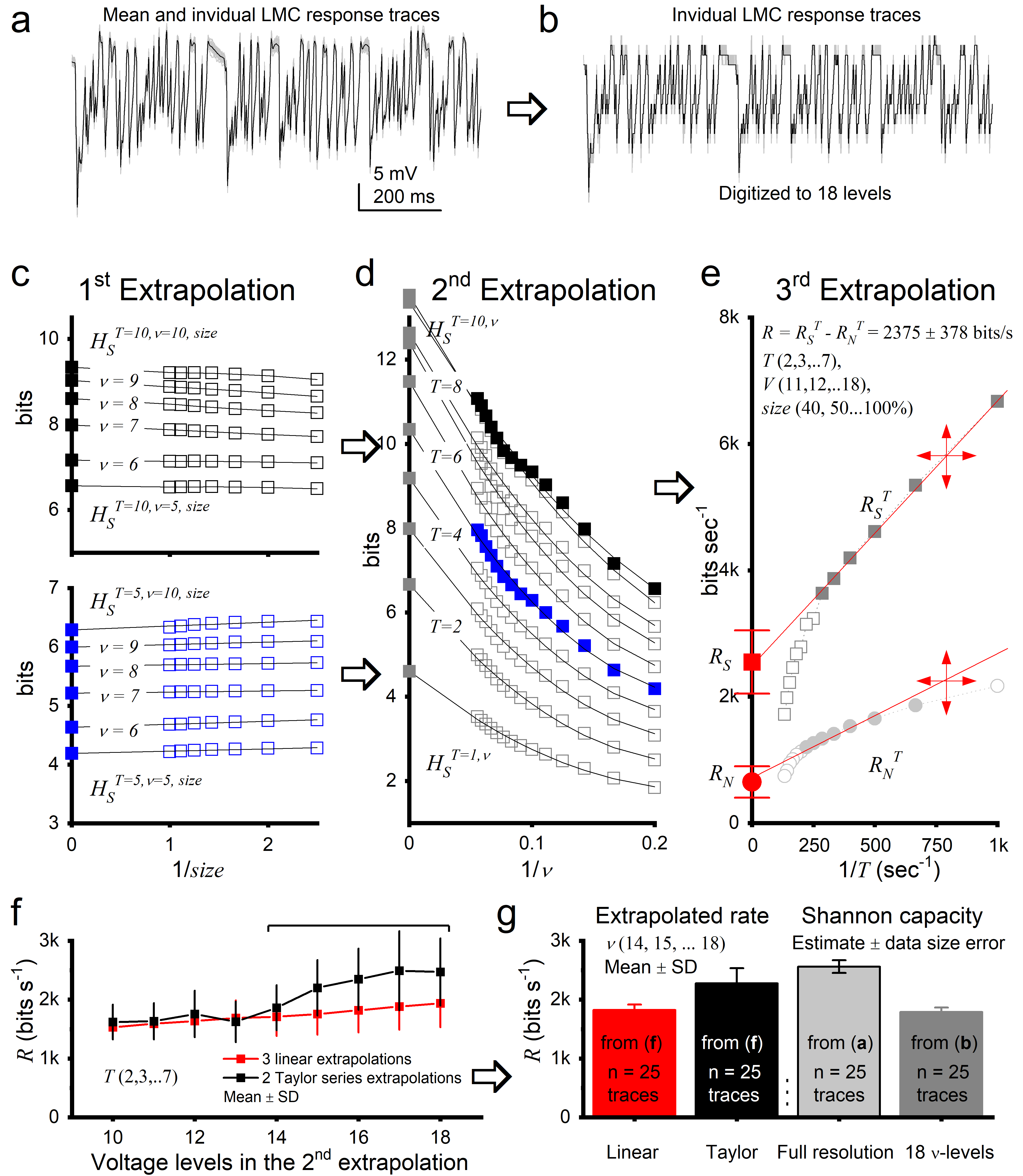

**Supplementary Fig. 8. Triple extrapolation method for estimating LMC information rates.**

(**a**) Mean (black) and 25 individual voltage responses (light grey) of a Musca LMC to a 1‑s 200‑Hz bursty stimulus.

(**b**) Responses digitised to 2-20 voltage levels ($\nu$); example shown for 18 levels. Total entropy ($H_{S}$) and noise entropy ($H_{N}$) were computed for $T$-letter words, where each 0.5‑ms letter corresponds to one voltage level, following Juusola and de Polavieja^35^.

(**c**) 1^st^ extrapolation: to infinite data size. Entropies of 10‑letter (top) and 5‑letter (bottom) words for 5-10 voltage levels fitted with linear trends. Extrapolation of $H_{S}^{T,\nu,size}$as 1/$size$ → 0 gives $H_{S}^{T=10,\nu,size} ($black) and $H_{S}^{T=5,\nu,size}$(blue). Size corrections are minor for 5‑letter words (probabilities similar across 40-100% of data) but more evident for 10‑letter words.

(**d**) 2^nd^ extrapolation: to infinite voltage resolution. $H_{S}^{T,\nu,size}$for words of 1-10 letters was fitted with 2nd‑order Taylor trends and extrapolated as 1/$\nu$ → 0. Squares: grey ($T$ = 5-10), blue ($T$ = 5), black ($T$ = 10).

(**e**) 3^rd^ extrapolation: to infinite word length. Total entropy rate ($R_{S}$, red square) and noise entropy rate ($R_{N}$, red circle) obtained by linear extrapolation as 1/$T$ → 0. When data are insufficient, both collapse to zero; here, linearity of the points allows reliable estimates of $R_{S}$, $R_{N}$, and information rate ($R$).

(**f**) Effect of voltage level number ($\nu$) on R. For $\nu$ ≥ 10, the first point of the 2nd extrapolation is set at the 5th level. Linear fits (red) and 2nd‑order Taylor fits (black) give similar estimates (<25% difference) for $\nu$ = 10-18.

(**g**) Comparison of $R$ estimates from the triple extrapolation method (linear fits, red; Taylor fits, black) and from the Shannon formula. Estimates from both methods are broadly consistent: Shannon gives ~12% higher rates for data in (**a**) and ~40% lower rates for 12‑level data in (**b**), indicating internal consistency of the approaches.

***Adaptation effect***: Moreover, data size and adaptation influence the estimated information transfer rates^35^. In our experiments, we typically recorded 30 repetitions of the same 2‑s bursty 200 Hz light stimulus from each LMC. To reduce the strongest adaptive effects, we discarded the first five responses and analysed the remaining 25. This step removes the most obvious drift in the recordings, but any residual adaptive trend still contributes apparent noise, thereby lowering the estimated information rate. As adaptation stabilises, this bias diminishes, and the estimates approach the system’s true synaptic transfer rate.

To assess the magnitude of this bias, we compared biological data with simulations of ergodic Gaussian white‑noise inputs^35^, which have additive signal-noise statistics but no adaptive components. For these simulated data, the Shannon method slightly overestimates information rate - by ~4.4% - when 25 repetitions of 1,000‑point segments are used compared with extrapolation to infinite data^35^. Thus, any large discrepancies in biological recordings are likely due to adaptive trends rather than to limitations of the Shannon formula itself.

We therefore systematically examined how removing early adaptive responses affects LMC information estimates (**S‑Figure 9**). Intracellular recordings show that early repetitions display a gradual adaptive drift, which broadly inflates frequency noise components in the signal‑to‑noise spectra (**a**-**b**). Sequentially discarding early responses - first five, then fifteen, and eventually twenty repetitions (**c**-**e**) - progressively flattens the noise traces and expands the frequency range over which the SNR remains above unity (**f**), extending reliable encoding towards 1 kHz. Correspondingly, Shannon‑based information transfer rates increase from conservative values to exceed 4,000 bits/s in the most stable segments (**g**).

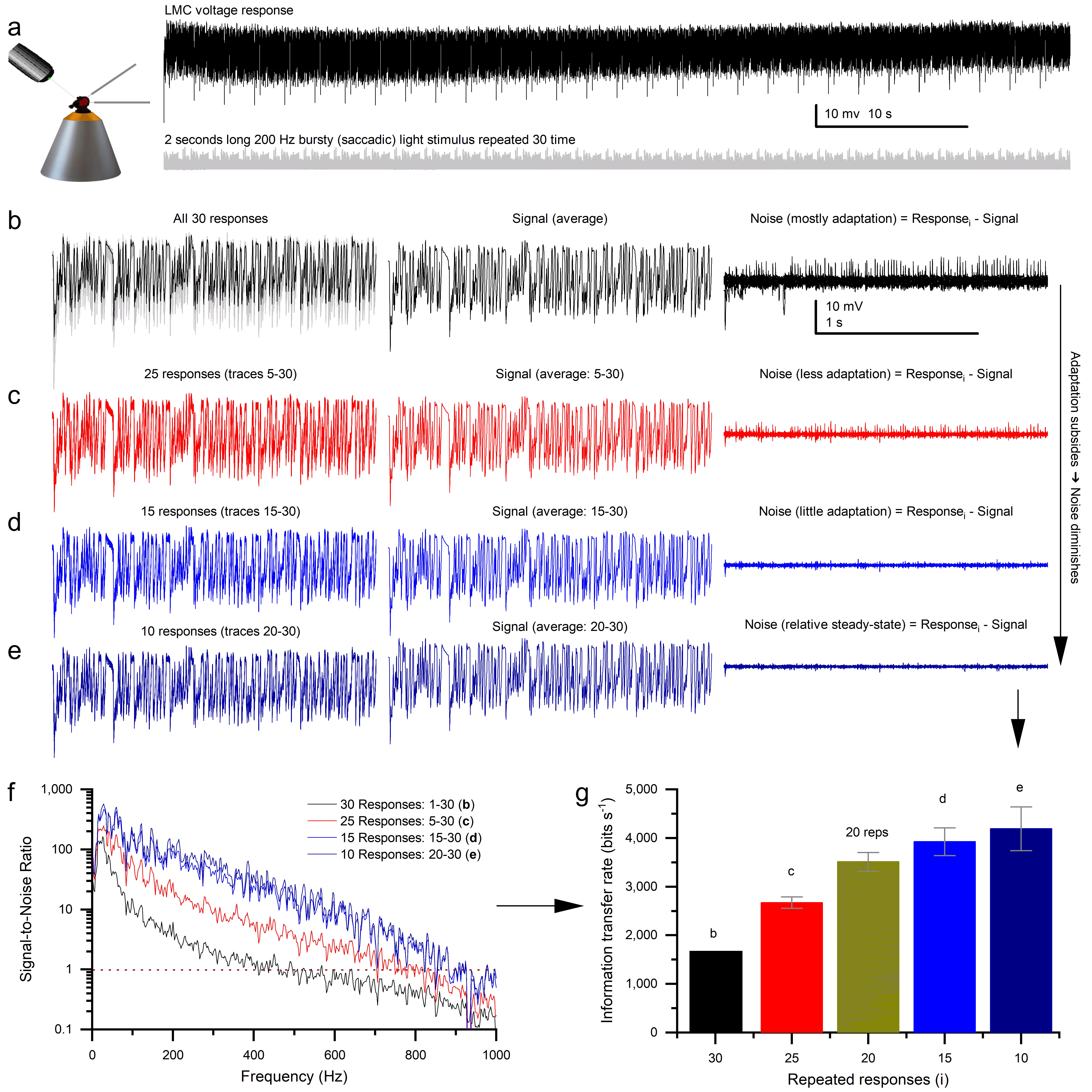

**Supplementary Fig. 9**. **Adaptive trends in repeated recordings bias Shannon information transfer rate estimates.**

Information transfer rates were calculated from repeated LMC responses to a fixed light stimulus using the Shannon formula applied to signal‑to‑noise spectra. Adaptive trends during the early repetitions elevate apparent noise, leading to systematic underestimation of the true synaptic information transfer rate.

(**a**) Minute‑long intracellular voltage recording (black) from an LMC responding to a 200‑Hz bursty contrast stimulus (grey) repeated 30 times (2‑s stimulus per repetition). The response shows a gradual adaptive drift across repetitions.

(**b**) All 30 voltage responses (left, superimposed), their mean (signal), and 30 noise traces derived by subtracting the mean from each individual response.

(**c**) Analysis after discarding the first five adaptive responses (25 remaining). Noise traces are flatter and show reduced variability compared with (b).

(**d**) Analysis after discarding the first 15 responses (15 remaining). Noise is further reduced.

(**e**) Analysis after discarding the first 20 responses (10 remaining). Responses closely align with the mean signal, and residual noise is minimal, indicating the system has reached a steady state.

(**f**) Signal‑to‑noise ratio (SNR) spectra from progressively trimmed datasets. Excluding early adaptive responses extends the frequency range of reliable encoding (SNR > 1) to ~1 kHz, indicating that the synapse can resolve events with ~0.5 ms precision.

(**g**) Corresponding Shannon information rate estimates exceed 4,000 bits/s once adaptive trends reach a relative steady state. Error bars indicate the estimation uncertainty due to data size, based on the number of repeated responses and assuming stationary data with Gaussian, additive signal and noise^35^.

These results demonstrate that adaptive trends contribute substantially to noise estimates in biological recordings; progressively excluding early responses reduces this bias and yields Shannon‑based information rates that more accurately reflect the true synaptic transfer capacity.

These analyses demonstrate that the estimates derived from 25 repetitions, after excluding the first five responses, are conservative rather than inflated; adaptive trends lead to underestimation rather than overestimation of synaptic information transfer capacity. Consequently, our reported values provide a robust lower bound on the true encoding precision of LMCs.

**Maximising Phasic Information Transfer**

Although both photoreceptors (**Supplementary Fig. 4d**) and LMCs (**Supplementary Fig. 7d**) encode saccadic contrast changes most efficiently, LMCs consistently exhibit higher information transfer rates (**Figure 3d**). This enhancement aligns with the data processing theorem^35,41,60^, which predicts that information throughput increases with the number of processed samples, provided other conditions remain constant^1,6,34,35^. Each LMC receives synaptic inputs from six photoreceptors - or seven in the frontal-dorsal acute zones known as the "love spots" of the male eye^39^ ( see **Section II**, **Supplementary Fig. 13**) - pooling these signals through morphodynamic neural superposition (**Figure 5**). This mechanism uses overcomplete tiling of moving receptive fields to enhance the eyes’ spatiotemporal resolution^4,52,61^, thereby improving the detection of moving objects well below the static resolution limit set by the interommatidial angles (${\Delta\phi}_{mean}={2.9}^{^{\circ}}$; see **Section II**) of the lens system.

During active vision, driven by the fly’s saccadic behaviours^62-64^, photoreceptors continuously adapt to bursty light contrast changes^1^, achieving exceptionally high signal-to-noise ratios (**Supplementary Fig. 4a**: 1,000-4,000 in bright conditions). This performance arises from stochastic refractory quantal sampling^1,6,27,34,40^ by ~54,000 microvilli per photoreceptor, which collectively absorb ~10^5^-10⁶ photons per second. The resulting flux of variable quantum bumps is integrated across the photoreceptor membrane into macroscopic voltage responses that effectively cancel residual noise^27,28^. Consequently, the synapses between photoreceptors and LMCs^8,9,16,27,49,65^ can respond reliably without delay to even the slightest changes in light contrast.

LMCs integrate ultrafast, synchronous histamine signals into transient, biphasic voltage responses. This collaborative synaptic encoding - driven by tonic feedforward-feedback interactions^1,7,8,11,16,54,65,66^ between photoreceptors and LMCs - efficiently expands and flattens the dynamic range of membrane potentials (**Supplementary Fig. 4f, 7f**), unlike the smaller, less structured responses to randomised control stimuli (Gaussian white noise, GWN). This process implements high-frequency jumping (described in the main paper and modelled in **Section IV**), redistributing information across a broader frequency spectrum (~1,000 Hz; cf. **Figure 2**). During synaptic transfer, the photoreceptor’s voltage range is clipped at both extremes because the activation range is narrower, squaring the signal and generating strong high-frequency components (>500 Hz) in the LMC. Consequently, signal bandwidth increases and transmission delays vanish (**Supplementary Fig. 1**), enabling ultrafast, reliable, and predictive visual processing.

**Adaptive Encoding and Signal Optimisation for High-Speed Inputs**

As saccadic stimulus speed increases, the distribution of LMC outputs first flattens (from 20 to 200 Hz bursts) before reaching a broad Gaussian shape at 500 Hz bursts (**Supplementary Fig. 7b**) - a hallmark of network-level adaptation^16,54^. In this adaptive framework, R1-R6 photoreceptors and LMCs, along with other cells of the morphodynamic neural superposition system^8,9,55^, dynamically balance synaptic feedforward and feedback loads to make full use of the LMCs’ output range under changing stimulus conditions.

Rapid adaptive signal flattening increases coding efficiency by approaching the ideal condition in which each output pattern (“symbol”) is used with equal probability^16,41^. Crucially, this dynamic encoding enhances the detection of underrepresented, novel, or salient features^16,27^ and requires continuous adjustment of synaptic gain - directly contradicting the static contrast normalisation model^53^, which assumes fixed gain and encodes contrast solely via maximum response amplitude.

However, LMC outputs consistently peak before the corresponding photoreceptor responses (e.g. **Supplementary Fig. 1** and **Figure 2d**), and both adapt over time - but in distinct ways (e.g. **Supplementary Fig. 10**). These observations confirm that the morphodynamic neural superposition system does not implement static contrast normalisation^53^, which would require a fixed synaptic (input-output) gain^8,16^.

**Benchmark in Neural Information Transfer**

LMCs sustain high information transfer rates across a broad range of stimuli (**Supplementary Fig. 10**), especially mid- and high-contrast 100 and 200 Hz bursts (**Supplementary Fig. 7c**, **d**). In contrast, photoreceptor sampling peaks only during the most intense 200 Hz bursts (**Supplementary Fig. 4c**, **d**). This superior performance - enabled by high-frequency jumping and minimal noise - yields a peak information transfer rate of ~4,100 bits per second, recorded from a single front-viewing LMC in a male housefly. To our knowledge, this is the highest rate reported for any neuron.

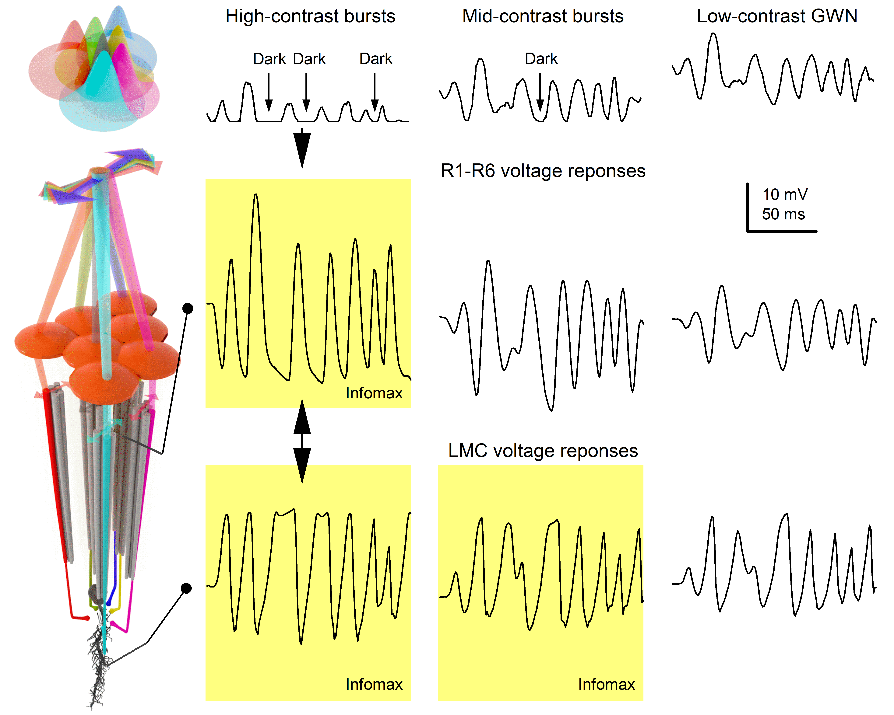
**Supplementary Fig. 10. The photoreceptor-LMC system performs dynamic contrast normalisation to maximise information flow (behavioural bursts drive efficient coding).**

**Left inset**: Similar to *Drosophila*^4,61^, the receptive fields of *Musca* R1-R6 photoreceptors from adjacent ommatidia - pooled into the same neural superposition system - form a rosette pattern, illustrated by transparent cones of different colours. Because the R1-R7/8 rhabdomere patterns are slightly rotated in neighbouring ommatidia, each receptive field shifts in distinct directions during microsaccades (indicated by double-headed arrows), which are triggered by local changes in photon flux. This spatially offset arrangement both minimises aliasing and enhances the detection of phasic (i.e. rapid) light contrast changes^4^.

The housefly visual system is adapted to efficiently encode bursty, high-speed temporal changes in the environment produced by rapid saccadic behaviours. The brief darker intervals between brighter moments allow microvilli - the photoreceptors’ photon-sampling units - to recover from refractoriness after prior photon absorptions, enabling significant changes in quantum bump rates^1,6^. As a result, both R1-R6 photoreceptors and LMCs make efficient use of their voltage ranges, with peak information transfer rates observed during bursty stimulation (yellow boxes). See also **Supplementary Fig. 25**, which schematically illustrates the core mechanistic principle of synaptic high-frequency jumping and its role in LMC signal normalisation

In contrast, Gaussian white noise (GWN) stimulation maintains photoreceptors at relatively steady brightness levels, causing many microvilli to remain refractory - thereby limiting maximal quantum bump rates and reducing LMC information throughput. Notably, LMC output is less contrast-specific than photoreceptor input, achieving similarly high information rates during both mid- and high-contrast bursts. This suggests dynamic synaptic contrast normalisation of signals routed to the brain.

As detailed in the **Methods** of the main paper, these values were obtained under restrictive experimental conditions that introduce instrumental noise and may cause some neuronal damage. Therefore, the actual encoding capacity of intact neurons during active vision in free flight is likely even higher.

**I.4. Differences in R1-R6 and LMC Signalling Performance and Sex-Based Variation**

LMCs (n = 6) consistently exhibit higher information transfer rates than photoreceptors (n = 20) when presented with identical stimuli (p = 0.0127). Supporting our initial hypothesis - that the housefly visual system and active behaviours are co-adapted to sample and process saccadic inputs with exceptional speed, accuracy, and minimal latency - both male and female flies responded most vigorously to rapid "saccadic" stimuli. However, male photoreceptors - and potentially LMCs - showed significantly higher information transfer rates (p = 0.0160; **Supplementary Fig. 7d**). This sex-related difference likely reflects specialised adaptations in the frontal-dorsal acute zones of male flies, known as “love spots"^24,32,67-69^, in which optical and neural architecture suggests enhanced visual resolution (**Section II**, **Supplementary Fig. 19**). These regions are absent in female flies.

Although our experiments did not intentionally target “love spots,” neuron selection was based solely on the quality of microelectrode recordings, independent of location. Due to the limited number of complete LMC recordings, we cannot statistically confirm sex-based differences in LMCs. However, the highest observed information transfer rate was recorded from a male LMC, supporting the plausibility of such distinctions.

During mating pursuits^64,70,71^, male flies execute rapid saccadic head and body movements - consistent with our findings and predictions for actively maintaining maximally efficient vision. This behaviour is likely further facilitated by specialised neural circuits in their "love spots." Intriguingly, in this region, central R7 photoreceptors - typically associated with medulla colour processing - project directly to LMCs^67,71-73^. While technically demanding, future studies could label individual LMC subtypes (L1-L3) to investigate their differential responses to saccadic stimuli. Notably, the largest L3 neurons are found specifically in the male “love spot”^74^, suggesting an additional anatomical adaptation for enhanced visual performance during fast pursuit behaviours.

**I.5.** **High-Frequency Jumping Reaches Highest Bandwidths during 50-200 Hz High-Contrast Bursts**

Natural visual behaviour in flies is dominated by brief, high-contrast intensity transients generated by rapid body, head, and retinal movements during saccades (e.g. **Figure 1**). The frequency content of the visual input experienced by individual photoreceptors can therefore vary widely, depending on the surrounding natural scene and the fly’s proximity to environmental structures. It is likely that flies adjust their flight speed in response to visual clutter, slowing down in crowded environments and speeding up in more open space, thereby dynamically shaping the temporal statistics of retinal input^1^.

To capture this wide range of possible stimulus patterns, we measured the signal-to-noise ratios (SNRs) of both the visual stimuli and the corresponding intracellular voltage responses of an exemplary R1-R6 photoreceptor and its postsynaptic LMC (**Supplementary Fig. 11a, b**). Recordings were obtained from cells exhibiting exceptional stability during high-contrast bursty stimulation with systematically varied cut-off frequencies, extending up to 750 Hz. As in other figures, effective bandwidth was defined conservatively as the frequency range over which *SNR*(*f*) > 1, ensuring that only reliably encoded signal components were considered. This approach enabled direct comparison of how stimulus bandwidth is transformed across the photoreceptor-LMC synapse under different temporal regimes.

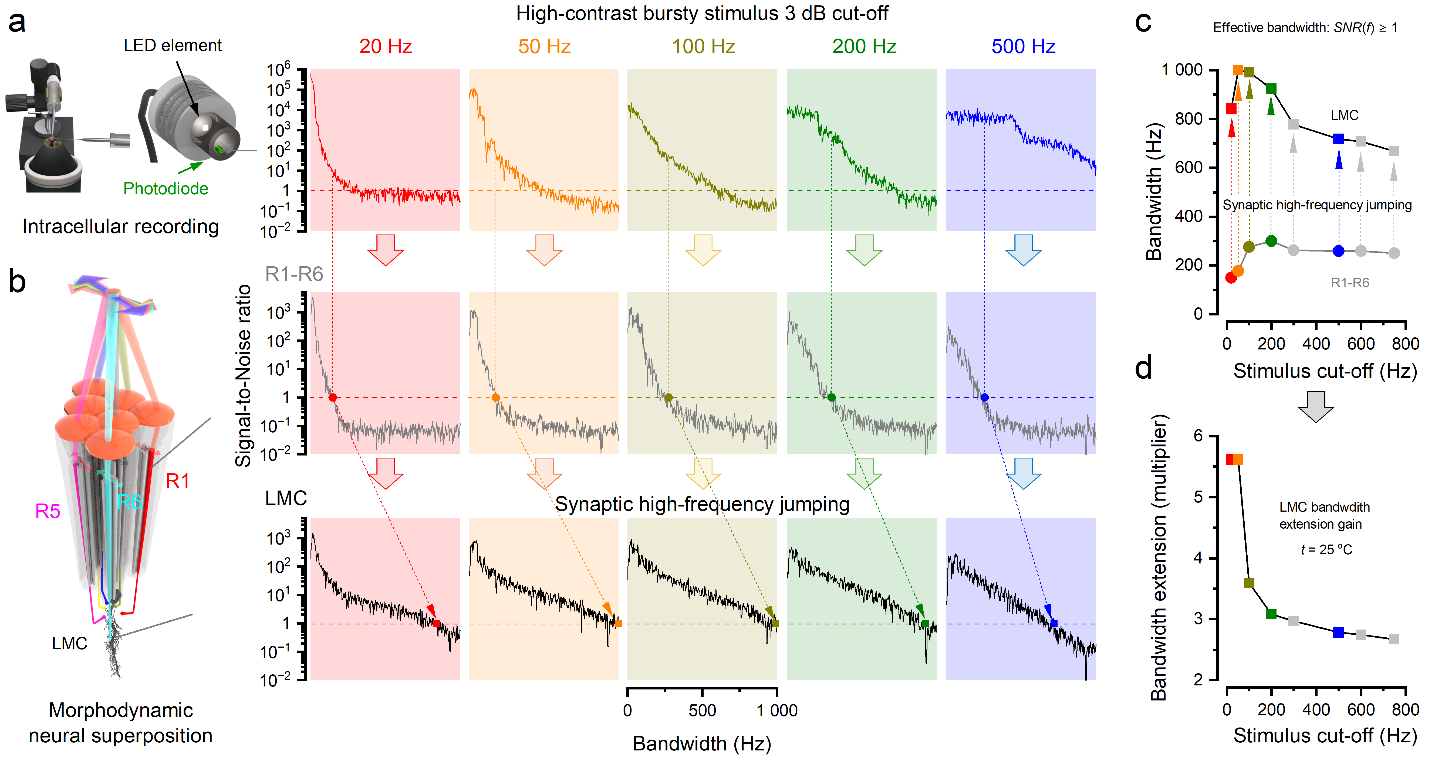

**Supplementary Fig. 11. Synaptic high-frequency jumping allocates LMC bandwidth most efficiently during high-contrast bursty (saccadic) stimulation in the 20-300 Hz range.**

(**a**) Signal-to-noise ratios (SNRs) of high-contrast bursty light stimuli with different 3-dB cut-off frequencies (20, 50, 100, 200, 300, 500, 600, and 750 Hz). Light output from the LED driver system was measured directly using a photodiode during intracellular recordings from R1-R6 photoreceptors and large monopolar cells (LMCs), sampled at 2 kHz. Dashed horizontal lines indicate the criterion for effective stimulus bandwidth, defined as frequencies for which *SNR*(*f*) > 1 (signal power exceeds noise).

(**b**) Effective bandwidths of an example R1-R6 photoreceptor and its postsynaptic LMC in response to 20, 50, 100, 200, and 500 Hz bursty stimulation. Plots show the SNR spectra of voltage responses; dashed horizontal lines again mark *SNR*(*f*) = 1, defining the frequency range over which neural responses reliably encode stimulus information. Owing to synaptic high-frequency jumping, the LMC’s effective bandwidth extends well beyond that of the presynaptic photoreceptor. This bandwidth extension is maximal for high-contrast bursty stimulation in the 50-200 Hz range.

(**c**) Summary of effective bandwidths for R1-R6 photoreceptors and LMCs as a function of stimulus cut-off frequency, illustrating the frequency-dependent emergence of synaptic high-frequency jumping.

(**d**) Photoreceptor-to-LMC bandwidth extension gain (ratio of LMC to R1-R6 effective bandwidth) for each bursty stimulus condition, showing that synaptic high-frequency jumping most efficiently enhances postsynaptic bandwidth under intermediate (20-300 Hz) high-contrast saccadic stimulation.

These measurements show that synaptic high-frequency jumping selectively extends LMC bandwidth beyond that of the presynaptic photoreceptor, with the largest relative gain occurring for bursty stimuli in the 20-300 Hz range (**Supplementary Fig. 11c, d**). Notably, for 50 Hz bursts, the efficient LMC bandwidth reached 1,000 Hz. At higher stimulus cut-off frequencies, the bandwidth extension diminishes. This reduction is consistent with increasingly frequent stimulus fluctuations providing fewer dark intervals to relieve microvillar refractoriness, thereby limiting changes in photoreceptor quantum bump rates. As a consequence, the macroscopic photoreceptor voltage response decreases and clips the synaptic operating range less effectively, reducing high-frequency synaptic jumping to LMCs. Thus, synaptic high-frequency jumping does not uniformly amplify bandwidth, but instead reallocates postsynaptic encoding capacity most efficiently to the temporal frequencies that dominate natural high-contrast saccadic inputs.

**I.6. Synaptic High-Frequency Jumping Increases with Light Intensity**

Natural visual input statistics change systematically with ambient light level. As illumination increases from twilight to daylight, photon flux rises, photoreceptor responses accelerate, and microvillar refractoriness increasingly shapes transduction dynamics. We therefore asked whether synaptic high-frequency jumping is gated by light intensity, and whether the enhanced postsynaptic bandwidth observed during bursty stimulation emerges progressively as illumination increases.

To address this, we analysed photoreceptor and LMC responses to identical high-frequency contrast stimuli across a wide range of light intensities, using both time-domain measures (**Supplementary Fig. 12**) and frequency-domain signal-to-noise analysis (**Supplementary Fig. 13**). In the time domain, increasing light intensity leads to larger and faster LMC voltage transients relative to those of the presynaptic photoreceptor, indicating enhanced transmission of rapid signal components. In the frequency domain, this effect is seen as a disproportionate expansion of LMC effective bandwidth with increasing light intensity, while photoreceptor bandwidth increases more gradually. Effective bandwidth was defined conservatively as the frequency range over which *SNR*(*f*) > 1, ensuring that only reliably encoded signal components were included.

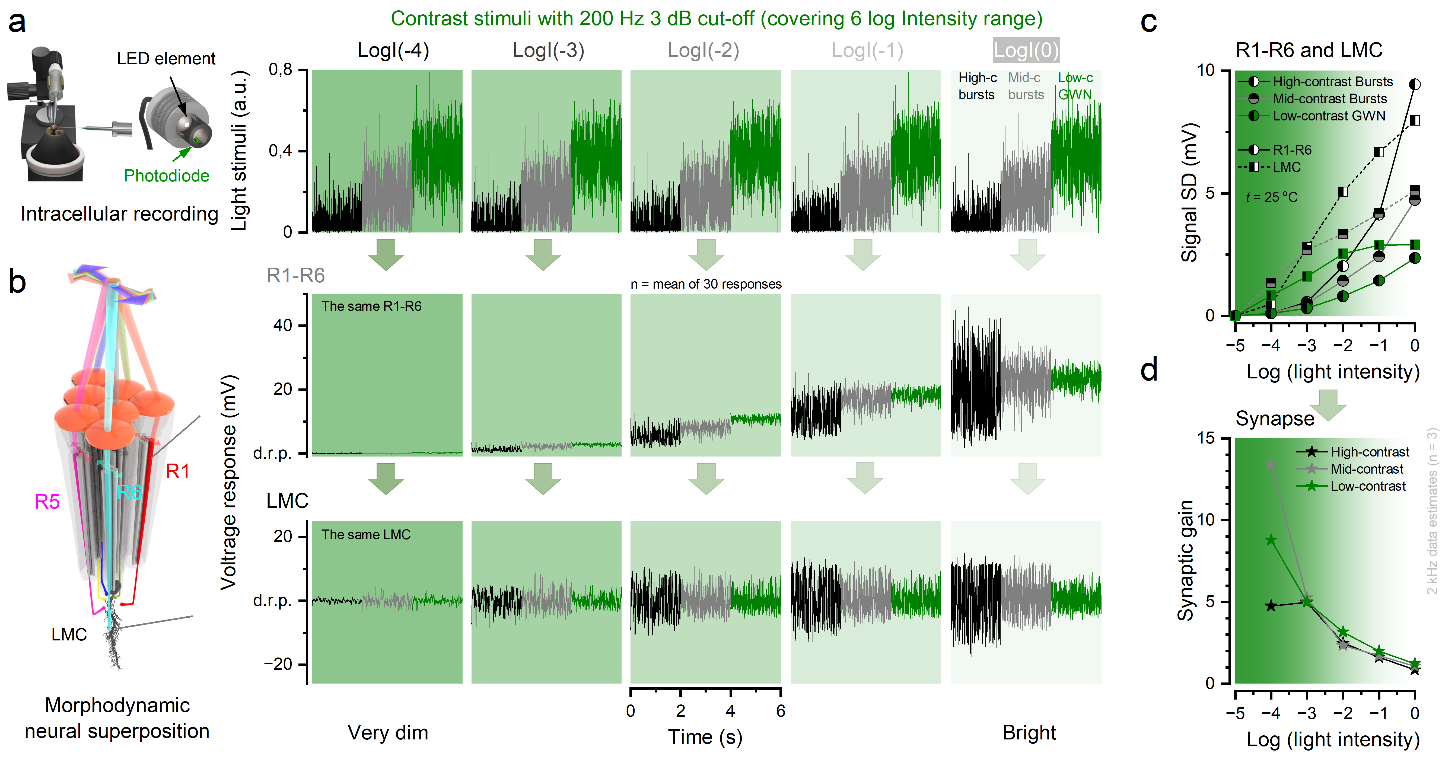

**Supplementary Fig. 12. High-frequency contrast bursts allocate photoreceptor and LMC voltage ranges most efficiently from twilight to daylight intensities.**

(**a**) Light output of three stimulus classes with a 200 Hz (3-dB) cut-off frequency: high-contrast bursty stimuli (black traces), mid-contrast bursty stimuli (grey traces), and low-contrast Gaussian white noise (GWN; green traces), measured directly using a photodiode. During intracellular recording experiments, stimulus intensity was attenuated over a million-fold range using a logarithmic series of neutral density filters (logI −5, −4, −3, −2, −1, and no filter = logI 0).

(**b**) Corresponding voltage responses of a single, highly stable R1-R6 photoreceptor (top) and its postsynaptic large monopolar cell (LMC; bottom) to the three stimulus classes across light intensities spanning very dim conditions [logI(−4)] to daylight [logI(0)]. At the lowest intensities [logI(−5) to logI(−3)], photoreceptor responses are not limited by microvillar refractoriness and effectively integrate all absorbed photons. Under these conditions, mid-contrast bursty and GWN stimuli evoke larger depolarisations than high-contrast bursty stimuli, which contain only ~9.43% of the photons present in GWN (see panel **a**). As light intensity increases, the dark intervals within bursty stimuli increasingly relieve microvillar refractoriness, thereby accentuating rapid intensity changes. Consequently, photoreceptor response variance (standard deviation) becomes largest for high-contrast bursty stimulation from twilight [logI(−3)] through daylight [logI(0)]. From logI(−3) onwards, LMC voltage responses are consistently largest for high-contrast bursty stimuli.

(**c**) Standard deviation of photoreceptor and LMC voltage responses as a function of light intensity, showing increased response variance with increasing illumination.

(**d**) Synaptic gain (photoreceptor-to-LMC transfer) decreases with increasing light intensity, reflecting compression of the synaptic operating range at higher photon flux.

The example photoreceptor and LMC shown were selected from the full intracellular recording datasets presented in **Supplementary Fig. 3** and **6**, respectively.

Together, these analyses show that synaptic high-frequency jumping is strongly light-intensity dependent (**Supplementary Fig. 13**). At very low intensities, both photoreceptor and LMC bandwidths are limited, and synaptic bandwidth extension is minimal. From twilight to daylight intensities, however, synaptic high-frequency jumping becomes increasingly pronounced, selectively reallocating postsynaptic encoding capacity toward higher frequencies. This light-dependent gating is consistent with bursty stimuli, in which intermittent dark periods relieve microvillar refractoriness, enabling larger changes in quantum bump rates and thus larger macroscopic voltage responses. As photoreceptor voltage increases, it exceeds the synaptic operating range and clips, leading to synaptic high-frequency jumping that progressively extends effective LMC bandwidth. Thus, synaptic high-frequency jumping is not a fixed property of the photoreceptor-LMC synapse, but an adaptive mechanism that becomes most effective under daylight conditions, when fast visual processing is behaviourally most relevant.

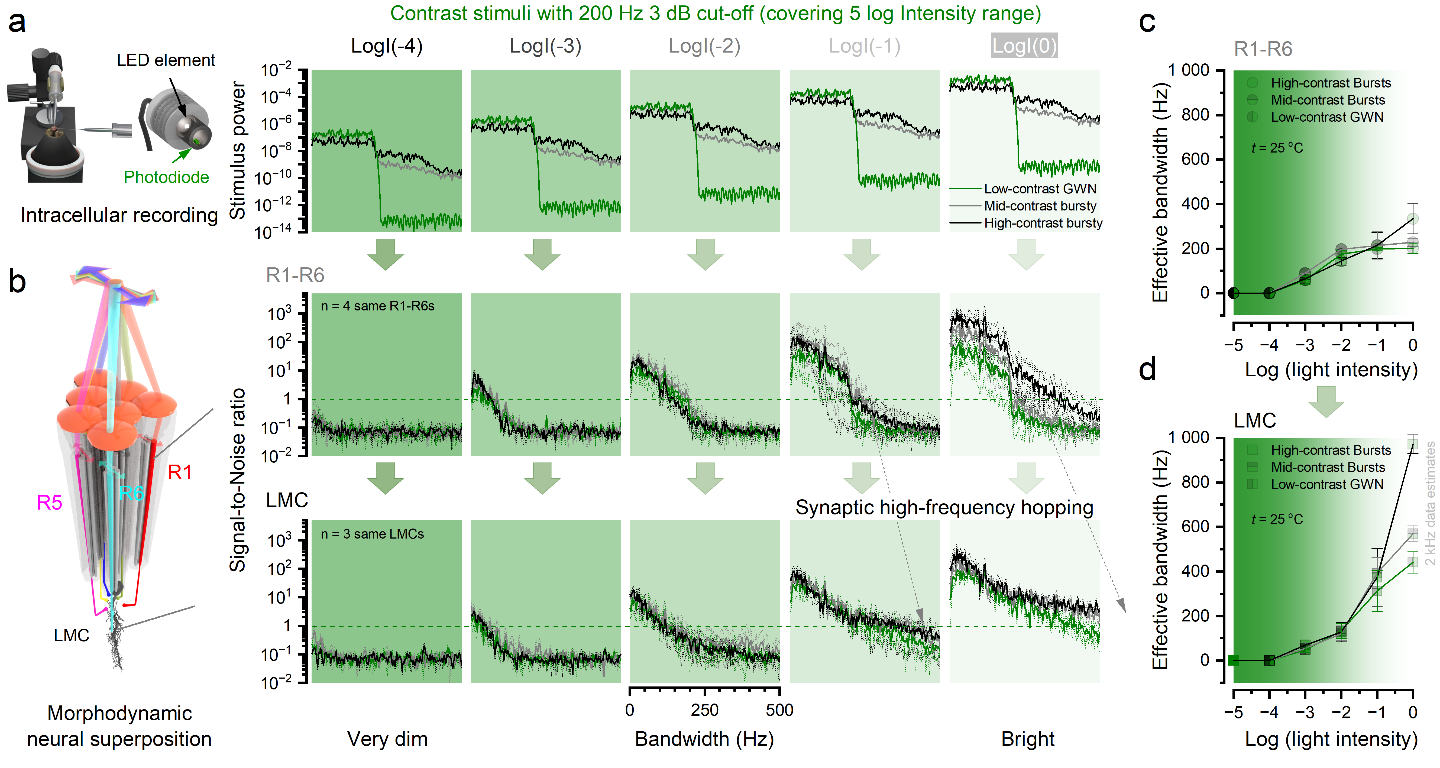

**Supplementary Fig. 13. Synaptic high-frequency jumping is light-intensity dependent in the frequency domain and is most effective at daylight intensities.**

(a) Power spectra of contrast stimuli with a fixed 200 Hz (3-dB) cut-off frequency, measured directly from the LED output using a photodiode. Three stimulus classes were used: high-contrast bursty stimuli (black), mid-contrast bursty stimuli (grey), and low-contrast Gaussian white noise (GWN; green). Stimuli were presented across a five-log-unit light intensity range (logI −5 to logI 0), spanning very dim conditions to bright daylight, using neutral density filters.

(b) Signal-to-noise ratios (SNRs) of intracellular voltage responses from R1-R6 photoreceptors (top) and their postsynaptic large monopolar cells (LMCs; bottom) to the same stimuli and light intensity range shown in (a). Dashed horizontal lines indicate *SNR*(*f*) = 1, defining the effective bandwidth over which neural responses reliably encode stimulus information. At very low light intensities, effective bandwidths are limited by low numbers of absorbed photons in both photoreceptors and LMCs. With increasing light intensity, photoreceptor bandwidth increases modestly, whereas LMC bandwidth increases disproportionately, revealing synaptic high-frequency jumping that selectively extends postsynaptic encoding at higher frequencies.

(c) Effective bandwidths of R1-R6 photoreceptors as a function of light intensity (logI), for the three stimulus classes, showing gradual bandwidth expansion with increasing photon flux.

(d) Effective bandwidths of LMCs across the same light intensity range. LMC bandwidth increases steeply from twilight to daylight intensities and exceeds photoreceptor bandwidth most strongly for high-contrast bursty stimuli, demonstrating light-intensity-dependent synaptic high-frequency jumping in the frequency domain.

Together with the time-domain analysis in **Supplementary Fig. 13**, these results show that synaptic high-frequency jumping is dynamically gated by light intensity and operates most efficiently under daylight conditions, when photoreceptor refractoriness and synaptic dynamics favour rapid, high-frequency signal transmission. These photoreceptor and LMC responses were recorded from some of the same cells presented in **Supplementary Fig. 3** and **6**, respectively.

**I.7. Light-Adapted R1-R6 Photoreceptors Exhibit Hyperacute Responses to Narrowing 2D Gratings**

To investigate the spatial resolution limits of *Musca* R1-R6 photoreceptors, we performed intracellular recordings while presenting a two-dimensional square-wave grating with progressively narrowing spatial features. Recordings were conducted at room temperature (~22-24 °C), without the Peltier temperature controller, to reduce contamination from external electronic noise (e.g., 50 Hz mains interference and harmonics).

**Grating Stimulus and Its Presentation**

A digital light projector (EKB DLP® LIGHTCRAFTER™ E4500 MKII™, EKB Technologies Ltd., Israel) was used to present the grating stimuli (**Supplementary Fig. 14a**). The projector was mounted on a rotatable Cardan-arm system to allow precise stimulus alignment. It featured three independent LED sources - ultraviolet (385 nm), blue (460 nm), and green (520 nm) - and projected its image onto a back-projection diffuser screen via a set of three close-up lenses (ZEIKOS, Japan; 65 mm, +10, +5, +1). The system operated at its native resolution of 912 × 1140 pixels with a refresh rate of 360 Hz. To minimise electrical noise during recordings, the projector was enclosed in a grounded copper mesh fabric.

The narrowing grating stimulus (**Supplementary Fig. 14b**) was programmed in MATLAB (MathWorks, USA) and defined by two variable and three constant parameters: grating speed ($s$), motion direction ($\theta$), initial angular distance between adjacent bars (ie. spatial wavelength; $\lambda_{0}$ = 5°), final spatial wavelength ($\lambda_{0}$ = 0.33°), and stimulus duration ($D$ = 40 s). The angular distance between adjacent bars varied over time according to the following equation:

$\lambda\left( t+dt \right)=\left( \frac{\lambda_{1}}{\lambda_{0}} \right)^{1/D}=\lambda_{0}\left( \frac{\lambda_{1}}{\lambda_{0}} \right)^{t/D}$	(4)

The stimulus was presented for 40 s, followed by a blank (black) screen. Intracellular voltage responses from R1-R6 photoreceptors were recorded for 45 s, with acquisition synchronised to stimulus onset.

Five stimulus speeds (6°/s, 20°/s, 40°/s, 60°/s, and 120°/s) were used to assess the effect of motion speed on spatial resolvability. Additionally, four cardinal motion directions were tested to evaluate directional sensitivity. These gratings were presented in randomised order. Additionally, each recording was repeated once more immediately after the first one had run, totalling two repeats per grating. This resulted in a total of 40 distinct recordings - approximately 30 minutes of recording time per photoreceptor.

The same stimulus projection system was used to generate moving two-dot stimulation with occlusion (**Figure 6**; **Supplementary Fig. 18**) to study the spatiotemporal acuity of LMC voltage responses.

***Musca* Preparation and Intracellular Recordings**

Adult houseflies were cold-anesthetised on ice, after which their wings and legs were removed to ensure stable mounting. Flies were then immobilised by applying low-melting-point beeswax to the head capsule and thorax within custom conical holders, consisting of a brass inner tube encased in a plastic cone. A small window (~6-10 ommatidia wide) was cut in the dorsal region of the left compound eye’s cornea to allow microelectrode access. To prevent desiccation during recordings, the opening was sealed with Vaseline.

The fly was positioned in the recording setup with its left eye facing the experimenter (**Supplementary Fig. 14a**). Sharp, filamented borosilicate glass microelectrodes (Sutter Instruments, USA; outer diameter 1.0 mm, inner diameter 0.5 mm) were pulled using a horizontal laser micropipette puller (P-2000, Sutter Instruments, USA), producing tip resistances of 100-250 MΩ. Reference electrodes were pulled separately using a dedicated program to produce blunt-tipped microelectrodes.

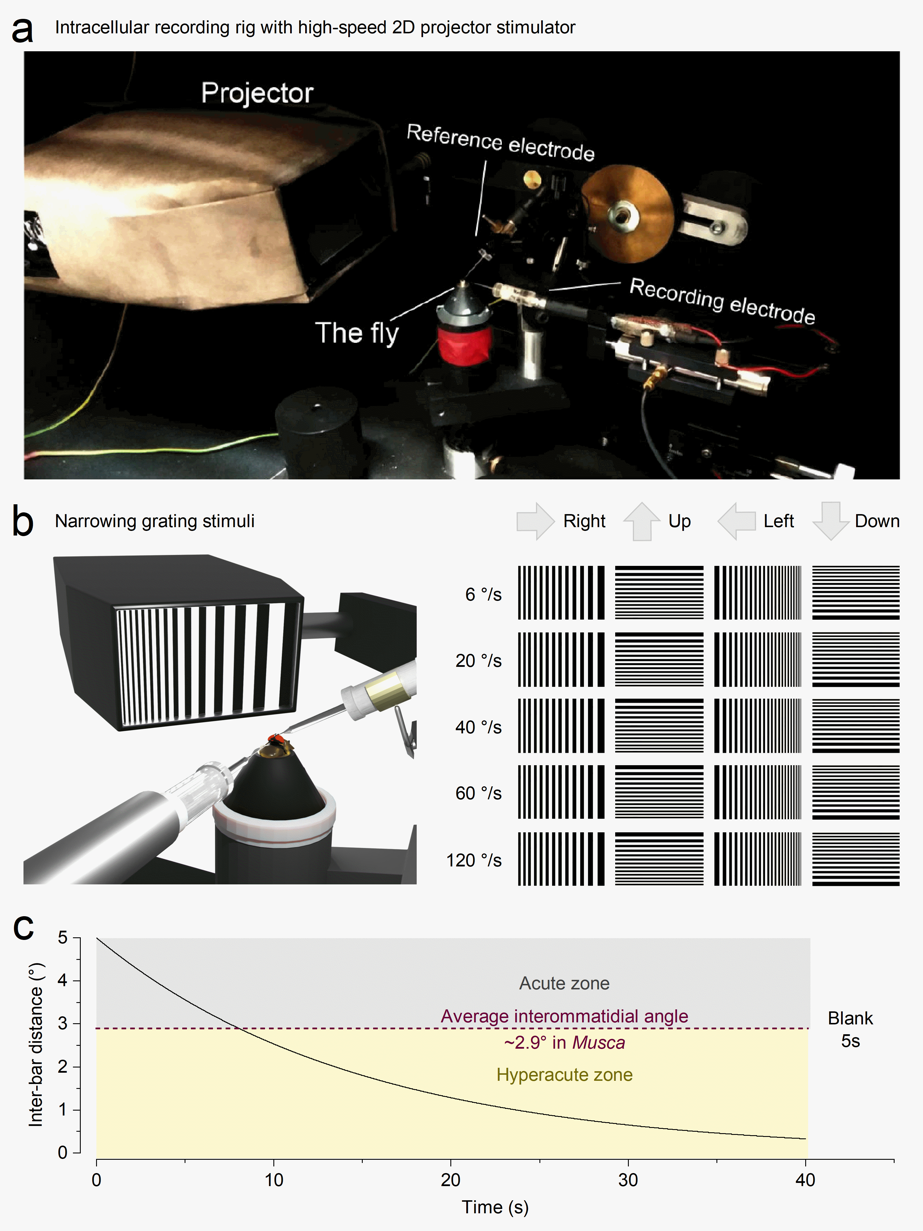
**Supplementary Fig. 14. Assessing *Musca* R1-R6 photoreceptors’ spatiotemporal resolution using intracellular recordings to moving and narrowing grating stimuli.**

(**a**) Experimental setup. The fly was mounted in a custom conical holder, and intracellular voltage recordings were made from R1-R6 photoreceptors using conventional sharp microelectrodes. A reference electrode was inserted through the ocellus in the head capsule. The fly and electrodes were positioned so that each recorded photoreceptor directly faced the centre of a back-projection screen, onto which stimuli were presented by a high-speed digital light projector. The screen was rotated using a Cardan Arm to match the receptive field’s optical axis.

(**b**) Stimulus design. A narrowing vertical grating was presented in four cardinal directions (Right, Left, Up, Down) and at five angular velocities (6°, 20°, 40°, 60°, and 120°/s). Each stimulus consisted of a pattern of black-and-white bars that moved steadily in the selected direction while the inter-bar spacing decreased over time.

(**c**) Stimulus profile. The angular separation between grating bars decreased exponentially throughout the stimulus period. The wine dashed line indicates *Musca*’s average interommatidial angle (~2.9°), which defines the theoretical optical resolution limit of the compound eye. The yellow-shaded region marks the “hyperacute zone,” where the stimulus features are finer than the sampling grid of ommatidia. By examining the voltage responses during this narrowing stimulus, we assessed how finely tuned R1-R6 photoreceptors are to dynamic, high-resolution spatiotemporal patterns.

Immediately prior to recordings, electrodes were backfilled as follows: recording electrodes were filled with 3 M KCl for photoreceptor recordings, and with 3 M potassium acetate containing 0.5 mM KCl for LMC recordings to maintain the chloride equilibrium potential. Reference electrodes, used for both photoreceptor and LMC recordings, were filled with fly Ringer solution (120 mM NaCl, 5 mM KCl, 5 mM TES, 1.5 mM CaCl₂, 4 mM MgCl₂, and 30 mM sucrose)^35,75^.

The blunt reference electrode was gently inserted into the ocelli (**Supplementary Fig. 14a**, **b**) using a three-axis micromanipulator (Narishige U-3C), while the recording electrode was advanced through the Vaseline-sealed corneal window using a remote-controlled micromanipulator (Märzhäuser DC-3K and PM 10 with two remote controllers). This process was guided visually using a stereomicroscope (Nikon SMZ645 with 30× eyepieces) illuminated by a halogen cold light source (Schott, with single gooseneck light guide).

The electrophysiology rig was mounted on a vibration-isolation table (Melles Griot frame with central-aperture breadboard - the aperture provided an entrance for a Peltier system^31^) and enclosed in a Faraday cage, equipped with two grounded, front-facing curtains for additional electrical shielding. Intracellular voltage responses were amplified using an npi SEC-10L amplifier (NPI Electronics, Germany), low-pass filtered at 500 Hz with custom-built analogue filters (Mick Swann, UK), and digitised at 1 kHz using a 12-bit A/D converter (NI PCI-MIO-16E-4, National Instruments, USA) connected via BNC interface blocks (NI BNC 2110).

The recording system - including the PC (running Windows 7), monitor, amplifier, and filters - was housed in a grounded instrument cabinet rack (Schroff 38U, nVent Schroff, Germany).

Visual stimuli were delivered using the open-source Psychophysics Toolbox (http://psychtoolbox.org) running in MATLAB (MathWorks, USA). Data acquisition was managed by Biosyst^35,75^ (M. Juusola, 1999-2020), interfaced with National Instruments data acquisition boards (Austin, TX) via the MATDAQ package (H.P.C. Robinson, 1997-2005). Biosyst was synchronised with the projector control software (LightCrafter), and a custom Biosyst extension was developed to enable automated batch recordings.

R1-R6 photoreceptors were identified by their characteristic voltage waveforms in response to light flashes, and by the electrode’s penetration angle within the retina. Only stable, high-quality cells were recorded. In darkness, resting membrane potentials were typically below -60 mV, and saturating bright pulses (100 ms) evoked responses of ≥45 mV. The projector screen was aligned to the photoreceptor’s optical axis by adjusting the Cardan-arm system while presenting a flashing dot stimulus. Once aligned, the recording site’s horizontal position - classified as front, mid-front, middle, mid-back, or back - was documented to provide a coarse localisation of the cell within the eye.

Due to the relatively long stimulus duration (40 s), short dark period (5 s), and brief inter-stimulus intervals (2-3 s), photoreceptors remained predominantly light-adapted throughout the experiment. Additionally, the projector’s blank (black) screen emitted ~0.1% of full-intensity light due to the device’s 1,000:1 contrast ratio, further contributing to sustained light adaptation.

**Algorithmic Resolvability Analysis of the Narrowing Grating Responses**

Algorithmic resolvability analysis of the narrowing grating data was implemented in Python 3.12 (Python Software Foundation, Netherlands). The raw photoreceptor voltage responses were used without preprocessing, as noise levels were minimal. Furthermore, the subsequent sequence similarity analysis benefits from the presence of small random fluctuations (0.1 - 0.5 mV) inherent in the recordings.

The primary parameter extracted from the narrowing grating responses was the smallest angular spacing that photoreceptors could reliably resolve. In theory, this finest resolved angle could be directly determined from the time axis (**Supplementary Fig. 14c**), since the grating’s angular wavelength (i.e., the inter-bar angular distance as perceived by the fly) at time $t$ follows **Equation 1**. However, in practice, the transition between resolved and unresolved stimuli occurs gradually over several seconds (**Supplementary Fig. 15a**), making precise determination challenging.

We developed an unbiased analysis algorithm to determine the final resolved grating angle that a photoreceptor could follow. The algorithm consists of three steps: (1) calculation of the sequence similarity of the photoreceptor’s voltage response over time, (2) fitting a horizontal noise baseline to the tail of the sequence similarity curve, and (3) identifying the first intersection point between the sequence similarity curve and the noise baseline. A point $s_{i}$ on the sequence similarity curve was calculated as:

$s_{i}=\frac{\sum_{i=a_{i}}^{b_{i}} \left| \left( v_{i}-v_{i+1} \right) \right|}{b_{i}-a_{i}}$	(5)

Here $a_{1}$ and $b_{i}$ denote the start and end indices of a sliding window centred at a time point $t_{i}$, while $v_{i}$ and $v_{i+1}$ represent the photoreceptor membrane voltages at time points $t_{i}$ and $t_{i+1}$. The window length ($b_{i}-a_{i}$) can be adjusted, balancing the trade-off between noise and accuracy: shorter windows (<100 samples) produce noisy sequence similarity curves, leading to overly pessimistic resolvability estimates, whereas excessively long windows (>1,000 samples) overly smooth the curve, leading to overly optimistic resolvability estimates (**Supplementary Fig. 15b**). We selected a window length $b_{i}-a_{i}=504$ for optimal performance (**Supplementary Fig. 15c** and **d**). Near the start $t_{i}$ and end $t_{45,000}$ of the recording, the window is truncated at the data boundaries to avoid exceeding available data points.

A horizontal noise baseline was fitted using data from the last quarter of the recording, excluding the final 5 seconds during which the projector displayed a blank screen. This exclusion was necessary because some photoreceptors showed decreased sequence similarity during the blank period, which would have biased the baseline fit and resulted in overly optimistic estimates of the final resolved angles. The baseline was computed as the mean sequence similarity between 33.75 seconds and 40 seconds. Importantly, this mean value is largely insensitive to the choice of window length, in contrast to metrics such as the minimum, maximum, or percentiles, which vary depending on the window size.

Finally, the last step of the algorithm was to iterate through data points to identify the moment when the sequence similarity curve first fell below the noise baseline. This straightforward approach worked well for all recordings except those with the slowest 6°/s grating. In these cases, an early, abrupt drop often occurred near the start, followed by the sequence similarity remaining consistently above the noise threshold. To avoid making erroneous estimates with these slow stimulus instances, the intersection point (i.e. the finest resolved angle) was constrained to occur only after 9 seconds. All recordings were subsequently checked to ensure that this constraint did not bias the results.

As shown in **Supplementary Fig. 15c**, this last step - identifying the intersection point - is sensitive to the chosen sequence similarity window length. An alternative approach involves fitting a line to the downward trend of the sequence similarity curve and determining its intersection with the noise baseline. While this method may appear attractive, it introduces its own trade-offs: most notably, the selection of fitting points has a strong influence on the outcome, particularly when the similarity curve declines gradually rather than sharply or when the decline is non-linear.

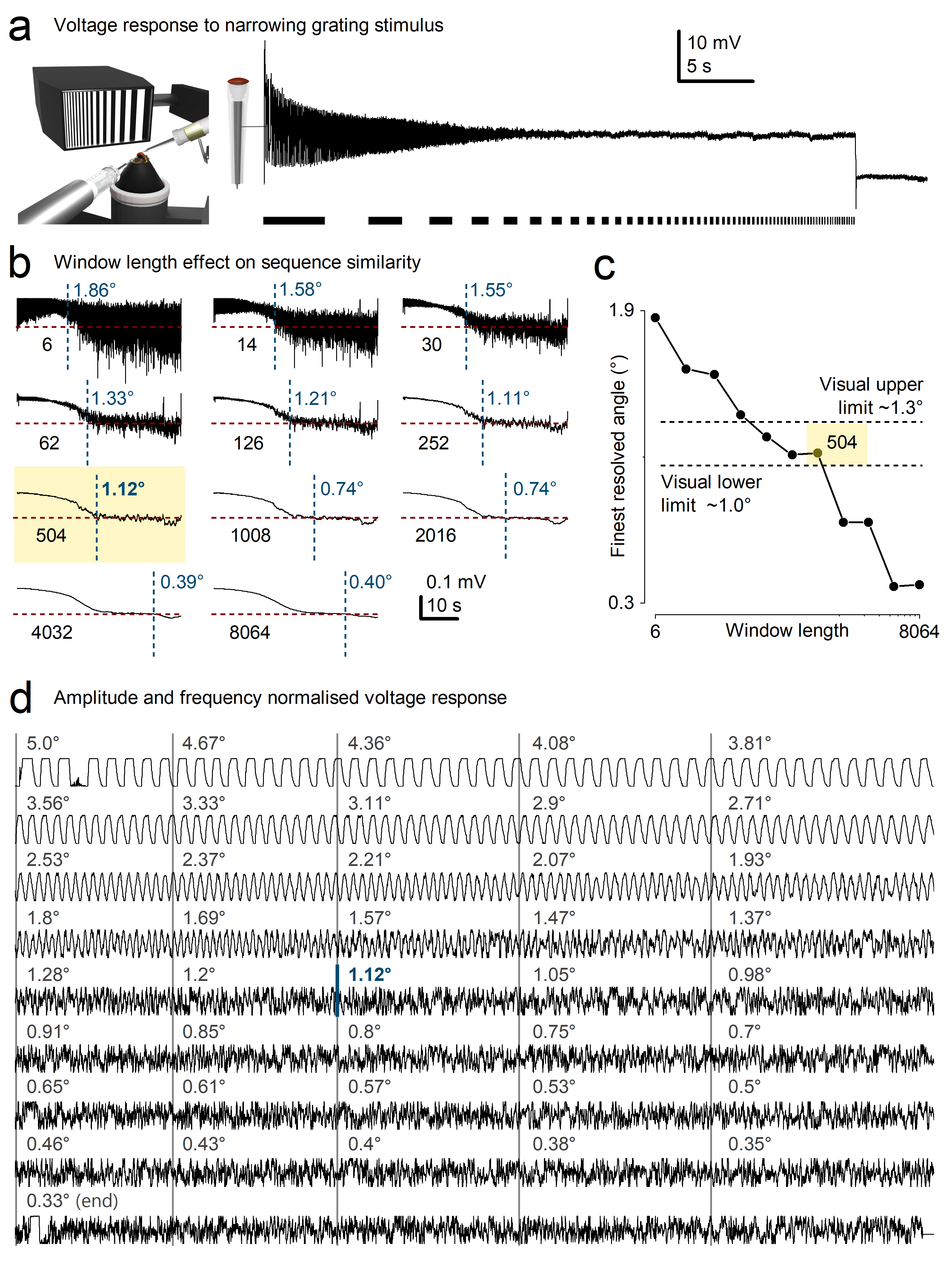
**Supplementary Fig. 15. Estimating the finest resolved angle from photoreceptor voltage responses using sequence similarity and stimulus narrowing dynamics.**

(**a**) In vivo intracellular voltage recording from a single *Musca* R1-R6 photoreceptor in response to a narrowing grating stimulus (black-and-white bar pattern shown below). The stimulus was presented along the cell’s optical axis, with spatial frequency increasing smoothly over time as the inter-bar distance decreased exponentially.

(**b**) Effect of window length on sequence similarity analysis. Voltage recordings were analysed using 11 different sliding window lengths (sample sizes indicated). Short windows produce noisy similarity curves, while excessively long windows smooth out important signal features. An optimal balance is required to accurately identify the transition point where the photoreceptor no longer tracks the stimulus, here resolved as 1.12° with a window size of 504 samples (highlighted in yellow).

(**c**) Plot of the finest resolved angle (°) versus window length. The selected window length (504 samples) lies within a robust range (visual resolution limit ~1.0°-1.3°) based on both algorithmic output and manual inspection of the traces shown in panel (**d**).

(**d**) Amplitude- and frequency-normalised voltage responses from the recording in (**a**), aligned to grating angles at each timepoint. As the inter-bar angle decreases from 5.0° to 0.33°, clear, periodic voltage modulations become increasingly attenuated and noisy. Visual inspection confirms that resolvability begins to degrade below ~1.12°, validating the algorithm’s output in (**b**-**c**).

**Uniform Presentation of the Narrowing Grating Responses**

The narrowing grating stimulus elicits photoreceptor responses with progressively decreasing amplitude and increasing temporal frequency, making it challenging to visually identify the final resolved angle directly from the raw membrane voltage traces. To address this, we developed a visualisation method that normalises response amplitude and applies a time-axis transformation to render the increasing stimulus frequency approximately constant. This approach facilitates intuitive comparison across conditions and enables visual validation of the algorithmically determined resolution limits.

To first amplitude normalise the responses, we selected a simple window-based algorithm, where the normalised voltage $u_{i}$ at a time point follows the equation:

$u_{i}=\frac{v_{i}-\underset{j=i}{\overset{i+w}{\min}}v_{j}}{\underset{j=i}{\overset{i+w}{\max}}v_{j}-\underset{j=i}{\overset{i+w}{\min}}v_{j}}$	(6)

Here, $v_{i}$ is the recorded membrane voltage at a time $t_{i}$, and $w=50$ denotes the window length, which can be freely selected. However, its choice involves trade-offs: excessively long windows result in poor normalisation, as they tend to capture low-frequency changes in membrane voltage, whereas overly short windows lead to noisy, jittery normalised signals. When appropriately chosen, the amplitude-normalised responses remain visually similar to the original data, although the waveform is not guaranteed to remain free from distortion by this simple algorithm.

The second normalisation step involved transforming the time axis. Because the grating stimulus angular size follows **Equation 1**, a simple iterative expression can be used to calculate the angular size $\lambda_{i}$ adjusted pseudo time points $\tau_{i}$ such that increasing stimulus frequency appears approximately constant across the new axis, facilitating uniform visual representation

$\tau_{i}=\frac{1}{t_{45,000}}\left( \frac{1}{\lambda_{i}}+\tau_{i-1} \right)$, $\tau_{0}=0$	(7)

Here $t_{45,000}=45 s$ denotes the final time point of the recording. Although the grating wavelength $\lambda_{i}$ is undefined beyond 40 seconds - when the projector transitions to a blank screen - pseudo-time values can still be extrapolated for the final 5 seconds to preserve continuity in the transformation. The resulting R1-R6 photoreceptor voltage trace, now both amplitude- and frequency-normalised, enables visual assessment of the final resolved grating angle with improved clarity (**Supplementary Fig. 15d**).

***Musca* R1-R6 Photoreceptors Resolve Hyperacute Details**

The housefly compound eyes have a sub-spherical architecture containing approximately 3,400-3,500 ommatidia per eye^76^, resulting in an average interommatidial angle of 2.9° (see **Section II**). If photoreceptors were static - as assumed in classical models of compound eye optics^43^ - this angular separation would impose a hard limit on spatial resolution, implying that houseflies could not resolve visual details finer than this sampling grid.

However, our experiments show that *Musca* R1-R6 photoreceptors can resolve hyperacute spatial details, well below this optical limit. Using a narrowing grating stimulus aligned with the photoreceptor’s optical axis (**Supplementary Fig. 16a**), we recorded intracellular voltage responses while systematically decreasing the grating bar width over time. As expected, response amplitude declined and frequency increased with spatial frequency (**Supplementary Fig. 16b**), yet robust periodic voltage modulations persisted even when bar widths dropped below 1° - a range traditionally considered unresolvable by compound eye optics.

To quantify the photoreceptors’ functional resolution limit, we applied a sequence similarity analysis to assess trial-to-trial reproducibility of voltage responses (**Supplementary Fig. 16c**). This revealed consistent signal structure down to 0.94°, beyond which responses dropped to the noise floor. Example traces (**Supplementary Fig. 15d**) show that although responses degrade with increasing spatial frequency, fine structure remains visible well below the interommatidial angle. These results highlight the photoreceptor’s capacity to extract features far finer than the eye’s static pixelation limit - consistent with predictions from morphodynamic information sampling theory^27^ and earlier findings from *Drosophila*^1,4^

Across the population, most photoreceptors resolved grating features between ~1° and 2° (**Supplementary Fig. 16e**-i), with resolution inversely related to recording noise (**e**-ii). Remarkably, resolution was robust across a wide range of stimulus speeds (**e**-iii), and showed modest regional variation (**e**-iv), with frontally located photoreceptors slightly outperforming those from the rear. Male photoreceptors (**e**-v) tended to outperform females, consistent with the anatomical specialisation of the male “love spot” (see **Section II**, **Supplementary Fig. 19**).

Together, these findings demonstrate that housefly photoreceptors exploit temporal sampling strategies - likely involving microsaccades and refractory quantal sampling^1^ - to resolve visual details far beyond what the eye’s geometric layout alone would suggest, supporting a functionally hyperacute, dynamic vision system.

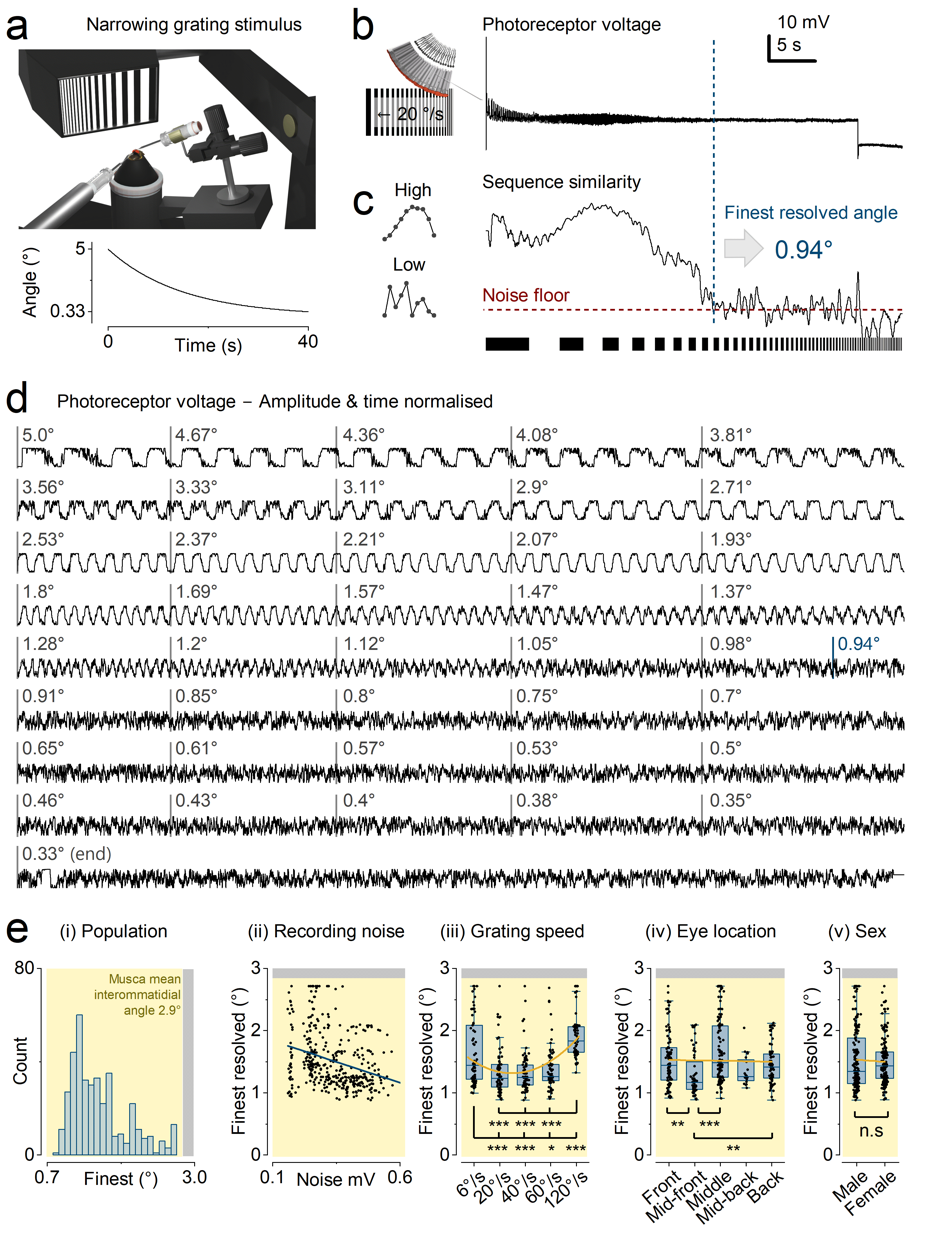
**Supplementary Fig. 16**. **R1-R6 photoreceptors in the *Musca* eye resolve hyperacute narrowing grating stimuli.**

(**a**) Schematic of the experimental setup. A sharp microelectrode is inserted into the dorsal region of the left compound eye to record from R1-R6 photoreceptors, while a blunt reference electrode is placed in the ocelli. Narrowing 2D grating stimuli are presented to the fly via a high-speed digital light projector, aligned with the photoreceptor’s optical axis. The lower plot shows how the angular width of the grating bars decreases exponentially over time.

(**b**) Example intracellular voltage response of a photoreceptor to a horizontally narrowing grating moving at 20°/s. As the bar width decreases, the response amplitude drops and the temporal frequency increases.

(**c**) Sequence similarity analysis for the response shown in panel (**b**). The curve shows a decline in similarity as spatial frequency increases, crossing the noise floor at the finest reliably resolved angle of 0.94° (vertical dashed line).

(**d**) Time- and amplitude-normalised voltage traces from a representative recording across a range of grating bar widths (from 5.0° to 0.33°). As the grating spatial frequency increases, the photoreceptor responses gradually degrade until they become indistinguishable from the noise recorded during blank screen presentation.

(**e**) Summary of photoreceptor resolution across multiple cells and conditions.

(i) Histogram showing the distribution of finest resolved angles across the photoreceptor population (n = 19); the average interommatidial angle (2.9°) is marked in yellow.

(ii) A negative correlation between recording noise and resolution: While the cells with the finest resolution also exhibited the lowest noise, an unexpected trend emerged - beyond this group, higher recording noise was paradoxically associated with finer resolved gratings.

(iii) Photoreceptor resolution as a function of stimulus speed; whilst resolution remains robust across all the tested speeds from 6°/s to 120°/s, the responses showed the highest resolvability with mid velocities: 20-60°/s.

(iv) Resolution grouped by recording location on the compound eye (front, mid-front, mid-back, back); cells from frontal regions tend to resolve finer gratings.

(v) Sex comparison; male photoreceptors, on average, resolve finer details than those from females.

**Supplementary Table 7** shows the significance of the statistical comparisons.

**Redefining Acuity in a Dynamic Visual System**

Visual acuity has traditionally been viewed as a static property of compound eyes^43^ - typically defined as the finest spatial wavelength that a regularly arranged retinal sampling matrix can resolve, limited by the eye’s interommatidial angles. However, photoreceptors and insect eyes are not passive sensors. Instead, they actively encode space in time^1,3,4^, responding most effectively to phasic changes in the visual environment. This dynamic encoding is co-adapted with active vision: top-down regulated body^77-79^, eye^80^, and intraocular muscle movements^1,38,81^, as well as photoreceptor microsaccades^1,3,4,27^, to maximise information capture.

In natural environments, pure sinusoidal wavelengths are rare. Instead, with sunlight typically falling from above, visual scenes are dominated by sharp contrast transitions caused by occluding 3D objects - such as edges and line elements - whose structure and spatial relationships (i.e. phase congruency^82^) are better defined by angular distances than by spatial wavelengths. Accordingly, in this study, as in our previous work^1,4^, we define visual acuity in terms of angular resolution.

**Directional Tuning and Hyperacute Motion Sensitivity of R1-R6 Photoreceptors**

To investigate how individual photoreceptors encode fine spatial details in moving stimuli, we mapped the directional tuning and resolvability of R1-R6 photoreceptors across the *Musca* compound eye using a narrowing 2D grating stimulus presented in four cardinal directions (Up, Down, Left, Right) at five different speeds (6-120°/s; **Supplementary Fig. 17a**). For each cell, intracellular voltage responses were recorded as the stimulus progressively increased in spatial frequency, enabling us to estimate the finest resolvable inter-bar angle in each direction.

The pooled mean resolvability plot from 19 photoreceptors revealed strong directional biases (**Supplementary Fig. 17b**). Most cells responded optimally to motion in one or two opposing directions - typically Left-Right or Up-Down - rather than uniformly across all directions. This directional tuning was preserved across stimulus speeds, although higher speeds (outer rings) generally produced broader tuning curves and slightly reduced acuity. This reduction likely reflects temporal and structural constraints on R1-R6 photoreceptors’ ability to sample quantal photon signals - set by the number of microvilli in each recorded rhabdomere, and by their refractory periods and latency distributions^6,27^.

Regional analysis of directional tuning (**Supplementary Fig. 17c**-**e**) revealed consistent spatial organisation across the eye. Photoreceptors in the front, middle, and back regions of the left eye exhibited distinct, location-specific tuning axes. These anisotropies are consistent with the known gradients of rhabdomere orientation and microsaccadic motion axes, which rotate systematically across the retina. As in *Drosophila*^3,4^, these axes are expected to be mirror-symmetric between the two eyes. In addition, structural asymmetries in R1-R6 rhabdomeres - such as differences in diameter and deviation from perfect sphericity - contribute to variation in receptive field size and shape (see **Section II**).

Together, these findings demonstrate that individual *Musca* photoreceptors exhibit hyperacute, directionally tuned responses to motion - shaped by both optical and morphodynamic features of the compound eye. These local anisotropies support efficient encoding of behaviourally relevant motion cues during active vision.

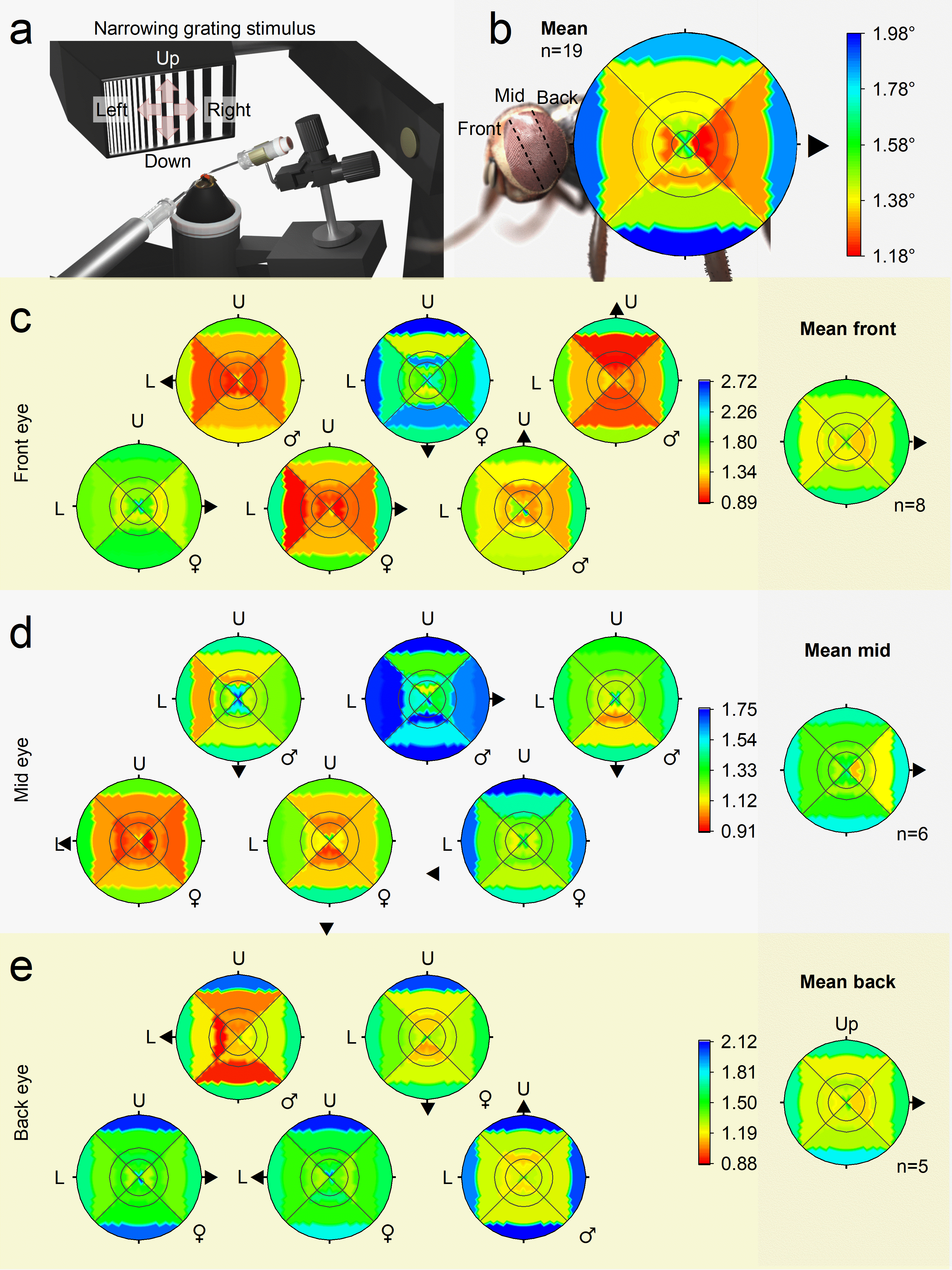
**Supplementary Fig. 17. Directional tuning and hyperacute motion sensitivity of R1-R6 photoreceptor receptive fields across the compound eye.**

(**a**) Experimental setup. A narrowing 2D grating stimulus was presented in four cardinal directions (Up, Down, Left, Right) using a high-speed digital light projector, precisely aligned with the optical axis of individual R1-R6 photoreceptors.

(**b**) Mean directional resolvability map of 19 R1-R6 photoreceptors. Each photoreceptor’s spatiotemporal acuity was tested using grating motion at five speeds (6, 20, 40, 60, and 120°/s) in the four cardinal directions. Polar plots were rotated to align each cell’s peak resolvability with the rightward (0°) axis for averaging. Colour indicates spatial resolution (inter-bar angle in degrees); lower values (warmer colours) indicate finer resolvability. The outer ring corresponds to the highest stimulus speed (120°/s), the inner ring to the slowest (6°/s).

(**c**-**e**) Individual resolvability plots from photoreceptors located in the front (**c**), mid (**d**), and back (**e**) regions of the left eye. Each subplot shows a single cell’s directionally tuned response, with separate panels for male (male) and female (female) flies. Arrows indicate the direction of peak sensitivity. Rightmost plots in each row show the regional means (n = 8 front, n = 6 mid, n = 5 back). Photoreceptors frequently exhibit strong directional tuning, responding best to motion in one or two opposing directions (e.g. Left-Right or Up-Down), consistent with local anisotropies in visual sampling. These anisotropies arise from photoreceptor microsaccades at each eye location, which follow specific motion axes determined by the systematic, gradual rotation of R1-R2-R3 rhabdomere arrangements within ommatidia across the eye. Like in *Drosophila*^1,3,4^, these motion axes are mirror-symmetric between the left and right eyes. Additionally, because R1-R6 rhabdomeres vary in size and their diameters deviate from perfect sphericity, this directly affects the size and shape of their receptive fields (see **Section II**).

**Supplementary Tables 8** and **9** show the significance of the statistical comparisons.

**Supplementary Table 7.** Finest resolved angles as a function of grating speed, using the data in **Supplementary Fig. 16**.

| **Group A** | **Group B** | **A: count of recordings (n)** | **B: count of recordings**  **(n)** | **Mean difference (A-B) (°)** | **Test** | **p-value (Holm-Sidak)** |  |
| --- | --- | --- | --- | --- | --- | --- | --- |
| 6 °/s | 20 °/s | 77 | 77 | 3.119 x 10^-1^ | Mann-Whitney | 3.811 x 10^-4^ | *** |
| 6 °/s | 40 °/s | 77 | 77 | 3.205 x 10^-1^ | Mann-Whitney | 5.162 x 10^-4^ | *** |
| 6 °/s | 60 °/s | 77 | 77 | 2.736 x 10^-1^ | Mann-Whitney | 1.696 x 10^-2^ | * |
| 6 °/s | 120 °/s | 77 | 77 | -2.541 x 10^-1^ | Mann-Whitney | 1.449 x 10^-4^ | *** |
| 20 °/s | 40 °/s | 78 | 78 | 8.551 x 10^-3^ | Mann-Whitney | 8.356 x 10^-1^ | ns |
| 20 °/s | 60 °/s | 78 | 78 | -3.836 x 10^-2^ | Mann-Whitney | 3.733 x 10^-1^ | ns |
| 20 °/s | 120 °/s | 78 | 78 | -5.661 x 10^-1^ | Mann-Whitney | 1.409 x 10^-16^ | *** |
| 40 °/s | 60 °/s | 77 | 77 | -4.691 x 10^-2^ | Mann-Whitney | 3.733 x 10^-1^ | ns |
| 40 °/s | 120 °/s | 77 | 77 | -5.746 x 10^-1^ | Mann-Whitney | 8.242 x 10^-20^ | *** |
| 60 °/s | 120 °/s | 78 | 78 | -5.277 x 10^-1^ | Mann-Whitney | 2.754 x 10^-18^ | *** |

We conducted statistical comparisons of the finest resolved grating angles across three experimental categories: stimulus speed (**Supplementary Table 7**), recording location (**Supplementary Table 8)**, and fly sex (**Supplementary Table 9**). For each comparison, we first tested for normality using the D’Agostino-Pearson normality test (α = 0.05). As none of the groups followed a normal distribution, we proceeded with the non-parametric Mann-Whitney U-test. To correct for multiple comparisons and control the family-wise Type I error rate, we applied the Holm-Šidák step-down method to adjust p-values within each table independently.

**Supplementary Table 8**. Finest resolved angles as a function of recording location, using the data in **Supplementary Fig. 17**.

| **Group A** | **Group B** | **A: count of recordings**  **(n)** | **B: count of recordings**  **(n)** | **Mean difference (A-B) (°)** | **Test** | **p-value (Holm-Sidak)** |  |
| --- | --- | --- | --- | --- | --- | --- | --- |
| Front | Mid-front | 100 | 60 | 2.335 x 10^-1^ | Mann-Whitney | 2.472 x 10^-3^ | ** |
| Front | Middle | 100 | 128 | -1.337 x 10^-1^ | Mann-Whitney | 2.156 x 10^-1^ | ns |
| Front | Mid-back | 100 | 20 | 1.773 x 10^-1^ | Mann-Whitney | 4.999 x 10^-1^ | ns |
| Front | Back | 100 | 80 | 8.856 x 10^-2^ | Mann-Whitney | 5.246 x 10^-1^ | ns |
| Mid-front | Middle | 60 | 128 | -3.672 x 10^-1^ | Mann-Whitney | 3.189 x 10^-7^ | *** |
| Mid-front | Mid-back | 60 | 20 | -5.613 x 10^-2^ | Mann-Whitney | 1.816 x 10^-1^ | ns |
| Mid-front | Back | 60 | 80 | -1.449 x 10^-1^ | Mann-Whitney | 1.415 x 10^-3^ | ** |
| Middle | Mid-back | 128 | 20 | 3.110 x 10^-1^ | Mann-Whitney | 7.371 x 10^-2^ | ns |
| Middle | Back | 128 | 80 | 2.223 x 10^-1^ | Mann-Whitney | 7.371 x 10^-2^ | ns |
| Mid-back | Back | 20 | 80 | -8.878 x 10^-2^ | Mann-Whitney | 4.999 x 10^-1^ | ns |

**Supplementary Table 9**. Finest resolved angles as a function of sex, using the data in **Supplementary Fig. 16** and **17**.

| **Group A** | **Group B** | **A: count of recordings**  **(n)** | **B: count of recordings**  **(n)** | **Mean difference (A-B) (°)** | **Test** | **p-value (Holm-Sidak)** |  |
| --- | --- | --- | --- | --- | --- | --- | --- |
| Male | Female | 204 | 184 | 3.585 x 10^-2^ | Mann-Whitney | 3.812 x 10^-1^ | ns |

**I.8. LMCs Evoke Ultra-Hyperacute Responses to Occluding Moving Stimuli**

To assess the relative contributions of specific pre- and postsynaptic mechanisms in the morphodynamic neural superposition system, we simulated LMC responses using progressively reduced model architectures and compared them directly with intracellular recordings obtained under identical stimulus conditions. To ensure analytical consistency, we present intracellular responses (**Fig. 6** and **Supplementary Fig. 18**) from a single LMC that maintained stable response characteristics throughout the prolonged recording procedures.

**Spatial stimulation and model configuration**

To model responses to spatially structured stimuli, we simulated a single LMC receiving input from its six presynaptic R1-R6 photoreceptors, each originating from a distinct ommatidium.

A virtual screen (15 × 15 mm; 150 × 150 pixels) was positioned 5 cm in front of the eye, with the visual axes of all six photoreceptors centred on the screen. For these experiments, we used the same intracellular recording and back-projection video stimulation system as described above (**Supplementary Fig. 14**).

Each photoreceptor was ray-traced individually, incorporating microsaccadic movements and voltage responses using the previously validated morphodynamic phototransduction model calibrated under bursty stimulation. The modelling approach follows that described in Kemppainen et al.^4^ for *Drosophila*. LMC responses were computed using the calibrated synaptic model (**Supplemental Notes IV**), with light intensities matched to the experimental conditions. The full morphodynamic neural superposition model was identical to that shown in **Fig. 5** of the main paper.

To probe spatial discrimination under demanding conditions, we used double-dot stimuli (e.g. white dots 1.4° wide × 0.7° high on a black background). These stimuli are more challenging to resolve than alternating black–white gratings because local adaptation and contrast interactions reduce separability.

**Model variants**

We evaluated four model configurations (as illustrated in **Supplementary Fig. 18 and Fig. 6**):

**c**, **Full morphodynamic neural superposition**

Included photoreceptor microsaccades, synaptic feedback, and stochastic refractory quantal sampling (see **Supplementary Notes IV**; **Supplementary Fig. 32**).

**d**, **Static neural superposition**

This model was otherwise identical to **c**, but lacked photomechanical photoreceptor microsaccades; LMC received input from perfectly aligned photoreceptor receptive fields of identical size (a classical assumption), while retaining synaptic feedback and stochastic refractory quantal sampling.

**e**, **Quasi-classical neural superposition**

Again, this model lacked microsaccades. In addition, LMCs received input from perfectly aligned photoreceptor receptive fields of identical size (a classical assumption), while retaining synaptic feedback and stochastic, refractory quantal sampling. We only show here the predicted responses for the full receptive field, as the responses of the clipped receptive field were broadly similar to those of the static neural superposition (**d**), lasting only slightly longer.

**f**, **Classic stationary neural superposition filter model**

Implements the assumptions of pioneering stationary filter models^46,47^ (i.e., LMC impulse response convolved with a stationary receptive field) without stochastic quantal sampling or morphodynamic mechanisms. The filter parameters of the photoreceptor and LMC are taken from the same recordings as the full model. For the spatial filter, a half-width of 2.6° was used (**Supplementary Table 15**).

**Occlusion paradigm**

**Supplementary Fig. 18** compares predicted and recorded LMC responses when moving objects become occluded.

In both recordings and simulations, the geometric centre of the LMC receptive field (RF) was positioned at the edge of a dark occluding plate covering 42% of the RF (RF centre 0.8° from the plate edge). One or two small bright dots moved laterally and disappeared behind the occluding plate, mimicking occlusion in cluttered natural environments.

Simulated responses are shown for dot velocities of 42 ° s^-1^ and 168 ° s^-1^ (saccadic velocity). For each velocity, responses are presented for one dot and for two dots separated by 0.7°, 2.1°, and 3.5°. Model predictions (coloured traces) are compared directly with corresponding intracellular recordings (black or dotted traces).

Resolvability (first-peak/second-peak ratio) and temporal delay (measured at peak response) were quantified as functions of dot separation for both velocities. Near-zero delay indicates accurate temporal prediction.

Some timing drift can occur in the experimental recordings. As reported previously^1,4,38,81^intraocular muscle activity can slowly shift photoreceptor receptive-field centres during prolonged experiments. Because these recordings required many minutes to complete, such drift was likely unavoidable. To correct for this, we temporally aligned the recorded voltage responses with the corresponding model simulations by matching the first-derivative peaks of the responses evoked by the first moving dot. This procedure enabled accurate comparison of LMC peak timings between recordings and simulations.

Because instrumental noise was relatively high in these intracellular recordings, the recording traces were up-scaled to facilitate waveform comparison with simulations; specifically, the smallest response (single dot at 42° s⁻¹) was scaled to approximately match the amplitude of the full morphodynamic model prediction.

**Ultra-hyperacute performance under occlusion**

The average interommatidial angle of the *Musca* compound eye is ~2.9° (**Supplementary Table 10**), defining the static anatomical sampling limit.

The full morphodynamic neural superposition model resolves dot separations well below this limit, even at saccadic velocities. This demonstrates that motion-coupled morphodynamic sampling preserves hyperacute discrimination when objects disappear behind occlusions.

Critically, the full morphodynamic model accurately reproduces the brief, precisely timed LMC responses observed experimentally - both when dots disappear (**Supplementary Fig. 18**) and when they appear (**Fig. 6**) behind the occluding plate. These results demonstrate that microsaccadic morphodynamics enhance temporal precision and spatial discriminability under occlusion, thereby reducing motion blur while supporting predictive coding in the photoreceptor–LMC circuitry during fast behaviour in natural three-dimensional environments.

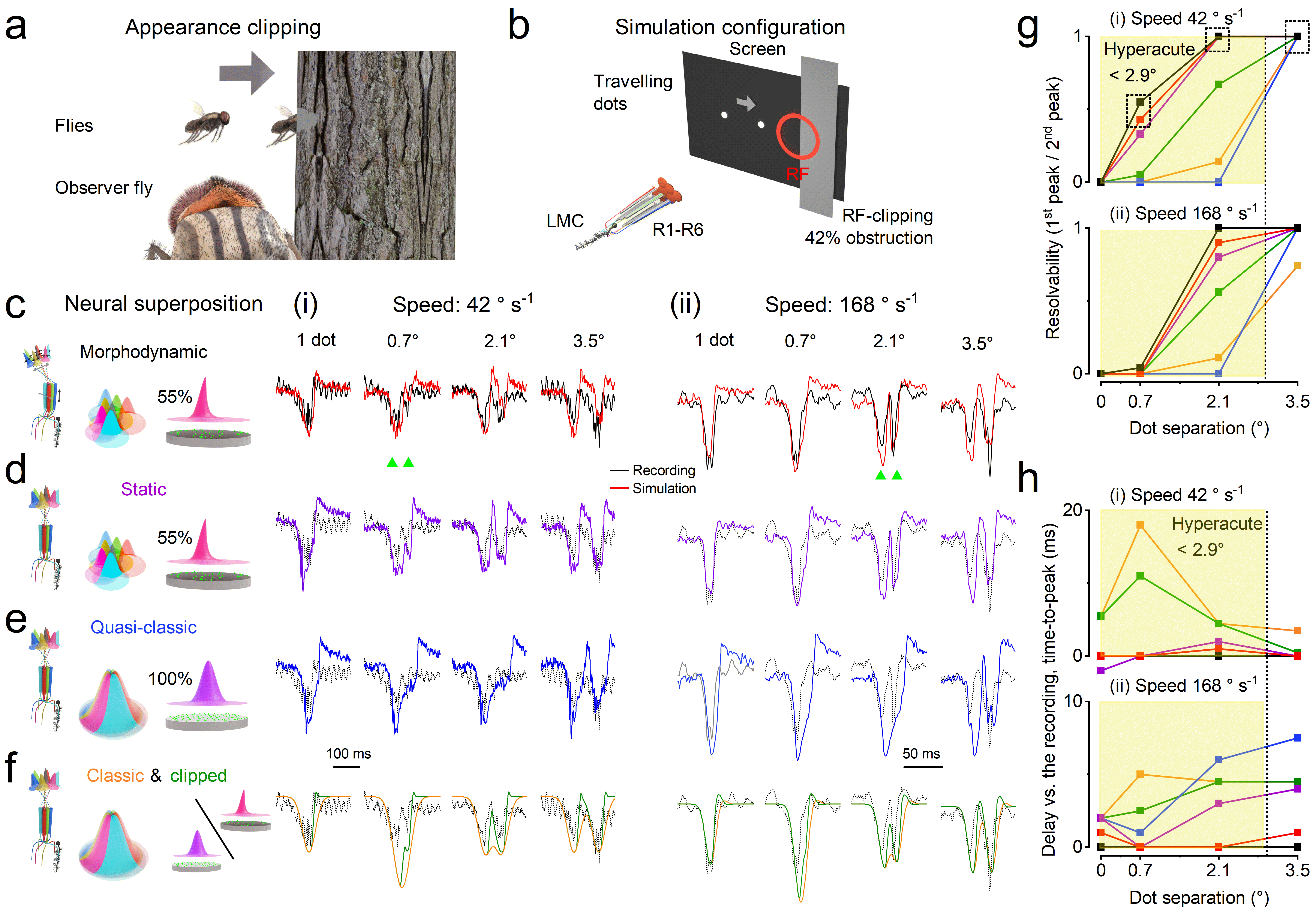

**Supplementary Fig. 18 | Testing predicted and recorded LMC responses to occluded moving objects under different neural superposition models.**

We compared simulated outputs of large monopolar cells (LMCs) under progressively reduced neural superposition models with corresponding intracellular LMC recordings during controlled visual stimulation. In both recordings and simulations, the geometric centre of the LMC receptive field (RF) was positioned at the edge of a dark occluding plate that covered 42% of the RF (the RF centre was 0.8° from the plate edge). One or two small dots, mimicking sunlit flying insects, suddenly disappeared behind the occluding plate, reproducing the visual situation in which motion is occluded in cluttered natural environments.

**a**, Conceptual illustration of the behavioural scenario: an observer fly detects moving objects as they disappear behind an occluding structure.

**b**, Experimental and simulation configuration. Travelling dots move laterally across the visual field and disappear behind an occluding screen, causing receptive-field clipping. LMC responses integrate inputs from R1-R6 photoreceptors.

**c-f**, Simulated LMC voltage responses under progressively reduced neural superposition models:

**c**, full morphodynamic neural superposition;

**d**, static neural superposition (no photoreceptor microsaccades) with overcompletely tiled photoreceptor receptive fields, synaptic feedback, and refractory stochastic quantal sampling;

**e**, quasi-classical neural superposition (no microsaccades), in which LMCs receive input from perfectly aligned photoreceptor receptive fields of identical size (classical assumption), while retaining synaptic feedback and stochastic, refractory quantal sampling;

**f**, classic stationary neural superposition filter model following the assumptions of pioneering studies.

Responses are shown for dots moving at 42° s⁻¹ (i, left) and 168° s⁻¹ (saccadic velocity; ii, right). For each speed, responses are shown for one dot and for two dots separated by 0.7°, 2.1°, and 3.5°. Coloured traces indicate model predictions; black (or dotted) traces show corresponding intracellular recordings.

**g**, Resolvability (first peak/second peak ratio) between simulated and recorded LMC responses as a function of dot separation for 42° s⁻¹ (top) and 168° s⁻¹ (bottom).

**h**, Delay (measured at the peak response) between simulated and recorded LMC responses as a function of dot separation for 42° s⁻¹ (top) and 168° s⁻¹ (bottom). Near-zero delays indicate accurate temporal prediction.

The average interommatidial angle of the *Musca* compound eye is ~2.9° (**Supplementary Table 10**), defining the static anatomical resolution limit. The morphodynamic neural superposition model resolves dot separations well below this limit, even at saccadic velocities, demonstrating that motion-coupled morphodynamic sampling preserves hyperacute discrimination when objects disappear behind occlusion. Notably, the morphodynamic model most accurately reproduces the brief, precisely timed LMC responses observed experimentally, indicating that photoreceptor microsaccades reduce motion blur and enhance predictive coding.

**Fig. 6** presents the complementary case, where dots appear from behind the occluding plate.

Final note: We have also verified experimentally and through simulations (data not shown) that photoreceptor - and consequently LMC - resolvability increases with increasing occlusion. This was tested by placing slits of different widths in front of the retinal lenses during moving stimuli. Because resolvability is determined by the convolution of the impulse response with the receptive field^46,47^ - and the receptive field width typically limits acuity - increasing occlusion effectively sharpens the sampling aperture, thereby improving LMC spatial discrimination beyond the 42% occlusion condition shown in **Supplementary Fig. 18** and **Fig. 6**. See also **Supplementary Note II**, **Supplementary Fig. 25**. To our knowledge, the potential benefit of receptive field clipping by photomechanical photoreceptor microscaccades, or by natural occlusion, for hyperacute discrimination has not previously been tested experimentally.

**II. Analysing Compound Eye Static and Morphodynamic Optics**

**II.1 SEM**

Adult *Musca* (**Supplementary Fig. 19**) and Drosophila (**Supplementary Fig. 20**) were anasthesised with CO_2_, the head removed and fixed in Davidson’s modified fluid^83^ for 24 hrs at room temperature, gently shaking, then dehydrated in steps of 30, 50, 70 and 3x 100% Ethanol for 24 hrs each, critical point dried in a Tousimis 931.GL Critical Point Dryer and mounted onto sticky carbon tabs on SEM stubs, gold coated (10 nm) and imaged in a Hitachi S-3400N SEM with secondary electrons at 5 kV.

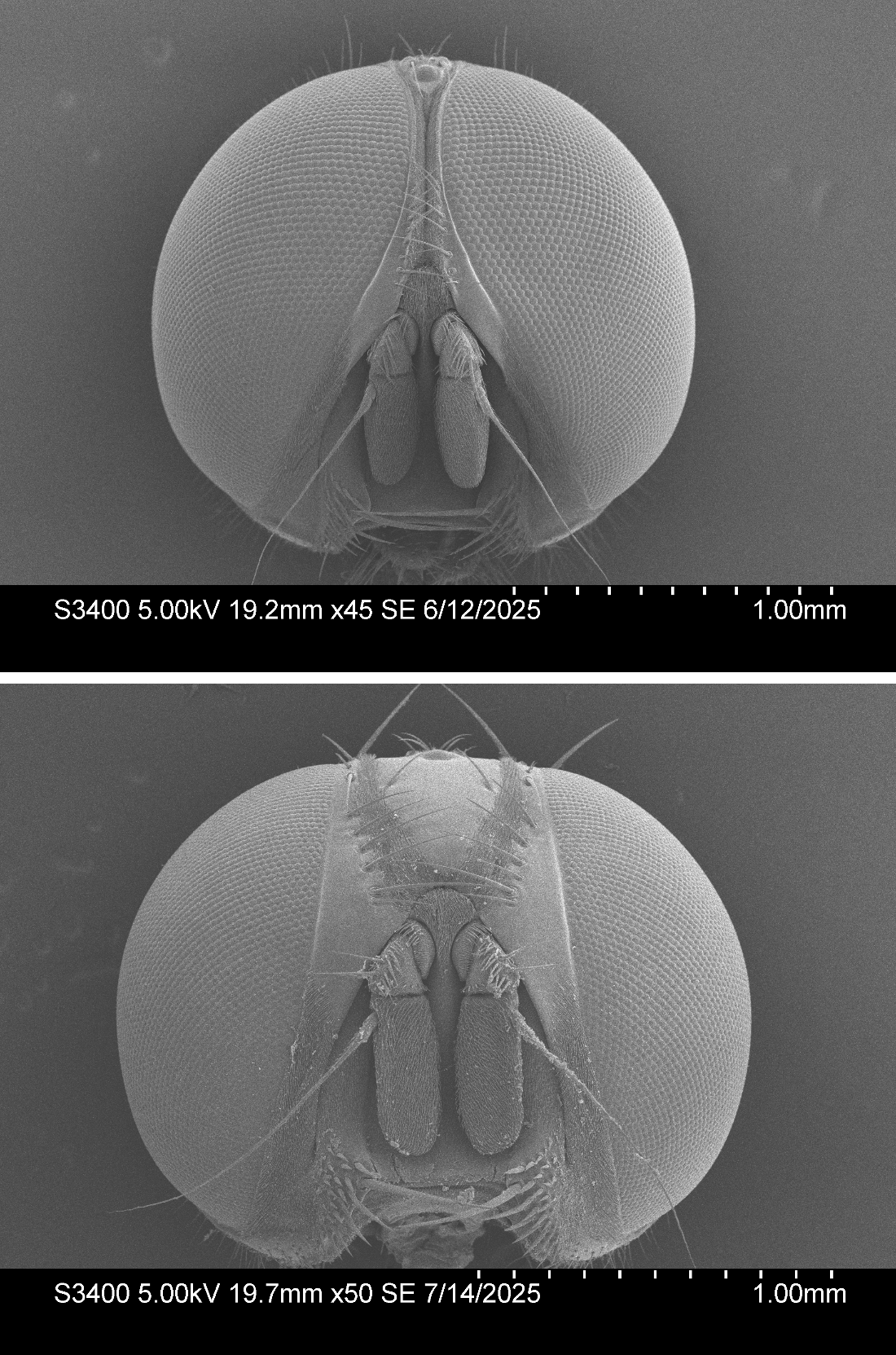
**Supplementary Fig. 19. Scanning electron micrographs (SEMs) of male (top) and female (bottom) *Musca domestica* heads showing sex-specific differences in compound eye morphology.** The tightly packed ommatidia form a hexagonal lattice across both eyes, with notable curvature and regional variations in facet size. In males, the compound eyes are larger and meet at the top (holoptic), featuring an anterior region with enlarged facets - commonly referred to as the “love spot” - believed to enhance high-acuity visual tracking during courtship. In contrast, female eyes remain dichoptic and lack this regional facet enlargement.

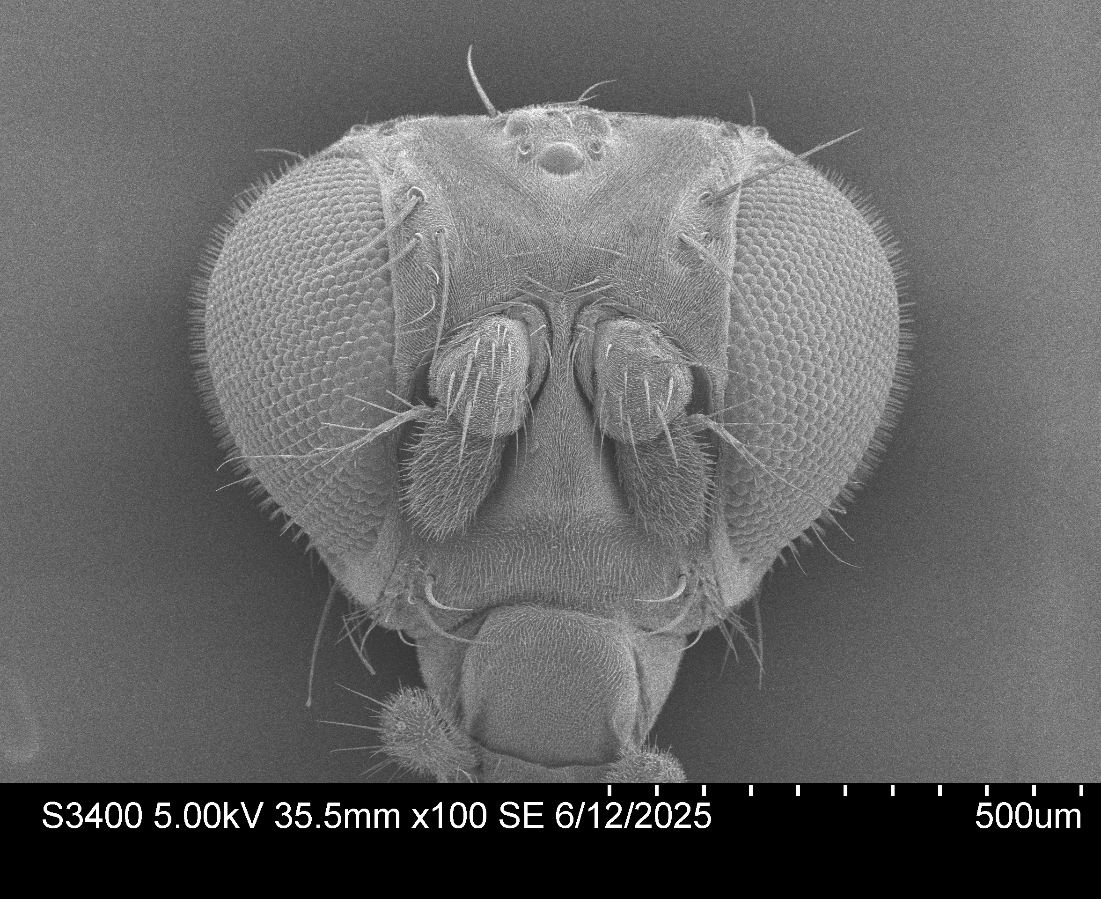
**Supplementary Fig. 20. Scanning electron micrograph (SEM) of a male *Drosophila melanogaster* head showing the structure of its compound eyes.** Compared to the larger *Musca domestica* eyes (**Supplementary Fig. 19**), *Drosophila* eyes are smaller, more hemispherical, and contain significantly fewer ommatidia (~750 per eye vs. ~4,000-5,000 in *Musca*). The ommatidia form a regular hexagonal lattice, but without the regional enlargement seen in the male *Musca* “love spot.” Unlike *Musca*, *Drosophila* males retain a dichoptic eye arrangement with a clearly separated frons, and the interommatidial angles are larger, contributing to lower spatial resolution. Nonetheless, the slower-flying *Drosophila* eyes remain highly effective for close-range 3D vision^4^ and visual behaviours. See also **Figure 5** in the main paper.

**II.2 Structural X-ray Imaging using HiTT**

The heavy metal-stained and resin-embedded fly heads were imaged using high-throughput imaging tomography (HiTT) at the P14 beamline in EMBL Hamburg. HiTT is a phase-contrast based X-ray imaging technique which has been optimised in both speed and quality for imaging biological samples; for full details of the imaging setup, please see^84^. The HiTT imaging is completely non-destructive, allowing downstream imaging with other techniques like SEM.

Here, the resin blocks were mounted vertically and imaged off-centre to enable 360˚ rotation of the sample, which provided an XY field of view of ~2 mm. Each fly head was imaged using four-distance holo-tomography^85^ at an energy of 23.5 keV using a 10X objective to achieve a nominal pixel size of 0.65 µm. Due to the size of the fly heads, a tiled acquisition in the z-plane was also required; two scans with an offset of 450 µm were acquired and stitched into one dataset using NRStitcher^86^. Each scan was acquired in 4.5 minutes and reconstructed in ~1 minute. The results are then visible live on the beamline. The 3D reconstructed data had an isotropic voxel size of 0.65 µm, which was sufficient to visualise the key anatomical features in the fly, as shown in **Supplementary Fig. 21**, including the eye as detailed below.

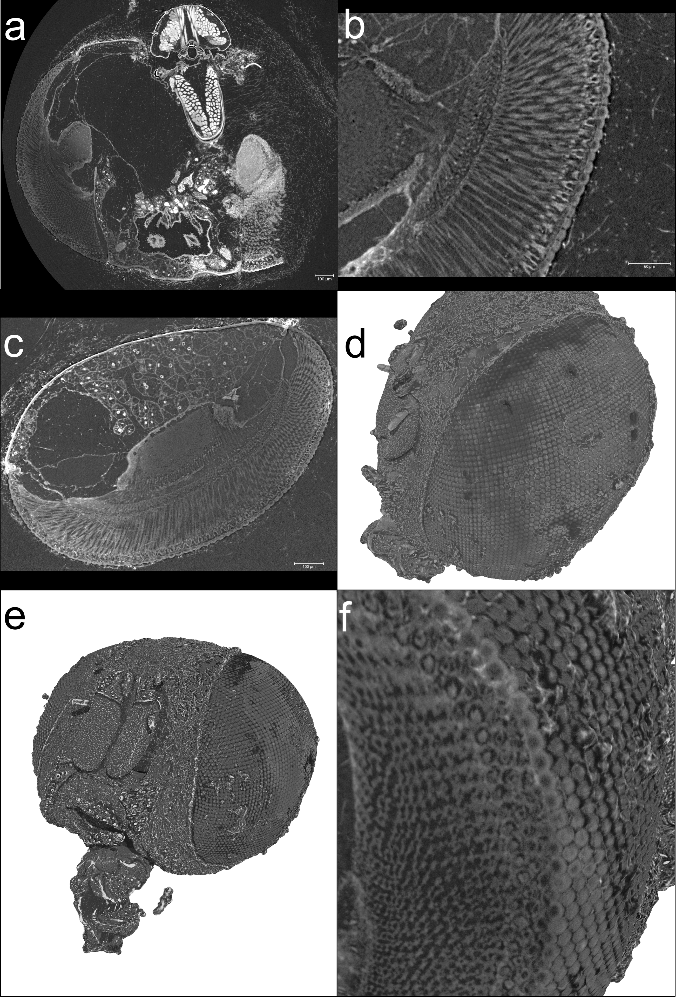
**Supplementary Fig. 21**. Synchrotron X-ray data of a female head imaged with high-throughput imaging tomography (HiTT). 3D volume rendering was created from reconstructed HiTT data in VG Studio Max (v2024.4, Volume Graphics, Germany) (**a**) A reconstructed 2D orthoslice through the head of the fly; here the structures of the brain, eyes and oesophagus can be seen. (**b**) A zoomed-in image of the eye showing the connections between the retinal R1-R7/8 photoreceptors and the lamina optic lobe in the brain. (**c**) A 2D cross-section of the eye viewed from a different angle to the one shown in (**b**, here the internal structure of the compound eye can be seen. (**d**) Volume rendering showing the surface of the eye from panel (**c**). (**e**) Volume rendering of the whole fly head viewed from the front, here both compound eyes are visible as well as the other features of the fly head. (**f**) Magnified image of the volume rendering of the fly eye, which has been digitally ‘clipped’ so that the internal ommatidial structures of the eye are visible in 3D.

**II.3 Analysing *Musca* Compound Eyes Static Optical Properties**

The *Musca* compound eye comprises thousands of nearly identical lens-capped ommatidia arranged in a regular hexagonal lattice (**Supplementary Fig. 19**). To quantify their optical and geometric properties, we used X-ray synchrotron imaging to reconstruct the female head (**Supplementary Fig. 21**) and rotated the 3D volume such that the head axis aligned with the image axis in 8-bit resolution. For each ommatidium along the equatorial (horizontal) line, we annotated six key structural points: three per lens (including corners at a fixed vertical height) and two per rhabdomere (marking both ends, each with a defined vertical coordinate) (**Figure 1**, **Supplementary Fig. 22a**). In total, ommatidia along this equatorial line were organised into 50 vertical columns, with minor variation in vertical position.

Inner ($r₁$) and outer ($r₂$) lens radii were estimated for each ommatidium using three annotated points per lens (**Supplementary Fig. 22b-c**, **Supplementary Table 10**). These radii showed no systematic variation across columns. Our measurements of the outer lens radius (~17 µm) agree with earlier estimates by Vowles^87^, while our inner radius measurements (~19 µm) appear clearer and more consistently defined than in previous studies. Lens thickness and diameter **Supplementary Fig. 22d-e**) were also computed from the annotated points (**Supplementary Fig. 22a**), with results consistent across columns and comparable to those reported by Vowles^87^ and Stavenga^88^.

Crystal cone and rhabdomere lengths were derived from 3D point annotations across the ommatidia (**Supplementary Fig. 22f-g**). Both showed the shortest values near the midline, with mean lengths reported in **Supplementary Table 10**. Our rhabdomere length estimates (~100-180 µm, mean ~132 µm) are in line with values reported by Boschek^89^ and Braitenberg^90^.

To estimate the interommatidial angle, we analysed the horizontal orientation of ommatidia using lens point coordinates (**Supplementary Fig. 22h**). Because the vertical columns of ommatidia are not perfectly aligned with the equatorial line, we corrected column indices by scaling the horizontal distance between ommatidia by the ratio 4/3, corresponding to the relationship between horizontal and total distances in a hexagonal lattice. A linear fit between the horizontal angle and corrected column values - excluding the first three outliers - yielded a *horizontal interommatidial angle*^22^ of 2.18 ± 0.02°. Applying the 4/3 correction factor gives a ***true interommatidial angle*** of **2.90 ± 0.03**° (**Supplementary Fig. 22i**), consistent with previous estimates. While Beersma et al.^22^ reported considerable regional variation in male *Musca*, particularly in the anterior “love spot,” our measurements from a female showed consistent interommatidial angles along the equator.

**Supplementary Table 10**. Optical properties of female *Musca* along the equator line

| **Property** | **Value** |
| --- | --- |
| Outer lens radius ($\boldsymbol{r}\boldsymbol{1}$) | 17.3 ± 1.5 µm |
| Inner lens radius ($\boldsymbol{r}\boldsymbol{2}$) | -19.4 ± 6.7 µm |
| Lens thickness | 8.7 ± 0.8 µm |
| Lens diameter | 22.2 ± 1.1 µm |
| Horizontal interommatidial angle | 2.18 ± 0.02 ° |
| Interommatidial angle | 2.90 ± 0.03 ° |
| Crystal cone length | 29.5 ± 4.2 µm |
| Rhabdomere length | 132 ± 20 µm |

We excluded the 2 outer columns on both sides from the distance calculations. From angle fit calculations, we excluded the 3 innermost ommatidia.

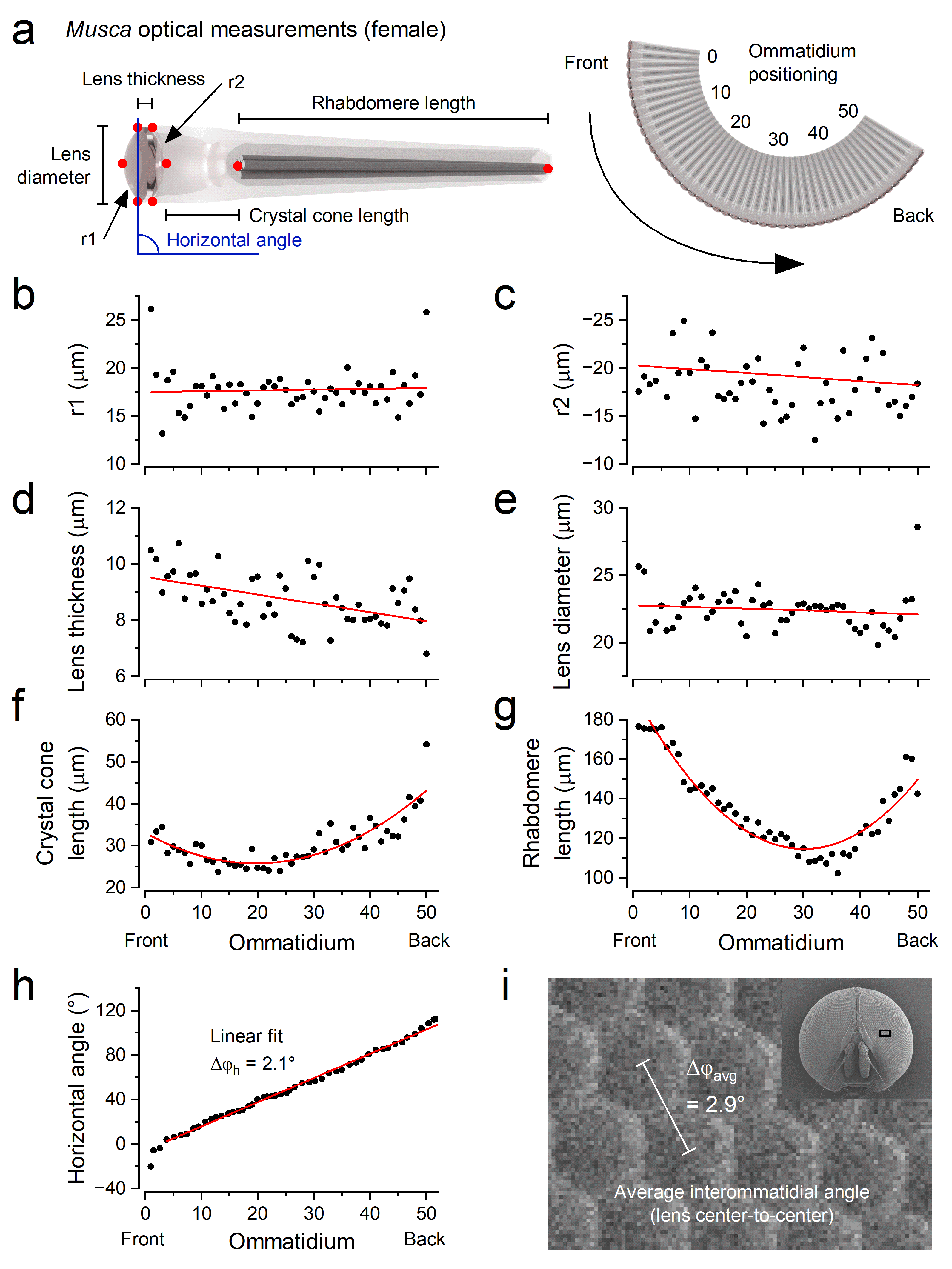
**Supplementary Fig. 22. Optical properties measured along the equator of the *Musca* compound eye**.

(**a**) The optical system of *Musca* comprises a lens, crystal cone, and rhabdomere. For each ommatidium along the equator - from the front to the back of the eye - we annotated six key points (red dots) to quantify structural parameters (**b**-**h**) in female *Musca*.

(**b**-**c**) Inner (r₁) and outer (r₂) lens radii were computed using a three-point algorithm applied along the equator.

(**d**-**e**) Lens thickness and diameter were similarly measured using the annotated surface points.

(**f**-**g**) Crystal cone and rhabdomere lengths were calculated in 3D from point annotations.

(**h**) The horizontal orientation of each ommatidium was estimated from lens positions. The blue angle in (**a**) illustrates the measurement. To compute the average horizontal interommatidial angle, ${\Delta\varphi}_{h}$, we fitted a line (red) through the central portion of the data, correcting column indices for vertical displacement.

(**i**) Following the tiling arrangement of corneal lenses both in female and male (insect) *Musca*, the average interommatidial angle, $\Delta\varphi$ - measured from the lens centre to the next lens centre - is about 2.9°.

**II.4 Serial Block-Face Scanning Electron Microscopy (SBF-SEM)**

Adult female *Musca* were anaesthetised with CO₂. Heads were removed, and most of one side of each head was excised with microscissors to allow fixative infiltration. Dissected heads were fixed overnight at 4 °C in 0.1 M sodium cacodylate buffer containing 0.75% formaldehyde and 0.75% glutaraldehyde, with gentle shaking.

After fixation, samples were washed in 0.1 M sodium cacodylate buffer and stained overnight at 4 °C with 2% osmium tetroxide in the same buffer, again with gentle shaking. The heads were then:

- Washed and incubated in 2.5% potassium ferrocyanide in 0.1 M buffer at room temperature for 2 h on a rocker
- Washed with deionised water (DI)
- Incubated in 1% thiocarbohydrazide (aqueous), washed
- Stained again with 2% aqueous osmium tetroxide overnight at 4 °C with gentle shaking
- Washed and stained with 1% aqueous uranyl acetate at room temperature on a rocker, then moved to 50 °C for 20 min
- Washed with DI water until clear
- Stained with 3% Reynolds lead citrate (Taab) at room temperature on a rocker for 1 h
- Washed and dehydrated through a graded ethanol series (30%, 50%, 70%, and 3× 100%), at least 4 h per step
- Incubated twice in 100% acetone for 4 h each

Samples were then infiltrated with 812 hard Epon resin in steps (30%, 50%, 70%, and 3× 100%) for 6 h each or overnight. Heads were finally embedded in 100% 812 Epon resin in block moulds and cured at 60 °C for 48 h.

Resin-embedded heads were trimmed out using a fine saw and razor blades, then mounted onto SBF-SEM stubs with conductive epoxy resin (Chemtronics) and hardened for 8 h at 70 °C. Sample sides were trimmed with a trimming knife, and the entire stub was sputter-coated with a 20 nm gold layer to reduce charging.

Images were acquired using a Merlin Compact SEM (Zeiss, Cambridge, UK) equipped with a Gatan 3View system and Gatan OnPoint backscatter detector. Acquisition settings were:

- Pixel size: 5 nm
- Dwell time: 1 μs
- Aperture: 20 nm
- Acceleration voltage: 2 kV
- Vacuum mode: high vacuum
- Charge compensation: 100% Zeiss Focal Charge Compensation
- Section thickness: 100 nm

**II.5 Transmission electron microscopy (TEM)**

Male Musca were sedated by incubation on ice. Heads were bisected and fixed in cold fixative (2.5% paraformaldehyde, 2.5% glutaraldehyde in 0.1 M phosphate buffer (PB), pH 7.3) for 12 h. After rinsing in PB, samples were post-fixed with reduced osmium for 2 h at room temperature (0.75% potassium ferrocyanide, 1.3% osmium tetroxide in 0.1 M PB). Following rinsing in water, tissues were dehydrated through a graded ethanol series and propylene oxide. Propylene oxide was then replaced by Durcupan (Sigma, formulation 3). Ultrathin sections (70 nm) were imaged using a Talos L120C electron microscope.

**II.6 Analysing Rhabdomere Sizes and Positions**

*Musca* rhabdomeres - densely packed with microvilli, the protruding compartmentalised photon-sampling units - exhibit a highly conserved spatial arrangement across ommatidia (**Supplementary Fig. 23**; **Figure 1a-iv**) The R1-R7/R8 photoreceptors are consistently organised, except for a developmentally programmed rotation that slightly offsets the rhabdomere pattern between neighbouring ommatidia. To quantify this architecture, we analysed the distal end positions and diameters of rhabdomeres in 14 ommatidia using SBF-SEM data.

For each ommatidium, measurements were taken from three distal-most image planes in the SBF-SEM stack. To improve image clarity, we averaged each 4 × 4 pixel block in the image plane. Ellipses were fitted to each rhabdomere cross-section to determine their major and minor axes.

Rhabdomere diameter was calculated as the diameter of a circle with the same area as the fitted ellipse (**Supplementary Table 11**). The position of each rhabdomere was defined by the centroid of its ellipse, measured relative to the R7/R8 rhabdomere within the same ommatidium (**Supplementary Table 11**). Our measurements align closely with those reported by Beersma et al.^22^, who found typical inter-rhabdomeral spacing of 1.7 µm horizontally and 0.99 µm vertically.

**Supplementary Table 11**. Position and diameter of rhabdomere tips (mean ± SD, n = 14 ommatidia)

| **Photoreceptor** | **Rhabdomere diameter (µm)** | **Rhabdomere X position (µm)** | **Rhabdomere Y position (µm)** |
| --- | --- | --- | --- |
| **R1** | 1.50 ± 0.13 | 1.09 ± 0.16 | -1.86 ± 0.16 |
| **R2** | 1.33 ± 0.11 | 1.47 ± 0.09 | -0.19 ± 0.05 |
| **R3** | 1.53 ± 0.11 | 1.65 ± 0.08 | 1.44 ± 0.09 |
| **R4** | 1.25 ± 0.11 | 0.10 ± 0.10 | 1.48 ± 0.13 |
| **R5** | 1.30 ± 0.07 | -1.37 ± 0.14 | 1.74 ± 0.13 |
| **R6** | 1.47 ± 0.18 | -1.61 ± 0.06 | 0.13 ± 0.13 |
| **R7/R8** | 1.15 ± 0.12 | 0 | 0 |

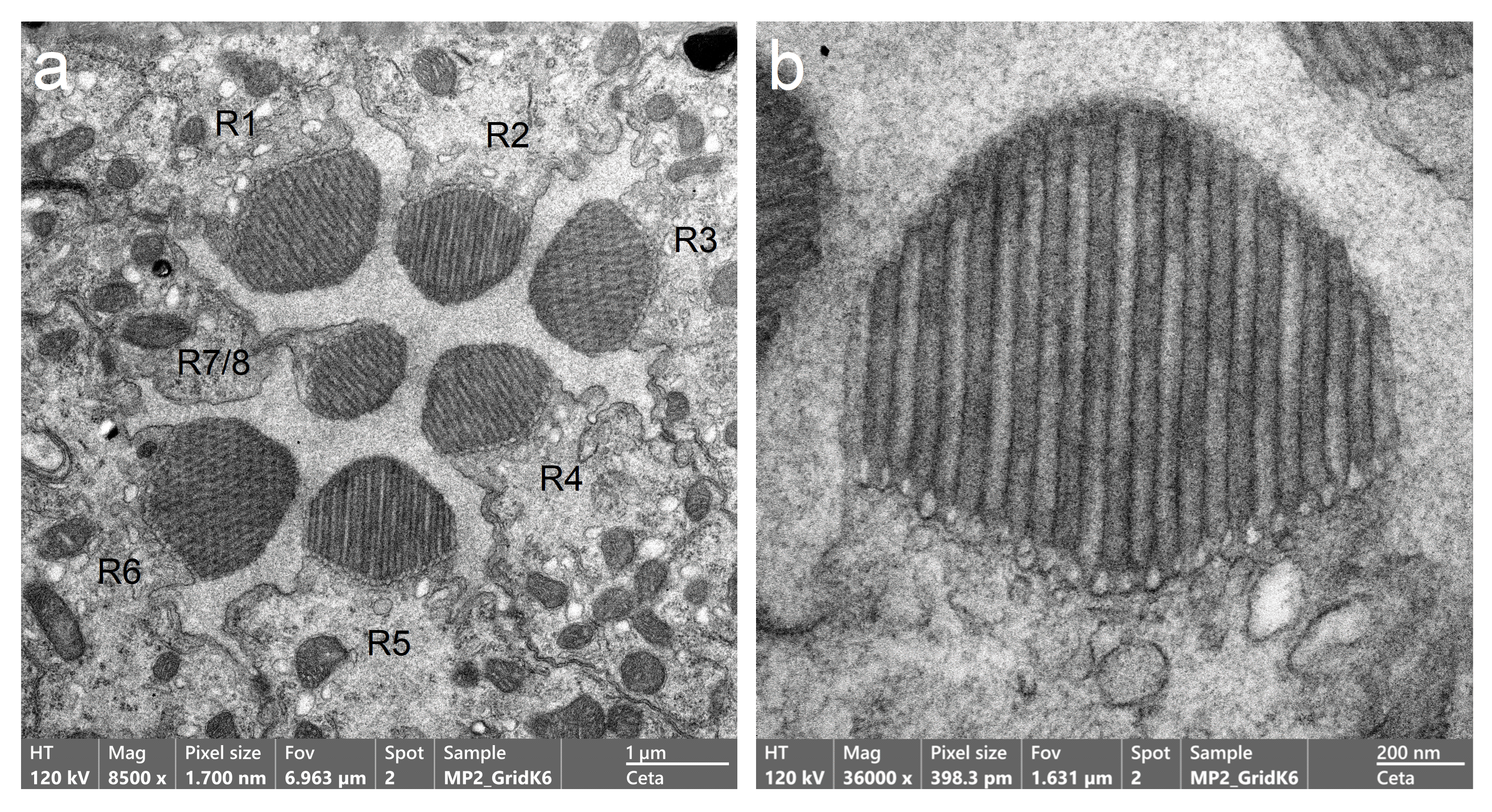
**Supplementary Fig. 23**. **Transmission electron micrographs of *Musca domestica* ommatidia reveal rhabdomere ultrastructure and organised microvillar arrays.**

(**a**) Cross-sectional view of a single ommatidia shows individual R1-R7/8 photoreceptor rhabdomeres with densely packed, radially arranged microvilli. Each rhabdomere appears as a distinct rod-shaped structure, varying in orientation due to sectioning angle.

(**b**) Higher-magnification image reveals the fine structure of microvilli within rhabdomeres, showing their parallel alignment and dense packing. These specialised membrane protrusions maximise the photoreceptive surface area and support the high-speed visual processing demands of the fly.

**II.7 Estimating R1-R7/8 Photoreceptors’ Varying Microvilli Counts**

Rhabdomere size varies depending on its position in the eye (**Supplementary Fig. 22g**) and its diameter (**Supplementary Fig. 23**; **Supplementary Table 11**), leading to differences in microvilli number across photoreceptors. Each microvillus has a diameter of 0.0675 µm^89^, and is hexagonally packed within the rhabdomere. An average R1-R6 rhabdomere, with a diameter of 1.4 µm and length of 132 µm (**Supplementary Tables 10** and **11**; see **Section II.5**), contains approximately 54,000 microvilli. Given that rhabdomere lengths range from about 100 µm to 180 µm (with average diameter constant), the microvilli count varies from roughly 41,000 (shortest) to 74,000 (longest). Within an ommatidium with average-length rhabdomeres, the microvilli count ranges from approximately 44,000 in R7/R8 to 59,000 in R3.

**II. 8 Morphodynamic Optics of the Compound Eye**

Insect compound eyes were traditionally assumed to be static structures, and the mathematics of their optics were formalised accordingly^19,91-95^. If both the eye’s structure and visual processing were static - as in classical models of dipteran flies, where R1-R6 photoreceptors from neighbouring ommatidia were assumed to align in neural superposition with perfectly overlapping receptive fields^52,96,97^ - then these models would suffice.

However, in both *Musca* and *Drosophila*, neural superposition is imperfect^4,61^. Their R1-R6 photoreceptors generate partially overlapping, overcompletely tiled receptive fields (e.g. **Figure 1** in the main article). In addition, insects actively sample visual information^27^ through saccadic head^98^, body^78,79,99,100^ and eye-muscle movements^1,37,38,81^, as well as through photoreceptor microsaccades^1,27,38,63,79,100^, supported by flexible morphodynamic eye structures^1,3,4,38,81^. As a result, classical models - based on sparse, static, pixel-like image formation^19,43^ - can no longer account for their true optical resolving power^1,4^ during active vision. In both *Musca* (see **Supplementary Fig. 15-17**, **Figure 4h**) and *Drosophila*^1,4^, visual acuity - even in head-fixed flies - is at least 3-4 times finer than the eye’s average interommatidial angle, which sets the classical static resolution limit.

In the following sections, we show how the asymmetric layout of R1-R7/8 photoreceptors - combined with their directionally varying morphodynamic (photomechanical) sampling motions, the moving interommatidial aperture^1,4^ (made by cone and pigment cells underneath the crystal cone) and the slight rotational offset of neighbouring ommatidia - enables the *Musca* visual system to resolve details of moving objects in cluttered, occluded environments, *even below the diffraction limit of their lens system*.

**Point-Spread Function (PSF) and Interommatidial Angle - the Absolute Spatial Resolution Limits?**

In *Musca*, the point-spread function ($PSF$ ≈ 623 nm; i.e. spatial impulse response function) of an average ommatidial lens ($D$ = 22.2 μm) for 350 nm UV light is only 41-54% of the rhabdomere diameter (1.15-1.53 μm) onto which the $PSF$ is projected.

Airy disc:

$\theta=1.22 \cdot\frac{\lambda}{D}=1.22\cdot\frac{\left( 350 nm \right)}{\left( 22.2 \mu m \right)}\approx0.0192\mathrm{rad}\approx\mathbf{1.1}^{\boldsymbol{^{\circ}}}$	(1)

Correcting for the refractive index, the effective focal length ($f$) is calculated using thick-lens equations^92^:

$$P_{l}=P_{1}+P_{2}{+P}_{3}$$

$P_{1}=\frac{n_{l}-n}{r_{1}}=0.026 \frac{1}{\mu m}$

$$P_{2}=\frac{n_{c}-n_{l}}{r_{2}}=0.0057 \frac{1}{\mu m}$$

$$P_{3}={-\frac{l_{t}}{n_{l}}P}_{1}P_{2}=0.000885 \frac{1}{\mu m}$$

Thus:

$f=\frac{n}{P_{l}}=32.47 \mu m$	(2)

Here, $P_{l}$ is the total lens power, and $P_{1}$, $P_{2}$ and $P_{3}$ are the respective powers of the three lens surfaces. Parameters used: refractive index of air ($n =1$), lens $n_{l} =1.45$, crystal cone $n_{c} =1.34$ from **Supplementary Table 13**), and lens diameter $l_{t} =8.7 \mu m$ from **Supplementary Table 10**.

The $PSF$, which sets the diffraction limit of the *Musca* compound eye, is then:

$PSF \approx f \times\theta=32.47 \mu m \times0.0192 rad\approx623 nm$	(3)

Even if each ommatidium were jam-packed with an infinite number of photoreceptors, the sharpest image this hypothetical sampling matrix could capture would still be limited by the $PSF$. The finest angular separation resolvable in visual space, by the Rayleigh criterion, would be the Airy disc angle itself ($\boldsymbol{\theta=1.1^{\circ}}$). Since each ommatidial lens projects a blurry image, adding more photoreceptors within an ommatidium offers no resolution gain. Based on this “poor optics” argument, it was concluded that eight photoreceptors - spread out within the ommatidium, with R7 stacked atop R8 at the centre - are sufficient for capturing the coarse image formed by the lens system.

Moreover, each R1-R7/8 rhabdomere acts as a waveguide with a diameter larger than the $PSF$, resulting in relatively broad receptive fields (half-widths in *Musca* vary from ~2.1° to ~2.6°; see **Section IV**, **Supplementary Figs. 29-30**). These fields are not well-suited for capturing fine spatial details. Because their size closely matches the interommatidial angle, (in *Musca*: $\Delta\varphi\approx2.9^{\circ}$; **Supplementary Table 10**), and it was previously assumed that in neural superposition the receptive fields of R1-R6 photoreceptors would effectively be identical in size and perfectly aligned in visual space^43^, the resolution limit of the compound eye has traditionally been equated with this angle.

And yet, our experiments show that *Musca* photoreceptors can resolve moving edges separated by 0.9°, and LMCs can resolve moving point objects separated by 0.7° - both well below the diffraction limit ($\theta=1.1^{\circ}$) and the interommatidial angle ($\Delta\varphi\approx2.9^{\circ}$), which defines the pixel-like spacing of their photoreceptor matrix. How is this possible?

***Musca* Eye Design Exploits Environmental Regularities and Motion to Enhance Acuity**

Occlusions in the optical pathway - such as those caused by the interommatidial aperture^1,4^ or by objects in the 3D environment that clip a photoreceptor’s receptive field - can, counterintuitively, improve visual acuity for detecting moving objects. Simulations show that even in a static compound eye - without photoreceptor microsaccades or retinal movements - the ommatidial optical architecture alone can resolve moving objects in 3D space with higher precision than classical 2D models predict. We illustrate this motion-occlusion principle in neural encoding using a photoreceptor’s Gaussian receptive field as an example (**Supplementary Fig. 24**).

**Case a**: The R2 photoreceptor’s receptive field (~2.1° half-width) is unobstructed, facing the clear sky. Two small, point-source-like, slow-flying flies pass horizontally across its view, separated by 1.1° (the diffraction limit). The model shows the photoreceptor cannot distinguish them as separate objects (**Supplementary Fig. 24bi**).

**Case b**: A dark ebony tree occludes two-thirds of the photoreceptor’s receptive field vertically, leaving a narrow slit (~0.7°) exposed to the sky. As the same two insects pass through this unoccluded portion and disappear behind the tree trunk, the model shows the photoreceptor can now resolve them as distinct entities (**Supplementary Fig. 24bii**).

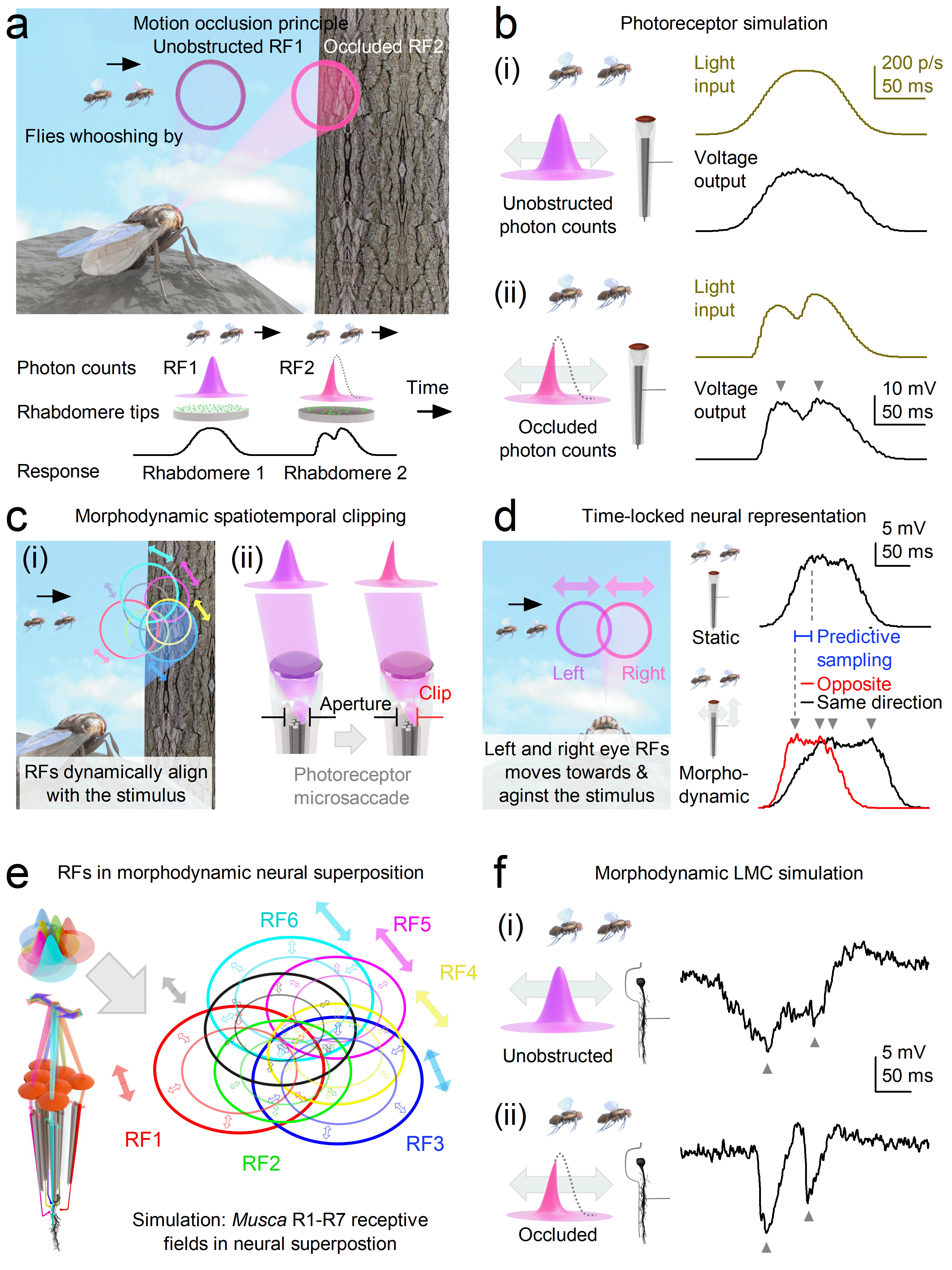
**Supplementary Fig. 24. Morphodynamic mechanisms in the *Musca* compound eye sharpen motion vision through occlusion, microsaccades, and overcompletely tiled neural superposition.**

Simulations are shown under relative dark adaptation with two closely spaced bright spots (“flies”) moving across a typical R1-R6 photoreceptor and a large monopolar cell (LMC) receptive field (RF). Under light adaptation, resolvability increases further (see **Supplementary Fig. 16**, **Figure 4h**). Grey arrowheads indicate when the two passing “flies” were neurally distinguished as separate events in the respective macroscopic voltage responses.

(**a**) **Exploit occlusion**: Two flies cross RFs at different eye positions. The unobstructed RF1 merges both “flies” into a blurred signal, while RF2, partially blocked by a tree trunk acting as a slit, receives fewer photons but gains sharper spatial and temporal resolution as this “slit effect” narrows its RF.

(**b**) **Sharpen photoreceptor output**: Occlusion shortens the integration window, producing crisper voltage responses. Even a static-eye model (as simulated here) with immobile photoreceptors yields hyperacute signals in natural 3D scenes with occluders. The “flies” were 1.1° apart, moving at 42° /s.

(**c**) **Clip with microsaccades**: Photomechanical microsaccades shift RFs relative to the intraocular aperture; edge alignment clips input and narrows RFs, boosting resolution.

(**d**) **Encode depth predictively**: In the frontal binocular zone, left and right RFs move in opposite directions (and narrow) during mirror-symmetric microsaccades. The resulting phase differences encode instantaneous depth and time-lock responses to motion. In contrast, in a hypothetical static eye (above), voltage responses of the left and right photoreceptors - with overlapping RFs - would be identical, appearing as a single trace. The “flies” were 1.4° apart, moving at 42° /s.

(**e**) **Densify sampling**: R1-R6 photoreceptors from neighbouring ommatidia overlap in visual space. Microsaccades shift (filled arrow) and narrow (unfilled arrows) these RFs, increasing sampling density and precision.

(**f**) **Refine LMC signals**: Occlusion sharpens LMC responses by combining RF clipping with redundant inputs from neural superposition. The “flies” were 1.1° apart, moving at 42° /s.

In **case b**, the occlusion clips the receptive field, narrowing it to a slit-like opening. Light from each fly is then sensed only briefly - as a sharp *pang*-*pang* signal. This “slit effect” reduces the effective receptive field half-width to about one-third of its full extent in **case a**, dynamically sharpening both spatial and temporal resolution^1,4^. In engineering terms, a photoreceptor’s spatiotemporal resolving power for moving objects can be approximated by convolving its receptive field with its impulse response^1,46,47^. Here, the narrow slit prevents reflected light from the two passing flies from being integrated (blurred) into a single broad hump. Instead, it transforms - effectively differentiates - the signal into a clear, double-humped light input to the photoreceptor.

Thus, despite their fixed x/y retinotopic positions in the eye, photoreceptors can dynamically encode finer spatial details of moving objects over time - resolving features far beyond classical optical limits, especially in cluttered natural scenes where objects frequently occlude one another. *Crucially, this enhancement depends on motion*. In the absence of movement - when both the eyes and the scene are static - compound eye acuity reverts to the lower limits predicted by classical models. With photoreceptors’ overcomplete receptive fields tiling almost the entire visual environment like a dome, even small body, head, or retinal movements can neurally enhance contrast and make occluding patterns “jump out,” revealing natural contours of overlapping objects in 3D space.

Importantly, this kind of dynamic receptive field narrowing is not produced by environmental occluders alone (as in **case b**); the fly’s own morphodynamic mechanisms - particularly photoreceptor microsaccades - can generate an equivalent slit effect. In the next section, we examine how these self-generated movements actively shift and narrow receptive fields to sharpen vision.

***Morphodynamic Receptive Field Acuity Enhancement*: Photoreceptor Microsaccades Shift and Narrow Receptive Fields to Sharpen Vision**

Insect eyes are not static, pixelated, low-resolution cameras, but morphodynamic sampling and processing organs that operate through stochastic, refractory quantal information flow. Environmental light contrast changes - caused by moving objects or self-motion (body^100^, head^98^, or whole-retina^38^ movements from intraocular muscle contractions) - drive photomechanical local photoreceptor microsaccades across the retinae^27^. During a microsaccade, a photoreceptor’s rhabdomere shifts laterally and moves axially in response to light^1-4^, morphodynamically shifting and narrowing its receptive field. Axial rhabdomere movement narrows the receptive field by moving the rhabdomere away from the ommatidial lens centre, reducing the angular extent from which it collects light. Lateral rhabdomere movement enhances the visual response to stimuli moving either with or against the receptive field’s motion.

**Supplementary Fig. 24d** illustrates these mechanisms using two overlapping receptive fields, one from the left and one from the right eye in the binocular frontal region. Mirror-symmetric microsaccadic motion^1,3,4^ causes one receptive field to move and narrow with the stimulus while the other moves and narrows against it. The resulting phase difference encodes instantaneous depth^4^, while neural representations across the retina become effectively time-locked to object motion.

Microsaccades can also transiently align a receptive field edge with a moving stimulus (**Supplementary Fig. 24ci**) or with the interommatidial aperture (**Supplementary Fig. 24cii**), narrowing the field by clipping the light path in the direction of rhabdomere motion. This morphodynamic spatiotemporal clipping - comparable to the “slit effect” (**Supplementary Fig. 24b**) - sharpens neural representations of moving objects^4^, accentuating edges and contours.

Importantly, all R1-R7/8 photoreceptors within an ommatidium are mechanically coupled^1,4^. When one rhabdomere is photomechanically activated, it induces a microsaccadic movement in the others. This coupling makes light sampling predictive: a photoreceptor can be mechanically drawn into the light path just before the stimulus would naturally reach it, reducing delays and enhancing temporal predictability^4^.

***Temporal “Impulse Response” Sharpening:* Morphodynamic Refractory Quantal Sampling Enhances Macroscopic Voltage Responses to Salient Contrast Changes**

Stochastic refractory quantal photon sampling gives photoreceptors an innate capacity to enhance stimulus salience. Adaptation occurring *after* phototransduction or synaptic transmission cannot increase the information transfer rate^6,35^. By contrast, adaptation *during* the sampling process can accentuate changes in quantum bump rate to new (surprising) stimuli, transiently increasing information transfer. Because the microvillar phototransduction cascade (PIP₂ cleavage^1,2,4,101^) triggers photoreceptor microsaccades at the point of quantum bump production, refractory quantal sampling is morphodynamically coupled to the continuous integration of macroscopic voltage responses to transient contrast changes. This intrinsic sampling mechanism accentuates rapid photon-rate fluctuations, enhancing the resolvability of moving objects and fast environmental light changes - prerequisites for synaptic high-frequency jumping (**Figure 4**).

Thus, beyond stimulus clipping - by occluders (**Supplementary Fig. 24b**), the intraocular aperture (**Supplementary Fig. 24c**), and photoreceptor microsaccades (**Supplementary Fig. 24e**) - refractory quantal sampling further sharpens photoreceptors’ macroscopic voltage responses in time^1,4,6^, improving the resolvability of environmental contrast changes and moving objects. As the number of absorbed photons increases with light adaptation, the signal-to-noise ratio of the morphodynamic neural superposition system also improves^6,13,36^. Consequently, compound-eye hyperacuity increases, as evidenced by the ultraprecise responses of R1-R6 photoreceptors to narrowing 2D gratings (**Supplementary Fig. 15-17**).

**LMC - Over-Complete Morphodynamic Neural Superposition**

In a neural superposition system, light from a small region of visual space is sampled by six photoreceptors (R1-R6) from neighbouring ommatidia, which converge onto shared downstream lamina neurons - large monopolar cells (LMCs: L1-L3) and an amacrine cell (am). These photoreceptors’ receptive fields overcompletely tile the same small visual area (**Supplementary Fig. 24e**: RF1-RF6) and undergo photomechanical shifts that dynamically adjust their spatial and temporal sampling. Due to the compound eye’s curvature and precise ommatidial optics, the visual scene is redundantly covered at high precision. Overlapping R1-R6 inputs are integrated by LMCs, refining the signal through morphodynamic processing. Although R7 and R8 photoreceptors (grey) do not synapse directly onto LMCs, they influence R1 and R6 via electrical coupling through gap junctions^11,12^, modulating the output.

**Supplementary Fig. 24e-f** show how morphodynamic neural superposition enhances LMC responses as the same two flies traverse the overcompletely tiled receptive fields, either unoccluded (**fi**) or occluded by the tree trunk (**fii**). **Supplementary Table 12** summarises intrinsic and extrinsic factors that jointly enhance compound-eye spatiotemporal resolution, enabling hyperacute vision.

Since rhabdomeres shift their x, y, and z positions within the ommatidium - within morphodynamic limits - during these microsaccadic movements, we modelled how such positional changes affect their photon absorption profiles and receptive field shapes relative to the ommatidial lens. These results are detailed in **Section IV.8**: **The Optical Models for Photoreceptor Receptive Fields Depend on Rhabdomere Position** and **Section IV.9**: **Photomechanic Microsaccades Narrow and Move Photoreceptor Receptive Fields**.

**Supplementary Table 12. Factors that enhance compound eyes’ hyperacute vision and predictive coding**

| **Process** | **Description** | **Figures & References** |
| --- | --- | --- |
| Image motion | Self-induced (e.g., from the animal’s own movements) or environmental motion. In animal vision, spatial information is encoded by photoreceptors as parallel temporal sequences of light-contrast changes. | **Supplementary Fig. 24**;  Ahissar & Arieli, 2001^102^  Juusola et al., 2017^1^  Rucci et al., 2025^103^ |
| Spatiotemporal acuity = receptive field $\boldsymbol{\bigotimes}$ impulse response | A photoreceptor’s ability to resolve moving stimuli is determined by the convolution of its receptive-field properties and its impulse response (how quickly it can respond to light changes). Acuity improves when the receptive field narrows or when the photoreceptor’s voltage response becomes faster. | Srinivasan &  Bernard, 1975^46^  Juusola & French 1997^47^  Juusola et al., 2017^1^  Kemppainen et al., 2022a^4^ |
| 3D structure of the world - environmental occlusions | Objects in a three-dimensional environment overlap in the visual field. Self-induced or environmental motion intermittently occludes reflected or emitted light, dynamically sharpening the light-intensity time series sampled by photoreceptors (the “slit-effect.”) | **Supplementary Fig. 24a** |
| Photoreceptor microsaccades - *axial movement* | Photomechanical axial motion of the rhabdomere away from the lens dynamically narrows the photoreceptor’s acceptance angle. This morphodynamic narrowing reduces spatial blur, enabling finer visual detail to be sampled. Although produced by a different physical mechanism, this receptive-field narrowing has a similar effect on temporal information capture as the “slit effect” caused by a physical object occluding the receptive field (see above).  **Supplementary Fig. 30** shows how the receptive fields of R1-R7/8 photoreceptors narrow as the rhabdomere moves to a more distal position. | **Supplementary Fig. 30**  Juusola et al., 2017^1^  Kemppainen et al., 2022a^4^  Kemppainen et al., 2022b^3^ |
| Photoreceptor microsaccades - *lateral movement* | **With stimulus motion**: When the rhabdomere moves in the same direction as a moving stimulus, the photoreceptor integrates over a longer period, improving resolution.  **Against stimulus motion**: When moving opposite to the stimulus, light changes become more transient; microvillar refractoriness enhances these transients, increasing acuity.  **Perpendicular motion**: No enhancement of photoreceptor output occurs when stimulus motion is perpendicular to rhabdomere movement.  **Supplementary Fig. 24d** shows how photoreceptor microsaccade motion - either with or against moving dot stimuli - alters the photoreceptor’s voltage response waveform. | **Supplementary Fig. 24d**  Juusola et al., 2017^1^  Kemppainen et al., 2022a^4^  Kemppainen et al., 2022b^3^ |
| Mechanical coupling of ommatidial R1-R7/8 photoreceptors | Inside an ommatidium, activation of one photoreceptor induces a microsaccade that mechanically pulls neighbouring photoreceptors into motion. As a result, a neighbouring photoreceptor that would not otherwise have been stimulated can be shifted to sample the same light stimulus, supporting predictive coding. | Kemppainen et al., 2022a^4^ |
| Intraommatidial aperture | Pigment and cone cells around the crystalline cone form an aperture aligned with the rhabdomere tips. Microsaccadic rhabdomere movements also shift this aperture, but with a slight phase delay. This movement clips the incoming light beam, producing a dynamic occlusion that sharpens (see the slit-effect” above) the light-intensity changes reaching the photoreceptors.photoreceptors. | **Supplementary Fig. 24cii**;  Juusola et al., 2017^1^  Kemppainen et al., 2022a^4^ |
| Intracellular photoreceptor pupil mechanism | After bright light exposure, tiny pigment granules within the cytoplasm of R1-R7/8 photoreceptors migrate toward their rhabdomeres, surrounding each rhabdomere with a light shield in about 10 seconds. This intracellular pupil mechanism is expected to narrow each photoreceptor’s receptive field (relative to its dark-adapted state) by approximately 10%. | Franceschini & Kirschfeld, 1976^104^  Juusola et al., 2017^1^ |
| Refractory quantal sampling | Refractory quantal sampling of photons by tens of thousands of microvilli, and of histamine molecules by thousands of LMC spines, accentuates contrast changes in the voltage response transients. This, in turn, drives high-frequency synaptic jumping and improves spatiotemporal resolution. | E.g. **Figure 4**;  Juusola et al. 2017^1^  Song & Juusola, 2014^34^ |
| Morphodynamic neural superposition  system | Neural superposition with overcomplete tiling: R1-R6 photoreceptors from adjacent ommatidia sample partially overlapping areas, dynamically shifting receptive fields through microsaccades. R1-R6 photoreceptors feed their respective information capture to the same LMC, where spatiotemporal resolution is enhanced by this morphodynamic sampling and processing. | **Supplementary Fig. 21e-f** |
| Binocular overlapping receptive fields of the left and right eye | Overlapping receptive fields between the two eyes allow the same object to be sampled by different photoreceptors from slightly different angles. The microsaccades of left- and right-eye photoreceptors are mirror-symmetric: when a moving object crosses an overlapping binocular receptive field, photoreceptors in one eye move with the stimulus, while those in the other move against it. When the positions of these photoreceptors are known (from the left and right eye sampling matrices), the resulting phase difference in encoding directly reveals the object’s distance. This binocular redundancy further enhances hyperacuity and supports predictive coding of object motion in depth. | **Supplementary Fig. 24d**  Kemppainen et al., 2022a^4^ |

**II. 9 Equivalent Encoding Principles: Hyperacuity vs. High-Frequency Jumping**

Although their mechanisms differ, the slit-effect from occlusion (underlying hyperacuity) and the synaptic clipping of saccadic photoreceptor signals (underlying high-frequency jumping) are mathematically and functionally analogous. In both cases, motion-driven spatial changes are transferred almost instantaneously into higher-frequency carrier bands, improving resolvability and minimising delays. **Supplementary Fig. 25** schematically illustrates these parallels.

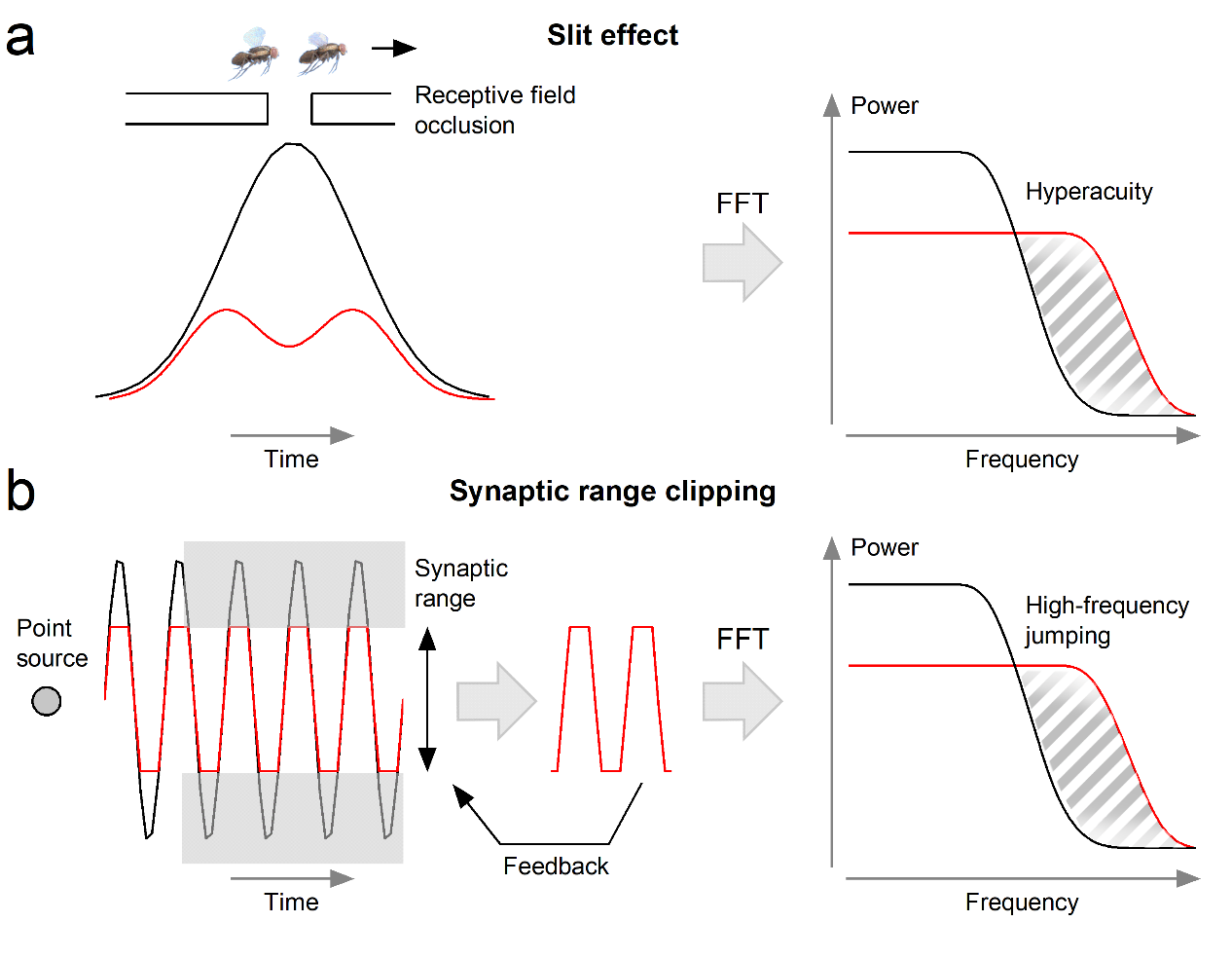

**Supplementary Fig. 25. Equivalent encoding principles: hyperacuity via the slit-effect and high-frequency jumping via synaptic clipping.**

(**a**) Slit-effect and hyperacuity. Two objects (flies) moving across the visual field are separated by less than the diffraction limit. When viewed through an unobstructed receptive field (Gaussian profile, centre), the signals merge. However, partial occlusion by a natural object (e.g., tree trunk in **Supplementary Fig. 24a**) or by the intraommatidial aperture narrows the effective receptive field into a “slit,” sharpening the temporal signal. In the frequency domain (FFT, right), this clipping shifts information into higher-frequency carrier bands, thereby enabling hyperacuity.

(**b**) Synaptic clipping and high-frequency jumping. Photoreceptor voltage responses to saccadic light fluctuations (from a point source, left) undergo synaptic transfer to LMCs, where the limited synaptic operating range clips the signal (centre). This feedforward-feedback interaction transforms smooth waveforms into sharper transients, redistributing information into higher-frequency bands (FFT, right). The outcome parallels the slit-effect: motion-driven spatial changes are encoded at higher frequencies, improving resolvability and minimising delays.

**III. In vivo high-speed optical imaging of photoreceptor microsaccades**

**III.1. High-Speed Imaging of Photoreceptor Microsaccades**

We recorded rapid, light-induced displacements of dark-adapted *Musca* rhabdomeres - photoreceptor microsaccades - in response to light flashes using high-speed infrared imaging, which does not activate the photoreceptors (**Supplementary Fig. 26a**). In this method, a water droplet was placed between the objective and the compound eye to neutralise the cornea, enabling direct visualisation of the R1-R7/8 rhabdomere tips^1,105^ inside ommatidia. This approach, previously applied to *Drosophila*^1,3,4^, revealed that both species exhibit photoreceptor microsaccades.

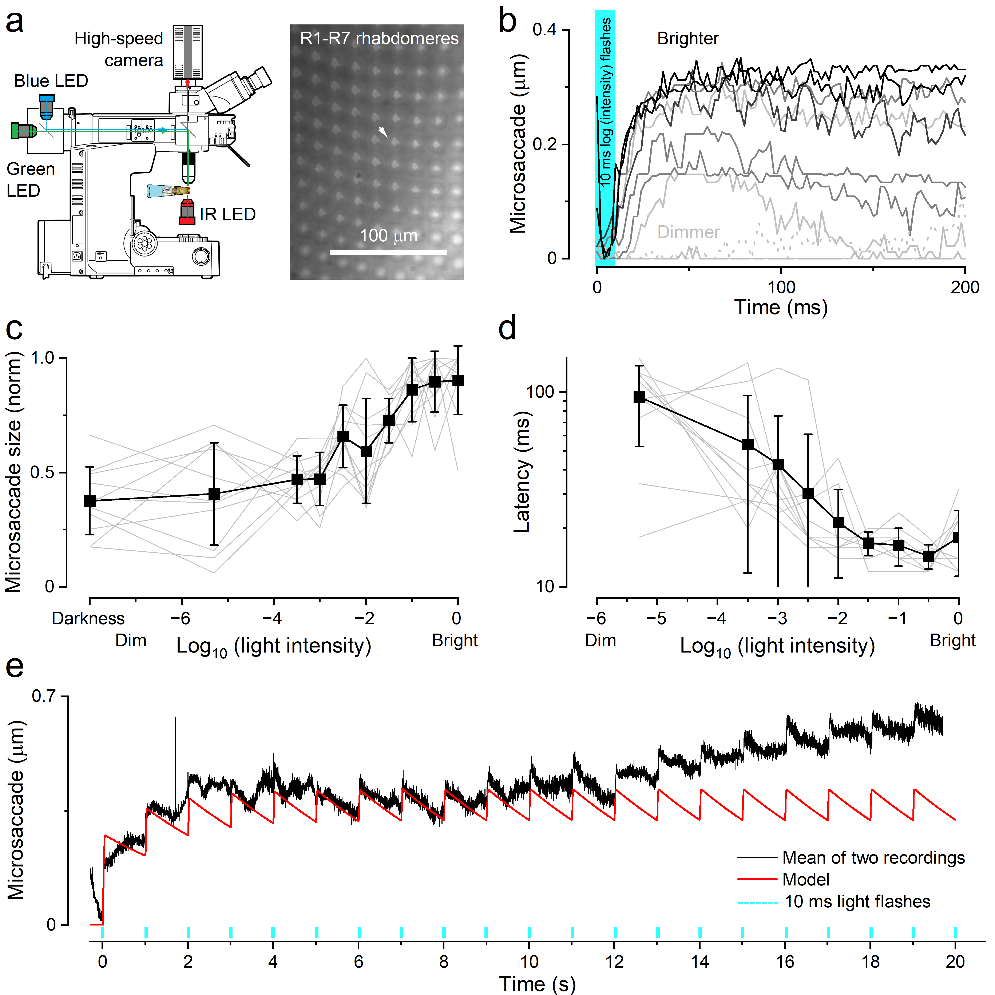
**Supplementary Fig. 26. Light flashes evoke photoreceptor microsaccades in dark-adapted *Musca* compound eyes, rapidly shifting rhabdomeres**.

(**a**) High-speed videography setup for imaging light-flash-induced rhabdomere movements - photoreceptor microsaccades. An opening in the back of the fly’s head cuticle allows infrared (IR) light to pass through the rhabdomere waveguides, visualised by the camera. Blue and green LED flashes, delivered through the objective and separated by a dichroic mirror, stimulate the photoreceptors and trigger microsaccades. Inset: Rhabdomere tip patterns from observed ommatidia. White arrow: similar to *Drosophila*^3,4^, R1-R2-R3-rhabdomere axis sets the local microsaccade movement direction.

(**b**) Microsaccade responses to 10-ms green-blue light flashes (green bar), recorded across a wide intensity range using neutral density filters (−5.3 to 0 log_10_ units). Dotted line: no-flash control.

(**c**) Microsaccade amplitude increases with flash intensity, following a sigmoidal activation curve - consistent with a photomechanical origin driven by phototransduction^1,2,101^.

(**d**) Microsaccade latency - measured as the delay to the point of maximal derivative - decreases with increasing flash intensity. Notably, responses to bright flashes begin well within the 10-ms stimulus duration (see panel **b**), closely matching the rapid rise of corresponding intracellular voltage responses (*cf*. **Supplementary Fig. 1a**). Grey lines: individual rhabdomere responses; black squares: mean ± SD.

(**e**) Repeated 10-ms flashes elicit repeated microsaccades. As in Drosophila, repeated stimulation leads to a gradual buildup of background tension, from which new microsaccades emerge. Black trace: mean response from two recordings; red trace: simulated response from the movement model (**Section IV.9ii**).

Approximately 4-8 ms after a bright 10 ms flash (**Supplementary Fig. 26b**), rhabdomeres facing the central light source began to shift rapidly in their genetically determined direction - along the R1-R2-R6 rhabdomere axis. As in *Drosophila*^3,4^, this axis is slightly rotated within each ommatidium relative to its neighbours. Consequently, the directional shifts observed in *Musca* were similar to those reported for *Drosophila* rhabdomeres^3,4^. The movements increased in amplitude (**Supplementary Fig. 26b-c**) and speed (**Supplementary Fig. 26d**) with flash intensity, reaching their intensity-dependent maxima (0.35-0.5 µm) within 30-50 ms before gradually returning to baseline. The microsaccades were reliably evoked by regularly timed flashes (**Supplementary Fig. 26e**).

According to morphodynamic information processing theory^27^, microsaccadic sampling should adapt to optimise spatiotemporal acuity. Consistent with this prediction, *Musca* photoreceptor microsaccades are faster and smaller than those in *Drosophila* (*cf*. **Figure 5** in the main paper), reflecting *Musca*'s faster visual lifestyle and its more densely faceted compound eyes (*i.e*. smaller interommatidial angles). Given that the mean tip diameter of R1-R6 rhabdomeres is 1.39 µm (**Supplementary Fig. 26a**), a bright flash displaced the tip by approximately one-quarter of its width.

Although these microsaccades are driven by phototransduction reactions - specifically PIP₂ cleavage^1,4,101,106^ - and show a similarly fast initial rise as the corresponding intracellular voltage responses (*cf*. **Supplementary Fig. 1a**), they reach their peak amplitudes and return to baseline more slowly, especially in dark-adapted eyes^3,4^. This overall slower waveform - and the trial-to-trial variability observed (**Supplementary Fig. 26e**) - is likely due to damping and inertia in the surrounding retinal tissue, including activity and adaptation state-dependent tension in the intraocular muscle system^27^.

Recording photoreceptor microsaccades in vivo in *Musca* is challenging due to large intraocular muscle contractions driven by “clock spikes” from the brain^37^. These movements can obscure the smaller, faster microsaccades and shift the imaged retina out of the microscope’s focal plane. Such contractions often occur several seconds apart. To capture usable data, we timed recordings to coincide with brief periods of retinal steadiness when muscle activity had subsided. Any recordings contaminated by large movements were excluded during offline analysis.

1. **Preparation**

- Flies with intact compound eyes were positioned atop a cut pipette tip, with their protruding heads securely fixed to the tip edge using beeswax.
- The head was tilted rightward, and a small window was cut in the posterior cuticle behind the left eye. This opening allowed back-illumination of the rhabdomere tips with low-power IR light, causing minimal eye damage.

2. **Illumination & Imaging**

- *Eye positioning*: The pipette-mounted fly was secured on a Sensapex micromanipulator (Finland) for precise alignment.
- *IR backlighting*: A 740 nm LED (Cairn OptoLED, UK), filtered at >720 nm, illuminated the retina through the cuticular window from below.
- *Objective*: A 40× water-immersion objective (Zeiss C Achroplan NIR 40×/0.8 W) was used on an Olympus BX51 upright microscope, with a water droplet for immersion.
- *High‑speed imaging*: Images were recorded at 500 Hz using an Andor Zyla camera (UK) through a 600 nm dichroic mirror, which transmitted the 740 nm light through the R1-R7/8 rhabdomeres while minimising rhodopsin activation.

3. **Light‑Flash Stimulation**

- *Activation LEDs*: Blue (470 nm) and green (545 nm) LEDs (Cairn OptoLED, UK) delivered flashes via a 495 nm dichroic mirror.
- *Control system*: A custom MATLAB program, interfaced with National Instruments data acquisition hardware, controlled flash timing and intensity.

Under these conditions, we observed photoreceptor microsaccades (**Supplementary Fig. 26**) - ultrafast photomechanical displacements of rhabdomeres that transiently narrowed and shifted each receptor’s receptive field, morphodynamically shaping its angular sensitivity.

**III.2 Photoreceptor Microsaccade Dynamics**

1. **Flash-Evoked Lateral Photoreceptor Displacement**

- We delivered a 10 ms flash (blue + green LEDs) and recorded rapid rhabdomere shifts with high-speed microscopy (see **Section III**: **Supplementary Fig. 26b**).
- Rhabdomere displacement during these microsaccades grew with flash intensity, peaking at 0.21 ± 0.09 µm (n = 11; **Supplementary Fig. 26c**).

2. **Latency and Speed**

- To find movement onset, we smoothed each trace with a 40-ms filter and extracted the maximum derivative following the flash onset (Fig. **Supplementary Fig. 23b**, **d**).
- Under bright flashes, the microsaccades reached their maximum velocity at 14 ± 2 ms (n = 10).
- Lower intensities delayed this peak, confirming slower microsaccades.
- This direct light-dependency is consistent with microsaccades being driven by phototransduction reactions.

3. **Adaptation Dynamics**

- We repeated 10 ms flashes every second at 0.5 log unit intensity (minimising artefacts) and tracked peak displacement over trials (**Supplementary Fig. 26e**).
- After an initial decline, movements stabilised at ≈ 0.05 µm in steady-state.

4. **Artefact Rejection**

- Intraocular “clock-spike” muscle twitches^37^ typically interfere with recordings, often shifting the whole retina out of frame.
- We, therefore, analysed only the recording series with minimal muscle interference (n = 2).

5. **Comparison to *Drosophila***

- Time-to-Peak: *Musca* 30-50 ms vs. *Drosophila* ≈100 ms
- Latency: *Musca* <5 ms vs. *Drosophila* 10-30 ms
- Amplitude: *Musca* ~0.2 µm vs. *Drosophila* up to >1 µm)^1^.
- Overall, *Musca* photoreceptor microsaccades are ~3-5× faster than in *Drosophila*.
- A *Musca* microsaccade (duration ~20 ms duration; amplitude = 0.4 µm; **Supplementary Fig. 26**) speed is approximately 37.2°/s (1.86°/µm (**see Section IV.8**) × 0.4 µm/20 ms). For comparison, in *Drosophila*, equivalent microsaccade speed^4^ is 45°/s (3°/µm × 1.5 µm/100 ms). This suggests that the spatiotemporal resolution of fly photoreceptors matches rather well, suggesting that the underlying phototransduction reaction cascade operates at similar speeds.

**IV. Modelling Morphodynamic Neural Superposition System with Adaptive Optics**

We developed a novel stochastic, multiscale modelling framework of the *Musca* compound eye’s early visual circuits, which we term the **morphodynamic neural superposition system model**. Meticulously built from the bottom up, it integrates biophysically realistic subcomponents to test and validate both qualitative and quantitative aspects of adaptive neural computation. This framework provides mechanistic insight into how quantal, refractory information sampling and processing have co-evolved with behaviour to maximise visual information.

This supplementary section details how the model - integrating the refractory, quantal nature of microvillar and synaptic information capture with rapid photomechanical interactions in superposition wiring - closely replicates the response dynamics of both photoreceptors and LMCs. Notably, this holistic approach reproduces intracellular voltage responses, adaptive dynamics, saccadic-behaviour-driven high-frequency jumping, response variability (noise), and information transmission rates across all tested stimuli - without free parameters.

Consistent with our recordings, the model shows how photon sampling and neurotransmission co-adapt during rapid movements to generate precise environmental representations. Rather than passively processing input, houseflies actively shape their visual world through rapid, self-induced manoeuvres - producing saccadic contrast bursts that drive *synaptic high-frequency jumping*, which abolishes transmission delays and facilitates efficient coding, hyperacute vision, and neural synchronisation, tightly linking behaviour to perception.

Thus, this modelling framework explains - and enables mechanistic investigation of - how adaptive form-function interactions embedded in the morphodynamic superposition system support ultrafast, efficient, and reliable early visual processing, revealing a highly effective neural strategy for high-speed, predictive coding.

**How we validated the model**

In the full morphodynamic neural superposition model, the vast majority of parameters are fixed by empirical measurements rather than freely fitted. Specifically, all static optical parameters, including ommatidial geometry, photoreceptor receptive-field structure (via Fourier beam propagation), as well as the photomechanical motion of R1-R6 receptive fields (**Supplementary Notes III**), are fully constrained by anatomical and optical measurements (**Supplementary Notes II**).

Similarly, the phototransduction model for each R1-R6 photoreceptor is derived directly from intracellular recordings. Each photoreceptor is modelled separately to reflect measured differences in rhabdomere size and microvillar number (**Supplementary Notes III**). The stochastic quantum bumps that generate the macroscopic voltage response are produced using empirically constrained distributions of quantum bump size, latency, and refractoriness, obtained from established systems and quantum-bump analyses of recorded photoreceptor signal and noise^1,4-6,13,27,34,36,40,42,75,107^.

Parameter adjustment during model construction followed a constructionist, stage-wise approach rather than global fitting. At each stage, model predictions were compared quantitatively with recorded photoreceptor and LMC responses, and the residual error dynamics were analysed in relation to known physiological processes and their published time constants. This procedure constrained synaptic output and feedback structure based on physiological plausibility rather than numerical optimisation.

Because the model is explicitly stochastic and refractory, we validated its reliability by computing signal-to-noise ratios and information transfer rates at each sampling and processing stage and comparing these directly with corresponding experimental measurements from photoreceptors and LMCs. The final model reproduces not only mean responses, but also response variability and frequency-dependent information transfer across a wide range of stimulus conditions.

Importantly, model behaviour - including bandwidth expansion, high-frequency jumping, and delay minimisation - proved robust to moderate parameter variation and emerged consistently across stimulus statistics, light intensities, and receptive-field configurations tested. Thus, at each stage of model building, sensitivity and robustness are intrinsically assessed through signal-to-noise and information-theoretical comparisons. Full parameter documentation is provided in **Supplementary Tables 13-16**.

**IV.1. Modelling Overview: Bigger Picture**

We have previously published a stochastic *Drosophila* photoreceptor model capable of generating realistic macroscopic voltage responses to a wide range of light stimuli, using fixed parameters calibrated to the tested mean light intensity^1,28,34,40^. This model consists of four modules:

1. **Random photon absorption model**

Photon absorption in microvilli was modelled as a Poisson process: light intensity time series, treated as photon fluxes, were absorbed by a population of 30,000 microvilli. Each microvillus absorbed photons discontinuously, producing a stochastic series of absorption events. Rarely, a microvillus would absorb more than one photon within a single 1 ms interval.

2. **Stochastic model of the phototransduction cascade in a single microvillus**

The signalling pathway was represented as a series of unidirectional reactions involving only unimolecular or bimolecular reactants. Each absorbed photon activated the photopigment rhodopsin (metarhodopsin, M*), triggering the phototransduction cascade. This led to the opening of light-gated ion channels and the influx of Ca²⁺ and Na⁺ ions, generating a light-induced current (LIC) quantum bump with a unique waveform.

3. **Integration of LIC across the rhabdomere**

Quantum bumps with varying waveforms from 30,000 microvilli were summed to produce a macroscopic light current response.

4. **Hodgkin-Huxley model of the photoreceptor membrane**

The *Drosophila* R1-R6 photoreceptor membrane was modelled as an electrical circuit: the membrane as a capacitor; voltage-gated channels as voltage-regulated conductances; leak channels as fixed conductances; and reversal potentials for different ion species as DC batteries.

Our current quantal sampling model for a *Musca* photoreceptor is **a probabilistic four-parameter model**^4^. It simplifies the detailed biochemistry of quantum bump generation by sampling from experimentally derived or theoretically estimated distributions. The key parameters are:

- Number of microvilli (~54,000 independent sampling units per rhabdomere)
- Quantum-bump waveform (average shape and duration at a given ambient intensity)
- Latency distribution (timing jitter before each bump)
- Refractoriness distribution (how long each microvillus stays unresponsive after a bump)

Although both models accurately simulate macroscopic voltage responses - with highly similar waveforms, signal and noise dynamics, and information transfer rates compared to intracellular recordings - the new model is vastly more efficient and does not require a computing cluster. Even on a standard desktop, it runs approximately 1,000-10,000 times faster. *This computational advantage enables us to integrate six individual photoreceptor models - each with slightly different microvilli counts (reflecting natural variation among R1-R6 cells that synaptically pool their signals to LMCs) - into the* **new morphodynamic neural superposition system model**.

However, unlike the previous model - where each photon absorption event, the activated microvillus, and the resulting quantum bump could be explicitly tracked and analysed - the new four-parameter model sacrifices this level of granularity. While it preserves the overall statistical properties of quantal sampling, individual events are no longer traceable in post hoc analysis.

**IV.1.i. Limits of the Morphodynamic Neural Superposition System Model**

The morphodynamic neural superposition system model adapts dynamically to visual input. The number of quantum bumps - unitary samples triggered by photon absorption or histamine-molecule binding - varies according to refractory sampling principles, as do the microsaccadic (morphodynamic) and synaptic feedback loops. However, the quantum bump waveforms and amplitudes - both in photoreceptors and LMCs - are fixed to their mean values at a given ambient light level, based on noise analyses from experimental recordings. While this simplification reduces biophysical realism, it does not pose a major limitation, for the following reasons:

**1. Realistic Neural Adaptability Without Bump Amplitude Modulation**

Although quantum bump waveform and size do not adapt dynamically, the model captures the core adaptive properties of the morphodynamic neural superposition system:

- **Refractory sampling** and **feedforward-feedback synaptic coupling** dynamically co-adjust the macroscopic response waveforms of photoreceptors and LMCs, keeping them within their respective biophysical voltage and frequency ranges across varying light intensities and temporal patterns.
- **Photoreceptor-LMC synapses** amplify LMC voltage transients, intrinsically implementing *high-frequency jumping* to sharpen saccadic contrast signals with minimal delay - shifting LMC signals to higher carrier frequencies and closely matching the dynamics observed in intracellular recordings.
- LMCs, in particular, exhibit ultrafast adaptation, dynamically adjusting their response range even without bump amplitude modulation.
- As a result, the system reliably tracks rapid stimulus changes - such as edges, object motion, or self-generated movement - under natural viewing conditions.

**2. High Information Rates Matching Recordings Despite Simplifications**

- The simplified stochastic sampling model retains the main sources of neural noise observed in photoreceptors and LMCs.
- Photoreceptors show additional low-frequency noise caused by rhabdomere microsaccades, which is replicated in the model via simulated microsaccadic photoreceptor movements.
- Simulated photoreceptor and LMC responses achieve high information rates across a broad frequency range, closely matching the encoding performance observed in intracellular recordings under equivalent light stimuli.
- Na⁺ channel-driven action potentials are not required for encoding; graded potentials suffice for high-throughput, reliable visual signalling.

**3. Biophysical Evidence Supports Model Assumptions**

- Optical stimulation (**Section III**) and anatomical measurements (**Section II**) show that the receptive field centre of an R1-R6 photoreceptor shifts laterally by ~~1.86°/µm during photomechanical rhabdomere movement (microsaccade) in response to a bright light flash.
- Receptive half-field widths (acceptance angle, $\Delta\rho$) vary across photoreceptor types: R1 has a half-width of ~2.62°, while R7 and R8 are narrower (~2.1°).
- Rhabdomere microsaccades are fast - reaching peak derivative at 14 ± 2 ms - but small in amplitude (~0.21 ± 0.09 µm), and show adaptation during repeated pulse stimulation (**Supplementary Fig. 26**).
- These small movements contribute to photoreceptor sampling noise, especially during small-dot or bursty stimuli, leading to a reduction in signal-to-noise ratio, ${SNR}_{PR}(f)$, especially below 10 Hz.

**4. Adaptive Feedback Mechanisms at the Synapse**

- As in *Drosophila*^8,16^, synaptic feedback from *Musca* LMCs to photoreceptors functions as an adaptive gain control mechanism.
- Nikolaev et al.^54^ showed that disrupting this feedback reduces ${SNR}_{LMC}(f)$ across all frequencies, leading to lower information transfer rates.
- This feedback dynamically adjusts the photoreceptor output range and shapes LMC responses, helping to maintain robust signalling under rapidly changing visual conditions.

**5. Neural Superposition Supports Spatial and Temporal Integration**

- R1-R6 photoreceptors inside an ommatidium differ slightly in microvilli count and optical alignment (see **Section II**), resulting in subtle variations in their macroscopic voltage responses, even when stimulated by the same spatiotemporal pattern.
- Due to differences in lens-to-rhabdomere geometry in the larger *Musca* eye, the receptive fields of photoreceptors from adjacent ommatidia - pooled into the same neural superposition system - do not overlap perfectly, but their individual half-widths, $\Delta\rho$, form a rosette pattern (*cf*. **Supplementary Fig. 24e**; **Figure 1**).
- Because each ommatidium is slightly rotated relative to its neighbour, photomechanical rhabdomere movements shift the receptive fields of R1-R6 in different directions when exposed to the same light stimulus (*cf*. **Supplementary Movie 2**). This morphodynamic summation enhances the spatial resolution (acuity) of LMC responses through precise synaptic integration.
- Additionally, differing waveguide properties and time constants across R1-R6 and R7/R8 photoreceptors further enrich spatial sampling diversity.
- Together, these features increase the robustness of the visual system and improve its capacity for predictive processing across both space and time.

Together, these powerful adaptive mechanisms ensure that the system maintains naturalistic high-fidelity visual encoding - even without stochastic dynamic quantum bump amplitude modulation - validating the model’s usefulness for simulating rapid, naturalistic visual processing.

**IV.1.ii. Morphodynamic Neural Superposition System Model Predicts Experimental Results**

This section highlights key examples where the morphodynamic model replicates experimentally observed photoreceptor and LMC responses. We organise these into validated phenomena, functional adaptations, and emerging predictions. Analytical explanations are detailed in subsequent sections.

**1. Core Functional Predictions Verified by the Model**

*1.1 High-Frequency Jumping and Delay Reduction in LMCs* (e.g. **Figure 1c**, **Figure 2d**)

- Example: 10 ms light pulse.
- Model shows synaptic probability $P(t)$ produces transient, biphasic LMC response.
- Validates: Fast adaptation through synaptic sampling.
- Confirms previous in vivo findings^13^.

*1.2 Light-Level Dependent Adaptation in LMCs* (tested - not shown)

- Examples:
- Flashes at two ambient light levels.
- Bursty and GWN stimuli at different light levels.
- Findings:
- Dim light → lowpass behaviour.
- Bright light → differentiated, frequency-enhanced responses.
- Model shows scalable adaptation through synaptic sampling.
- Limitation: No quantum bump adaptation in photoreceptors; simulated range smaller than in vivo.

**2. Emergent Properties and Model Robustness**

*2.1 Neural Superposition and Natural Variation* (**Supplementary Fig. 4b**)

- Example: Slightly different R1-R6 responses to the same stimulus.
- Cause: Overlapping but non-identical receptive fields.
- Confirms: Natural variability across ommatidia^1^.

*2.2 Feedback-Driven Saturation Prevention* (**Supplementary Fig. 32f**)

- Example: Two 10 ms light pulses with and without feedback.
- Model shows enhanced synaptic probability, $P(t)$, peak and greater LMC voltage response with feedback.
- Feedback resembles derivative of LMC response → regulates inhibitory and excitatory synaptic loads.

*2.3 Microsaccades and Refractory Noise Modulation* (**Figures 4c** and **4f**)

- Examples:
- 200 Hz bursty stimulus with vs. without microsaccades.
- Repeated pulse stimulus.
- Findings:
- Microsaccades introduce low-frequency noise.
- Drop in low-frequency SNR and information rate.
- Consistent with findings from *Drosophila*^1^: slight stimulus-dependent low-frequency response variability.

*2.4. Energy Efficiency and Energy-Information Trade-off* (tested through simulations - not shown)

While direct metabolic recordings at the relevant temporal and spatial scales are not currently available in the literature, energetic costs in fly photoreceptors can be quantitatively estimated from well-established physiological principles. In previous work, we used biophysically realistic photoreceptor models to calculate ATP consumption under different stimulation regimes, including Gaussian white noise (GWN) and naturalistic, behaviourally relevant input statistics^34^. In these models, ATP usage is dominated by the activity of the Na⁺/K⁺ exchanger required to counterbalance light-induced ionic fluxes, and thus depends strongly on the mean depolarisation level of the photoreceptor membrane^108^.

Under GWN stimulation, photoreceptors are driven into a sustained, high mean depolarisation state (typically ~20–30 mV above dark resting potential), which imposes a large and continuous ionic load and consequently high ATP demand. By contrast, under bursty, saccade-like stimulation, mean depolarisation remains substantially lower (typically ~10–15 mV above resting potential), while preserving high temporal responsiveness through transient, high-frequency events. As a result, maintaining responsiveness under bursty stimulation is predicted to require substantially less ATP than under GWN, despite comparable or higher information throughput. For *Musca* photoreceptors, comparable depolarisation regimes are observed in the present data (e.g. **Supplementary Fig. 12b**).

Accordingly, our use of the terms thermodynamic constraints and energy-efficient coding refers to mechanistically motivated predictions derived from established ionic and biophysical relationships, rather than to new metabolic measurements.

- Examples:
- Stochastic photoreceptor model with/without microvilli refractoriness.
- Stimulus types: bursty vs. broadband GWN.
- Findings:
- Maintaining graded voltage responses consumes ATP (as proposed earlier^108^).
- Refractoriness reduces membrane depolarisation by ~15-20 mV (see also^34^).
- ATP consumption can drop by >40% (see also^34^).
- Energy cost per bit can fall by ~35% (see also^34^).
- Encoding bursty stimulation is energetically cheaper than encoding GWN, which drives photoreceptors into a high-voltage, poor-SNR regime.
- GWN underestimates neural performance and overestimates energy cost, calling into question early proposed energy/information trade-off trends^109,110^.
- In microvillar photoreceptors of different insect species, variation in sampling unit (microvillus) number and quantum bump statistics (size, latency, refractory distributions) produces distinct response waveforms to the same light stimulus, leading to different information transfer rates and energy consumptions, as confirmed by simulations matching recordings^6,28,34,40^.

*2.5. Refractoriness, Information Transfer and Saliency*

- Examples:
- Simulation vs. recorded data.
- Responses to sudden light increments and decrements (see also^1,6,35^).
- Findings:
- Refractoriness increases initial changes in neural (quantal) sample rate after large light transitions (see also^1,6,35^).
- Early on-responses (increments) have higher SNR due to more sampling units (microvilli/synaptic spines) being active.
- Early off-responses (decrements) are larger because more units remain quiescent initially.
- Polarity not critical - larger, faster transitions carry more information.
- Information transfer peaks at large dim-to-bright or bright-to-dim transitions, then declines with adaptation (cf. Juusola & de Polavieja^35^, Fig. 4).
- Adaptive trends invariably lead to the underestimation of neural information (**Supplementary Fig. 9**).
- *Because of refractory quantal sampling, the first response is the most salient and carries the most information*.

**3. Higher-Order Visual Properties Captured by the Model**

*3.1 Spatial High-frequency jumping*

- Example: Wide bars or small dots sweeping across receptive fields (**Supplementary Fig. 24**).
- Observation: Photoreceptors and LMCs adapt during the stimulus motion.
- Stronger with flash-like (fast onset) stimuli.
- Limitation: No quantum bump adaptation; the model assumes constant adaptation to ambient light level.
- Including dynamic quantum bump size adaptation may increase the resolvability further.

*3.2 Hyperacuity and Receptive Field Matching*

- Example: Bars or dots moving over receptive fields with matched ommatidial motion (~50 ms rise phase) (**Supplementary Fig. 24**).
- Observation: LMC responses show fine oscillations depending on receptive field half-widths.
- Modelling challenges: the synaptic probability $P(t)$ operating range represents a sensitive dynamic balance between feedforward and feedback loads; responses may overshoot or undershoot.

**4. Predictions Not Yet Fully Tested**

*4.1 Time-Scale Invariance*

- Example: The same light intensity time series stimulus presented at different speeds^35^.
- Prediction: Multi-scale adaptation makes the amplitude and timing of LMC responses show self-similarity, similarly paced patterning (when normalised over time)^27^.
- Not yet validated in experiments.

*4.2 Whitening of LMC Responses*

- Example: Repeating the same stimulus three times.
- Observation: Burst responses adapt within ~1 second; in vivo adaptation is slower. This is likely due to quantum bump size reduction, which is expected to reflect gradual calcium accumulation in the cell body.
- Notes: Simulation time window (2 s) longer than prior studies (e.g., Zheng et al^16^: 0.5 s).
- Hypothesis: LMC response flattening occurs only within the narrow photoreceptor voltage range.

**5. Summary of Model Assumptions**

- No dynamic adaptation of quantum bump size was included in the simulations.
- Simulations assume the system is in a relatively steady state, mimicking intracellular voltage responses to a 2-second light stimulus pattern after three repetitions.
- Synaptic sampling $P(t)$ was used to model dynamic adaptation and transient shaping.
- As a control, photoreceptor microsaccades were not explicitly modelled in all cases.

**IV.2. Modelling Overview: Multiscale Stochastic Information Processing in Compound Eyes**

The compound eye of *Musca domestica* consists of a matrix of hexagonally arranged, lens-capped units called ommatidia (**Supplementary Fig. 27a**), each housing eight photoreceptors (R1-R8)^87,89,111^. Incoming light, focused by the corneal lens, is guided onto the photoreceptors’ microvillar, light-sensitive regions called rhabdomeres. These structures act as optical waveguides, absorbing photons as they pass through. Each rhabdomere contains approximately 54,000 microvilli - thin, cylindrical membrane protrusions that serve as discrete photon-sampling units (see **Section II**). Each microvillus contains the complete phototransduction machinery, enabling it to convert absorbed photons into discrete electrical signals called quantum bumps^40,101^.

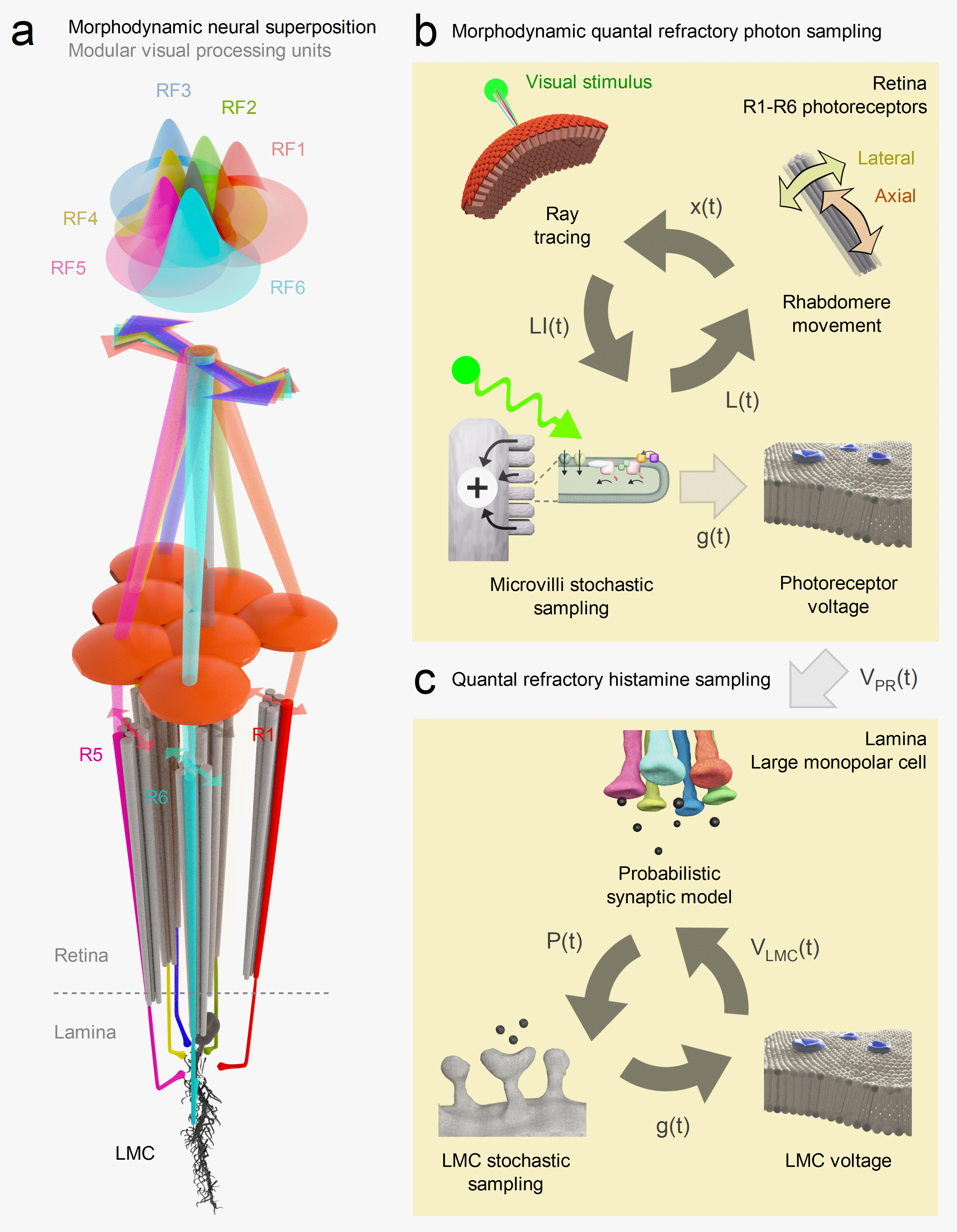
**Supplementary Fig. 27.** **Multiscale morphodynamic modelling of peripheral visual processing in the *Musca* compound eye**.

(**a**) **Morphodynamic neural superposition**. Modular visual processing units in the *Musca* compound eye integrate information via neural superposition, whereby six R1-R6 photoreceptors from different ommatidia (receptive fields RF1-RF6) converge onto shared downstream neurons. Each photoreceptor samples the same point in space from a slightly different angle, with a receptive field that shifts photomechanically in space and time. Due to the eye's curvature and the optical arrangement of ommatidia, these spatiotemporally overlapping inputs form an overcomplete tiling of visual space. Their signals are pooled by large monopolar cells (LMCs) in the lamina, enabling morphodynamic information processing that sharpens neural representations. R7 and R8 photoreceptors (grey) contribute indirectly via gap junctions to R1 and R6 terminals but do not form direct synapses with LMCs.

(**b**) **Morphodynamic quantal refractory photon sampling** - **photomechanical feedback**. The photoreceptor model integrates optical ray-tracing with dynamic rhabdomere motion - comprising lateral and axial microsaccade components - to calculate the effective light input 𝐿(𝑡), shaped by local illumination $LI(t$) and stochastic photon sampling by 35,000-70,000 microvilli (individual photon-sampling units that form the photoreceptor’s light-sensitive part, the rhabdomere). This input generates a quantal, refractory phototransduction response $g(t)$, which drives the photoreceptor membrane voltage $V_{PR}\left( t \right),$via a Hodgkin-Huxley-type model of the photoreceptor plasma membrane.

(**c**) **Quantal refractory histamine sampling** - **synaptic feedback**. At the synapse, a probabilistic model converts presynaptic voltage into a neurotransmitter release probability $P(t)$, which is stochastically sampled by LMC dendrites to generate quantal postsynaptic events. These are integrated by the LMC membrane model to produce $V_{LMC}\left( t \right).$ Feedback synapses from LMCs to photoreceptor terminals close the morphodynamic processing loop from visual input to postsynaptic output.

Like in *Drosophila*, *Musca* photoreceptors R1-R6 (**Supplementary Fig. 27a**), located in neighbouring ommatidia and transmitting visual information to the same lamina cartridge, are optically slightly misaligned, sampling light from similar - but not identical - directions^4,61^. This arrangement, known as neural superposition^61,112,113^, allows each photoreceptor to capture partly overlapping information. These photoreceptors synapse onto shared lamina neurons - large monopolar cells (L1-L3) and amacrine cells (AMs) - via histaminergic, hyperpolarising (inhibitory) synapses^48,114,115^. LMCs and AMs integrate these inputs and relay them to the brain, while also providing tonic depolarising (excitatory) feedback to the same photoreceptors^8,16,54,66,116,117^.

Without this feedback, R1-R6 photoreceptors rest near the potassium reverse potential (-85 to -90 mv), as seen in the temperature-sensitive *shibire^ts^*-mutants at 30 ^o^C, where synaptic transmission in the photoreceptor-LMC pathway is blocked^8^. Similar hyperpolarised resting potentials occur in dissociated photoreceptors lacking all synaptic feedback^8,66^. By contrast, in vivo recordings from intact wild-type flies at 30 °C show that continuous excitatory synaptic feedback raises the resting potential of R1-R6 photoreceptors in darkness by approximately 10-25 mV, stabilising it around -65 to -77 mV^8^.

**Modelling photoreceptors’ morphodynamic quantal photon sampling**

We model photoreceptor optics by simulating photon sampling through ray tracing from a virtual environment^4^ (**Supplementary Fig. 27a**). Individual photoreceptor receptive fields are computed using optical simulations that integrate ray tracing with the Fourier beam method^4^. Additionally, photoreceptor microsaccades - small, rapid photomechanical movements of the rhabdomeres - are modelled based on noninvasively measured microsaccades (see **Section III**) using high-speed infrared microscopy, which does not activate phototransduction^1,3,4^.

Beneath the ommatidial lenses, dynamic changes in light intensity cause photoreceptors to rapidly contract and elongate in and out of their focal plane, and shift laterally, executing a complex piston-like motion. These microsaccades dynamically reshape and reposition photoreceptor receptive fields in response to visual stimuli^1,3,4^. This optomechanical feedback mechanism continuously adjusts photoreceptor alignment and geometry based on local contrast changes, significantly enhancing the spatiotemporal resolution of the compound eye^1,4^. As a result, flies achieve morphodynamic hyperacuity - driven by motion-induced physical sampling - enabling them to resolve visual details far finer than the theoretical limits imposed by the fixed spatial sampling of the ommatidial array (**Supplementary Table 10**; **Supplementary Fig. 24**).

Following photon absorption, photoreceptor voltage responses are simulated using a stochastic sampling model coupled with an electrical membrane model for each photoreceptor^1,4,40^ (**Supplementary Fig. 27b**). Each microvillus generates a discrete conductance response, or quantum bump, upon photon absorption^101^, characterised by a latency period ("dead time"^118^) between photon capture and the onset of the quantal event^36,42,75^, and a conductance waveform (shape and amplitude) determined through noise analysis^42,75,119^.

After generating a response, the activated microvillus enters a refractory period during which it cannot respond to additional absorbed photons^34,36,40^. Individual quantal conductance events are summed within the membrane model to simulate the macroscopic photoreceptor voltage output^34,36,40,42,75^. This refractory quantal encoding allows photoreceptors to rapidly adapt to large changes in light intensity - providing a wide (logarithmic) operational range - and to actively enhance phasic changes in light input^6,27^.

**Modelling LMCs’ quantal stochastic neurotransmitter sampling**

The stochastic sampling process describes LMC function mathematically (**Supplementary Fig. 27b**). Unlike photoreceptors, which rely on microvilli, LMCs receive input through distinct histaminergic synaptic sites from R1-R6 photoreceptors arranged in neural superposition^4,9,49,61,74,112^. Each synaptic site independently generates a quantal conductance response upon synaptic activation, following binomial statistics, whereas photoreceptors follow Poisson statistics^120^. In LMCs, activation probability is determined by photoreceptor voltage, and a refractory period is calculated for each synaptic site after activation.

The binomial postsynaptic site activation probability, linking the photoreceptor membrane model^121^ to postsynaptic activation, is modelled as a differential equation. This equation takes photoreceptor membrane voltage as an input, with LMC membrane voltage providing feedback input, along with additional tonic activity^65^, modelled as a small probability of generating a miniature LMC response. The LMC electrical membrane model then sums the quantal synaptic conductances, followed by a simulation of the resulting membrane voltage.

**Application: Simulating High-Information-Rate Conditions**

To validate the model’s predictive capabilities, we simulate conditions generating the highest measured information rates in *Musca* photoreceptors and LMCs under bursty stimulation (contrast of 1.29 and a bandwidth of 200 Hz). Parameters derived from in vivo neuron recordings inform model simulations. Then, we run simulations to verify that the model accurately reproduces the temporal dynamics and information rates observed in the recordings. These simulations successfully reproduce observed temporal dynamics and information rates, using the modelled photoreceptor outputs as inputs for the LMC simulations.

**IV.3. Why a Multiscale Stochastic Sampling Model for LMC Visual Information Flow?**

Photoreceptors have been extensively characterised by combining stochastic sampling and electrical membrane models^1,4,6,34,40^. LMCs exhibit closely parallel functional motifs^8,13,16,21,48,122,123^, making the same multiscale stochastic framework an ideal choice. Key advantages include:

- **Stochastic independence:** Photoreceptor microvilli are structurally isolated - connected to the cell body only through narrow openings at their base - allowing them to function as independent stochastic photon-sampling units^6,34,40^. Similarly, LMC postsynaptic sites, located on elongated dendritic spines, likely operate independently as stochastic (histamine-molecule) sampling units^9,120^.
- **Adaptive response to bright light:** Photoreceptors exhibit an adapting differential response at the onset of a bright light pulse due to the refractory period limiting available samples^6,40^. Likewise, LMCs demonstrate a derivative response to bright stimuli, attributable to refractoriness at synaptic sites^8,13,16,54,124^.
- **Lack of refractoriness in dim light:** With ample microvilli, practically every absorbed photon triggers a bump; thus, photoreceptors do not exhibit refractory adaptation^6,40^. Correspondingly, LMCs show an absence of transient derivative responses in low-light conditions, lacking refractory adaptation^6,16,40,125^.
- **Wide dynamic range:** Photoreceptors operate across an extensive (logarithmic) illumination range, primarily due to adaptive mechanisms such as refractory photon sampling^1,6,34-36,40,101,126,127^. Similarly, LMCs effectively adapt their responses across a broad range of lighting conditions^8,13,16,54,124,128^.
- **Primary noise sources:** Photoreceptor response variability to repeated stimulation (noise) - though minute relative to the mean response (signal; *cf*. **Supplementary Fig. 4a**) - primarily arises from stochastic photon sampling^6,28,40^. Similarly, LMC variability largely stems from stochastic quantal histamine release^13,56,65,129^. For example, strong hyperpolarisation of photoreceptors - such as that induced by an ultra-bright stimulus activating the Na⁺/K⁺ exchanger in dark-adapted cells - halts presynaptic histamine release^65^. This effectively “silences the synapse,” leading to LMC depolarisation and a marked reduction in response variability^65^.
- **Refractory-based cycling:** Photoreceptor microvilli undergo cycles governed by refractory periods, analogous to refractory-based synaptic vesicle recycling in LMCs^121^.
- **Nonlinear summation:** LMC membranes nonlinearly integrate synaptic inputs^122^, similar to the photoreceptor membrane summation of microvillar responses^7,40,123,130^.
- **Emergent multiscale dynamics**: Conventional, narrowly focused modelling methods cannot capture the complex, emergent response properties and dynamics arising from interactions across scales - such as *high-frequency jumping*, *efficient encoding*, *hyperacute vision*, *fast adaptive gain control*, and *predictive time-locking* - that remain obscured when neural components are studied in isolation. A multiscale stochastic sampling model provides insights into these integrated mechanisms.

**IV.4. Noise Analysis of Photoreceptor and LMC Voltage Responses**

Noise analysis of voltage recordings (**Figure S24**) was performed for two distinct cases:

- Light stimulus (photon flux) to the photoreceptor membrane voltage^42,75,119,131,132^
- Photoreceptor membrane voltage to LMC membrane voltage, reflecting synaptic transmission^8,13,120,133^ .

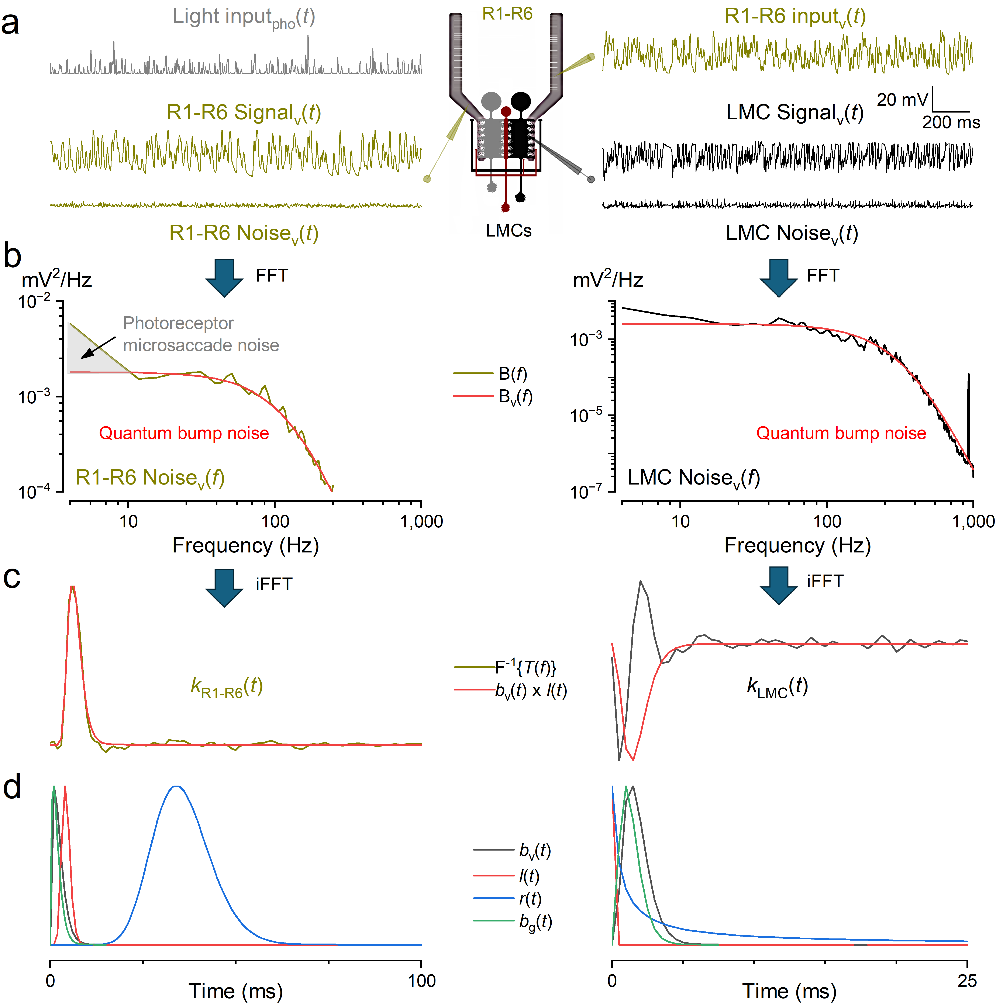
**Supplementary Fig. 28.** **Noise analysis of photoreceptor and LMC voltage responses.**

Left panels illustrate the analysis of voltage noise between light input and photoreceptor membrane voltage.

Right panels show the corresponding analysis between photoreceptor voltage input and LMC membrane voltage, reflecting synaptic transmission.

(**a**) Mean voltage responses (Signals) and example noise traces from R1-R6 photoreceptors (left) and LMCs (right) to a repeated bursty stimulus (Light input). Noise traces are obtained by subtracting the mean from each individual trial.

(**b**) Power spectra of the photoreceptor (left) and LMC (right) voltage noise signals, with fitted Lorentzian functions (red) capturing the quantal frequency characteristics (quantum bump noise). Photoreceptor voltage noise also contains low-frequency jitter caused by photomechanical photoreceptor microsaccades^1^ (gray area).

(**c**) Normalised impulse responses $k(t)$, calculated via inverse Fourier transform of the transfer function between input $x_{t}$ and mean output $\mu_{t}$, showing the temporal profile of photoreceptor (left) and LMC (right) transmission.

(**d**) Normalised voltage quantum bump waveform, $b_{v}(t)$, latency distribution, $l(t)$, refractory distribution $r(t)$ and conductance quantum bump waveform, $b_{g}\left( t \right),$used for fitting model parameters listed in **Supplementary Table 12**. For LMCs, the fit is aligned to the negative-going voltage bump.

For a repeated stimulus pattern over time, denoted as $x_{t}$, voltage traces, $y_{td}$ (where $t$ represents time and $d$ indicates repeats), were analysed after subtracting their DC offset (**Supplementary Fig. 27a**). For photoreceptors, $x_{t}$ represents the light stimulus; for LMCs, $x_{t}$ is the average photoreceptor voltage response.

We first calculated the mean response across repeats:

$\mu_{t}=E_{d}(y_{td})$	(1)

and the variance of individual traces by subtracting mean response from trace and then taking variance over time:

$\sigma_{d}^{2}=Var(y_{td}-\mu_{t})$	(2)

Traces with variances exceeding 1.75 times the lowest observed variance discarded, typically corresponding to initial recordings where neurons are adapting to new stimuli. Then repeat the mean and variance calculations with the rest of the traces. After removing these initial adaptation traces, we recalculated the mean and variance.

The final mean response for photoreceptors was calculated as:

$\mu=E_{t}\left( \mu_{t} \right)-E_{rest}$	(3)

where $E_{rest}$ is the resting potential. For LMCs, due to tonic histamine release and hyperpolarised resting voltage, the calculation differs:

$\mu=E_{t}\left( \mu_{t} \right)-max\left( \mu_{t} \right)$	(4)

This distinction reflects the fundamental difference in photoreceptor and LMC resting conditions and is consistent across analyses^13^.

We assumed the elementary voltage response (quantum bump) follows a gamma function^42,75,119^ (**Supplementary Fig. 28c**):

$b_{v}\left( t \right)\propto\frac{1}{\left( n_{v}!\tau_{v} \right)}\left( \frac{t}{\tau_{v}} \right)^{n_{v}}e^{-t/\tau_{v}}$	(5)

where $\tau$ is the time constant of the filter, and $n$ the order (or shape parameter) of the gamma function, and v stands for voltage. Consequently, the bump's power spectrum in the frequency domain follows a Lorentzian curve (**Supplementary Fig. 28b**):

$b_{v}\left( f \right)\propto{[1+\left( 2\pi\tau f \right)^{2}]}^{-\left( n_{v}+1 \right)}$	(6)

Here $f$ indicates the frequency. For LMC responses, the mean noise power spectrum ($B\left( f \right)$) is computed as:

$B^{LMC}\left( f \right)=E_{d}[F\left( y_{td}-\mu_{t} \right)]$	(7)

Where $F(\cdot)$ represents the Fourier transform.

Photoreceptor power spectra, however, can be refined by subtracting separately measured dark noise, $N^{D}(f)$, which also includes any residual instrumental noise:

$B^{PR}\left( f \right)=E_{d}[F\left( y_{td}-\mu_{t} \right)]-N^{D}(f)$	(8)

Note, the subtraction is not feasible for LMCs due to tonic synaptic activation.

Following Campbell’s theorem, $B(f)$ is assumed to consist of independent, additive quantal responses with similar shapes, represented by fitting a Lorentzian curve to the central frequency range of $B(f)$.

**Supplementary Table 13. Parameters for simulating voltage responses**

| **Model parameter** | **Photoreceptor value** | **LMC value** |
| --- | --- | --- |
| Mean response, $\mu$ | 13.8 mV | -11.8 mV |
| Number of quantal sampling units, $N$ | 54,000 | 2,340 |
| Voltage quantal response gamma decay, $\tau_{v}$ | 0.71 ms | 0.45 ms |
| Voltage quantal response gamma exponent, $n_{v}$ | 2.4 | 3.0048 |
| Voltage quantal response average duration, $T_{v}$ | 4.1 ms | 2.9 ms |
| Area of voltage quantal response, $s_{v}$ | 8.2*10^-5^ mVs | 1.15*10^-4^ mVs |
| Conductance quantal response gamma decay, $\tau_{g}$ | 0.58 ms | 0.44 ms |
| Conductance quantal response gamma exponent, $n_{g}$ | 1.24 | 2.7135 |
| Conductance quantal response average duration, $T_{g}$ | 2.5 ms | 2.7 ms |
| Area of voltage quantal response, $s_{g}$ | 2.50*10^-14^ Ss | 1.15*10^-13^ Ss |
| Quantal bump rate, (λ) | 186*10^3^ quanta/s | 142*10^3^ quanta/s |
| Latency gamma decay, $\tau_{l}$ | 0.37 ms | 0.023 ms |
| Latency gamma exponent, $n_{l}$ | 10.4 | 1 |
| Refractory gamma decay, $\tau_{r}$ | 3.5 ms | 585 ms |
| Refractory gamma exponent, $n_{r}$ | 10.5 | 0.0268 |
| Specific conductance, $C_{m}$ | 0.9 μF/cm^2^ | 0.9 μF/cm^2^ |
| Membrane capacitance, $C$ | 17.9 pF | 18 pF |
| Leak conductance, $g_{rest}$ | 24.9 nS | 45.7 nS |
| Input reverse potential, $E_{in}$ | 10 mV | -85 mV |
| Passive reverse potential, $E_{rest}$ | -76.8 mV | -29.3 mV |

Parameters $\tau_{v}$ and $n_{v}$ extracted from this fit define the gamma function of the miniature response (**Supplementary Table 13**), following the formalism of Wong et al.^119^. The gamma bump duration $T_{v}$ is calculated as:

$T_{v}=\frac{\tau_{v}\left( n_{v}! \right)^{2}2^{2n_{v}+1}}{\left( 2n_{v} \right)!}$	(9)

From noise traces, we compute the quantal response amplitude $h$ and the quantum bump rate $\lambda$:

$h= \frac{E_{d}\left( \sigma_{d}^{2} \right)}{\mu}$	(10)

$\lambda={\frac{1}{T}}_{v}\times\frac{\mu^{2}}{E_{d}\left( \sigma_{d}^{2} \right)}$	(11)

The quantal response’s total area $s$ is calculated as:

$s=\int b_{v}\left( t \right)dt=hT_{v}$	(12)

Note that LMC responses follow binomial rather than Poisson statistics, making $h$ and $\lambda$ approximations^120^.

The latency $l(t)$ of the miniature response is derived from the impulse response, calculated from the transfer function^42,75^ $T_{xy}\left( x_{t},\mu_{t} \right)$:

$T_{xy}\left( x_{t},\mu_{t} \right)=\frac{\left\langle X\left( f \right)\underline{Y(f)} \right\rangle}{\left\langle X\left( f \right)\underline{X(f)} \right\rangle}$	, $X\left( f \right)=F\left\{ x_{t} \right\}$ and $Y\left( f \right)=F\left\{ \mu_{t} \right\}$	(13)

, where the <> denotes average.

The impulse response, $k(t)$ (**Supplementary Fig. 28c**) is obtained by inverse Fourier transform:

$k\left( t \right)=F^{-1}\left\{ T_{xy}\left( x_{t},\mu_{t} \right) \right\}$	(14)

This impulse response represents the convolution of the miniature response gamma function and the latency gamma function$l(t)=\Gamma(nl, l)$:

$k\left( t \right)= b_{v}(t)\otimes l(t)$	(15)

We performed a least-squares fit of the convolved function to $k\left( t \right)$. For LMCs, where responses are inverted, fitting is performed to the prominent negative peak indicating zero latency (**Supplementary Fig. 28d**, **Supplementary Table 13**).

**IV.5. *Musca* Photoreceptor and LMC Membrane Properties**

Both R1-R6 photoreceptors and LMCs possess three principal classes of ion channels^21,122,123,126,134,135^ (**Supplementary Fig. 27a**): (**1**) ***input channels***, gated by light (photoreceptors) or neurotransmitter (LMCs); (**2**) ***delayed‐rectifier* potassium channels**; and (**3**) ***leak (passive) channels****.*

**1. Input Conductances**

- Photoreceptors express light‐gated TRP/TRPL channels^106^, whose stochastic, quantal openings generate the time‐varying input conductance $g_{in}(t)$. This light-induced conductance^28,40,130^ has a reversal potential

$E_{in}^{PR}=10 mV$.

- LMCs bear histamine-gated ligand channels^129^ with the reverse potential

$E_{in}^{LMC}= -85 mV$.

**2. Passive Membrane Parameters**

**Photoreceptors (R1-R6)**

- Resting potential:

$E_{rest}^{PR}=-76.8 mV$.

- Membrane capacitance $C^{Rk}$ and conductance $g_{rest}^{Rk}$, where $k=1-6$ is the photoreceptor index, are obtained by fitting the exponential charging curve to current‑injection pulses in current‑clamp recordings (**Supplementary Table 13**).
- Mebrane surface area is then

$S_{m}^{Rk}=\frac{C^{Rk}}{C_{m}}$,	(16)

where $C_{m}$ is the specific membrane capacitance.

**LMCs**

- Resting potential is chosen so that, in the absence of histaminergic input, the membrane sits at the $\max(\mu_{t})$ peak of its subthreshold response:

$E_{rest}^{LMC}=-29.3 mV$.

- Capacitance $C^{LMC}$ , conductance $g_{rest}^{LMC}$, and the surface area $S_{m}^{LMC}$ are fit to reproduce “bursty” voltage recordings (**Supplementary Table 12**). These values closely match those measured in blowfly, *Calliphora* LMCs^123^ (synaptic zone $\approx$ ~0.45 x 10^-5^ cm^2^ and $C_{m}^{LMC,calliphora}$= 1-3 μF/cm^2^).

**3. Delayed-Rectified Potassium Channels**

Both cell types express voltage‐activated $K^{+}$channels^122,130,135^,$K_{s}$ for photoreceptor and $K_{d}$ for LMC, described by Hodgkin-Huxley-type kinetics:

$\frac{dx}{dt}=\frac{x_{\infty}(V)-x}{\tau_{x}(V)},x\in\left\{ m,n,h_{p},h_{l} \right\}$,	(17)

Where $x_{\infty}(V)$ is the voltage-dependent steady-state and $\tau_{x}(V)$ the time constant of the gate $x$.

| **Cell type** | **Channel model** | $\boldsymbol{E}_{\boldsymbol{K}^{\boldsymbol{+}}}$ **(mV)** | $\boldsymbol{g}_{\boldsymbol{K}}$ **(mS/cm^2^)** | **Gates** |
| --- | --- | --- | --- | --- |
| R1-R6 photoreceptor | *Drosophila* *Shab* channel^28,40,130,134^ | -85 | 10 | 2 activation $(n$)  1 inactivation $(h_{p}$) |
| LMC | in Rusanen et al.^123^ (*Calliphora*) | -65 | 6 | 1 activation (m)  1 inactivation $(h_{l}$) |

**4. Membrane Voltage Equations**

The total membrane current balances capacitive charging.

- **Photoreceptor**:

$C_{m}\frac{dV^{PR}}{dt}=\frac{g_{in}^{PR}\left( t \right)}{S_{m}^{Rk}}\left( V^{PR}-E_{in}^{PR} \right)+\frac{g_{rest}^{Rk}}{S_{m}^{Rk}}\left( V^{PR}-E_{rest}^{PR} \right)+g_{Ks}n^{2} h_{p} (V^{PR}-E_{K}^{Pr})$	(18)

- **LMC**:

$C_{m}\frac{dV^{LMC}}{dt}=\frac{g_{in}\left( t \right)}{S_{m}^{LMC}}\left( V^{LMC}-E_{in}^{LMC} \right)+\frac{g_{rest}^{LMC}}{S_{m}^{LMC}}\left( V^{LMC}-E_{rest}^{LMC} \right)+g_{Kd}m h_{l} (V^{LMC}-E_{K}^{LMC})$	(19)

**IV.6. Estimating Quantum Bump Conductance Responses**

Quantum bump, or miniature conductance responses, $g_{g}^{y}(t)$, where y is either PR for photoreceptor or LMC for LMC (**Supplementary Fig. 28b**), are derived from the estimated quantum bump voltage responses via the membrane’s input-input conductance transfer function:

$H^{y}\left( f \right)= \frac{G_{g}^{y}(f)}{V^{y}(f)}$ ,	(20)

where $V^{y}(f)$ and $G_{g}^{y}(f)$ are the Fourier transforms of the voltage response and the miniature conductance (i.e., the average quantum bump), respectively.

**1. Decomposition of Total Conductance**

The instantaneous input conductance $g_{in}^{y}$ is decomposed into a time‑varying quantum bump component and a steady offset:

$g_{in}^{y}(t)=g_{g}^{y}(t)+g_{mean}^{y}$	(21)

Here, $g_{mean}$, is the constant conductance that, in isolation, holds the membrane at its mean voltage.

- **Photoreceptor:**

$\bar{V}^{PR}=E_{rest}^{PR}+\mu$	(22)

- **LMC:**

$\bar{V}^{LMC}=max(\mu_{t})+\mu$	(23)

In these expressions, $\mu$ (or $\mu_{t}$) denotes the average light‑induced depolarisation.

**2. Solving for the Average Conductance**

At steady state ($\dot{dV/dt}=0$) with no dynamic stimulus-driven quantum bump fluctuations ($g_{g}\left( t \right)=0$ ), the membrane equation reduces to a balance of leak and potassium currents. Solving for $g_{mean}$ gives:

- **Photoreceptor:**

$g_{mean}^{PR}=g_{rest}^{Rk}E_{rest}^{PR}+n_{\infty}\left( \bar{V}^{PR} \right)^{2}h_{p\infty}\left( \bar{V}^{PR} \right)g_{Ks}E_{K}^{Pr}-\frac{\bar{V}^{PR}(g_{rest}^{Rk}+n_{\infty}\left( \bar{V}^{PR} \right)^{2}h_{p\infty}(\bar{V}^{PR})g_{Ks})}{(\bar{V}^{PR}-E_{in}^{PR})}$	(24)

- **LMC:**

$g_{mean}^{LMC}=g_{rest}^{LMC}E_{rest}^{LMC}+m_{\infty}(\bar{V}^{LMC})h_{l\infty}\left( \bar{V}^{LMC} \right)g_{Kd}E_{K}^{LMC}-\frac{\bar{V}^{LMC}(g_{rest}^{LMC}+m_{\infty}(\bar{V}^{LMC})h_{l\infty}(\bar{V}^{LMC})g_{Kd})}{(\bar{V}^{LMC}-E_{in}^{LMC})}$	(25)

**3. Linearised Transfer Function**

Linearising around the mean voltage and taking the Laplace transform of the membrane equation yields the frequency‑domain transfer function:

- **Photoreceptor:**

(26)

$H^{PR}\left( f \right)=(C^{Rk}2\pi if+g_{rest}^{Rk}+g_{mean}^{PR}+{n_{\infty}(V)}^{2}h_{p\infty}(V)g_{Ks}-\left( g_{rest}^{Rk}E_{rest}^{PR}+g_{mean}^{PR}E_{in}^{PR}+g_{Ks}{n_{\infty}(\bar{V}^{PR})}^{2}h_{p\infty}(\bar{V}^{PR})E_{K}^{Pr} \right)/\bar{V}^{PR})/(E_{in}^{PR}-\bar{V}^{PR})$

- **LMC:**

(27)

$$H^{LMC}\left( f \right)=(C^{LMC}2\pi if+g_{rest}^{LMC}+g_{mean}^{LMC}+{m_{\infty}(\bar{V}^{LMC})h_{l\infty}(\bar{V}^{LMC})g}_{Kd}-\left( g_{rest}^{LMC}E_{rest}^{LMC}+g_{mean}^{LMC}E_{in}^{LMC}+g_{Kd}m_{\infty}(\bar{V}^{LMC})h_{l\infty}(\bar{V}^{LMC})E_{K}^{LMC} \right)/\bar{V}^{LMC})/(E_{in}^{LMC}-\bar{V}^{LMC})$$

**4. Conductance Noise Spectrum**

Given the measured voltage‑noise power spectrum $B^{y}(f)$ (of a voltage noise trace composed of uniformly sized quantum bumps, $b_{v}\left( t \right)$, Eq. 5), the corresponding conductance noise spectrum is:

$B_{g}^{y}\left( f \right)=B^{y}(f)\times\left| H^{y}\left( f \right) \right|^{2}$	(28)

A Lorentz curve is then fitted to the central portion of $B_{g}^{y}\left( f \right)$, yielding the bump shape parameters, $n_{g}^{y}$ and $\tau_{g}^{y}$ (**Supplementary Table 13**).

**5. Quantum Bump Duration and Area**

From the Lorentz fit:

- **Average conductance‑bump duration** (compare to Eq. 9)

$T_{g}^{y}=\tau_{g}^{y}\left( n_{g}^{y}! \right)^{2}2^{2n_{g}^{y}+1}/\left( 2n_{g}^{y} \right)!$	(29)

- **Conductance bump area** (derived from Eq. 12)

$s_{g}^{y}=h^{y}\times H^{y}(0)\times T_{g}^{y}$.	(30)

Here, $h$ is the quantal response amplitude of voltage and $H(0)$ is the zero‑frequency gain of the transfer function.

**6. Methods and Data Sources**

- **Stimulation**: All values were derived under “bursty” light‐contrast stimuli delivered at 200 Hz.
- **Photoreceptor passive properties**: Membrane capacitance and conductance were measured directly via current‑clamp recordings.
- **LMC membrane parameters**: Electrical parameters were obtained by fitting the model to the same bursty voltage responses.
- **Photoreceptor quantum bumps**: Voltage bump amplitudes and kinetics came from photoreceptor high‑input‑resistance recordings, ensuring minimal attenuation.
- **Photoreceptor refractoriness**: Refractory time constants were extracted by fitting the response to prolonged light pulses.
- **LMC quantum bumps and refractoriness**: Quantal bump properties and refractoriness were fitted for the “bursty” stimulus.

**IV.7. Stochastic Sampling Models**

Building on the noise analysis of photoreceptor and LMC temporal dynamics (**Supplementary Table 12**), we implement a stochastic sampling framework^1,40^ (**Supplementary Fig. 27**) in which each quantal unit - microvillus in a photoreceptor or synaptic site in an LMC - generates an identical miniature conductance waveform $g_{g}(t)$ after a random latency drawn from a gamma distribution $\Gamma\left( n_{l},\tau_{l} \right)$.

**1. Quantal Sampling Unit Counts**

- **Photoreceptor (microvilli)**	$N_{PR}\approx54,000$

Estimated from a R1-R6 rhabdomere of diameter 1.4 µm and average length 132 µm (**Supplementary Tables 10** and **11**), with microvillar diameter $\approx$ 0.065 µm^89^, where microvilli are hexagonally packed.

- **LMC (dendritic spines)**	$N_{LMC}\approx390 x 6\approx2,340$

Fitted to reproduce bursty voltage responses under the 200 Hz contrast stimulus.

**2. Simulation Loop**

We advance in discrete steps $t_{i} \in t=k\Delta t$ ($k=0,1,2,\ldots$. ), where $\Delta t$ is the time step, tracking three pools:

1. Active units: $A(t)$
2. Latency-delayed activations:$L(t)$
3. Refractory returns: $I(t)$

Initialisation :	$A\left( 0 \right)=N, L\left( t \right)=0, I\left( t \right)=0, g\left( t \right)=0$

At each step:

1. Draw activations $R\left( t \right)$ (Eqs 31-34)
2. Update active pool $A\left( t \right), L\left( t \right) and I(t)$ (Eqs 35-38)
3. Sum conductance $g\left( t \right)$(Eqs 39-41)

**2.1 Quantal activation**

- **Photoreceptor**

1. Convert light intensity, $LI(t_{i})$ (see **Section IV.9.ii**) to absorbed photons

$PA\left( t_{i} \right) \sim Poisson(LI\left( t_{i} \right)*\Delta t)$	(31)

2. Distribute $P_{a}$ across $N$ microvilli:

$$\left\{ M_{n} \right\} \sim multinomial(n=PA(t_{i}),p_{m}=1/N)$$

3. Binary activation in each microvillus $m$

$R_{m}(t_{i},m)=\left\{ \begin{aligned} 1, M_{m}>0, \\ 0, otherwise \end{aligned} \right.$	(32)

4. Sum over active microvilli:

$R\left( t_{i} \right)=\sum_{m=1}^{A\left( t_{i} \right)} R_{m}\left( t_{i},m \right)$	(33)

- **LMC**

$R\left( t_{i} \right) \sim Binomial\left( n= A\left( t_{i} \right),p=P\left( t_{i} \right) \right)$	(34)

where $P\left( t \right)$ (see **Section IV.11**) is the instantaneous release probability from the synaptic model.

**2.2 Update pools**

- Enqueue latency:

$L\left( t \right) +=\sum_{j=1}^{R(t_{i})} \delta(t-t_{i}-\Gamma(n_{l}, \tau_{l}))$	(35)

, where $\delta(t)$ is the Dirac delta function.

- Compute refractory returns $I\left( t \right)$ after latency and bump duration $\tau_{g}$ (photoreceptor) or immediately (LMC):
- **Photoreceptor**

$I\left( t \right) +=\sum_{k=1}^{L\left( t_{i} \right)} \delta\left( t-t_{i}-\Gamma\left( n_{r}, \tau_{r} \right)-\tau_{g} \right)$	(36)

- **LMC**

$I\left( t \right) +=\sum_{k=1}^{R\left( t_{i} \right)} \delta\left( t-t_{i}-\Gamma\left( n_{r}, \tau_{r} \right) \right)$	(37)

- Update active pool for next round:

$A\left( t_{i}+\Delta t \right)=A\left( t_{i} \right)-R\left( t_{i} \right)+I(t_{i})$	(38)

**2.3. Conductance Summation**

- **Photoreceptor macroscopic conductance**

$g_{in}^{PR}\left( t \right) +=L\left( t_{i} \right) g_{g}\left( t-t_{i} \right)$	(39)

- **LMC total conductance**

1. First assign each latency event (count: $L\left( t_{i} \right))$to a random synapse rs $\in\left\{ 1,\ldots,N \right\}$, ensuring that one bump is only assigned by permuting the synapse order and taking the $L\left( t_{i} \right)$first synapses in the permuted series:

$g_{s}\left( t,rs \right) +=\sum_{k=permute(N)}^{L\left( t_{i} \right)} g_{g}\left( t-t_{i} \right) \delta\left( rs-k \right)$	(40)

2. Then cap each synaptic site to maximum $g_{g}$ and sum:

$g_{in}^{LMC}\left( t_{i} \right) =\sum_{n=1}^{N} \min[g_{s}\left( t_{i},n \right),max(g_{g}\left( t \right))].$	(41)

**Key Points**

- **Latency** $\Gamma\left( n_{l}, \tau_{l} \right)$ and **refractoriness** $\Gamma\left( n_{r}, \tau_{r} \right)$ are fitted separately for photoreceptors (hand-fit to 1 second pulses) and LMCs (fit to 200 Hz bursts)
- **Photoreceptor** $\tau_{g}$ is measured from the voltage-bump duration; **LMC** $\tau_{g}$ is set to zero, as multiple vesicles can overlap.

**IV.8. The Optical Models for Photoreceptor Receptive Fields Depend on Rhabdomere Position**

We simulate light capture within a single ommatidium by combining geometric ray-tracing through the corneal lens with the *Fourier‑Transform Beam Propagation Method* (FTBPM)^4^ to model light propagation through the crystal cone and rhabdomere. These simulations are based on the static *Musca* compound eye morphology, with parameters obtained from EM and X-ray imaging data (**Section II**). Optically, each ommatidium comprises three main elements:

1. **Corneal lens** (front surface)
2. **Crystal cone** (clear zone)
3. **Rhabdomere** (photosensitive waveguide)

**Methodology**

1. **Lens Raytracing (Geometric Optics).**

- Emit a parallel bundle of rays from a distant point source, uniformly sampling the lens aperture.
- Trace each ray through the corneal lens surface (lens properties in **Supplementary Table 10** and **11**) to compute the transmitted wavefront at the lens exit.

2. **FTBPM Propagation.**

- Initialise the electric field at the lens exit plane.
- Propagate this field through the crystal cone, aperture, and rhabdomeric waveguide using the split‑step Fourier method^4^ (optical properties in **Supplementary Table 10** and **11**).
- Compute local absorption by integrating field intensity losses along the rhabdomere’s length.

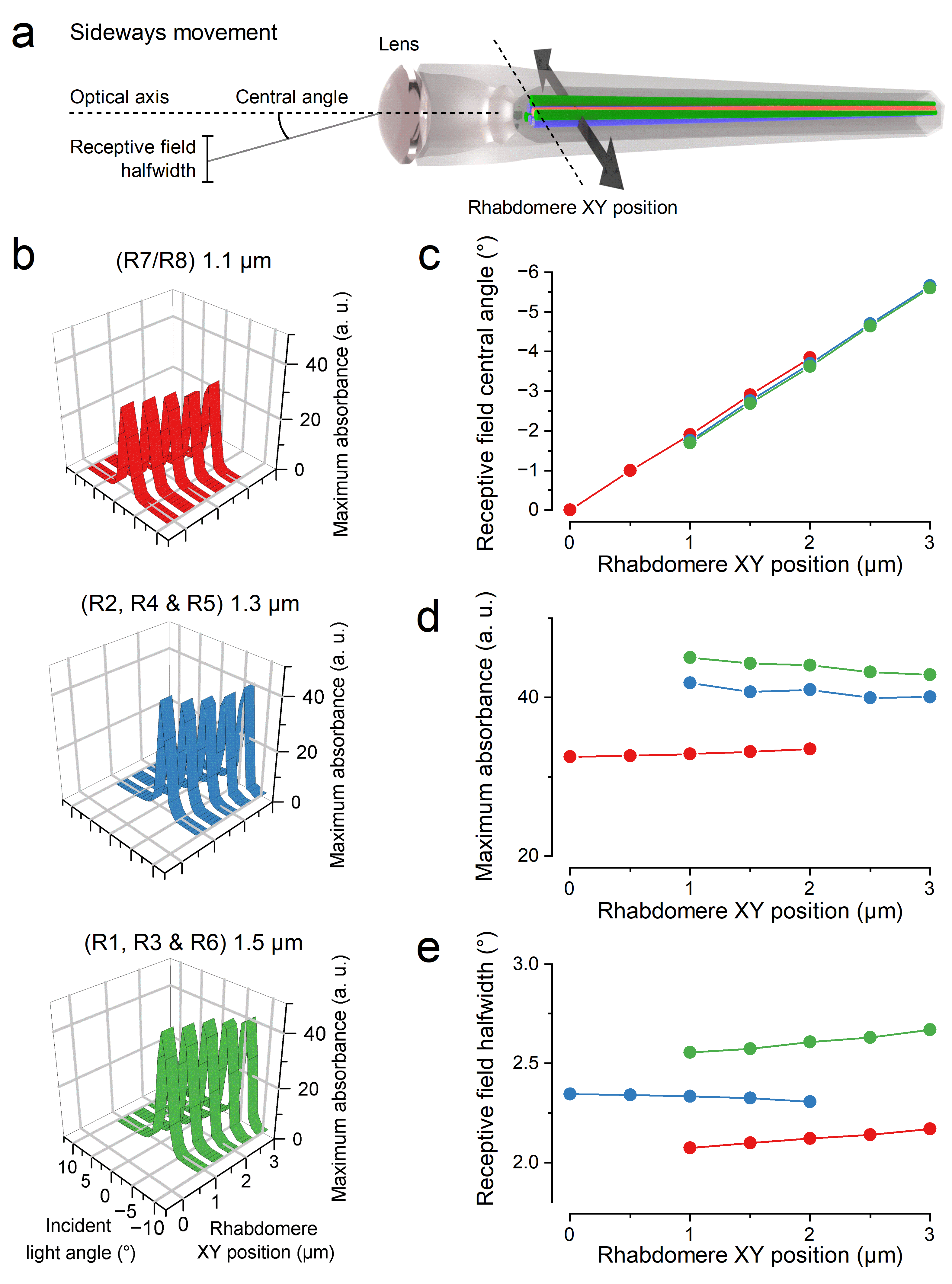
**Supplementary Fig. 29. Static optical simulations reveal how lateral shifts in rhabdomere position modulate spatial light absorption and receptive field tuning in *Musca* ommatidia**

(**a**) Diagram of an ommatidium illustrating how lateral (XY) displacements of rhabdomeres from the lens's optical axis alter the angle and width of their receptive fields.

(**b**) Simulated angular photon absorption profiles for rhabdomeres of different diameters: 1.1 µm (R7/R8, red), 1.3 µm (R2, R4, R5, blue), and 1.5 µm (R1, R3, R6, green). Each panel shows how the profiles change with increasing lateral rhabdomere displacement.

(**c**) Receptive field central angle (°) shifts linearly with rhabdomere XY position.

(**d**) Maximum photon absorption amplitude declines slightly with increasing lateral displacement, especially for wider rhabdomeres.

(**e**) Receptive field half-width (°) broadens with displacement, particularly in wider rhabdomeres, indicating a resolution trade-off.

These simulations show that even small submicron rhabdomere displacements within an ommatidium can dynamically adjust receptive field properties, offering a potential mechanism for adaptive visual tuning.

**Lateral Displacement Analysis**

- To reproduce the effect of the asymmetric R1-R6 rhabdomere positions underneath the ommatidium lens, we laterally offset the rhabdomere axis from the lens centre by up to ±3 µm (**Supplementary Fig. 29**).
- Simulations run for three rhabdomere diameters (rounded from **Section II**: **Supplementary Table 11**): 1.1 µm (R1), 1.3 µm (R2, R4 & R5) and 1.5 µm (R1, R3 & R6).
- Measured output:
- Receptive‑field centre shift of -1.86° per µm (negative indicates inversion by the lens) (**Supplementary Fig. 29b**).
- Peak amplitude of the angular sensitivity curve declines with offset (**Supplementary Fig. 29c**).
- Half‑width broadens as offset increases (**Supplementary Fig. 29d**).

**Axial Displacement Analysis**

- To reproduce the effect of R1-R6 rhabdomere tips resting different distances away from the ommatidium lens at different eye locations, we varied the rhabdomere’s axial distance from the lens exit between 19 µm and 32 µm (resting = 29.5 µm (**Supplementary Table 11**)), for the same set of diameters as lateral displacement.
- Results:
- Half‑width of the angular response narrows as the rhabdomere moves farther away (**Supplementary Fig. 30b**).
- Peak amplitude increases with increasing distance (**Supplementary Fig. 30c**).
- Numerical values for R1-R7/R8 receptive fields are provided in **Supplementary Table 15**.

**Validation against Experimental Data**

- Burton and Laughlin^67^ report acceptance angles of 2.94° (males) and 5.77° (females), though these appear to reflect standard deviations rather than true half‑widths.
- Vowles^87^ measured light‑adapted half‑widths of 3.0° (horizontal) and 2.5° (vertical), expanding to 8.5° and 4.5°, respectively, under dark adaptation.
- Our simulated half‑widths fall within these ranges, confirming that our optical model accurately predicts *Musca*’s angular sampling.

**Comparison to *Drosophila***

- *Musca* shares a similar lens diameter but features a larger overall eye, flatter curvature, slightly smaller rhabdomeres, longer lens‑to‑rhabdomere distance, and consequently narrower angular sensitivities than *Drosophila*^4,92^.

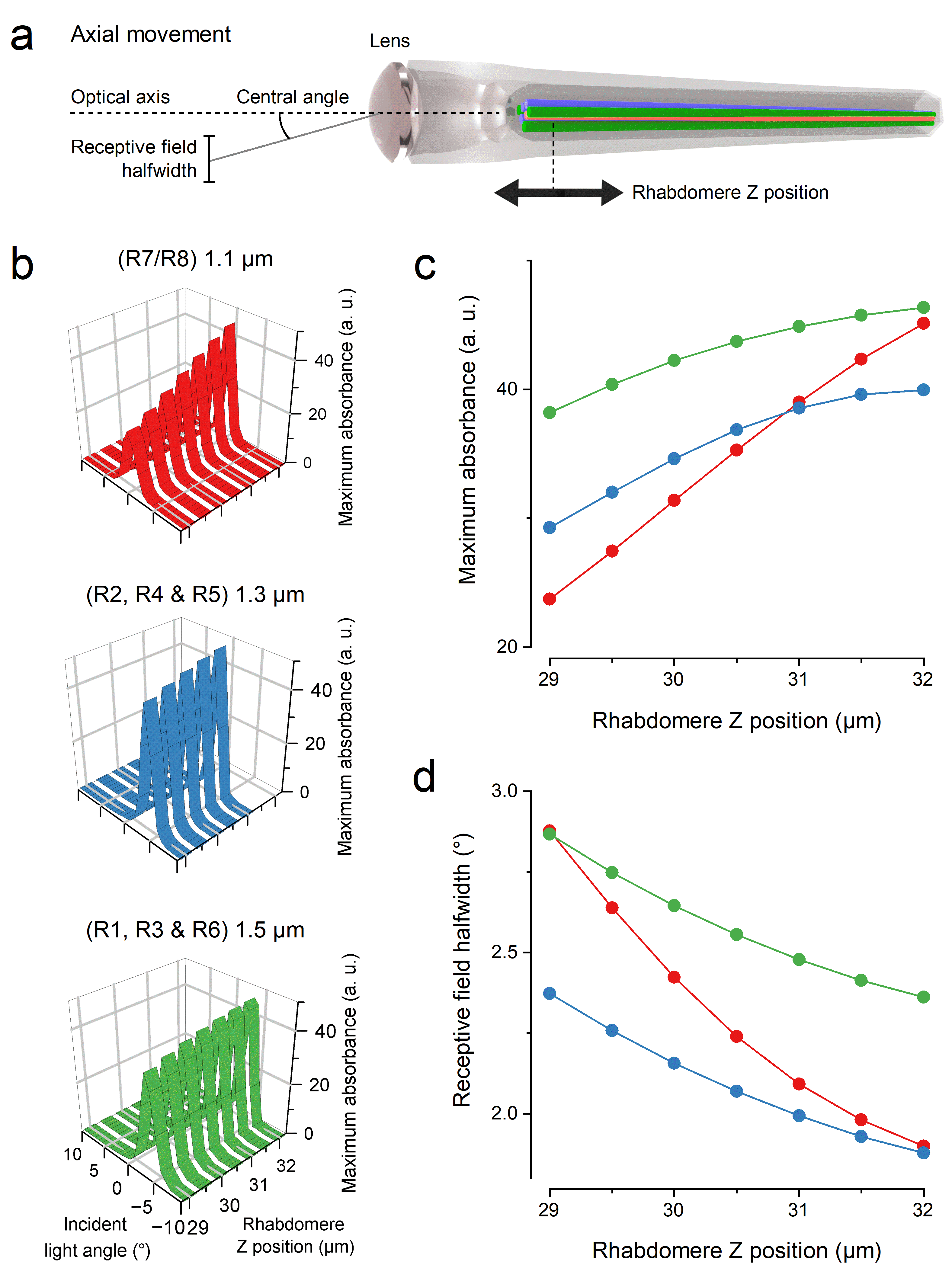
**Supplementary Fig. 30.** **Static optical simulations show how axial rhabdomere displacements modulate light absorption and receptive field sharpening in *Musca* ommatidia.**

(**a**) Schematic of an ommatidium illustrating how axial (Z-axis) shifts in rhabdomere position relative to the lens affect the incident angle and spatial tuning of light absorption.

(**b**) Simulated angular absorption profiles for rhabdomeres of varying diameters - 1.1 µm (R7/R8, red), 1.3 µm (R2, R4, R5, blue), and 1.5 µm (R1, R3, R6, green) - as their axial position is varied.

(**c**) Maximum photon absorption increases as rhabdomeres are shifted axially closer to the lens focal plane, with stronger gains observed for wider rhabdomeres.

(**d**) Receptive field half-width (°) narrows significantly with axial displacement, especially in narrower rhabdomeres, indicating improved spatial selectivity.

These simulations highlight how submicron axial repositioning of rhabdomeres can dynamically sharpen visual sampling and modulate signal gain within individual ommatidia.

**Supplementary Table 14.** Optical parameters of *Musca domestica* ommatidium.

| **Ommatidium parameter** | **Value** | **Reference** |
| --- | --- | --- |
| Cone/pigment-cell aperture diameter | 5 µm* | Chi and Carlson^111^ |
| Cone/pigment-cell aperture thickness | 2 μm* | Chi and Carlson^111^ |
| Cone/pigment-cell aperture total transmittance | 2.8%* | Kemppainen et al.^4^ |
| Ommatidial lens refractive index | 1.45* | Stavenga^92^ |
| Crystal cone refractive index and outside rhabdomere | 1.34* | Stavenga^92^ |
| Rhabdomere refractive index | 1.363* | Stavenga^92^ |
| Rhabdomere absorbance | 0.005/μm* | Kirschfeld^136^, Warrant and Nilsson^137^ |

*Same as *Drosophila melanogaster* simulations^4^.

**Supplementary Table 15.** Calculated receptive fields of R1-R7/8 photoreceptors and their changes as a function of their rhabdomere’s ommatidial position based on optical simulations where the light beam backpropagates through the rhabdomere and the ommatidial optics to the visual space.

| **Rhabdomere** | **Receptive field half width (**°) | **Receptive field half width change (**°/µm) | **Receptive field maximal amplitude** | **Receptive field maximal amplitude change** (1/µm) |
| --- | --- | --- | --- | --- |
| R1* | 2.62 | -0.17 | 41.7 | 2.7 |
| R2* | 2.10 | -0.16 | 38.9 | 3.7 |
| R3* | 2.62 | -0.17 | 41.7 | 2.7 |
| R4* | 2.10 | -0.16 | 38.9 | 3.7 |
| R5* | 2.13 | -0.16 | 35.5 | 3.7 |
| R6* | 2.57 | -0.17 | 43.2 | 2.7 |
| R7/R8 | 2.34 | -0.33 | 33.0 | 7.3 |

The distance from the centre of the ommatidium was considered in the parameter values.

*5 μm aperture, formed by the surrounding pigment cells, touching the rhabdomere tip’s outside edge.

**IV.9. Photomechanic Microsaccades Narrow and Move Photoreceptor Receptive Fields**

**IV.9.i. Photoreceptor Microsaccade Force Model**

We model each rhabdomere’s lateral (microsaccadic) movement inside an ommatidium (**Supplementary Fig. 27b**) as a spring-damper system driven by light-activated force^4^. The net acceleration of the rhabdomere is given by:

$\frac{d^{2}x}{{dt}^{2}}=F_{light}(t)-F_{d}(\dot{x})-k_{tot}(t)*x$	(42)

where:

1. **Light-activated force**

$F_{light}(t)=F_{light}^{max}\frac{{[act\left( t \right)]}^{n_{act}}}{F_{50}^{n_{act}}+{[act\left( t \right)]}^{n_{act}}}$	(43)

- $act\left( t \right)$is the delayed quantum-absorption signal, $L\left( t \right)$
- Single photoreceptor: $act\left( t \right)=L\left( t \right)$
- Whole ommatidium ($R1\ldots R7/R8$): $act\left( t \right)= \frac{1}{7}\sum_{R1}^{\frac{R7}{R8}} L\left( t \right)$
- Parameters (*Musca*-specific, fitted to repeated 10 ms pulses):
- $F_{light}=9.81\times{10}^{-4}\frac{\mu m}{{ms}^{2}}$
- Hill exponent $n_{act}=1$
- Half-max constant $F_{50}=25.5$microvilli activation per 0.5 ms

2. **Damping force**

$F_{d}(\dot{x})=-D_{coeff}(D_{base}^{-D_{exp}\dot{x}}-1)$	(44)

- Fitted values:
- $D_{coeff}=2.4\times{10}^{-4}\frac{\mu m}{{ms}^{2}}$
- $D_{base}=2$
- $D_{exp}=2100\frac{ms}{\mu m}$

3. **Spring stiffness**

$k_{tot}(t)=k_{0}+k_{coeff}\frac{[{act(t)]}^{n_{act}}}{F_{50}^{n_{act}}+{[act\left( t \right)]}^{n_{act}}}$	(45)

- Fitted values:
- $k_{0}=1\times{10}^{-4} {ms}^{-2}$
- $k_{coeff}=1\times{10}^{-3} {ms}^{-2}$

**Simulation details**

- We excluded the corneal lens from the mechanical model (recordings used a water droplet immersion).
- To mimic in vivo bursts, we applied 5.38 kph in 10 ms light pulses with 1 second intervals, using our single-photoreceptor stochastic sampling model (**Supplementary Fig. 31a**).
- All parameters were optimised by fitting simulated trajectories to empirically measured microsaccade bursts (**Supplementary Fig. 26e)**.

**IV.9.ii. Ray-Tracing Spatiotemporal Stimuli to Photoreceptors**

We compute each photoreceptor’s input light intensity $L_{i}(t)$ by tracing rays through its receptive field footprint (**Supplementary Fig. 31a**). The procedure^4^ is:

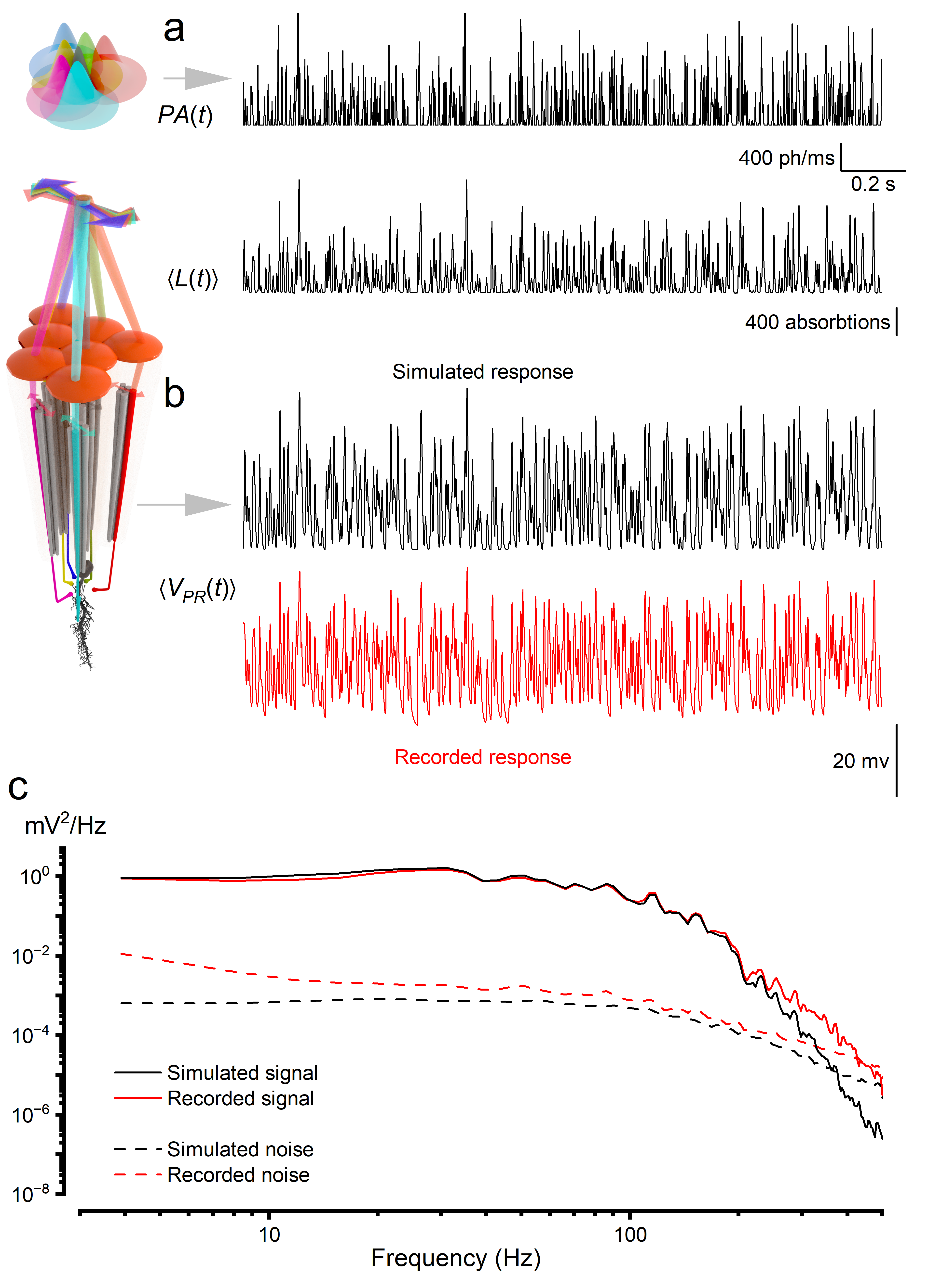
**Supplementary Fig. 31. Simulated R3 photoreceptor responses closely match in vivo recordings for bursty saccadic stimuli.**

(**a**) A R3 photoreceptor sample the light stimulus (photon rate changes) through a photomechanically moving and narrowing receptive field. It makes one morphodynamic “petal” of the neural superposition sampling matrix: R1-R6 photorceptors from neighbouring ommatidia forming the “flower pattern” of partially overlapping receptive fields. A high-contrast ($c$ = 1.29), 200 Hz bandwidth saccadic “bursty” light stimulus was simulated using the stochastic sampling model. The time-varying absorbed photon input $PA(t)$ drove quantal responses $\left\langle L(t) \right\rangle$ across 15 repetitions, following the removal of the first non-stationary trial. Each absorbed photon triggered a latency-distributed bump event, governed by parameters derived from Musca microvillar kinetics.

(**b**) Quantal conductance events were integrated by a Hodgkin-Huxley-style membrane model to produce the macroscopic voltage response $\left\langle V(t) \right\rangle$. The simulated voltage trace (black) shows strong temporal alignment with the in vivo recorded R1-R6 photoreceptor response (red) to the same bursty stimulus.

(**c**) Power spectral densities of the simulated (black) and recorded (red) voltage responses show close correspondence in both signal (solid lines) and noise (dashed lines) components, validating the model’s temporal and frequency-domain accuracy. The simulation included dynamic receptive field shifts due to microsaccades. These did not alter the signal spectrum but slightly elevated low-frequency noise, consistent with experimental data from both *Musca* and *Drosophila* photoreceptors.

1. **Generate rays**

- For each receptor (R1-R7/R8), emit a uniform 41×41 grid of rays in 20°× 20° box with varying angle from the lens centre.
- Assign ray angles by sampling a Gaussian distribution whose mean and width come from the receptor’s half-width and angular displacement (**Supplementary Tables 14** and **15**).

2. **Apply ommatidial sift**

- For receptors in neighbouring ommatidia, shift the ray‐grid origin by the interommatidial shift so that each sees the appropriate portion of the scene. The interommatidial sift (3.1°) is calculated based on R1-R6 average rhabdomere position sift (**Supplementary Tables 11**) multiplied by receptive‑field centre shift (**Section IV.9.i)**, while correcting for the hexagonal pattern. The sift is close to the interommatidial angle, but the specific position of the ommatidium dictates a slight shift. **Figure 1a** shows the receptive fields (marked as circles) of multiple ommatidia after ray tracing.

3. **Project onto the stimulus canvas**

- Intersect each ray with the pixelated stimulus screen (image or video).
- Record which pixel each ray hits and compute any angular offset from lateral displacement $x$ by photoreceptor microsaccades using the model in **Section IV.9.i** (**Supplementary Table 15**).

4. **Compute pixel weights**

- For each pixel, interpolate the fraction of rays (i.e. solid‐angle fraction) that fall within its bounds, adjusting for angular differences.

5. **Integrate intensity**

- Multiply each pixel’s luminance by its weight and sum over all pixels.
- Normalise by the total number of rays and scale by the maximum light level to yield $LI(t)$.

**IV.10. Simulating Photoreceptor Responses to Saccadic Bursty Contrast Stimuli**

To validate our *Musca* R1-R6 photoreceptor model (**Supplementary Fig. 31b**), we recreated the high‐contrast, saccadic bursty stimulus ($c$ = 1.29, 200 Hz bandwidth) known to drive maximal information rates in vivo (**Supplementary Fig. 4c**).

We first computed the time‐varying light input $L_{i}(t)$ by ray tracing a 3° patch (an 8 mm target at 15 cm, rendered on a 9×9 pixel grid) through each photoreceptor’s receptive field (**Section** **IV.3**). These fluctuations in $L_{i}(t)$ drove our two‐stage model:

1. **Stochastic sampling**: Each absorbed photon generates a quantal bump via the single‐microvillus sampler.
2. **Membrane integration**: Conductance bumps feed into a Hodgkin-Huxley-style electrical membrane, yielding the receptor voltage$V(t)$.

We also incorporated our microsaccade submodel (**Section** **IV.9**) so that the receptive‐field centre and width dynamically shift in response to each flash. All model parameters (latencies, refractoriness, membrane resistance) were drawn from *Musca* measurements (**Supplementary Table 13**, **Supplementary Figs. 22-23**), reflecting their rapid kinetics and low impedance.

**Simulation protocol:**

- Time step $\Delta t=1 ms$
- 16 repeats of the same burst sequence
- First repeat discarded to remove nonstationary adaptation effects

**Key findings:**

- After discarding the first trial, the mean bump rate $\lambda$ matched in vivo recordings (**Supplementary Fig. 31a**).
- The simulated voltage waveform $V(t)$ overlaid tightly with recorded photoreceptor traces (**Supplementary Fig. 31a**).
- Signal and noise power spectra of $V(t)$ matched experimental spectra across frequencies (**Supplementary Fig. 31b**).
- The simulated information rate was ≈ 2, 256 bits/s, closely matching the in vivo value of 2,422 bits/s.

Although large microsaccades (~0.8 µm; **Supplementary Fig. 26e**) can move the receptive field partly outside the 3° patch, their variable small‐scale nature leaves the signal spectrum essentially unchanged; they do, however, elevate low‐frequency noise, just as observed in both *Musca* and *Drosophila* photoreceptors^1^.

**IV.11. Synaptic Connection Model: Postsynaptic Activation Probability**

The synaptic model (**Supplementary Fig. 32b**) computes the LMC’s postsynaptic activation probability, $P(t)$, as a binomial release process whose instantaneous success rate is biophysically driven by presynaptic photoreceptor voltage, $V^{PR}(t)$, and feedback from the LMC membrane potential, $V^{LMC}(t)$^7,8,16,54,116^.

We model the LMC’s postsynaptic activation probability, $P(t)$, as the sum of a tonic component, $p_{ss}$, and a dynamic component, $p_{d}$ (**Supplementary Fig. 32b**):

$P\left( t \right)=p_{ss}+(1-p_{ss})\times p_{d}(t)$	(46)

1. **Tonic Activation** ($p_{ss}$)

- Baseline probability that a postsynaptic site is active in darkness, set to $p_{ss}=0.0529$, when $\Delta t=0.5 ms$ so the model reproduces the LMC’s dark‐adapted resting potential (**Supplementary Table 12**).
- This spontaneous, tonic release drives the characteristic voltage noise observed in LMCs under dark conditions^13,138^.

2. **Dynamic Activation (**$\boldsymbol{p}_{\boldsymbol{d}}\mathbf{(}\boldsymbol{t}\mathbf{)}$**)**

Captures rapid, voltage‐dependent increases in postsynaptic activation probability midpoint ($V_{0}=-55.9483 mV$, slope $V_{d}=2.5 mV$) via a simplified biophysical cascade:

1. **Photoreceptor depolarisation** $V_{PR}(t)$ opens voltage‐gated Ca²⁺ channels
2. **Ca²⁺ influx** triggers histamine vesicle release.
3. **Histamine binding** converts to postsynaptic activation in the LMC.

We express this as:

$\frac{{dp}_{d}}{dt}={\frac{1}{\tau_{p}}e}^{\frac{V^{PR}(t)-a_{f}p_{f}(t)-V_{0}}{V_{d}}}\left( 1-p_{d} \right) - \frac{1}{\tau_{n}}\frac{1}{1+(0.05/p_{d})^{2}}$	(47)

Voltage-driven activation	Hill-type decay (constant rate)

- **Activation time constant**: $\tau_{p}=0.1 ms$
- **Decay time constant**: $\tau_{n}=7.5 ms$ (models a near‐constant decay rate via Na⁺/Ca²⁺ exchange)
- **Voltage feedback**: $a_{f}=7 mV$ scales how much the LMC’s own feedback term $p_{f}\left( t \right)$ shapes (accentuates) the effective photoreceptor voltage.

3. **LMC Voltage Feedback** $p_{f}\left( t \right)$

Reflects slower modulation of release probability by the LMC’s membrane potential:

$\frac{dp_{f}}{dt}={\frac{1}{\tau_{f}}(e}^{\frac{V^{LMC}(t)-V_{f0}}{V_{fd}}}\left( 1-p_{f} \right)-p_{f})$	(48)

- **Midpoint:** $V_{f0}=-56 mV$
- **Slope:** $V_{fd}=2.5 mV$
- **Time constant:** $\tau_{f}=0.8 ms$

**Summary:**

- $p_{ss}$ sets the dark‐noise floor of postsynaptic activation.
- $p_{d}(t)$ drives rapid, contrast‐dependent increases via a Hill‐type activation and decay.
- $p_{f}(t)$ provides negative feedback from the LMC’s own voltage onto activation probability.

This model reproduces both the steady‐state noise and the fast, adaptive synaptic transmission observed experimentally (**Figs. 3-5**, **Supplementary Fig. 32b**).

4. **Observations, Clarifications and Explanations**

*Tonic Activation*: The parameter $p_{ss}$ represents the baseline probability that an LMC postsynaptic site is active in darkness^65,138^. We set $p_{ss}$ so that, in the absence of light, the model’s mean LMC membrane voltage matches its dark-adapted resting potential (**Supplementary Table 13**). This constant, stochastic release of histamine causes the characteristic voltage noise observed in LMCs under dark conditions. Importantly, because this tonic (random) activation hyperpolarises the cell to its resting level, any negative contrast step produces a transient depolarisation above rest.

*Dynamic Activation*: The time-varying component, $p_{d}(t)$, captures rapid, voltage-driven changes in release probability. In our model, photoreceptor depolarisation opens voltage-gated Ca²⁺ channels^121,139^, raising intracellular calcium and triggering histamine vesicle fusion at the synapse^115^. This process is modelled so that small voltage changes in the photoreceptor translate into corresponding increases in $p_{d}$, while a separate decay mechanism returns $p_{d}$ toward zero at a constant rate - mimicking the action of the Na⁺/Ca²⁺ exchanger, which clears calcium at its maximum rate^140^.

*Synaptic Kinetics*: Rather than using a simple exponential rise and decay, we implement a biophysically inspired activation function, combined with a rapid activation time constant. The decay back to baseline follows a Hill-type relationship that ensures a constant relaxation rate across most of the activation range, matching in vivo observations that LMC voltage returns to rest without overshoot or action-potential-like behaviour (**Supplementary Fig. 32c**).

*Feedback from LMC to Photoreceptor*: Finally, we include feedback^7,8,16,54,66,116^ term, $0\leq p_{f}(t)\leq1$, driven by the LMC’s membrane potential. When the LMC hyperpolarises, this feedback ($a_{f}$) reduces the effective photoreceptor voltage by up to 7 mV, raising the threshold for further dynamic activation and preventing runaway saturation. The feedback mechanism itself follows a voltage-dependent activation, capturing the moderate latency and persistence of this modulatory effect.

**IV.11.i. Neural Superposition of Neighbouring Ommatidia**

1. **Six-to-One Convergence**

- Each LMC pools input from six distinctive *morphodynamic* (photomechanically moving) photoreceptors: R1, R2, R3, R4, R5 and R6 - one from each of six adjacent ommatidia - that share broadly the same visual axis (**Supplementary Fig. 32a**)^4,52,61,141^.
- Each photoreceptor has an independent synaptic activation probability, $P(t)$.
- Each photoreceptor ($p$) contributes via one-sixth of the LMC’s synaptic sites ($N$), which sum is $g_{in}^{P}$ calculated according to Equation 41; the total conductance:

$g_{in}(t)=\sum_{p=1}^{6} g_{in}^{P}(t)$	(49)

2. **Receptive-Field & Absorption Scaling**

- Variations in R1-R6 photoreceptor morphology (**Supplementary Table 11**) affect their photon sampling properties.
- A photoreceptor’s spatially absorbed photon count, $L_{i}(t_{i})$, is determined by its receptive field.
- Receptive field shape is set by each rhabdomere diameter (i.e. its microvillus array) (**Supplementary Table 10**), rhabdomere position (**Supplementary Tables 11**), and lens geometry (**Supplementary Table 10**).
- Because R1-R6 rhabdomeres differ in size and optical‐axis offset - and the eye’s curvature is not a perfect sphere - their neural‐superposition receptive fields do not align precisely. Instead, they move photomechanically to slightly different directions while forming an overcomplete rosette tiling of a small visual patch^3,4,61^.
- Photon count $L_{i}\left( t_{i} \right)$follows directly from that field via ray tracing (**Section IV.3**).

3. **Rhabdomere Diameter Effects**

- Microvilli number, $N$: scales linearly with rhabdomere diameter (constant microvillus ∅ = 0.065 µm)³⁹.
- Membrane area: scales with diameter² (area ∝ diameter²), as an increase in diameter results in more microvilli and longer microvilli.
- The microvillar folding surface is significantly larger than the soma cylinder, meaning that 25% of the total photoreceptor membrane does not scale with rhabdomere diameter.
- Since specific membrane capacitance remains constant (**Supplementary Table 13**), capacitance is linearly related to the membrane area: $C$: ∝ total membrane area.

4. **Conductance Scaling**

- Synaptic bumps $g_{g}(t)$: Each bump amplitude is independent of diameter; larger rhabdomeres simply generate more bumps per flash.
- Leak & K⁺ channels ($g_{Rest},g_{Ks}$) are located in the soma and thus scale only with somatic area (the 25%).

5. **Parameter Assignment**

- We base all R1-R6 values on their average rhabdomere diameter (**Supplementary Table 13**): corresponding to the photoreceptor voltage response recording in vivo that we analysed for noise.
- Under neural superposition, we then adjust the mean bump rate to match in vivo recordings.

**IV.11.ii. Modelling High-Frequency Jumping LMC Responses**

We combined six neural-superposed *morphodynamic* (photomechanically moving) photoreceptor inputs, synaptic release, membrane integration, and feedback into a single LMC model to reproduce the characteristic high-frequency jumping under a 200 Hz bursty stimulus. The complete R1-R6→LMC framework (**Supplementary Fig. 32a**) has four modules:

- **R1-R6 photoreceptors**: Six independent stochastic inputs driven by the same light pattern.
- **Synapse**: Tonic + dynamic activation probabilities feeding into stochastic quantal release.
- **Electrical membrane**: Summation of synaptic conductances into the LMC voltage.
- **Feedback**: Voltage-dependent modulation of presynaptic activation.

We hand-tuned all free parameters to ensure the simulated LMC waveform, power spectra, and information rates match in vivo bursty-contrast recordings.

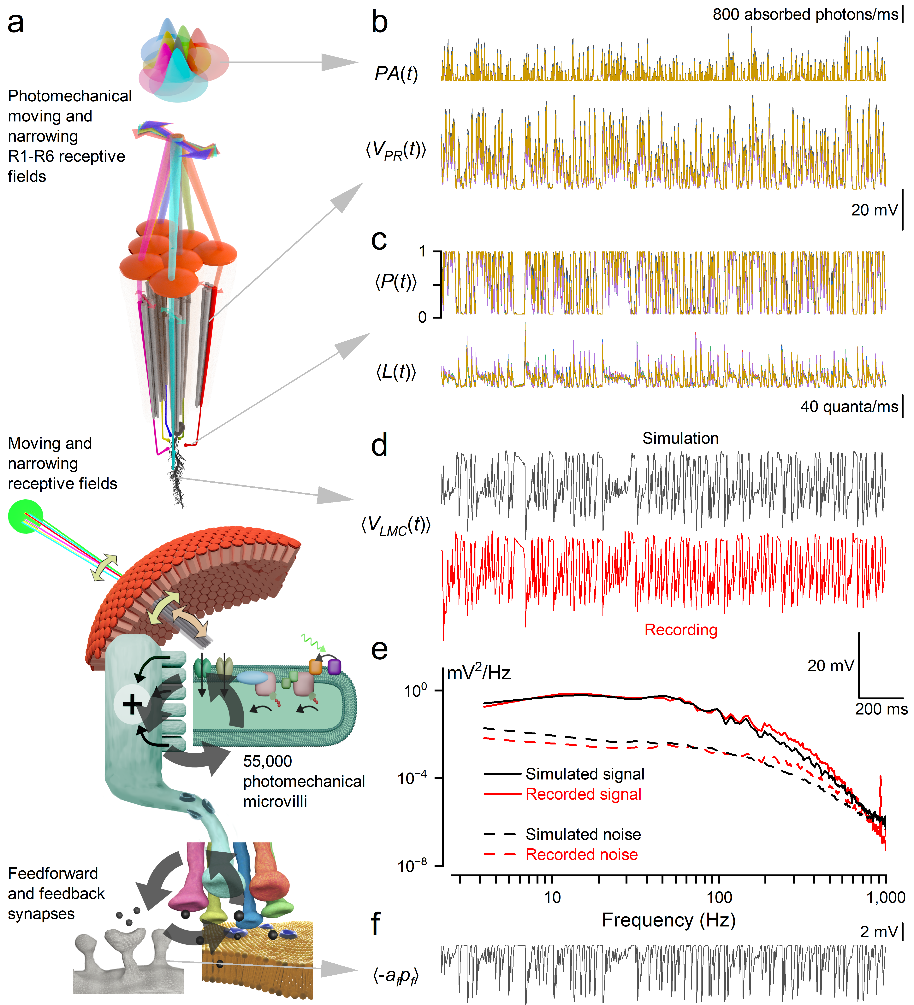
**Supplementary Fig. 32.** **Simulated *Musca* LMC responses of a morphodynamic neural superposition system closely match in vivo recordings for bursty visual stimuli.**

(**a**) Top: R1-R6 photoreceptors from neighbouring ommatidia sample the light stimulus through a photomechanically moving and narrowing “flower pattern” of partially overlapping receptive fields.

Bottom: The model includes two major feedback loops (gray cuircular arrows)

Feedback loop 1: Each receptive field responds to light changes with individual microsaccades, producing variable sequences of quantum bumps and resulting voltage signals. These responses adapt the receptive fields’s movements over time.

Feedback loop 2: Voltage differences between R1-R6 photoreceptors trigger histamine release, which binds to receptors on LMCs, causing chloride influx and hyperpolarising their responses. This, in turn, activates depolarising synaptic feedback to photoreceptors, balancing inhibitory and excitatory synaptic activity and enabling fast, phasic responses with minimal delay.

(**b**) A bursty light contrast stimulus - mimicking a saccadic visual input - is delivered to six R1-R6 photoreceptors (shown in different colours), each from neighbouring ommatidia within a neural superposition cartridge. Due to slight differences in optical axes and photomechanical receptive field movements, the photoreceptors receive slightly different light intensity profiles $PA(t)$, producing distinct mean voltage responses $\left\langle V_{PR}(t) \right\rangle$.

(**c**) The synaptic model computes the postsynaptic activation probability $\left\langle P(t) \right\rangle$, which is used by a sampling model to determine the number of activation events $\left\langle L(t) \right\rangle$at the LMC.

(**d**) Simulated (black) and in vivo recorded (red) average LMC voltage responses $\left\langle V_{LMC}(t) \right\rangle$show strong temporal correlation, capturing key features of the real response dynamics.

(**e**) Power spectral densities of signal (solid lines) and noise (dashed lines) for LMC responses demonstrate close correspondence between simulation (black) and in vivo data (red), validating the model’s frequency-domain accuracy.

(**f**) A feedback term from the LMC to the photoreceptors, denoted $\langle-a_{f} p_{f}\rangle$, dynamically modulates the photoreceptor output, reducing the effective synaptic input during high activity periods.

1. **Stimulus and Protocol**

- **Bursty input**: 3° × 3° dot on a 9 × 9 pixel grid, driven by a 200 Hz, high-contrast sequence.
- **Repeats**: 16 two-second trials, discarding the first to avoid nonstationary adaptation.
- **Ray tracing**: Computed each photoreceptor’s $LI(t)$ (**Section IV.3**).

2. **Photoreceptor Inputs**

- **Stochastic sampler**: Six R1-R6 models, using previously validated parameters (**Supplementary Fig. 32a**).
- **R1-R6 inputs show amplitude and speed differences**: Due to varying rhabdomere sizes and ommatidial positioning (Juusola et al^1^).
- **Output**: Six distinct activation probability traces $P(t)$ (**Supplementary Fig. 32a, b**).

3. **Synaptic Module**: Simulates the probability of postsynaptic site activation from each R1-R6 input.

- **Local independence**: Because each R1-R6 photoreceptor has a unique membrane voltage, we simulate its synapse to the LMC independently (**Supplementary Fig. 32b**)
- **Tonic + Dynamic**: $P\left( t \right)=p_{ss}+(1-p_{ss})\times p_{d}(t)$ (Eq. 46; **Section IV.11**).
- **All-or-nothing bursts**: Bright inputs drive a bimodal $P\left( t \right)$ with peaks at $p_{ss}$ and 1, clipping the extremes of $V^{PR}$.
- **High-frequency jumping**: Squared-off $P\left( t \right)$ injects high-frequency (>500 Hz) components.
- **Dim light**: No saturation - $P\left( t \right)$ remains graded.
- **Decay**: Modelled by a Hill‐type function (half-activation at $p_{d}=0.05$, yielding a constant relaxation rate and linear LMC depolarisation (**Supplementary Fig. 32b**).

4. **Stochastic Sampling of Release**

- **Sites per photoreceptor**: Fitted to ~ 390 (vs. ~ 250 in *Drosophila*¹⁹), reflecting tonic silencing of many sites.
- **Microsaccade-induced overshoot**: Tonic release causes transient dips below $p_{ss}$ after negative contrast.
- **Fast quantal kinetics**: Zero-latency bumps with brief refractory periods (mean ~10 ms; $n_{r}<0.1$, $\tau_{r}\approx800 ms$; **Supplementary Table 13**).
- **Empirical LMC recordings constrain model parameters**:
- **Noise analysis** of bursty‐stimulus responses yields quantal voltage‐bump amplitudes and latency distributions (**Section IV.4.**, **Supplementary Fig. 28**).
- **Current‐clamp recordings** from other LMCs provide initial estimates of membrane channel conductances, which are then refined by fitting.
- **Bursty‐stimulus comparisons**, with and without histaminergic input, isolate the membrane potential dynamics and constrain the leak and synaptic conductance values.

5. **Membrane Integration**

- **Voltage saturation**: Large quantal conductances quickly drive $V^{LMC}$ toward its saturated hyperpolarisation, explaining constant amplitude across stimulus sizes.
- **Time constant**: Small enough to pass high-frequency “jumps.”

6. **LMC→Photoreceptors (Synaptic Feedback)**

- **LMC hyperpolarisation-driven**: $p_{f}(t)$ reduces effective $V^{PR}$, raising the release threshold and preventing early saturation.
- **Dynamics**: Follows the derivative of LMC activity, but is low-pass filtered by $\tau_{f}$ (**Supplementary Fig. 32e**).

7. **Model Fitting and Validation**

We hand-tuned all modules together - because each influences the others - to reproduce key in vivo LMC metrics:

- **Synaptic model**: postsynaptic activation probability parameters
- **Stochastic sampling**: number of synaptic sites and refractory distribution
- **Membrane model**: capacitance, leak conductance, and delayed-rectifier $K^{+}$conductance
- **Feedback loop**: all voltage-dependent feedback parameters

We adjusted these parameters until the simulations simultaneously matched:

- LMC voltage waveforms (**Supplementary Fig. 32c**)
- LMC signal and noise power spectra (**Supplementary Fig. 32d**)
- LMC information rates (**Supplementary Fig. 38**)
- LMC mean quantal bump rates

This integrative fit naturally produces the “high-frequency jumping” behaviour - fast, clipped synaptic input driving a rapid membrane response with adaptive feedback.

8. **Mechanical Explanations,** **Observations and Clarifications**

***A key factor underlying high-frequency jumping is the nonlinear clipping of photoreceptor signals during synaptic transfer***. Because the synaptic activation range is narrower than the photoreceptor voltage range ($V_{PR}$), both the lowest and highest voltages are truncated (see **Supplementary Fig. 25** for a schematic). This squaring of the signal generates strong high-frequency components (>500 Hz) in LMC responses, driving high-frequency jumping. For small photoreceptor voltage fluctuations to bursty stimuli, the synaptic activation probability $P(t)$ takes intermediate values. Synaptic feedback to photoreceptors, together with pre- and postsynaptic refractory processes, continuously reshapes the effective activation range, dynamically matching it to incoming signals.

Under dim conditions, the synaptic model does not saturate, thereby avoiding clipping effects. The best fit for the decay of $P(t)$ back to $p_{ss}$ was achieved using a Hill decay function, which maintains a constant relaxation rate even for small $P(t)$ values when using a saturated Hill function (half-activation at $p_{d}=0.05$). This results in linear LMC depolarisation (**Supplementary Fig. 32b**).

The LMC stochastic sampling model simulates conductance quantal releases from $P(t)$ for each of the six photoreceptors. For the best fit, we found that each photoreceptor has approximately 390 postsynaptic sites - significantly larger than the ~250 per photoreceptor observed in fruit flies^9^. This discrepancy arises because tonic release ensures that the number of active synaptic sites remains substantially smaller than the total number of available sites.

Tonic release enables the depolarised overshoot following negative light contrast. This occurs because the number of active sites is lower immediately after deactivation than in complete darkness, while $P=p_{ss}$ remains unchanged, leading to fewer quantal activations. Noise analysis reveals a fast conductance quantal response with no latency (**Supplementary Table 13**).

**Supplementary Fig. 28c** illustrates the LMC biphasic impulse response, which arises due to steady-state release and refractoriness (see the short pulse response to bright light, **Supplementary Fig. 1a**). The rapid depletion of postsynaptic sites results in a derivative-like effect in bright light. The shape of the refractory distribution determines how quickly this depletion occurs. Under dim conditions, the number of active sites is not depleted, resulting in a low-pass response, as the refractory process is sufficiently fast to restore activations.

LMC refractoriness is characterised by a significantly low n value (<0.1), whereas $\tau_{r}$ is large (~800 ms). The mean refractory time, calculated as $n_{r}\times\tau_{r}$, is approximately 10 ms (**Supplementary Table 13**). This prolonged refractory period occurs because the postsynaptic side of the synapse can be reactivated by histamine vesicle fusion at the presynaptic terminal. When multiple vesicles are available presynaptically, the refractory effect is immediate. However, when the activation pool is depleted, pool replenishment occurs at a much slower rate^142^.

The LMC electrical membrane integrates conductance quantal responses to simulate LMC membrane voltage (**Supplementary Fig. 32C**). We found that the membrane potential saturates under reasonable stochastic sampling activation because the quantal histamine-induced conductance response is large relative to passive resistance. This results in the LMC amplitude response remaining constant across different stimulus sizes, while the response speed increases with faster activation. Additionally, the resting membrane time constant is small enough to support high-frequency jumping.

The feedback model, based on the synaptic response to LMC membrane potential, activates upon LMC hyperpolarisation, reducing $P(t)$. The primary effect of feedback is synaptic activation within a usable range, preventing $P(t)$ from saturating too easily by adjusting the clipping point (**Supplementary Fig. 32e**). Under bright, bursty stimulation, the feedback follows the derivative of LMC statistics, but with a slightly reduced spectrum due to the feedback time constant limiting response speed.

From a functional and mathematical perspective, both synchronised vesicle release and short-term synaptic depression impose state-dependent constraints on synaptic transmission that are formally included to the refractory synaptic information sampling framework described in this study. In all these cases, recent transmission events transiently reduce synaptic availability, thereby shaping the timing, frequency content, and gain of postsynaptic responses. Our model explicitly captures these effects at the level of information flow, without committing to a single microscopic implementation.

Importantly, the present work is deliberately framed at the level of effective synaptic dynamics, showing how refractory, state-dependent transmission - regardless of its specific molecular origin - can redistribute power toward higher frequencies under bursty, saccade-like input. Thus, synchronised vesicle release or synaptic depression are not alternative explanations to high-frequency jumping, but rather plausible mechanistic substrates through which the same functional principle could be realised.

**IV.11.iii. Model parameter effects on LMC output**

Using the same bursty light stimulus, we examined how different biophysical parameters influence the LMC voltage response by comparing each case to the fixed morphodynamic neural superposition control model (**Supplementary Fig. 33a**).

**Case 1: No Microsaccades** (**Supplementary Fig. 33b**)

Setting the maximum microsaccade activation to zero ($F_{max}=0$) fixes photoreceptor receptive fields in place. Without microsaccades, low-frequency photoreceptor noise decreases, as these movements typically induce stochastic shifts^1^. High-frequency jumping redistributes LMC noise towards higher frequencies. Overall, both photoreceptor and LMC information rates are slightly elevated.

**Case 2: Single Photoreceptor** (**Supplementary Fig. 33c**)

Here, the LMC receives input from only one R3 photoreceptor, reducing the total number of synaptic inputs from 6 × 390 to just 390. This diminishes the LMC voltage response and lowers the information rate.

**Case 3: No LMC Feedback** (**Supplementary Fig. 33d**)

LMC feedback to photoreceptors is removed by setting $a_{f}=0 mV$. Without feedback, the simulated LMC potential deviates more from in vivo recordings, as LMC activation becomes more saturated. This saturation results in slightly lower information rates compared to normal conditions.

**Case 4: Synaptic Model with Shallow Steady-State Activation** (**Supplementary Fig. 33e**)

To demonstrate the role of postsynaptic activation in high-frequency jumping, we reduce the synaptic model's steady-state activation slope by setting $V_{0}=-40.9 mV and V_{d}=7.5 mV$, adjusting $V_{0}$ to maintain the midpoint of the steady-state equation. This results in LMC membrane responses that resemble photoreceptors, with lower information rates.

**Case 5: No Tonic LMC Activation** (**Supplementary Fig. 33f**)

In the synaptic model, tonic postsynaptic activation probability drives a sustained LMC voltage response. Silencing this activation by setting the tonic postsynaptic activation probability to zero ($p_{ss}=0$), the LMC membrane potential stabilises at a depolarised level, determined by passive and delayed rectifier conductance at rest. As a result, the LMC loses its ability to generate a negative contrast response to bursty stimuli while retaining the positive contrast response.

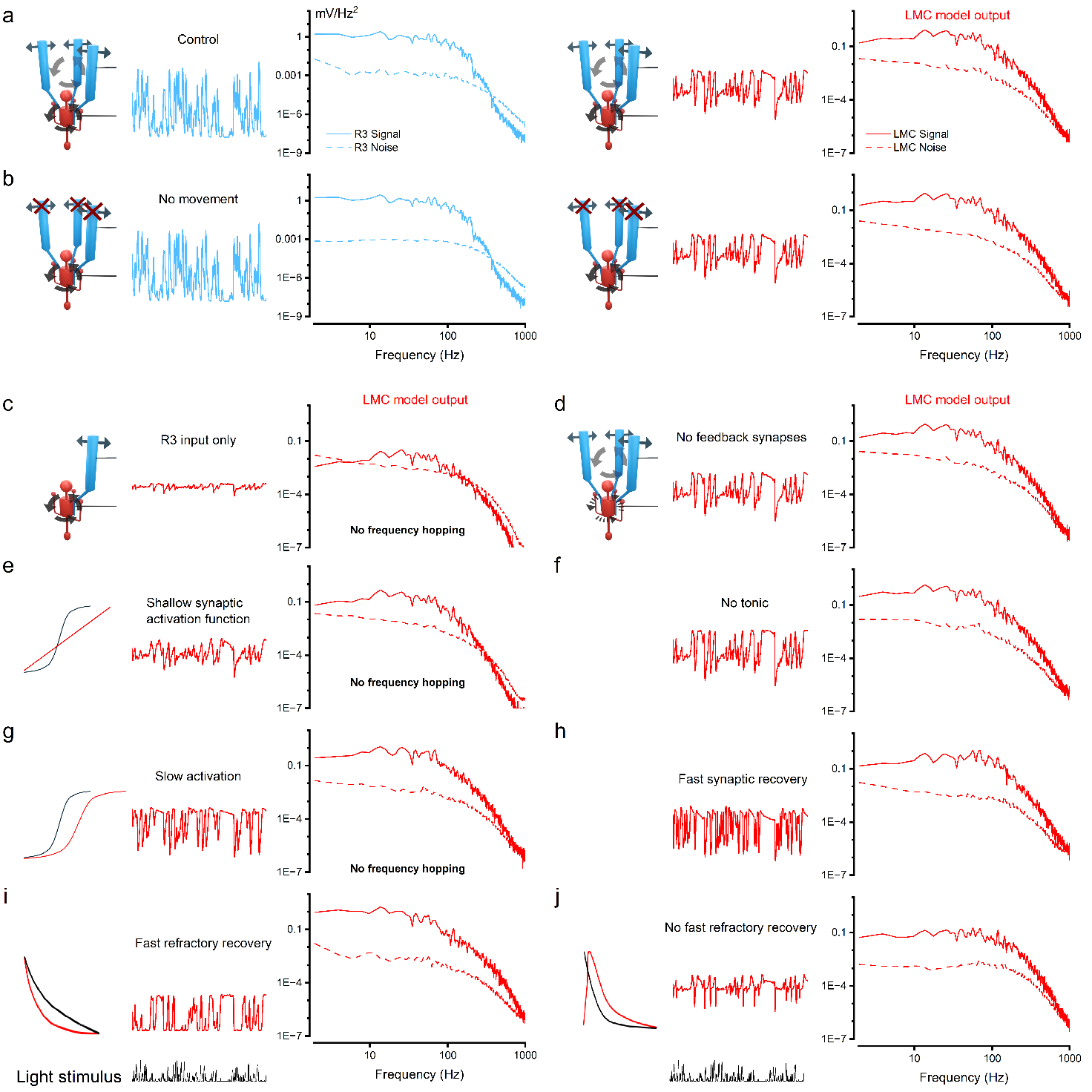
**Supplementary Fig. 33. Effects of modifying biophysical components on LMC output.** Simulated responses of the LMC model to bursty light stimuli under different biophysical configurations, compared to the control model (**a**), which includes morphodynamic neural superposition, feedback, and microsaccades. Each case isolates a key component to assess its contribution to LMC signalling dynamics.

(**a**) Control: Full model with fitted parameters and six-input morphodynamic superposition.

(**b**) No Microsaccades: Receptive fields are static ($Fₘₐₓ$ = 0). Without microsaccades, low-frequency photoreceptor noise is reduced, and high-frequency jumping shifts noise to higher frequencies. Slightly higher information rates are observed.

(**c**) Single Photoreceptor Input: LMC receives input from only one R3 photoreceptor. With fewer synapses (390 vs. 6 × 390), the voltage response and information rate are both reduced.

(**d**) No LMC Feedback: Feedback to photoreceptors is removed ($a_{f}$ = 0 mV), increasing LMC saturation and reducing response fidelity and information rate.

(**e**) Shallow Synaptic Activation: Postsynaptic activation slope is reduced. LMC responses become more photoreceptor-like, with diminished gain and lower information rates.

(**f**) No Tonic Activation: Tonic postsynaptic activation is silenced ($pₛₛ = 0$), causing the LMC membrane to stabilise at a depolarised potential and lose negative contrast responses.

(**g**) Slow Synaptic Activation: Activation time constant is increased ($\tau_{p} = 10 ms$), delaying response onset and increasing low-frequency contributions to the signal spectrum.

(**h**) Fast Synaptic Recovery: Recovery time constant is shortened ($\tau_{n} = 1 ms$), enhancing high-frequency signal content and increasing the saccadic character of LMC responses.

(**i**) Fast Refractory LMC Sampling: LMC refractory time is reduced ($\tau_{r} = 5.4 ms$), leading to a squarer, more synaptic-like voltage profile.

(**j**) Delayed Refractory Recovery: Gamma-distributed refractory response ($n_{r} = 1.6$, $\tau_{r} = 9 ms$) alters spike timing and reduces low-frequency signal power, increasing spike sharpness during contrast changes.

Simulations of various modifications for the LMC model compared to the control case with the fitted model. See text for description of each model.

**Case 6: Slow Activation of the Synaptic Model** (**Supplementary Fig. 33g**)

Increasing the synaptic activation time constant ($\tau_{p}=10 ms$) slows response onset: the synaptic model activates more slowly while still exhibiting high-frequency jumping. The increased activation time constant shifts the midpoint of steady-state activation to a higher photoreceptor voltage, prolonging steady periods with spikes in $P(t)$ and LMC voltage. As a result, the signal spectrum shows an increased low-frequency contribution.

**Case 7: Fast Recovery of the Synaptic Model** (**Supplementary Fig. 33h**)

By decreasing the synaptic recovery time constant ($\tau_{n}=1 ms$), accelerates recovery, resulting in more saccadic $P(t)$ and LMC voltage profiles and enhanced high-frequency signal content (high-frequency jumping). Like Case 6, this also shifts steady-state activation to higher photoreceptor voltages.

**Case 8: LMC Sampling Model with Fast Refractory** (**Supplementary Fig. 33i**)

Reducing the LMC refractory time constant ($\tau_{r}=5.4 ms$) lessens adaptation, making the LMC voltage response resemble that of the synaptic model $P(t)$, becoming more square-like in time.

**Case 9: LMC Sampling Model Refractory Without Immediate Recovery** (**Supplementary Fig. 33j**)

Adjusting the gamma distribution parameters of the LMC refractory period ($n_{r}=1.6077$, $\tau_{r}=9 ms$) preserves the mean refractory period ($n_{r}\times\tau_{r}$) of the original model but delays recovery. This produces spikier voltage traces in response to contrast changes and reduces low-frequency signal power.

**IV.11.iv. Testing the Model with Different Stimulus Patterns**

We tested the model using fifteen different light intensity time series stimulus patterns, spanning a wide range of contrasts and temporal bandwidths, similar to those used in the in vivo recordings (*cf*. **Supplementary Fig. 2** and **5**). These simulations, performed using the full morphodynamic neural superposition model with no free parameters, closely replicated the recorded photoreceptor and LMC response dynamics. Because the mean light level changes between the stimulus types, photoreceptor model parameters were fixed using the same type of recording as needed in simulations (**Supplementary Tables 13** and **16**).

**Supplementary Fig. 31** and **33** show simulated voltage responses of R1-R6 photoreceptors and LMCs, respectively, to repeated stimulation with all 15 tested light-intensity time series. **Supplementary Fig. 33** and **34** present the corresponding signal-to-noise ratios, probability density functions, and information transfer rates**.** These simulations, constrained entirely by the above-described biophysical functions without free parameters, reproduce encoding dynamics of R1-R6 and LMCs that closely match those observed in the intracellular recording (cf. **Supplementary Fig. 2-7**).

**Supplementary Table 16. Parameters for simulating voltage responses in middle bursty and GWN conditions**

| Mean response, $\mu$ | 16.9 mV | 15.1 mV |
| --- | --- | --- |
| Conductance quantal response gamma decay, $\tau_{g}$ | 0.30 ms | 0.30 ms |
| Conductance quantal response gamma exponent, $n_{g}$ | 4.00 | 4.00 |
| Conductance quantal response average duration, $T_{g}$ | 2.2 ms | 2.2 ms |
| Area of voltage quantal response, $s_{g}$ | 1.91*10^-14^ Ss | 1.58*10^-14^ Ss |
| Quantal bump rate, (λ) | 309,000 quanta/s | 351,000 quanta/s |
| Passive reverse potential, $E_{rest}$ | -73 mV | -70 mV |
| **LMC model parameter** |  |  |
| Synaptic model half activation, V0 | -48.9 mV | -46.9 mV |
| Refractory gamma decay, $\tau_{r}$ | 292.5 ms | 292.5 ms |
| Postsynaptic sites active in darkness, $p_{ss}$ | 0.0132 | 0.0132 |

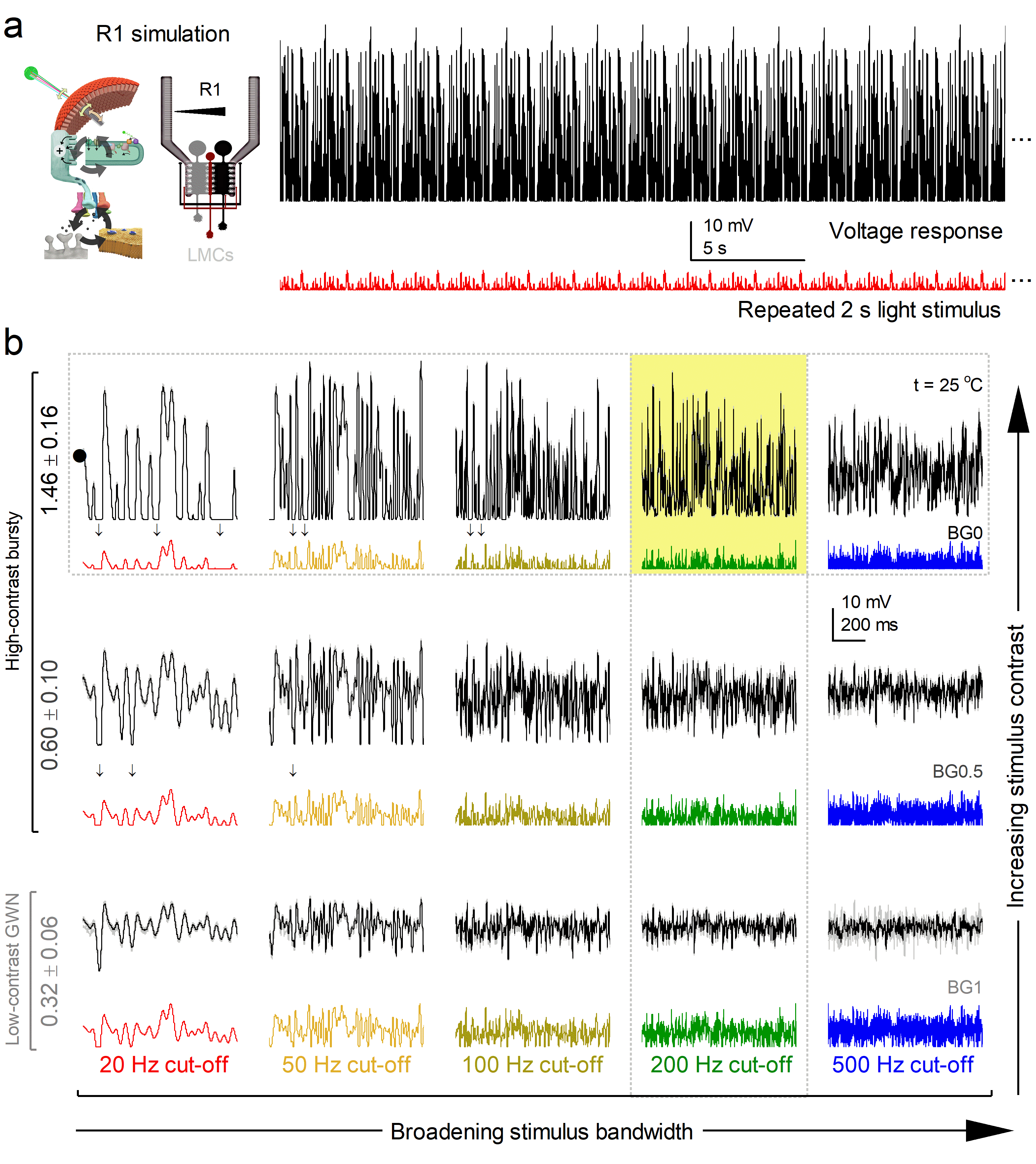
**Supplementary Fig. 34. High-contrast “saccadic” bursts maximise R1 photoreceptor responses with minimal noise.**

(**a**) Left: Schematic of a multiscale morphodynamic neural superposition system model for an R1 photoreceptor in the *Musca* eye. Right: Example trace showing the repeated high-contrast bursty stimulus (red) and the corresponding photoreceptor voltage response simulation (black) at 20 Hz bandwidth.

(**b**) Simulated R1 responses to a range of stimuli varying in both contrast and temporal bandwidth. Stimuli range from high-contrast, saccade-like bursts (BG0) to low-contrast Gaussian white noise (GWN, BG1), each delivered at cut-off frequencies of 20, 50, 100, 200, and 500 Hz. Black and grey traces show the mean responses; thin, light grey traces represent 20 individual responses to different stimulus exemplars. Coloured traces below indicate the 15 different stimulus waveforms. The yellow box highlights the stimulus condition (200 Hz bandwidth) that elicits maximal information transmission. Arrows indicate some dark intervals embedded in the saccadic stimuli. The dotted horizontal and vertical boxes indicate stimulus conditions used in **Supplementary Fig. 33a-d** and **32e-h**, respectively. All data were recorded from the same photoreceptor cell, in a systematic sequence starting with low-contrast GWN (20-500 Hz) and ending with high-contrast bursts (20-500 Hz).

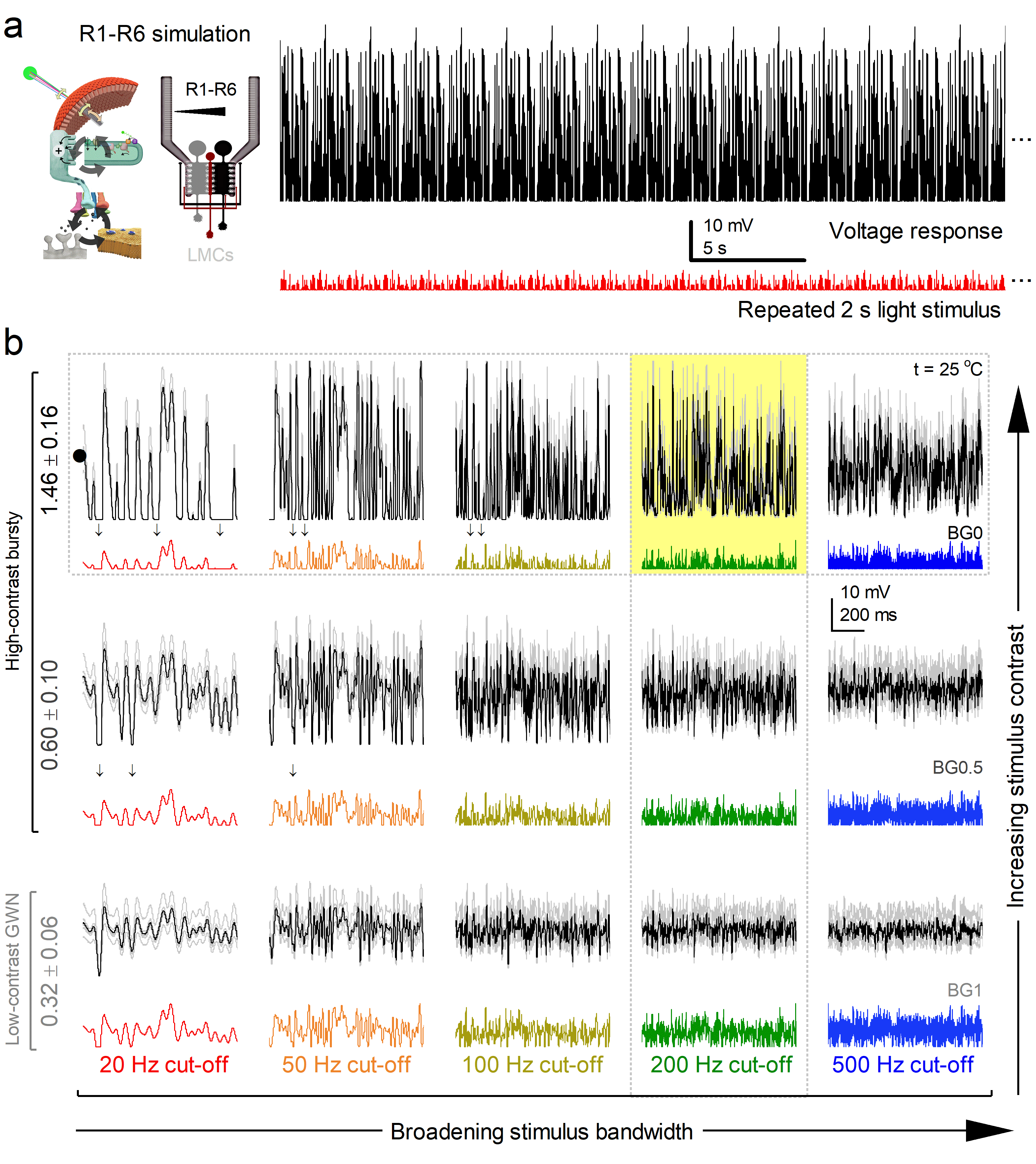
**Supplementary Fig. 35.** **R1-R6 photoreceptors respond most effectively to high-contrast saccadic stimuli.**

(**a**) Left: Schematic of a multiscale morphodynamic neural superposition system model for R1-R6 photoreceptors in the *Musca* eye. Right: Example trace showing a repeated high-contrast saccadic stimulus (red) and the averaged simulated voltage response (black trace) of R1-R6 (grey), from the neighbouring ommatidia at 20 Hz bandwidth.

(**b**) Simulated R1-R6 responses to a stimulus set varying in contrast and temporal bandwidth, ranging from high-contrast saccadic bursts (BG0) to low-contrast Gaussian white noise (GWN, BG1), delivered at 20, 50, 100, 200, and 500 Hz. Black and grey traces show the mean responses across all photoreceptors, while thin, light grey lines represent individual responses. Coloured traces below indicate the 15 different stimulus waveforms. The yellow box highlights the stimulus condition (200 Hz bandwidth) that elicits maximal information transmission across the population. Arrows mark dark intervals in the saccadic stimuli.

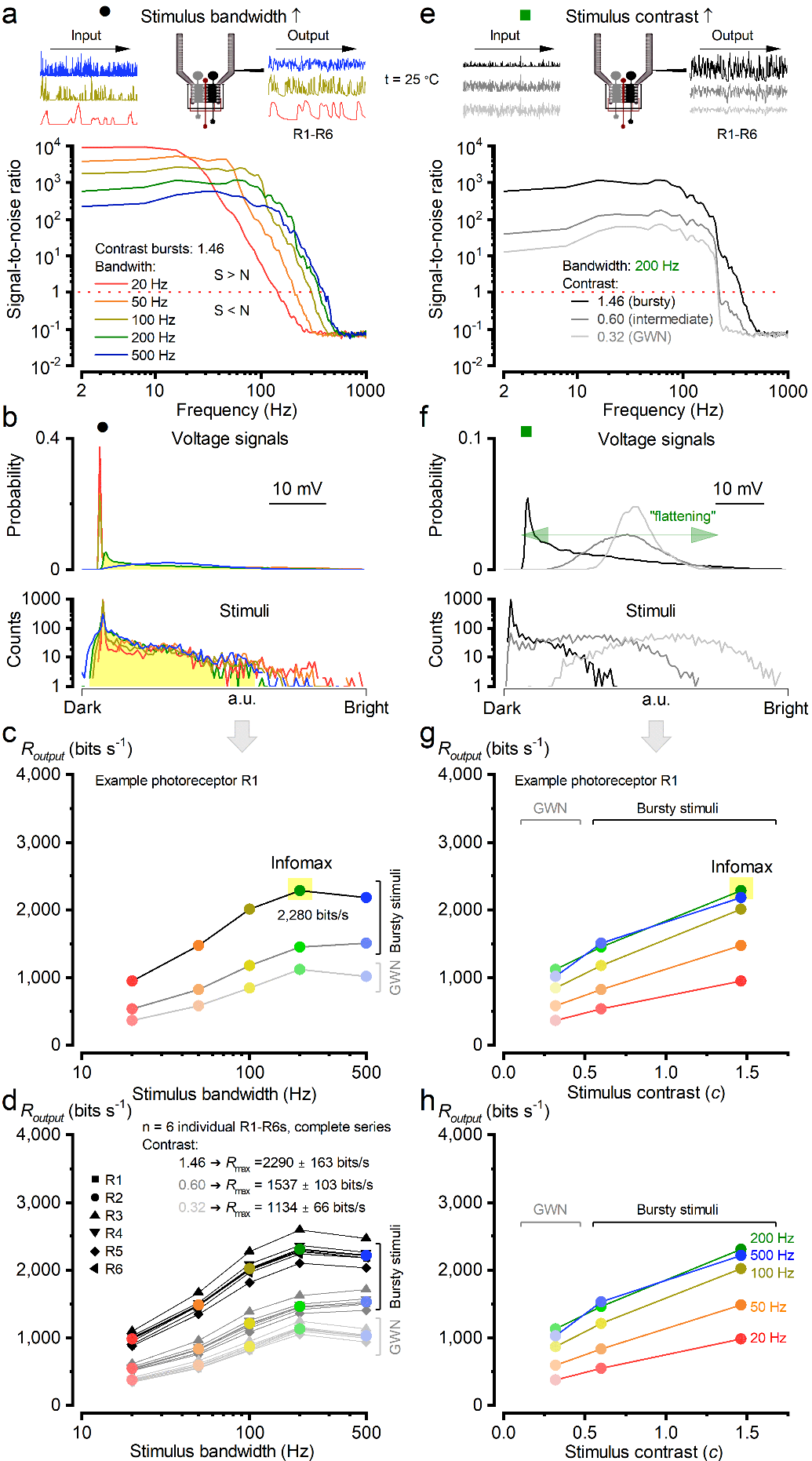
**Supplementary Fig. 36. SNR analyses of the simulated R1-R6 responses**.

(**a**) Simulations’ signal-to-noise ratio, ${SNR}_{PR}(f)$, when increasing the bandwidth from 20 Hz to 500 Hz.

(**b**) Top: All simulations probability density functions, PDFs, are skewed except for 500 Hz. Bottom: Stimulus intensity distributions are skewed. At 200 Hz, the photoreceptor had the broadest frequency and voltage distributions.

(**c**) The information transfer rates, $R_{output}$ (from a representative photoreceptor simulation in **Supplementary Fig. 34**) were best for high-contrast "saccadic" stimuli, and it peaked at 200 Hz bandwidth (marked with a yellow box). The information transfer rates were calculated using the Shannon formula^41^.

(**d**) $R_{output}$ was calculated for all the simulations (n = 14) to all the stimulus bandwidths. "Saccadic" bursts drove the maximal information transfer, which peaked at 200 Hz. We provide $R_{output}$ for R1-R6, each differing in size and in the number of microvilli.

(**e**) Response ${SNR}_{PR}(f)$ at 200 Hz stimulus bandwidth when increasing the contrast from *c* ~0.32 to *c* ~1.46, being the highest for high-contrast bursty stimuli.

(**f**) Top: Response PDFs are approximately Gaussian for low- and mid-contrast stimuli, but become skewed under high-contrast bursty stimulation. This skew expands and flattens the photoreceptor output range, enhancing signalling efficiency. Bottom: Stimulus intensity distribution is Gaussian for low-contrast stimuli but skewed for mid and high-contrast stimuli.

(**g**) Same as in **c**, but comparing against different contrast levels.

The simulations reproduce encoding dynamics closely matching the intracellular recordings (**Supplementary Fig.** **4**).

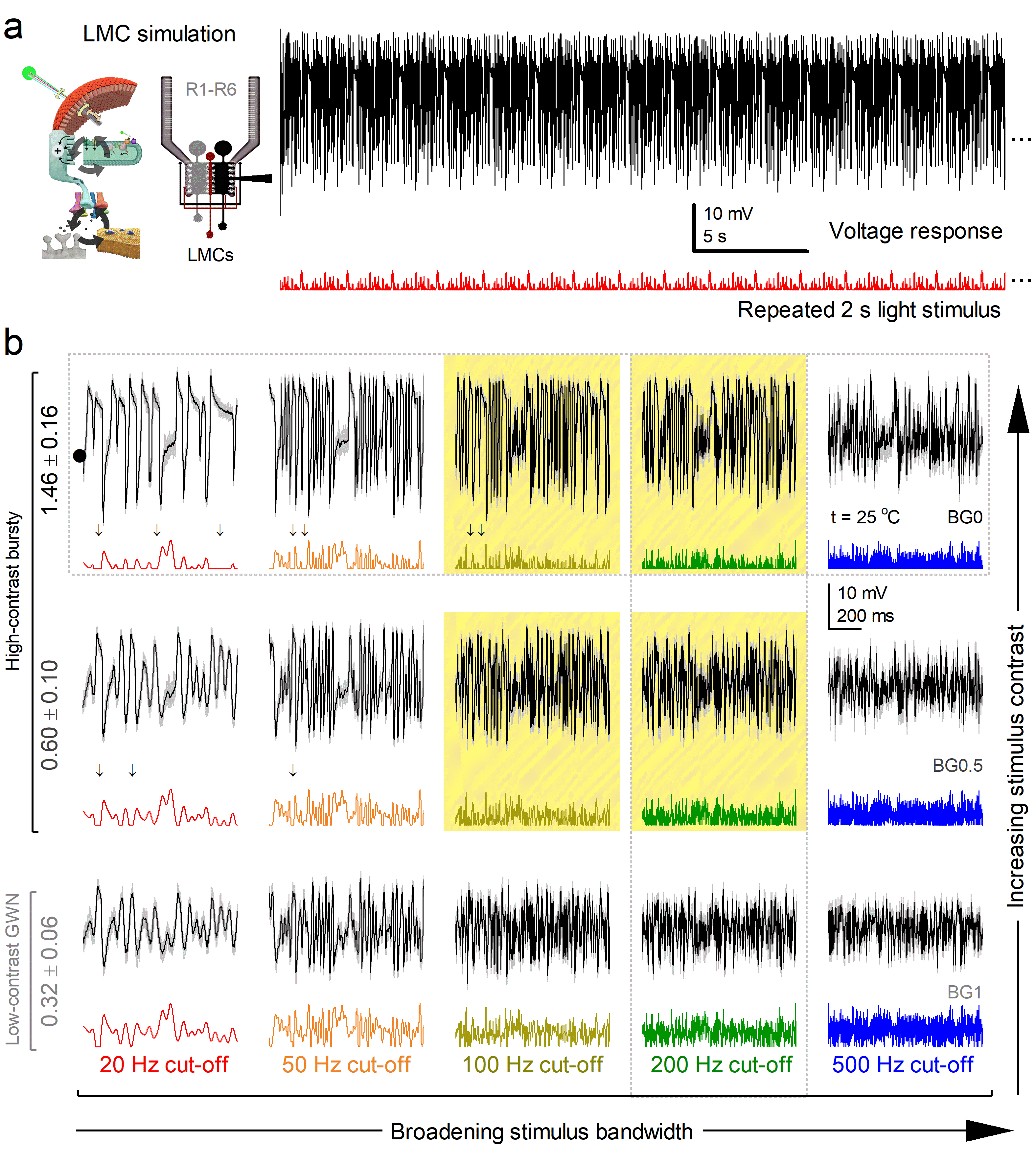
**Supplementary Fig. 37.** **Mid and high-contrast "saccadic" bursts maximise LMC's response.**

(**a**) Left: Schematic of a multiscale morphodynamic neural superposition system model for an LMC. Right: An example of a repeated high-contrast bursty stimulus (red trace) and a response (black trace) at 20 Hz bandwidth.

(**b**) Simulated LMC voltage responses ranging from high-contrast "saccadic" bursts (BG0) to low-contrast GWN stimuli (BG1) in different cut-off frequencies, i.e. bandwidth patterns (20, 50, 100, 200 and 500 Hz). Mean (thick black and grey traces) and 25 individual responses (thin, lightly coloured) to 15 different stimuli (colourful traces beneath the responses). Yellow boxes: similar maximum information rates were carried by three sets of responses to different bursty stimulus patterns. Arrows: dark intervals in saccadic stimuli. Vertical dotted and horizontal rectangle: responses for bandwidth and contrast used in **Supplementary Fig. 35a** and **34e**, respectively. The recordings are from the same LMC.

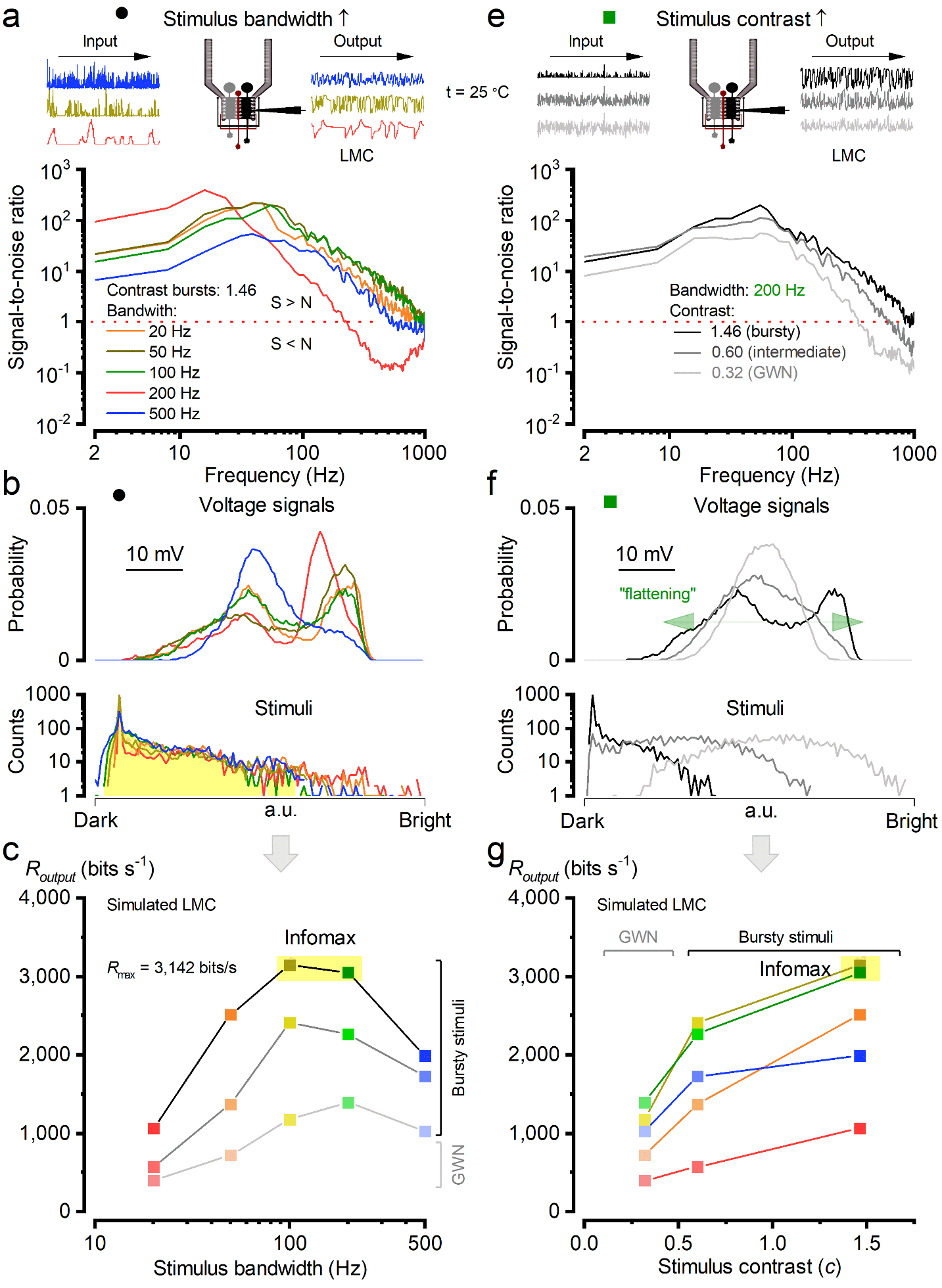
**Supplementary Fig. 38. SNR analyses of the simulated LMC responses**.

(**a**) Simulation signal-to-noise ratio, ${SNR}_{LMC}(f)$, to high-contrast bursty stimuli when increasing the bandwidth from 20 Hz to 500 Hz.

(**b**) Top: Simulation probability density functions (PDFs) to high-contrast bursty stimuli are not Gaussian except broadly for 500 Hz. Bottom: Stimulus intensity distributions are massively skewed.

(**c**) The information transfer rates, $R_{output}$, from a representative LMC simulation in **Supplementary Fig. 37**) were best for mid-contrast and high-contrast "saccadic" stimuli, and they peaked at 100 Hz and 200 Hz (marked with a yellow box). $R_{output}$was calculated using the Shannon formula^41^.

(**d**) Simulated ${SNR}_{LMC}(f)$ at 200 Hz stimulus bandwidth when increasing the contrast from *c* ~0.33 to *c* ~1.29. ${SNR}_{LMC}(f)$ is the highest for high-contrast bursty stimuli.

(**e**) Top: Response PDFs are approximately Gaussian for low- and mid-contrast stimuli, but become double-peaked under high-contrast bursty stimulation. This “skew” expands and flattens the LMC output range, enhancing signalling efficiency. Bottom: Stimulus intensity distribution is Gaussian for low-contrast stimuli but skewed for mid and high-contrast stimuli.

(**g**) Same as in **c**, but now comparing against different contrast levels.

The simulations reproduce remarkably similar encoding dynamics to those seen in the corresponding intracellular recordings (**Supplementary Fig.** **7**).

**Pulse Stimuli**

Pulse stimuli are commonly used to characterise the response properties of visual neurons. We simulated R1-R6 and LMC voltage responses to light pulses of varying duration and intensity using the model validated with “bursty” stimulus (**Supplementary Fig. 32**).

A short bright pulse (**Figure 1c**, main paper) (10 ms, total photon count: 7,800) evokes a biphasic LMC response with distinct on- and off-transients, reflecting the LMC's differentiating function^8,13,16,143^. The synaptic model $P(t)$ induces a transient LMC voltage response, causing the LMC to reach its peak faster than the photoreceptor. Following the initial response, the synaptic sampling model rapidly reduces postsynaptic activations, leading to LMC depolarisation.

Similarly, a longer bright pulse (1,200 photons/ms, 1 second duration) elicits the characteristic differentiating response for both light onset and offset (tested - no figure included here).

For dim pulses (120 photons/ms), the adaptive behaviour of the LMC sampling model weakens. As a result, pulse responses revert to a more amplitude-based response profile for both short and long pulses^16,125^ (tested - no figure included here).

**Demonstrating LMC Feedback (tested: No figure, No current simulation)**

To show the functional role of synaptic feedback in preventing LMC saturation, we simulated responses to two short light pulses (10 ms) separated by a 10 ms interval, with and without feedback. With feedback enabled, $P(t)$ shows a more pronounced dip between the pulses, leading to greater voltage differences in the LMC response. Notably, feedback mirrors the shape of the LMC voltage trace, consistent with its differentiating role.

**Spatial Stimulation**

To model responses to spatially structured stimuli, we simulated a single LMC and its six presynaptic photoreceptors, each from a distinct ommatidium. A virtual screen (15 × 15 mm, 150 × 150 pixels) was placed 5 cm in front of the eye, where the field of view of all six photoreceptors were at centre of the screen, if not stated otherwise.

We raytraced all photoreceptors in six ommatidia, calculating their microsaccadic movements and voltage responses using a model validated under bursty stimulation (**Supplementary Fig. 32**). LMC responses were computed using the same calibrated model, ensuring light levels were matched to previous conditions. Consequently, both photoreceptors and LMCs continuously adapted to changing light conditions due to the model's inherent properties when processing bursty saccadic stimuli.

**Hyperacuity in the model**

To evaluate hyperacuity (**Figure 4h**, main paper), we simulated responses to small dots (0.7° × 0.7°) moving across the centre of the screen at various angles, replicating the in situ experimental setup. The distance between dots was varied in 0.7° increments, and the dot speeds matched those used in the recordings. The resulting responses closely matched in vivo data.

For clipping experiments - where the moving stimulus becomes occluded, photoreceptor receptive fields were shifted laterally by 7.6 mm. This placed their centres 0.1 mm outside the visible screen area, although parts of the receptive fields remained within view. As a result of this occlusion, the effective receptive field size was “physically reduced” (see also **Supplementary Fig. 24**). We confirmed this by simulating a case without micromovement by setting $F_{max}=0$.

In this configuration, the microsaccades of the six ommatidia became synchronised, since all six photoreceptors in neural superposition viewed the same central stimulus. In typical spatial stimulation, however, photoreceptor microsaccades are not synchronised because each ommatidium samples a different region of visual space.

**Model Limitations and Excluded Properties**

The model does not account for the following properties:

#### Ion Channels

- **LMC Na^+^ Channel^56^**: This channel exhibits the potential for action potentials following extreme hyperpolarisation. However, our model does not generate action potentials in response to bursty stimuli that lack action potentials, even though such responses are theoretically possible.
- **Photoreceptor Na^+^/Ca^2+^ Exchanger and Na^+^/K^+^ Pump^65^**: The model does not incorporate these mechanisms, which results in the absence of photoreceptor hyperpolarisation and the silencing of the synapse.
- **Feedback Channels**: Feedback channels are not directly connected to the photoreceptor model and only influence the synaptic model.
- **Histamine Receptor Channels^21^**: Studies show that histamine receptor currents decrease with prolonged histamine stimulation. However, because synaptic activation occurs over a short duration (2.7 ms, **Supplementary Table 13**), the effect of receptor deactivation is negligible.

#### Connectomics

This model only considers the direct connection from R1-R6 photoreceptors to the LMC and its feedback, while omitting:

- **Internal connections within the lamina cartridges^9^**
- **Interconnections between neighbouring cartridges^9^**
- **Gap junctions between photoreceptors^12^**
- **Gap junctions between L1 and L2 neurons^58^**

At present, the global synaptic feedback from LMCs to photoreceptors serves as a functional umbrella for all local synaptic feedback pathways within a lamina cartridge, tuning information flow from a small region of visual space. As more detailed knowledge emerges about the roles of specific local top-down and lateral feedback connections, this global feedback can be reorganised in the model into finer subprocesses.

#### Photoreceptor Adaptation and Noise Analysis

- **Bump Size Adaptation:** We have shown in *Drosophila* photoreceptors that during bright light stimuli, such as contrast bursts or GWN at a bright background, the average quantum bump size remains relatively constant^1^ (i.e. it has light-adapted to a very small size and may only change a little with the light contrast fluctuation). Hence, the model does not account for bump size adaptation in photoreceptors, which leads to inaccuracies under dim light conditions^40^. Each light level requires a separate noise analysis for bump parameters.

#### Feedback Mechanisms

- **Synaptic Feedback:** Simulations indicate that the model lacks synaptic feedback mechanisms necessary to scale prolonged light activation.
  - LMC responses are suppressed during extended background light exposure.
  - LMC feedback to the photoreceptor follows the LMC voltage profile (**Supplementary Fig. 32f**). The highest feedback values occur with large light level changes, meaning feedback is most pronounced at high frequencies.

#### Synaptic and Motor Considerations

- **Simplified Synaptic Model:** The model is inspired by biological mechanisms but does not fully replicate their complexity.
- **Muscle Movements:** Although significant muscle movement was observed in microsaccade recordings, no clear rules were identified for implementation in the model^38^.

**Analysis and simulation methods**

Analysis and simulation are done with MATLAB (Mathworks, USA). The power spectra were done using the Welch method to calculate power spectral density with the Blackman-Harris window (256 points long). All the fits were done using the minimum least square error. Various random values for distributions were generated with pseudo-random MATLAB generators. Image analysis was done with Fiji^144^.

**IV.11.v. Additional Factors to Consider**

**1. Dynamic Encoding by the Morphodynamic Neural Superposition System**

In bright ambient illumination, the morphodynamic neural superposition system functions primarily as a differentiator. Rather than encoding absolute luminance values, it highlights dynamic changes in contrast. If the visual input remains constant, LMC output swiftly returns to baseline, thereby suppressing redundant signals and enhancing transient information.

This temporal processing begins early in the visual pathway. Although ON-OFF opponency is typically attributed to downstream processing in the medulla, our findings suggest that LMC dendritic responses in the lamina already encode both light increments (ON) and decrements (OFF). Thus, what is often modelled as binary ON-OFF signalling in motion detection emerges from a continuous, analogue pre-processes rooted in synaptic dynamics.

**2. Temporal Precision and Synaptic Dynamics**

**High-Frequency Transfer and Reduced Latency**

The morphodynamic synapse enhances temporal resolution by enabling high-frequency information transfer and low-latency responses:

- **Synaptic delay**: effectively 0 ms.
- **Activation time constant**: ~0.1 ms.

Such fast kinetics allow small photoreceptor voltage changes to generate disproportionately large synaptic responses - a hallmark of differentiator circuits that prioritise transients over steady inputs.

**Vesicle Recycling and Release Modelling**

Unlike classical synaptic models that assume vesicle release as a Poisson process or simple exponential decay^121,145^, our synaptic model supports refractory-driven release governed by a gamma distribution. This approach:

- Allows versatile synaptic site replenishment dynamics.
- Eliminates the need for multiple vesicle pools to sustain high release rates (*cf*. Rizzoli^142^).
- Captures realistic short-term synaptic dynamics, including saturation and recovery.

Bird^146^ and Rosenbaum^147^ discuss vesicle depletion in the synapse with spike coding. The stochastic sampling process with steady-state (tonic^65^) postsynaptic activation models the vesicle depletion in the postsynaptic neuron.

Moreover, while some models posit a one-to-one mapping between presynaptic and postsynaptic spike trains^148^, we observe that synaptic transmission in our system exhibits significant processing, including nonlinear gain modulation and frequency-dependent transmission enhancement.

**3. Relevance of Spike Train Statistics**

Traditional rate coding models and Poisson spike train inputs^145,149,150^ are not applicable as gradient LMC codes , thus the model does not have Na^2+^ channels to replicate measured LMC voltages. In vivo sensory spike statistics often deviate from Poisson assumptions^146^, especially under naturalistic input where $1/f$ statistics dominate. As Rosenbaum^147^ noted, the broad spectrum of a Poisson process may overwhelm synaptic filtering capacities. Our model accounts for this by incorporating a more selective and structured refractory release dynamic.

**4. Spatial Aspects of Superposition and Stimulus Geometry**

The spatial configuration of the rhabdomeres in R1-R6 photoreceptors leads to distinct receptive field overlaps and motion parallax. Photoreceptor microsaccades - minute, photomechanical (rapid) shifts in rhabdomere positions - occur asynchronously and vary across photoreceptors due to their differing anatomical orientations. This motion enhances edge localisation and precise time-locking of the LMC response but introduces temporal jitter in photoreceptor activation timing.

**Effects of Spatial Stimulus Geometry**

- Microsaccades triggered by stimulus edges cause effective receptive field narrowing ("clipping"), sharpening spatial resolution^4^ (see **Section II**, **Supplementary Fig. 24**).
- Double-dot stimuli (e.g., white dots 1.4° wide × 0.7° high on a black background) are harder to discriminate than alternating black-white patterns due to adaptation and contrast limitations (**see Section I, Supplementary Fig. 18**).
- Wide bar patterns (~10°) allow adaptation at both photoreceptor and LMC levels, resembling a long pulse stimulus.
- Constant background illumination and absence of microvilli adaptation limit high-frequency jumping under such conditions.

Microsaccades prove more effective in response to brief flashes (rapid onset followed by decay), as these better align with the temporal dynamics of photoreceptor adaptation (microvilli refractoriness).

**5. High-frequency jumping Enables Fast Neural Coding**

LMCs implement high-frequency jumping - a shift in the dominant frequency of information transmission - when stimulated by high-contrast, temporally structured inputs such as saccades or fast flashes.

**Stimulus Dependence**

- High-frequency jumping is observed with both middle-contrast and high-contrast bursty stimuli.
- High-frequency jumping is observed with both short and long bright pulses^8,13,16^.
- Not seen with white noise (limited photoreceptor voltage range)^13,124^ or naturalistic $1/f$ stimuli (owing to its lack of high-frequency components)^8,16^.

**Modelling Limitations of High-Pass Filters**

Simple black-box high-pass filter models^13-15,47,124,151-155^ fail to replicate LMC behaviour due to:

- Inability to implement high-frequency jumping.
- High amplification of photoreceptor noise.
- Lack of dynamic gain adaptation.

Nonlinear kernel-based models with simple feedback structures also fall short^47,124,156^, highlighting the need for multiscale (multicellular) morphodynamic models that couple refractoriness, adaptive feedback, and structural adaptation mechanisms.

**LMC Model Fitting and Parameter Effects**

The LMC model responses were obtained after adjusting several biophysical and synaptic parameters, which collectively influence the voltage waveform, frequency spectrum, noise characteristics, and information rate of the cell’s output. Below, we summarise the role of each key parameter and how it affects LMC voltage responses.

**1. Postsynaptic Structure and Synaptic Dynamics**

- **Number of postsynaptic sites** ($N$):
  - The number of functional postsynaptic release sites (sampling units) modulates both signal amplitude and noise.
  - With fewer sites (small $N$), stochastic variability increases because there are fewer inputs to average out.
  - As $N$ increases, the number of quantal samples rises, boosting the signal-to-noise ratio and information throughput^34^.
  - $N$ may therefore vary across the eye to match regional differences in natural image statistics - for example, in frontal versus dorsal fields during active behaviour.
  - The number of sites also constrains the maximum hyperpolarisation achievable during a light pulse.
- **Rate of quantal bumps**
  - determined by *N* and the mean refractory interval ($\tau_{r}\times n_{r}$)
- **Refractory distribution shape** $(n_{r}):$
  - This shape parameter controls adaptation dynamics.
  - A smaller $n_{r} ,$ while keeping $\tau_{r}\times n_{r}$ constant, slows down adaptation, prolonging hyperpolarisation.
- **Refractory Distribution Shape** $(\tau_{r}):$
  - Smaller $\tau_{r}$ leads to faster depolarisation from overshoot back to resting potential after the off stimulus.

**2. Synaptic Activation Function Parameters**

- **slope of the synaptic activation function** $\boldsymbol{p(t)}$ $(V_{d}$):
  - Smaller $V_{d}$ makes it LMC voltage square-shaped.
  - Squaring the activation function shifts the LMC voltage power spectrum toward higher frequencies.
  - Increasing $V_{d}$ (e.g. by 0.25 mV) requires a compensatory increase in $V_{0}$ (e.g. by 1 mV) to maintain the midpoint of the activation curve.
- **midpoint (threshold) of the activation function** $\boldsymbol{p(t)}{(V}_{0})$:
  - Sets the midpoint (threshold) of the activation function $p(t)$.
  - For optimal behaviour, this value should lie within the dynamic range of R1-R6 photoreceptor voltages.
  - When R1-R6 are near their resting potential, $p(t)$ should approach zero.
- **Activation time constant of** $\boldsymbol{p}\left( \boldsymbol{t} \right) (\tau_{p}$):
  - Primarily affects the speed of hyperpolarisation onset in the LMC response.
- **Recovery time constant of** $\boldsymbol{p(t)}$ ($\tau_{n}$):
  - Determines how quickly the LMC depolarises after a stimulus.
  - Higher values lead to slower recovery and reduced information transfer rates.

**3. Membrane Properties**

- **Membrane capacitance** ($C$):
  - Determines the amplitude of voltage responses via the cell’s surface area.
  - Higher capacitance leads to smaller voltage changes for the same synaptic input.
- **Membrane RC time constant:**

The effective low-pass filtering of the membrane.

- Cannot be too large, as this would attenuate high-frequency signals and distort power spectral content.
- **Reversal Potential** ($E_{in}$):
  - Mainly determines the maximum hyperpolarisation achievable during a light pulse.

**4. Feedback Effects**

- **Synaptic Feedback:**
  - The feedback loop from the LMC to the photoreceptor synapse reduces synaptic activation when the LMC is hyperpolarised, effectively implementing a gain control mechanism.

**V Behavioural Experiments**

**V.1. Housefly methods for leg- and antenna-response videos**

**Animal model**

Adult houseflies (Musca domestica) were obtained from Blades Biological Ltd. (Edenbridge, UK). Upon arrival, flies were transferred to plastic fly boxes with netted tops for ventilation and maintained at room temperature on a 12 h light/dark cycle. No specific licences or permits were required for these experiments, which were conducted in accordance with the ASAB/ABS guidelines for the use of animals in research.

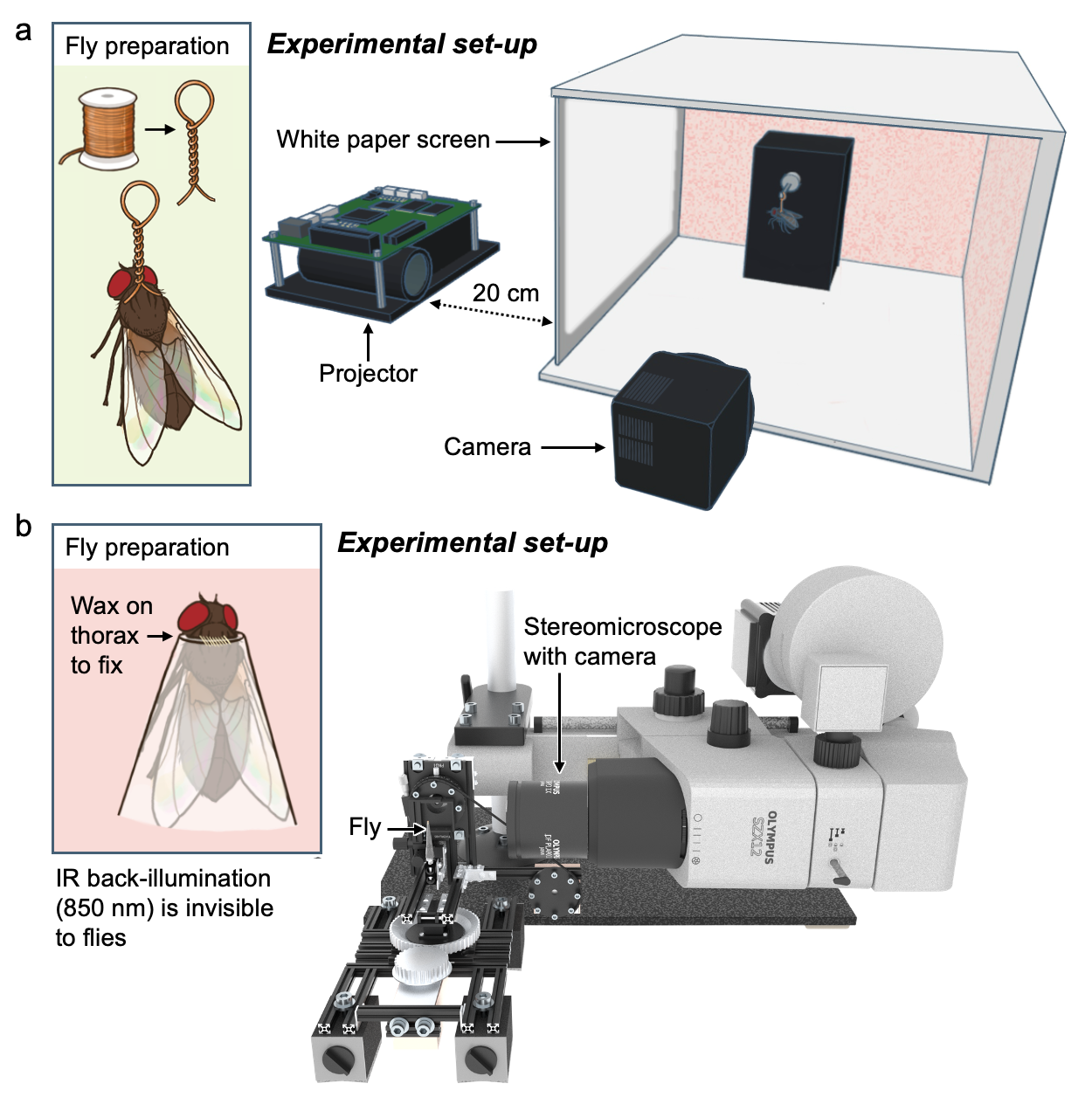
**Supplementary Fig. 39. Experimental set-ups for leg and antenna response assays.**

(**a**) **Leg response experimental set-up.** A length of copper wire was wound into a loop and attached to the thorax of the housefly (see inset). The loop was threaded onto a metal hook in the arena, and pinched so that the fly was facing in the direction of the the projector screen (made of white printer paper). The projector (0.45” DLP Lightcrafter E4500 MKII projector) displayed the looming stimulus using green and blue channels (520 and 460 nm wavelength, respectively) onto this screen. The high-speed Kinetix camera recorded the behaviour at 2500 fps.

(**b**) **Antenna response experimental set-up.** Flies were placed in pipette tips, waxed on the thorax to hold them in place, and mounted on the imaging stage of the MUSA system. This system comprises a sideways-mounted stereomicroscope, which delivers stimuli in the form of UV light flashes from a 365 nm UV LED mounted in the ocular slot. A high-speed camera, mounted to the microscope, was used to trigger the stimulus and acquire images at 1,000 fps.

**Leg response**

*Experimental set-up.* Experiments were conducted in a custom-built fly arena (**Supplementary Fig. 39a**). The stimuli were presented on a 0.45” DLP Lightcrafter E4500 MKII UV (385 nm) + Blue (460 nm) + Green (520 nm) projector (EKB Technologies, Ltd., Bat Yam, Israel), positioned ~20 cm from the paper screen. Green and blue channels only were used to present the stimulus, as the UV would fail to pass through the paper. Videos were recorded at 2500 fps using a Kinetix camera (Teledyne Vision Solutions, USA) and Micro-Manager software^157^. All experiments were conducted at room temperature, under standard white LED lights.

*Stimulus design.* The looming stimulus consisted of 24 black and white images. These images were generated in MATLAB, and uploaded to the projector using Lightcrafter 4500 software (EKB Technologies, Ltd.). The timing sequence for the images was also set using this software. The first image had the smallest black square, appearing as a pin-point, and was shown as a ‘pre-stimulus’ for 80 ms. Following this, the remaining 23 images were shown in sequence lasting for one of two possible durations: either 20.4 ms (one image every 0.9 ms), or 33.6 ms (one image every 1.5 ms).

*Experimental protocol.* Individual houseflies were put on ice to temporarily stun them. Once the fly stopped moving, a small loop made from 0.1-diameter copper wire was fixed to the dorsal thorax with a thin layer of cyanoacrylate glue (**Supplementary Fig. 39a**, inset panel). Care was taken to avoid impacting the range of motion of the wings or legs, so that these could all move freely during the experiment. After a brief recovery period, during which the glue dried and the fly recovered from the ice, the fly was placed in the arena. The wire loop was placed over the hook, and pinched so that the orientation of the suspended fly (facing the projector) would be secured.

The projector and the camera were synchronised in such a way that when the camera was triggered to start recording, the projector would play the selected stimulus. Each fly was presented with both stimuli, with each stimulus presented to each fly at least three times. Between presentations of the stimuli, flies were given a ~2 min rest period, to return to their normal behaviour after responding.

**Antenna response**

*Experimental set-up.* Experiments were conducted using a Multisensory Apparatus (MUSA) system; described previously as the Goniometric high-speed deep pseudopupil (GHS-DPP) imaging system by Kemppainen *et al*. (2022). Briefly, the MUSA system (**Supplementary Fig. 39b**) utilises a sideways-mounted stereomicroscope (Olympus SZX12) with a high-intensity UV LED (365 nm UV-OptoLED, Cairn Research, Faversham, UK) in the ocular slot, which delivers the stimulus. A high-speed camera (Orca Flash 4.0 C13440, Hamamatsu Photonics, Shizuoka, Japan), capable of recording from 100-1000 fps, is also mounted to the microscope. To allow the camera to record at 1000 fps, we used a crop of 1024 x 24 (width, height), with 2x2 sensor binning. Transparent IR and opaque UV filters in the camera prevent the UV light from the stimulus from polluting the image sensor.

The MUSA set-up is enclosed in a black metal cubicle, with the one open side covered in black curtains to ensure conditions within remain dark. The set-up within is back-illuminated by two 850 nm IR LEDs (IR-OptoLED, Cairn Research, UK), to allow images to be obtained. A PC running Windows 7 and Gonio-imsoft v0.1.0 software were used to obtain the images. As previously described by Kemppainen et al.^3,4^, Gonio-imsoft interfaces with MicroManager to control the camera, using the NI-DAQmx module to control data acquisition. All software are available under a free and open (GPLv3) software license in a GitHub repository: [https://github.com/musaprog](https://github.com/musparog) .

*Stimulus design.* Each trial began with 200 ms perceptual darkness (with IR back-illumination only). This was followed immediately by the stimulus, which comprised a 40 ms-long UV flash. The LED driver control value (brightness) was 1 V for all but two flies (named alicetest3 and muscatest in the original data, where it was 2 V). IR illumination remained consistent during the flash, and the stimulus was followed by a further 200 ms of perceptual darkness with IR illumination only.

*Experimental protocol.* Individual houseflies were secured in a pipette tip that was trimmed to make the opening wide enough to allow their heads to emerge from the tip without damage, but narrow enough that their thorax could not pass through (**Supplementary Fig. 39b**, inset panel). Melted beeswax was then applied around the thorax to secure them to the pipette tip, which was mounted on the imaging stage of the MUSA system. For each fly, the stimulus was repeated either 10 or 100 times, with each trial separated by a 10-second interval.

**Video analysis**

The leg response videos had 625 frames, with each frame representing 0.4 ms; making each video 250 ms long. At frame 2, the pre-looming stimulus light appeared, and the looming stimuli began at frame 200. The looming stimuli ended at either frame 251 (for the 20.4 ms stimulus) or 284 (for the 33.6 ms stimulus). Meanwhile, the antenna response videos had 440 frames, with each frame representing 1.0 ms, meaning each video was 440 ms long. The light stimulus began at frame 201 and continued until frame 241: a duration of 40 ms. See **Supplementary Movie 3**.

Videos were analysed manually using ImageJ software, where they were viewed as .tif stacks (version 1.54g^144^). The analysis was done according to the same principle for both types of video. First, videos where the fly failed to move at all were excluded from the analysis. Next, videos where the fly moved *prior* to the presentation of the stimulus were removed: even if this movement was slight and more substantial movements occurred post-stimulus. This helped ensure that the only videos included in the analysis involved responses to the stimulus, specifically. Of the remaining videos, the first frame where either a leg or an antenna shifted position (if this occurred) was taken as the start of the response (with the latency being the difference between this frame and the first frame of the the stimulus, converted to ms).

To validate the manual analysis, we also performed automated tracking using a two-dimensional cross-correlation technique previously described by Kemppainen et al.^3,4^, where it was used to measure photoreceptor microsaccade kinematics. This method was applied to antennal motion using the Gonio-Analysis software (v0.8.2, available at <https://github.com/musaprog>). Rectangular regions of interest (ROIs) were manually drawn around the antennal arista, partially enclosing it. The ROIs were approximately square, with side lengths ranging from 10 to 20 pixels, and typically spanned most of the vertical extent of the image (24 pixels).

Motion analysis was performed using two-dimensional cross-correlation; however, the results were constrained to the x-dimension for two primary reasons. First, in the narrow 24-pixel-high images, the ROI frequently approached the horizontal image borders (or with a sharp oblique from the horizontal), occasionally touching them. Handling such out-of-bounds cases would have required additional algorithmic complexity with minimal gain in useful information. Second, the arista was typically oriented vertically and appeared as a line-like structure in the images. The effectiveness of cross-correlation - its ability to track features while rejecting noise and spurious matches - depends on the structural characteristics of the target. For line-like features, correlation is weak along the axis of the line (predominantly the y-direction), but strong perpendicular to it (x-direction). Thus, restricting analysis to the x-dimension improved tracking reliability.

Following motion analysis, reaction times were extracted from the time-motion traces using a visual inspection protocol. In low-noise cases with abrupt motion onset, reaction time was directly read from the time axis at the point of onset. In noisier traces or when the onset was more gradual, reaction time was estimated as the intersection of two fitted lines: a baseline fit to the pre-stimulus period and a linear fit to the initial phase of motion.

Overall, the manual and automated analyses yielded comparable reaction times. For example, in *fly001*, the manual method produced a mean reaction time of 25.8 ms (median: 26 ms), while the automated method gave a mean of 26.4 ms (median: 25 ms). The manual approach successfully estimated reaction times in 79 out of 100 trials, compared to 59 out of 100 for the automated method. Although we did not systematically investigate this difference any further, the reduced success rate of the automated analysis was possibly due to instances where the arista intermittently exited the field of view or the ROI - a situation the manual analysis could accommodate but the automated algorithm could not.

Two-dimensional cross-correlation was initially applied to the leg response data. However, it failed to produce consistent reaction time estimates in most cases, due to several confounding factors: (1) the complex and variable trajectories of leg movements, (2) frequent leg-leg occlusions and body-induced masking, (3) image noise stemming from high frame rates and limited illumination, (4) insufficient spatial resolution, and (5) the use of a pseudorandom background, which interfered with correlation tracking by partially engaging the algorithm's grip.

To address these limitations, we developed a custom linear interpolation-based motion analysis, implemented within the Movemeter software (v0.8.0, available at <https://github.com/musaprog>). In this approach, the user marks line segments along the legs in selected frames, and the software interpolates their positions across the remaining frames. By manually inspecting the video across frames, the user can visually identify time points where the interpolated line diverges from actual leg motion and correct it by adding new line segments. This process results in an interpolated path that closely and reliably tracks leg motion over time. An inherent limitation - though potentially an advantage depending on the research question - of the linear interpolation approach is its inability to capture the fine-scale motion fluctuations present in the cross-correlation analysis. Some of these fluctuations may reflect true high-frequency motion dynamics, while others may arise from image noise sources such as photon shot noise or sensor-related artefacts.

**Statistical analyses**

To quantify the fly’s fastest responses robustly, we summarised the lower tail of the response-time distribution using the 10^th^ percentile (q10; fastest 10% of trials). Uncertainty was estimated via a non-parametric bootstrap (50,000 resamples with replacement), yielding a bootstrap distribution for q10. We then defined a critical fast-latency threshold T* as the smallest response-time bound for which the data support, at α = 0.05, that q10 lies below it. Operationally, T* was taken as the 95^th^ percentile of the bootstrap q10 distribution (equivalently, the smallest T such that Pboot(q10 ≥ T) ≤ 0.05. For interpretability, T* was reported rounded up to the nearest millisecond.

For rapid six-leg lift responses (**Supplementary Fig. 40**), reaction times were variable and right-skewed (n = 35). The fastest-capable performance, quantified by the 10th percentile, was q10 = 17.76 ms (bootstrap 95% CI [16.80, 23.20] ms). From the bootstrap distribution of q10, we estimated the critical fast-latency threshold T* (α = 0.05) as ~21.9 ms (one-sided bootstrap test, p = 0.026), indicating that the fastest 10% of responses occur below ~22 ms.

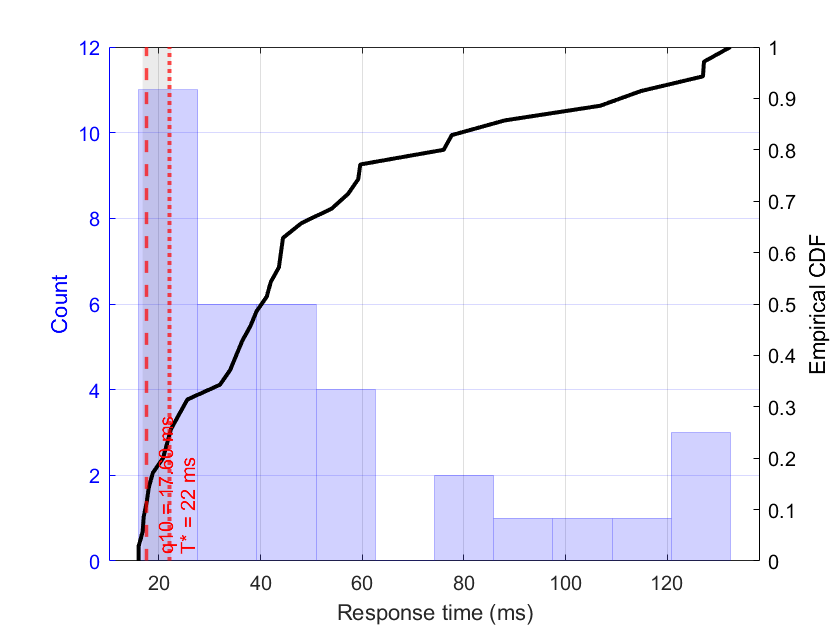

**Supplementary Fig. 40**. **Distribution of fly response times to looming stimuli and fastest-tail estimate.** Blue bars (left y-axis) show the histogram of raw response times (counts). The black curve (right y-axis) is the empirical cumulative distribution function (ECDF). The red-dashed vertical line marks the 10th percentile latency (q10; the fastest 10% of trials). The grey shaded band indicates the 95% bootstrap confidence interval for q10 (50,000 resamples). The black dotted vertical line marks the critical fast-latency threshold T* (reported as 22 ms), defined as the smallest upper bound for which the data support at α = 0.05 that q10 lies below T*.

Antenna movement response times (**Supplementary Fig. 41**) were variable and right-skewed (n = 35). The fastest-capable performance, quantified by the 10th percentile, was q10 = 22.00 ms with a bootstrap 95% CI [19.00, 23.00] ms (n=79). Using the bootstrap distribution of q10, we estimated the critical fast-latency threshold T* (α = 0.05) as approximately 23.00 ms (one-sided bootstrap test, p = 0.025). At this threshold, the one-sided bootstrap tail probability satisfied p ≤ 0.05, indicating that the data support the statement that the fastest 10% of responses occur below ~23 ms.

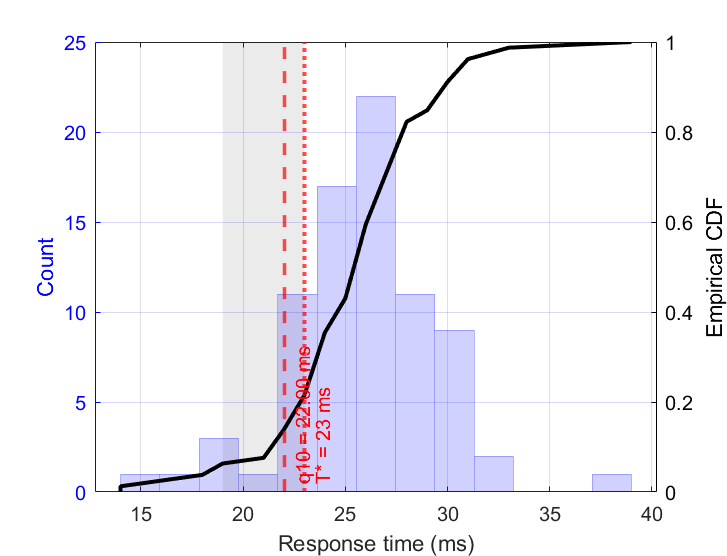

**Supplementary Fig. 41.** Leg movement analysis. Response times were variable and right-skewed (n = 35). The fastest-capable performance, quantified by the 10th percentile, was q10 = 17.76 ms with a bootstrap 95% CI [16.80, 23.20] ms. Using the bootstrap distribution of q10, we estimated the critical fast-latency threshold T* (α = 0.05) as approximately 21.9 ms (one-sided bootstrap test, p = 0.026). At this threshold, the one-sided bootstrap tail probability satisfied p ≤ 0.05, indicating that the data support the statement that the fastest 10% of responses occur below ~22 ms.

**VI Functional Connectomics**

### VI.1: Workflow for exploring the functional logic of *Drosophila* circuits in the Fruit Fly Brain Observatory

The Fruit Fly Brain Observatory (FFBO) is an open-source ecosystem of tools to explore experimental data and computational models of the fruit fly brain^5^. Within the FFBO ecosystem, FlyBrainLab (FBL) is an interactive computational platform to programmatically interact with FFBO tools^158^. As depicted in **Supplementary Fig. 42**, FlyBrainLab enables a principled workflow to (i) visually explore the 3D morphology of neuropils, neurons and synapses in the connectome using natural language queries via the [NeuroNLP](https://flywire.neuronlp.fruitflybrain.org/) user interface (<https://www.fruitflybrain.org/#/brainmapsviz>) (ii) construct and subsequently analyze graph abstractions of morphological connectivity data using the NeuroGraphBench tool (iii) use the NeuroCircuitDesk tool to create executable circuits from the abstracted graphs by assigning executable code to each component of the graph (iv) use the NeuroGFX tool to interactively explore the functional logic of executable circuits for various inputs.

While the workflow described here is implemented using the *Drosophila melanogaster* connectome, it is used in this study as a reference framework for identifying minimal feedforward architectures and conservative lower-bound transmission constraints, rather than as a claim of numerical or anatomical equivalence across dipteran species.

**Supplementary Fig. 42.** FlyBrainLab workflow from structure to function. Columns: (far left) 3D exploration and visualization of fly brain connectomic/synaptomic datasets (with NeuroNLP), (left) creation of graph abstractions and analysis (with NeuroGraphBench), (right) creation of executable circuits (with NeuroCircuitDesk), (far right) interactive exploration of the functional logic of the chosen circuits (with NeuroGFX).
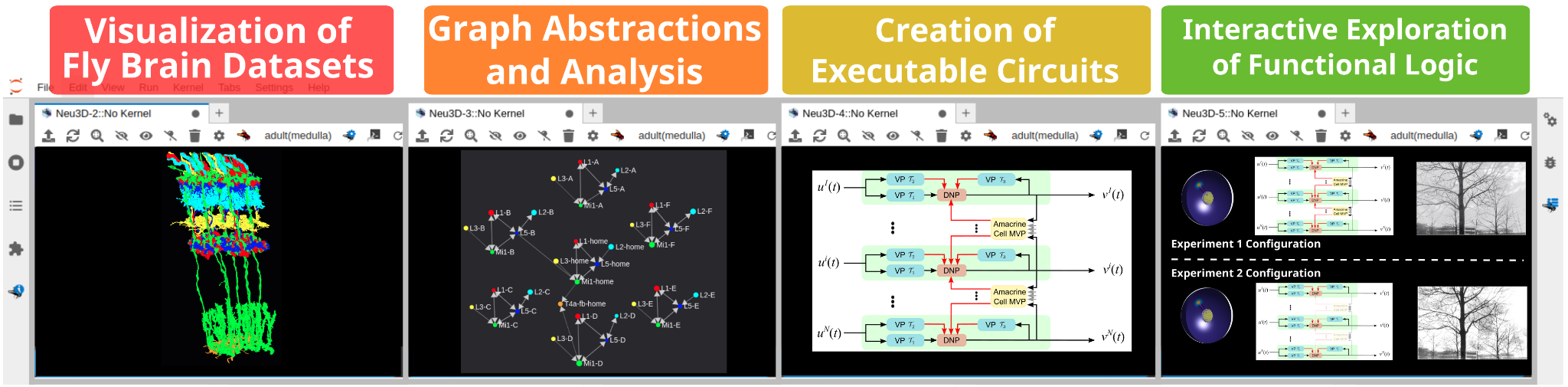

#### An example of using the NeuroGraphBench tool to construct graph abstractions of the fruit fly connectome

In this section, we discuss a graph abstraction of the connectome that enables quick discovery of reflexive feedforward pathways. A naive graph abstraction of the connectome entails constructing a directed graph $G_{NS}$ with *neurons as nodes* and the *synapses that connect them as edges*. However, given that the Drosophila brain has over 200,000 neurons searching for pathways over $G_{NS}$ is very computationally expensive.

Instead, if we restrict ourselves to reflexive feedforward pathways that do not traverse local neurons (that are not strictly feedforward and often serve as feedback neurons), we can instead construct a coarse-grained level directed graph $G_{NT}$ with *neuropils as nodes* and *tracts of neurons as edges*. Each neuropil node $R$ in $G_{NT}$ comprises the set of neurons $N_{R}$ that innervate it, while the tract from a neuropil $R_{A}$ to another neuropil $R_{B}$ comprises the set of neurons with input synapses in $R_{A}$ and output synapses in $R_{B}$. More formally, let $N_{i}(R), N_{o}(R)\subset N_{R}$ respectively be the set of neurons with input and output synapses in neuropil $R$. Then the tract from neuropil $R_{A}$ to neuropil $R_{B}$ is defined as $T(R_{A},R_{B}) = N_{i}(R_{A}) \cap N_{o}(R_{B})$. Furthermore, let $S(N_{1}, N_{2}, R)$ denote the subset of neurons $N_{1}$ that have synaptic outputs onto the set of neurons $N_{2}$ in neuropil $R$. The set of valid pathways over $G_{NT}$ with $N$ synaptic hops from a source neuropil $R_{S}$ to a set of target neurons $N_{T}$ innervating the target neuropil $R_{T}$ are then given as: $S(N_{S}, T_{1}, R_{S}) \to S(T_{1}, T_{2}, R_{1})\to S(T_{1}, T_{2}, R_{1})\to\cdots S(T_{N-1}, N_{T}, R_{T})$, where $R_{1}, R_{2}\cdots R_{N}$ and $T_{1}, T_{2},\cdots T_{N}$ are respectively the intermediate neuropils and tracts along the N-hop pathways.

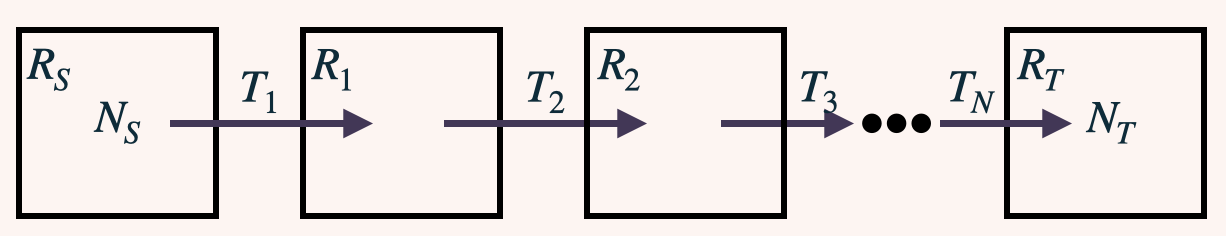

**Supplementary Fig. 43**. A diagram illustrating a valid pathway from source neurons $N_{T}$ in neuropil $N_{S}$ to target neurons $N_{T}$ in neuropil $R_{T}$.

### VI.2: Using NeuroGraphBench to estimate response times for visuomotor pathways

#### This analysis is explicitly restricted to identifying the shortest feedforward visuomotor pathways, thereby providing conservative lower-bound estimates on pathway depth and transmission delay that are largely insensitive to species-specific differences in overall brain size, neuron number, or neuropil volume.

#### VI.2.1: Finding and visualizing pathways in Drosophila

In subsequent analysis, we wish to find the shortest feedforward pathways from the retina to some motoneurons of interest. As discussed in the previous section, pathfinding on the naive graph abstraction $G_{NS}$ is computationally prohibitive, and we thus instead use NeuroGraphBench to look for shortest paths along $G_{NT}$ instead. Given a source neuropil $R_{S}$ (with the set of neurons $N_{S}$ innervating it) and a set of target neurons $N_{T}$ (that innervate a target neuropil $R_{T}$), we can use Dijkstra’s shortest path algorithm over $G_{NT}$ by setting the source and target nodes to respectively be $R_{S}$ and $R_{T}$ and then filtering for pathways $P$ that satisfy $N_{T} \subset P$. We further filter the neurons by neurotransmitter type, only retaining pathways comprising fast excitatory acetylcholinergic neurons (**Supplementary Fig. 43**).

##### VI.2.1.i Visuomotor Pathway to Antennae

Unlike the giant-fibre-mediated escape pathway, the visuomotor circuitry controlling antennal movements has not yet been anatomically reconstructed in *Musca* or other dipteran species. Accordingly, the pathways identified here should be interpreted as hypothesis-generating estimates rather than definitive circuit reconstructions.

The antennae of *Drosophila* are controlled by 4 muscles inserted at the interior portion of the second segment^159^. Some of the motoneurons that synapse onto these muscles have been identified through GAL4 lines^159^ and subsequently labeled on FlyWire dataset^160^ by community effort. **Supplementary Fig. 44** shows NeuroNLP visualization of the 5 antennal motoneurons on the FlyWire dataset (CB0723 cyan, CB0750 yellow, CB0810 green, CB0873 blue and CB0886 red), whose axons are all bundled in one track together with a retinal motoneuron (CB0804 black) that innervate the muscles controlling retinal movement^38^.

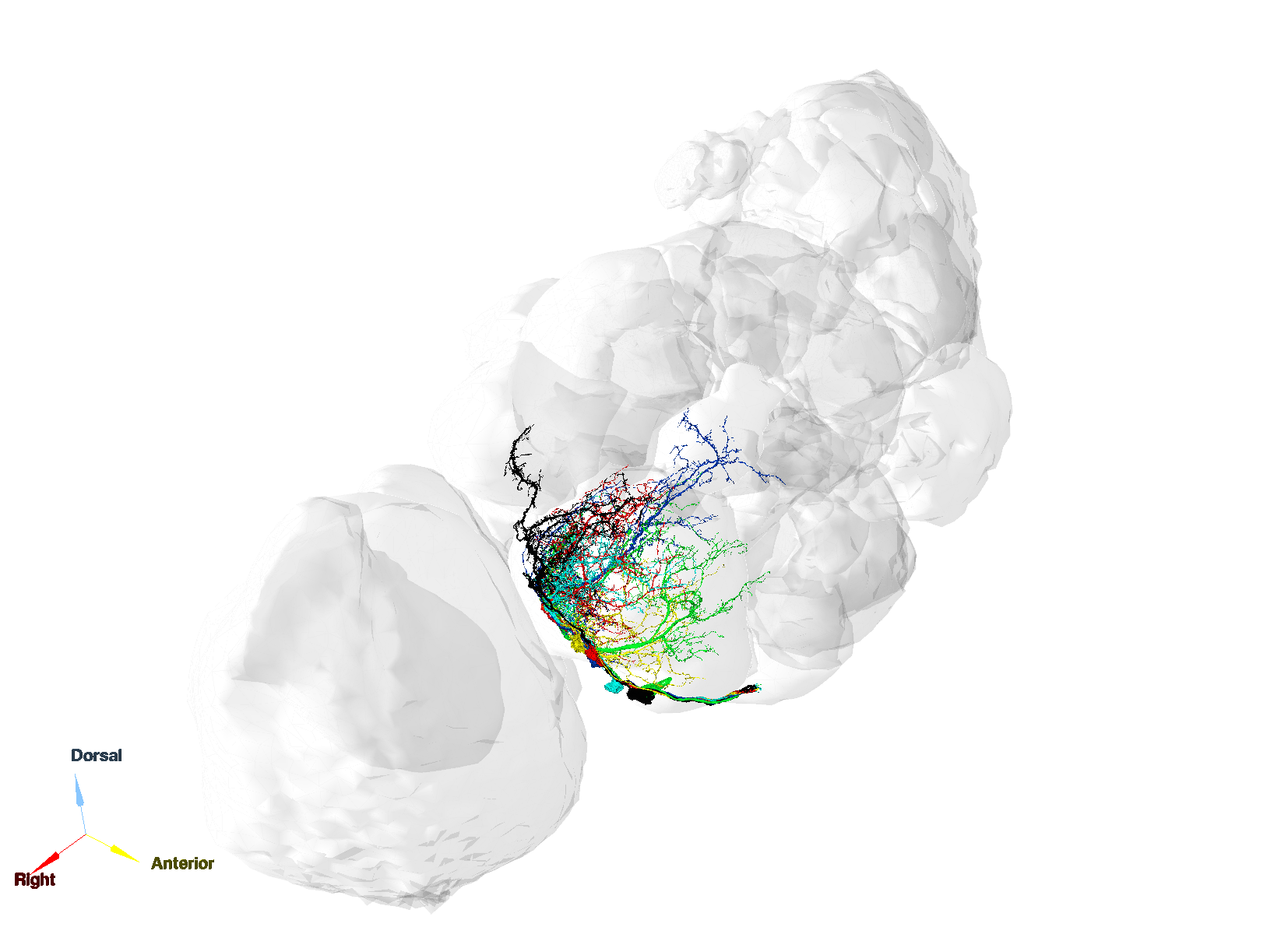

**Supplementary Fig. 44**. Antenna motoneurons ([tag](https://flywire.neuronlp.fruitflybrain.org/?tag=yiyin_20250709_ant_motoneurons))

Per the formalism described above, to find pathways from the retina to the antennae, we set the photoreceptors R1-R6 (in the retina, $R_{S})$ as source neuron $N_{S}$, and the 5 antennal motoneurons of interest (in the Antennal Mechanosensory and Motor Center, $R_{T}$) as target neurons $N_{T}$. The corresponding shortest paths over $G_{NT}$ obtained from NeuroGraphBench, correspond to multiple directed paths of neurons in $G_{NS}$ with the least number of (synaptic) hops that receive synaptic input in the retina and provide synaptic output to the 5 antennal motoneurons.

One such shortest pathway identified from the retina to antenna muscles is shown in **Figure 4**. This circuit begins with photoreceptors R1-R6, which responds to light input and drives the lamina monopolar cell L1; L1, in turn, projects to the transmedullar neuron Tm3, which arborizes onto LC4, a visual projection neuron that is involved in looming escape response; LC4 then synapse onto the descending neuron DNp05, which drives the antenna motoneuron CB0810 in the GNG that directly controls the antenna muscle (see also **Supplementary Movie 5**).

##### VI.2.1.ii Visuomotor Pathway to Legs

For leg-lift response described in **Supplementary Fig. 46e-g**, we consider well-known cylindrical tergotrochanteral muscles (TTMs) that are innervated by the tergotrochanteral motoneurons (TTMns)^161,162^. The giant fibers (GFs) that mediate escape response in *Drosophila* form chemical and electrical synapses with the TTMns^162^. The TTMs have been observed to elicit fast esponse from brains stimulation in *Musca*^161,163^. To identify the shortest path to this TTMn motoneuron via the GF, we again used NeuroGraphBench to estimate the shortest pathway from the retina to the GF in the FlyWire dataset, and from the GF to the TTMns in the VNC dataset. We thus demonstrate finding shortest pathways by combining information from two complementing connectomic datasets.

One of the shortest pathways identified from the retina to leg motor control is shown in **Supplementary Fig. 44** and **Supplementary Fig. 46**, reconstructed from the FlyWire (brain) and MANC (VNC) datasets, respectively; together they reconstruct a looming-induced escape circuit by combining data from the brain and the ventral nerve cord. This circuit begins with photoreceptors R1-R6, which responds to light input and drives the lamina monopolar cell L3; L3, in turn, projects to the transmedullar neuron Tm20, which arborizes onto LPLC2, another visual projection neuron that is involved in looming escape response; LPLC2 then synapse onto the descending neuron GF, which directly drives the motoneuron MNwm34_T2 in the ventral nerve cord (VNC) that directly controls the leg muscle (see also **Supplementary Movie 3**).

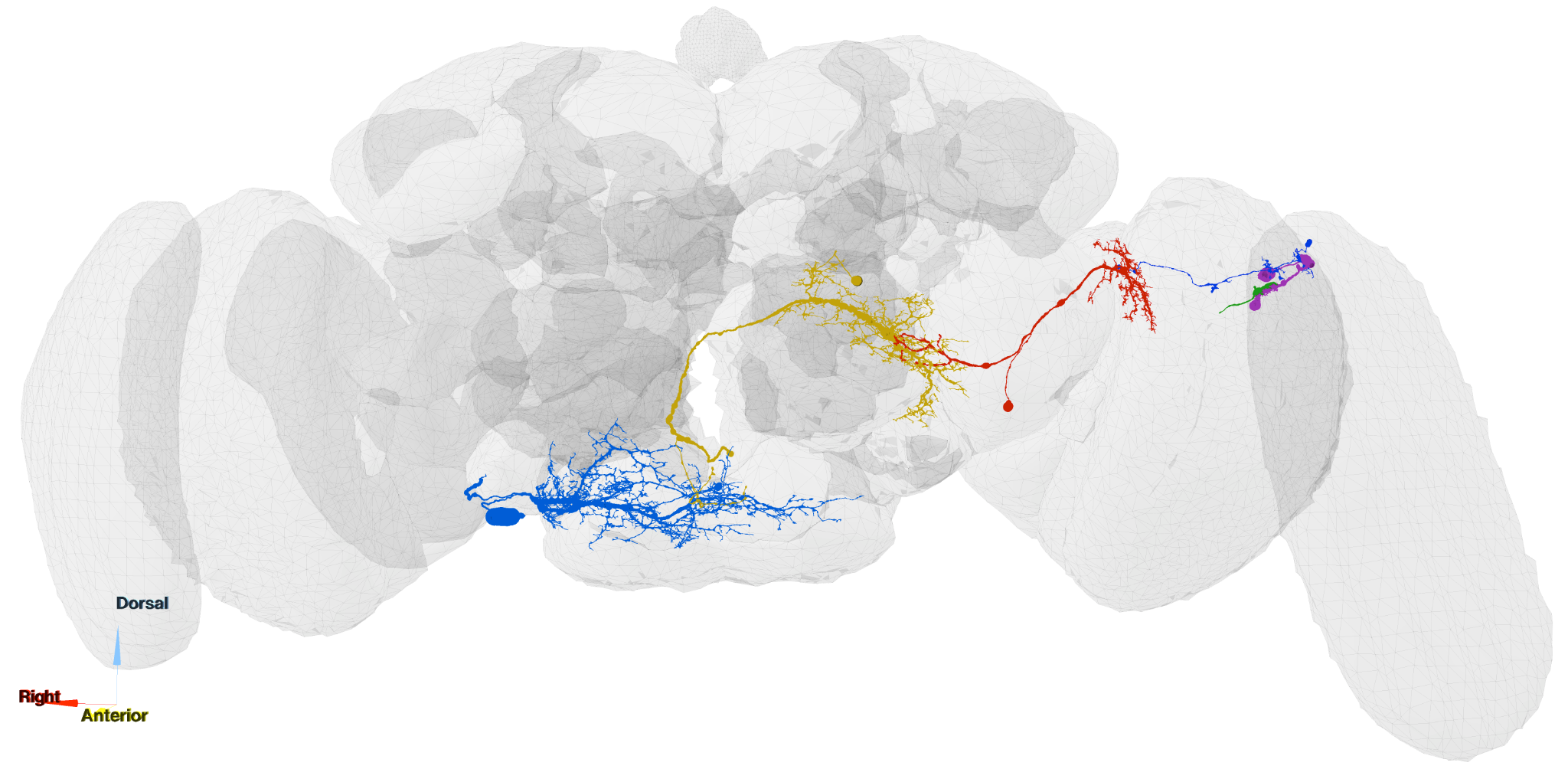

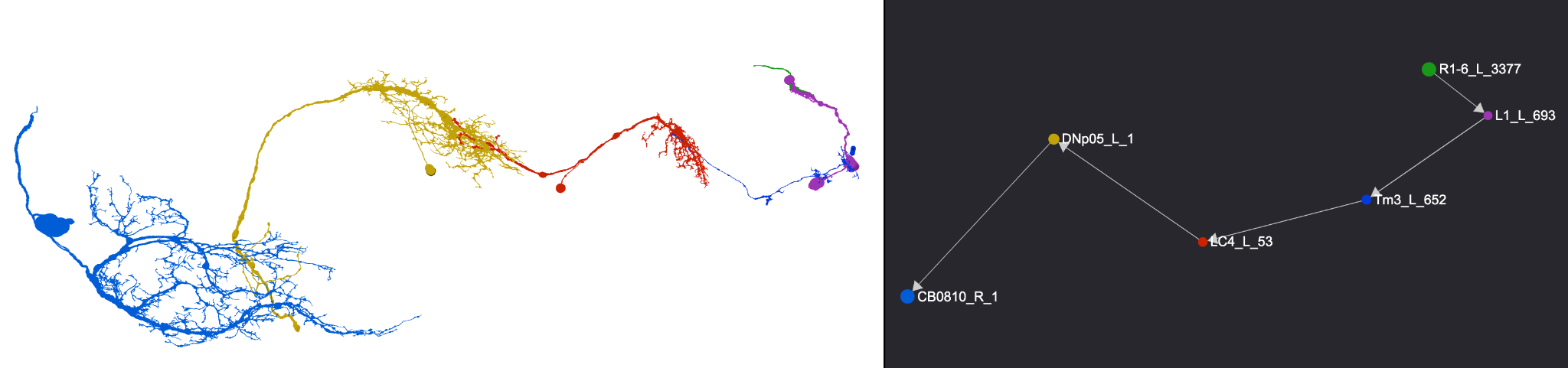

**Supplementary Fig. 45.** Visualisation of a shortest path circuit for light-induced antenna movement response, from the retina to the gnathal ganglia (GNG) in Flywire dataset using (<https://flywire.neuronlp.fruitflybrain.org/?tag=supplementary_retina_antenna>). *Top*: Full anatomical view of the circuit overlaid on the brain neuropils. *Bottom left*: Isolated view of the neurons involved in the pathway. *Bottom right*: Graph representation of the circuit, showing connectivity and directionality between nodes.

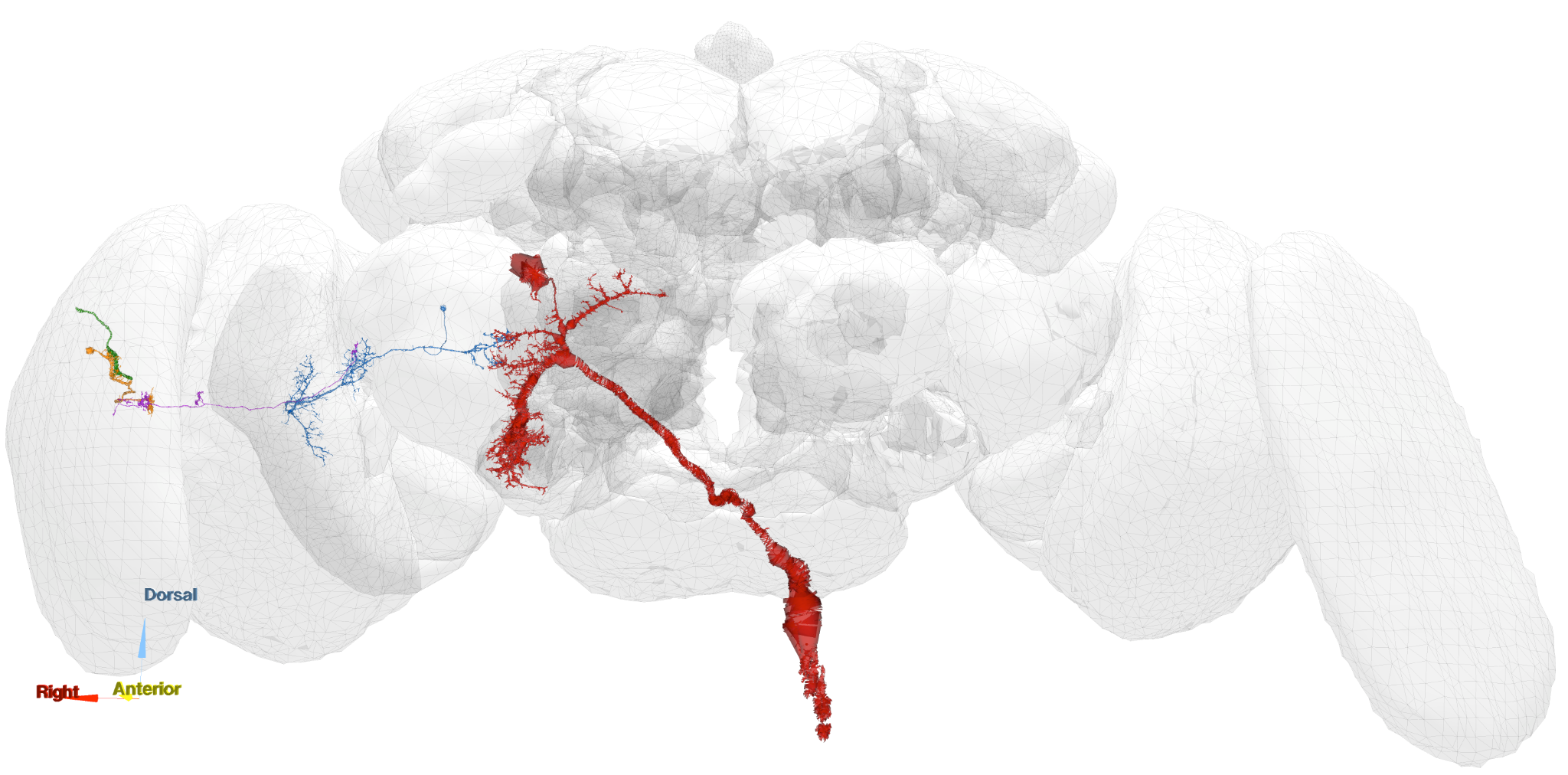

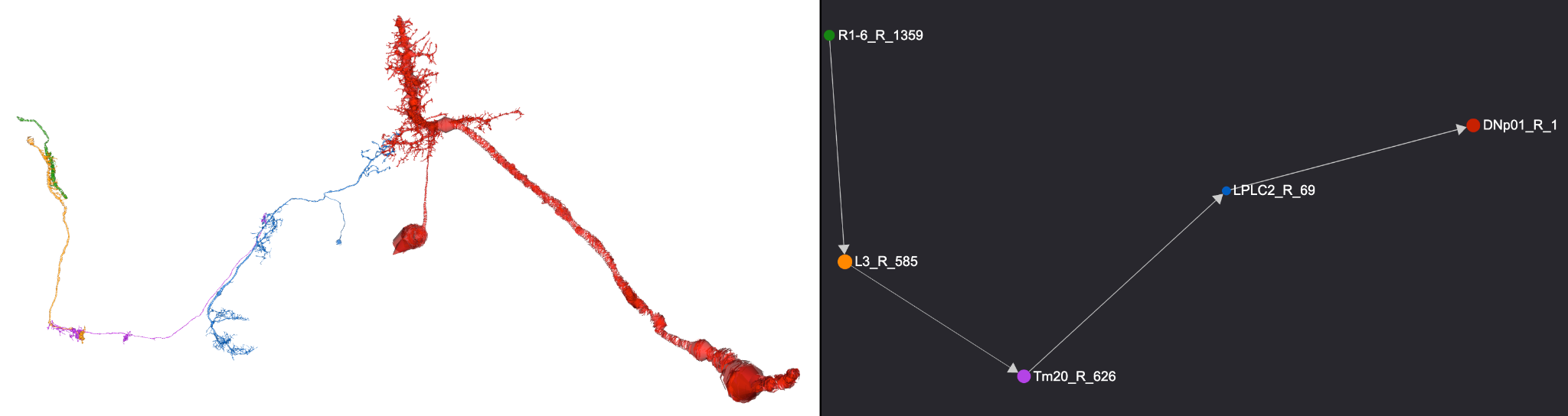

**Supplementary Fig. 46.** Visualisation of the brain segment of a shortest path circuit mediating looming-induced escape, from the retina to the GF neuron (DNp01 in Flywire convention), reconstructed from the FlyWire dataset using [NeuroNLP](https://flywire.neuronlp.fruitflybrain.org/?tag=supplementary_retina_leg) (<https://flywire.neuronlp.fruitflybrain.org/?tag=supplementary_retina_leg>). *Top*: Full anatomical view of the circuit overlaid on the brain. *Bottom left*: Isolated view of the neurons involved in the pathway. *Bottom right*: Graph representation of the circuit, showing connectivity and directionality between nodes.

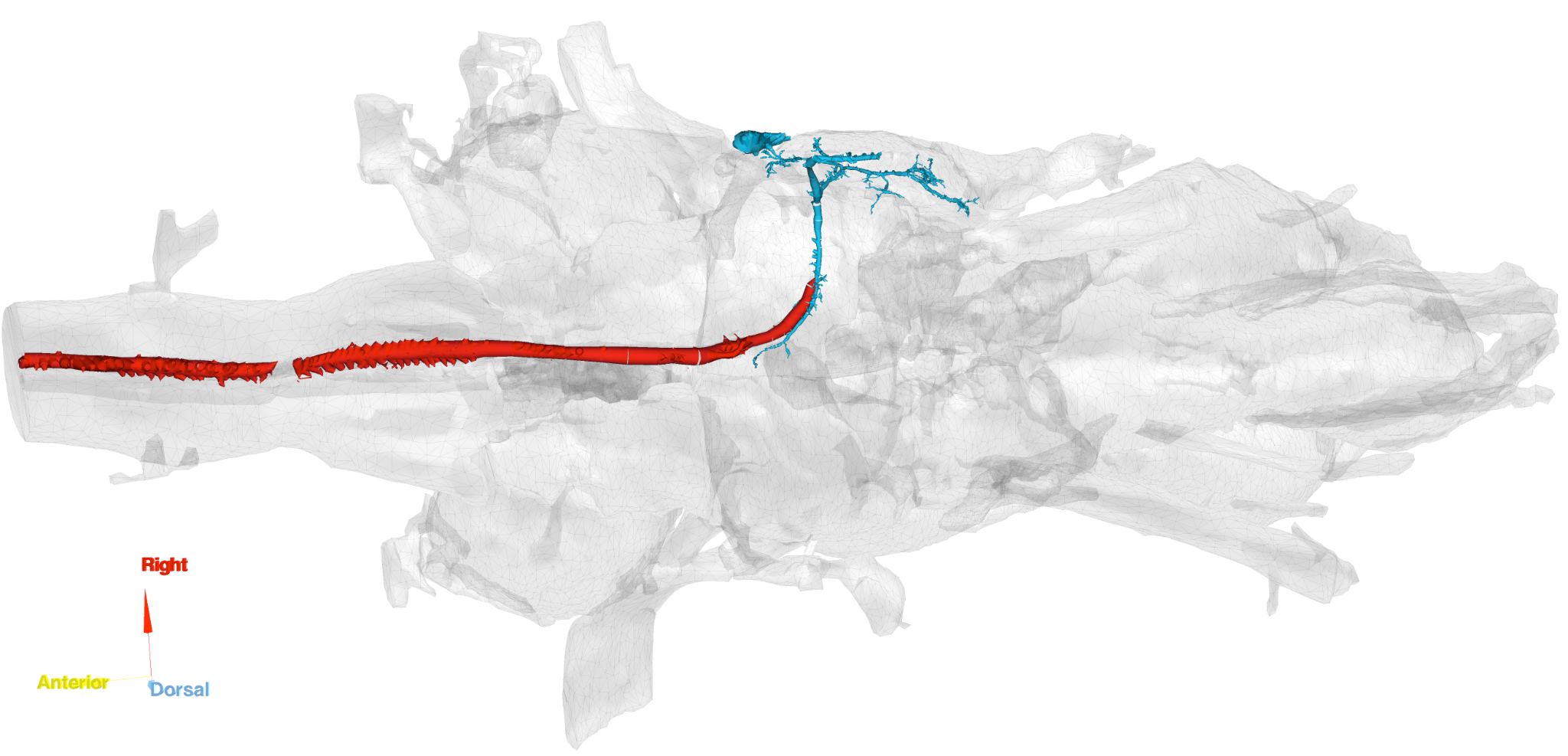

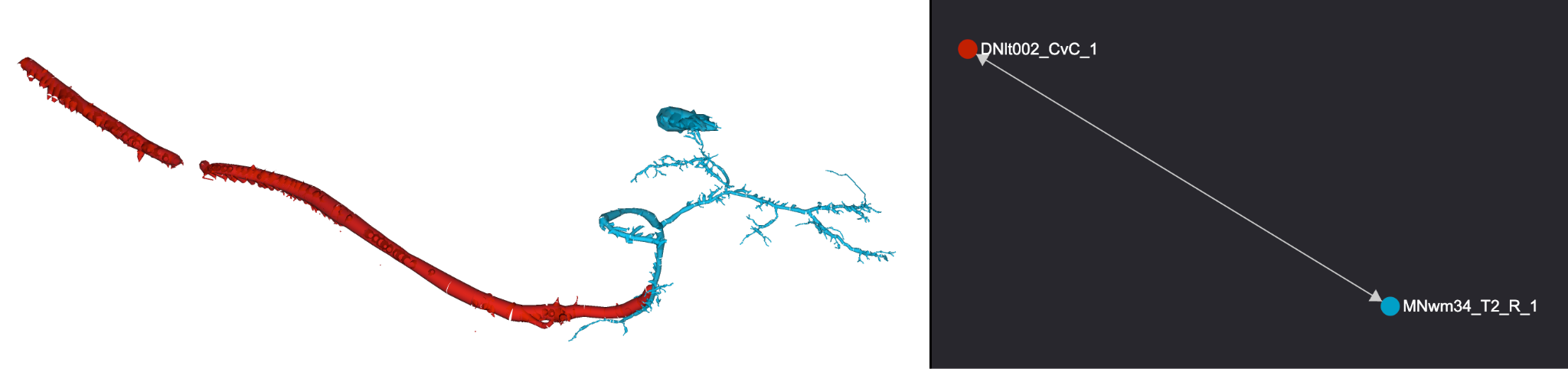

**Supplementary Fig. 47.** [NeuroNLP](https://flywire.neuronlp.fruitflybrain.org/?tag=supplementary_retina_leg) visualisation of the ventral nerve cord (VNC) segment of the looming-induced escape circuit, from the GF neuron (DNlt002 in MANC convention) to a leg motoneuron, reconstructed from the MANC dataset (<https://flywire.neuronlp.fruitflybrain.org/?tag=supplementary_retina_leg>). *Top*: Anatomical view of the pathway within the VNC. *Bottom left*: Neurons isolated from surrounding structures. *Bottom right*: Graph representation showing the downstream connectivity from the giant fiber to motor output.

#### VI.2.ii: Estimating time delays along visuomotor pathways in *Drosophila* and *Musca*

Having found the shortest pathways using NeuroGraphBench in **Section VI.2**, we also estimate an upper bound for the conduction lengths along these pathways. We do this by finding the cable length of the longest path along each neuron in a pathway, and then sum up these cable lengths for each neuron in the pathway to obtain an upper bound for the conduction length along the pathway. These conductance length estimates will be used in subsequent conductance time calculations, assuming a uniform conduction speed along the pathway.

##### VI.2.ii.a: *Drosophila*

To estimate the minimal reaction time of *Drosophila*’s antenna and leg movement in response to visual stimuli, we identified the shortest reflex circuits linking the retina to the relevant motoneurons (as shown in **Supplementary Fig. 45-46**). These fast sensorimotor pathways can be broken down into processing stages each associated with specific delays:

1. **Phototransduction and Voltage integration**:

Photoreceptors initially convert a flash of light into a graded voltage response. This transformation is governed primarily by the phototransduction cascade and the subsequent integration of the signal, and typically occurs over approximately 10 ms (see **Figure 1** in the main text).

2. **Neural Conduction and Synaptic Delays**:

Signal propagation speed has been shown to be approximately 0.13 m/s in peripheral sensory neuron in *Drosophila* larva^164^ which is thinner than the housefly sensory neurons and can be served as a lower bound of condution speed. The higher bound can be established by the conduction speed in *Drosophila* giant fiber that is approximately 2 m/s^165^.

In *Drosophila*, neurons are very small with short axonal lengths, resulting in minimal conduction delays, typically in the sub-millisecond range (approximately 0.1-0.5 ms per segment). Even when accounting for several such segments, these delays remain relatively low.

3. **Synaptic Transmission & Membrane Charging**:

Each synapse introduces a delay due to the time required for neurotransmitter release, receptor activation, and the subsequent charging of the postsynaptic membrane toward threshold. It has been observed that the synaptic transmission delay from an excitatory projection neuron to a local neuron in the antennal lobe is about 1 ms^166^. The latency from a Giant Fiber spike to TTM muscle activation had been measured to be approximately 0.81 ms^167^. The time required for membrane charging, determined by the cell’s membrane time constant is effectively included within the synaptic delay estimates.

With these delay estimates established, we can derive a lower-bound approximation of the total neural processing time for fast visual reflexes in *Drosophila*. By summing the component delays along the shortest identified pathways from the retina to motoneurons, we estimate the minimal time required to initiate motor responses in two representative circuits: one controlling the antenna and the other the legs. Even when using conservative estimates, the total sensorimotor delay consistently falls within a narrow range that serves as a robust approximation of the lower-bound reaction time

**Supplementary Table 17. Light-induced Antenna Reflex Pathway in *Drosophila***

| **Stage** | **Description** | **Estimated Delay** |
| --- | --- | --- |
| Phototransduction | Phototransduction by R1-R6 photoreceptors | ~15 ms |
| Synaptic Transmission | 6 synapses, as shown in graph edges in **Supplementary Fig. 45**,  at ~1-2 ms each | 6 ms |
| Conduction Delay | Across short axonal segments (~2,100 µm) | >1.76-4.4 ms |
| Total Estimated Delay |  | 22.76-25.4 ms |

**Supplementary Table 18. Looming-Induced Leg Reflex Pathway**

| **Stage** | **Description** | **Estimated Delay** |
| --- | --- | --- |
| Phototransduction | Phototransduction by R1-R6 photoreceptors | ~15 ms |
| Synaptic Transmission | 4 synapses, as shown in graph edges in **Figure 3** and **4**, at ~1-2 ms each | ~4 ms |
| Conduction Delay (up to the GF) | Across short axonal segments (360 µm) with a conduction speed of 0.2-0.5 m/s | 0.72-1.8ms |
| Transmission from GF to TTM motor activation | Measured to be 0.81 ms | 0.81 ms |
| Total Estimated Delay |  | 20.53-21.61ms |

Accounting for possible additional minor overheads in neural processing, realistic estimates range from 18 to 21 ms for the antenna light reflex and 16-17 ms the full looming-triggered avoidance response. These estimates are based on highly optimised and compact neural circuits and assume the shortest unidirectional, reflex-like pathways relying on conventional fast synaptic transmission.

##### VI.2.ii.b: Adapting *Drosophila* visuomotor pathways to *Musca*

As depicted in **Supplementary Fig. 46**, Buschbeck and Straussfield^168^ experimentally verified the existence of an evolutionarily conserved Diptera pathway from the retina to leg motoneurons that is functionally contiguous despite taxon-specific variations.

Although *Musca domestica* has a substantially larger brain and higher absolute neuron count than *Drosophila* melanogaster, comparative anatomical and physiological studies indicate that early visual processing stages and fast visuomotor escape pathways in dipterans follow a conserved organisational plan, with differences primarily reflecting scaling rather than fundamental changes in circuit depth or logic.

Given this evolutionary correspondence, we postulate the existence of a shortest pathway from the retina to the giant fibre in *Musca* that is comparable to the one we found in *Drosophila*, as described in **Section VI.2.ii.a**.

The experimentally verified homologue of the fruit fly GF in the housefly is the Giant Descending Neuron (GDN)^169^, which also initiates escape behaviours in response to looming stimuli. Behaviorally, houseflies and fruit flies exhibit qualitatively similar escape behaviours with rapid leg-driven launch and take-off to avoid a collision. *Musca* escapes are very fast and high-thrust^170^, befitting a larger fly that can cover more distance in a jump. The housefly’s threshold-trigger mechanism emphasises speed and robustness: once the looming object’s shadow reaches a critical intensity, the fly commits to an escape with minimal delay. *Drosophila* escapes, while only marginally slower, exhibit more steering and planning. Fruit flies devote ~200 ms to aim their jump, as evidenced by the preflight postural adjustments and oriented take-off direction. This difference may partly reflect body size and ecology: a 2 mm fruit fly might benefit from ensuring a correct escape direction (since it cannot leap far), whereas a 6-7 mm housefly’s more powerful jump might carry it out of harm’s immediate reach regardless of direction. Nevertheless, it is likely that houseflies also perform some threat localisation - observations of flies jumping away from a looming stimulus suggest they, too, can aim their escape. Despite these differences, both insects achieve reaction times on the order of a few hundred milliseconds or less from threat detection to take-off.

On the other hand, as past literature on both *Drosophila* and *Musca* does not discuss reflexive pathways from the retina to antennae, the shortest pathway discussed in **Section VI.2.ii.b** is a novel pathway that we discovered from connectomic analysis, warranting further experimental investigation.

To adapt the estimate from *Drosophila* to *Musca*, we modify two measures. One is that we triple the conduction length as a housefly is approximately 3 times larger than a fruit fly. The other is that the latency from GF activation to TTM activation is measured to be 2.2 ms^161^.

These adaptations are intentionally conservative and are expected to overestimate, rather than underestimate, total transmission delay, ensuring that the resulting values represent upper bounds for reflex-like visuomotor responses in *Musca*.

The following tables summarise the delay estimation for the two pathways in *Musca*.

**Supplementary Table 19. Light-induced Antenna Reflex Pathway in *Musca***

| **Stage** | **Description** | **Estimated Delay** |
| --- | --- | --- |
| Phototransduction | Phototransduction by R1-R6 photoreceptors | ~10 ms |
| Synaptic Transmission | 6 synapses, as shown in graph edges in **Supplementary Fig. 45**,  at ~1 ms each | ~ 6 ms |
| Conduction Delay | Across short axonal segments   >2,640 µm) with a conduction speed of 0.2-0.5 m/s | >5.28-13.2 ms |
| Total Estimated Delay |  | ~21.28-29.2 ms |

**Supplementary Table 20. Looming-induced Leg Reflex Pathway in *Musca***

| **Stage** | **Description** | **Estimated Delay** |
| --- | --- | --- |
| Phototransduction | Phototransduction by R1-R6 photoreceptors | ~10 ms |
| Synaptic Transmission | 4 synapses, as shown in graph edges in **Supplementary Fig. 46**,  at ~1 ms each | ~4  ms |
| Conduction Delay (up to the GF) | Across short axonal segments (~1,080 µm) with a conduction speed of 0.2-0.5 m/s | ~2.16-5.4 ms |
| Transmission from GF to TTM motor activation | Measured to be 2.2 ms^161^ | 2.2 ms |
| Total Estimated Delay |  | ~ 18.36 - 21.6 ms |

Accounting for possible additional minor overheads in neural processing, realistic estimates range from 22.5 to 29.5 ms for the antenna light reflex and from 18.6 to 22 ms for the full looming-triggered avoidance responses.

We stress that these estimates are not intended to assert precise numerical equivalence between *Drosophila* and *Musca* circuits, but to demonstrate that the experimentally observed behavioural latencies in houseflies are compatible, at the order-of-magnitude level, with known dipteran visuomotor architectures operating under conventional synaptic and conduction constraints.

**Glossary**

**This glossary defines key technical terms used throughout the supplementary text, with cross-references to Supplementary Fig. and Supplementary Tables where they are exemplified.**

**Adaptive optics**

Real-time photomechanical changes of ommatidial optics to enhance resolution during high-speed imaging of photoreceptor microsaccades and synaptic morphodynamics (**Supplement Section IV**; **Supplementary Fig. 25-26**).

**Antialiasing hypothesis**

Proposes that small structural differences among photoreceptors that combat aliasing artefacts and improve reliability of visual information processing (**Supplementary Fig. 2-3**; **Supplementary Tables 1-2**).

**Biphasic response**

Voltage waveform with two phases (hyperpolarisation then depolarisation, or vice versa), characteristic of LMC responses to contrast transients (**Supplementary Fig. 5b**; **Supplementary Fig. 7a**).

**Bursty stimulus / saccadic stimulus**

High-contrast, temporally clustered light input mimicking rapid light intensity time series generated by saccadic movements, used to probe high-frequency encoding (**Supplementary Fig. 2**, **4**, **5**, **7**).

**Centre-surround antagonism**

Classical receptive-field model where central excitation is opposed by peripheral inhibition; contrasts with dynamic predictive coding observed in LMCs (**Supplementary Fig. 1**).

**Compound eye**

Insect visual organ composed of ommatidia; provides high temporal and spatial resolution (**Supplement Section II**; **Supplementary Fig. 13-15**).

**Contrast normalisation (dynamic)**

Synaptic adjustment of gain to maintain encoding efficiency across contrast levels; observed in LMC responses (**Supplementary Fig. 4f**, **7f**, **10**).

**Data processing theorem**

Information-theoretic principle that information cannot increase after processing; applied to compare photoreceptor and LMC information throughput (**Supplementary Fig. 4**, **7**).

**Diffraction limit**

The fundamental limit to the resolution of an optical system, determined by the wave nature of light and the aperture size of the imaging system (e.g., ommatidial lens). It sets the smallest angular separation between two objects that can be resolved without interference. In insect vision, this corresponds to the airy‑disk angle of individual facets (e.g., 1.1° in *Musca domestica*; see **Supplement Section II.8**, **Supplementary Fig. 23**). Cross‑referenced with related terms: **interommatidial angle** and **hyperacuity** (resolving below interommatidial angle).

**Electrical synapse / gap junction**

Direct neural connection allowing ultrafast signal transmission; complements histaminergic synapses in the lamina (**Supplement Section I.1**, **Supplementary Fig. 1**).

**Entropy**
Measure of uncertainty or information content in a signal; the difference between total entropy ($H_{S}$) and noise entropy ($H_{N}$) gives the system’s rate of information transfer (**Supplementary Fig. 8b-e**).

**Flicker-fusion limit**

Temporal frequency at which flicker appears continuous; ~230 Hz in *Musca*, exceeded by LMC high-frequency jumping (~920 Hz) (**Supplementary Fig. 2e**; **Supplementary Fig. 7a**).

**Gaussian white noise (GWN)**

Control stimulus with flat power spectrum; compared against bursty naturalistic stimuli (**Supplementary Fig. 2-3**, **5-6**, **7**).

**High-frequency jumping**

Encoding strategy where synaptic responses shift into higher-frequency carrier bands during rapid input, extending bandwidth and minimising delay (**Supplement Section IV**; **Supplementary Fig. 7d**; main **Figure 2**).

**Histaminergic synapse**

Chemical synapse where photoreceptors release histamine, hyperpolarising LMCs during light increments and, through reduction in histamine release, depolarising them during decrements (**Supplement Section I.3**; **Supplementary Fig. 5a**).

**Hyperacuity**

Visual ability to resolve spatial details finer than the optical sampling grid, achieved here via photoreceptor microsaccadic sampling (**Supplement Section II**; **Supplementary Fig. 13**) and morphodynamic pooling of slightly offset photoreceptor receptive fields (**Supplement Section II**; **Supplementary Fig. 20**).

**Information transfer rate (bits/s)**

Rate at which visual information is conveyed; measured via Shannon formula or triple-extrapolation methods (**Supplementary Fig. 4c-d**, **7c-d**; **Supplementary Fig. 8**).

**Interommatidial angle (**$\Delta\varphi$**)**

Angular spacing between optical axes of neighbouring ommatidia; defines static spatial resolution (~2.9°) of the compound eye (**Supplement Section II**; **Supplementary Fig. 13**).

**Large monopolar cells (LMCs)**

First-order interneurons pooling inputs from six (or seven in male love spots) photoreceptors; perform predictive, high-frequency encoding (**Supplementary Fig. 5-7**; **Supplementary Tables 5-6**).

**Latency distribution**

Spread of delays between photon absorption and quantum bump generation; key parameter in stochastic microvillar models (**Supplement Section IV**).

**Local field potential (LFP)**

Extracellular voltage reflecting summed synaptic activity; used to track motion processing in optic lobes (**Supplementary Fig. 1d**).

**Love spot**

Sexually dimorphic acute zone in male *Musca*’s dorsal-frontal eye, throught to be specialised for mate pursuit (**Supplement Section II**; **Supplementary Fig. 15**).

**Microelectrode intracellular recording**

Sharp-electrode method for measuring membrane potentials in single photoreceptors or LMCs in vivo (**Supplementary Fig. 2a**; **5a**).

**Microvillus (plural: microvilli)**

Actin-rich phototransductive projections in rhabdomeres (~54,000 per photoreceptor) acting as independent photon-sampling units (**Supplement Section II**; **Supplementary Fig. 14g**).

**Microsaccade / photomechanical movement**

Rapid submicron movement of rhabdomeres shifting receptive fields to enhance spatial sampling (**Supplement Section III**; **Supplementary Fig. 21-22**).

**Morphodynamic neural superposition system**

Integrated framework combining optical, mechanical, electrical, and synaptic dynamics to explain predictive visual processing (**Supplement Section IV**; **Supplementary Fig. 21**).

**Noise entropy (**$H_{N}$**)**

Entropy component representing signal variability unrelated to stimulus; subtracted from total entropy to estimate information rate (**Supplementary Fig. 8b-e**).

**Ommatidium (plural: ommatidia)**

Single optical unit of the compound eye containing photoreceptors R1-R8 and support cells (**Supplement Section II**; **Supplementary Fig. 16**).

**Photon shot noise**

Intrinsic variability in photon arrival and detection, imposing a fundamental noise limit on photoreceptor signals (**Supplementary Fig. 2**; **4a**).

**Photoreceptor (R1-R6)**

Outer photoreceptors specialised for motion detection; converge onto LMCs in the lamina (**Supplementary Fig. 2-4**; **Supplementary Tables 1-3**).

**Phototransduction**

Biochemical conversion of photons into electrical signals via quantum bumps in microvilli (**Supplement Section IV**; **Supplementary Fig. 4**).

**Predictive coding**

Neural strategy anticipating future inputs; here, LMCs respond earlier than photoreceptors to moving stimuli (**Supplementary Fig. 1**; main **Figure 2d**).

**Quantum bump (quantal response)**

Elementary voltage response to single-photon absorption; stochastic in timing and amplitude (**Supplementary Fig. 2-3**; **Supplement Section IV models**).

**Quantal sampling theory**

Framework describing vision as the sum of stochastic quantal events across thousands of microvilli (**Supplement Section IV**; **Supplementary Fig. 4**).

**Refractory period**

Interval after a quantum bump during which a microvillus cannot respond; limits maximal sampling rate (4‑parameter model in **Supplement Section IV**).

**Rosette pattern (neural superposition)**

Arrangement where neighbouring ommatidial inputs partly overlap onto a single LMC, enhancing spatiotemporal resolution (**Supplementary Fig. 10 inset**).

**Saccade**

Rapid stereotyped movement of head or body causing transient visual motion; drives bursty stimuli in experiments (**Supplement Section V**).

**Shannon formula (information theory)**

Equation for computing mutual information from $SNR$ spectra; widely used in photoreceptor and LMC analyses (**Supplementary Fig. 4c-d; 7c-d**).

**Signal-to-noise ratio (**$SNR$**)**

Metric comparing signal power to noise power; higher values indicate more reliable encoding (**Supplementary Fig. 4a**; **7a**).

**Synaptic feedback**

Top-down modulation where LMC activity influences photoreceptor output; contributes to dynamic gain control and predictive coding (**Supplement Section I.3**; **Supplementary Fig. 5**).

**Tonic vs. phasic response**

Tonic responses are sustained during constant stimuli; phasic responses are transient, emphasising rapid changes (**Supplementary Fig. 5b; 7b**).

**Triple extrapolation method**

Entropy-based approach for estimating information rates, correcting for finite data length, voltage resolution, and word length (**Supplementary Fig. 8**).

**Ultrafast infrared imaging**

High-speed noninvasive optical method capturing photoreceptor microsaccade dynamics at sub-millisecond resolution in vivo (**Supplement Section III**; **Supplementary Fig. 21**).

**Supplementary Movie Legends**

**Supplementary Movie 1 | LMC response dynamics reveal synaptic high-frequency jumping.**

Intracellular voltage recordings from *Musca* first visual interneurons, the large monopolar cells (LMCs) during controlled light stimulation with sharp microelectrodes. Morphodynamic photoreceptor inputs, evoked by high-contrast bursty stimuli mimicking saccadic light fluctuations, are transformed into virtually noise-free LMC responses with minimal delay. These responses exhibit an effective signalling bandwidth of ~920 Hz (signal-to-noise ratio >1), showing how synaptic high-frequency jumping enables rapid, high-fidelity vision at ~0.5 ms resolution.

**Supplementary Movie 2** **| photomechanical photoreceptor microsaccades drive hyperacute vision.**

High-speed infrared imaging of photoreceptor microsaccades, combined with intracellular recordings to high-resolution moving gratings, reveals how *Musca* photoreceptors resolve fine image detail. Each compound eye (~3,500 ommatidia; cf. 800 in *Drosophila*) is wired in neural superposition. Light changes trigger photomechanical microsaccades - tiny receptor contractions that shift receptive fields and actively sample the scene. These movements, smaller and faster in *Musca*, follow evolutionary scaling laws. Grating tests show that Musca photoreceptors resolve features down to 0.9°, far beyond their 2.9° interommatidial angle, with directional tuning. Thus, insect eyes enhance acuity and speed by encoding space dynamically through time.

**Supplementary Movie 3 | Ultrafast behavioural responses to looming stimuli.**

This video shows how Musca react to looming stimuli. A custom arena with high-speed projection and video recording revealed that tethered flies often initiated movements within ~20 ms, with the fastest at 16.8 ms - far shorter than predicted by models requiring ≥4 synapses before motor activation. Connectomic simulations, using conventional conduction and synaptic transmission values for the shortest pathway (photoreceptors → LMCs → optic lobes → giant fibre → TTM “jump muscle”), predict transmission completing in ~22 ms. Most responses, however, occurred later (40-100 ms), implying additional integration and feedback. These results highlight synaptic high-frequency jumping, predictive coding, and rapid neural synchronisation.

**Supplementary Movie 4 | Simulated conduction of the light-induced antenna reflex pathway in *Musca*.**

This video shows a connectome-based simulation of signal transmission from photoreceptors to antennal motor neurons, using conventional conduction and synaptic values from the literature. Light is first transduced by R1-R6 photoreceptors (~10 ms), followed by transmission across six synapses (~6 ms; cf. **Supplementary Fig. 43**). The signal then propagates along >2,640 µm of axon, with conduction speeds of 0.2-0.5 m/s, adding >5.3-13.2 ms. These stages yield a total predicted reflex delay of ~21-29 ms. By contrast, experiments show the fastest light-pulse-triggered antennal movements within 13 ms.
